# Orthosteric-allosteric dual inhibitors of PfHT1 as selective anti-malarial agents

**DOI:** 10.1101/2020.08.25.260232

**Authors:** Jian Huang, Yafei Yuan, Na Zhao, Debing Pu, Qingxuan Tang, Shuo Zhang, Shuchen Luo, Xikang Yang, Nan Wang, Yu Xiao, Tuan Zhang, Zhuoyi Liu, Tomoyo Sakata-Kato, Xin Jiang, Nobutaka Kato, Nieng Yan, Hang Yin

## Abstract

Artemisinin-resistant malaria parasites have emerged and been spreading, posing a significant public health challenge. Anti-malarial drugs with novel mechanisms of action are therefore urgently needed. In this report, we exploit a “selective starvation” strategy by selectively inhibiting *Plasmodium falciparum* hexose transporter 1 (PfHT1), the sole hexose transporter in *Plasmodium falciparum*, over human glucose transporter 1 (hGLUT1), providing an alternative approach to fight against multidrug-resistant malaria parasites. Comparison of the crystal structures of human GLUT3 and PfHT1 bound to C3361, a PfHT1-specific moderate inhibitor, revealed an inhibitor binding-induced pocket that presented a promising druggable site. We thereby designed small-molecules to simultaneously block the orthosteric and allosteric pockets of PfHT1. Through extensive structure-activity relationship (SAR) studies, the TH-PF series was identified to selectively inhibit PfHT1 over GLUT1 and potent against multiple strains of the blood-stage *P. falciparum*. Our findings shed light on the next-generation chemotherapeutics with a paradigm-shifting structure-based design strategy to simultaneously targeting the orthosteric and allosteric sites of a transporter.

**Significance statement:** Blocking sugar uptake in *P. falciparum* by selectively inhibiting the hexose transporter PfHT1 kills the blood-stage parasites without affecting the host cells, indicating PfHT1 as a promising therapeutic target. Here, we report the development of novel small-molecule inhibitors that are selectively potent to the malaria parasites over human cell lines by simultaneously targeting the orthosteric and the allosteric binding sites of PfHT1. Our findings established the basis for the rational design of next-generation anti-malarial drugs.

## Introduction

*Plasmodium falciparum* is the deadliest species of *Plasmodium* responsible for around 50% of human malaria cases and nearly all malarial death (1). Despite intensive malaria-eradication efforts to control the spread of this disease, malaria prevalence remains alarmingly high, with 228 million cases and a fatality tally of 405,000 in 2018 alone (2). The situation has become even more daunting as resistance to the first-line anti-malarial agents has emerged and is rapidly spreading. For instance, artemisinin resistance, primarily mediated by *P. falciparum* K13 (Pfkelch13) propeller domain (*PF3D7_1343700*) mutations (3, 4), severely compromised the campaign of anti-malarial chemotherapy (5–9). Novel anti-malarial agents overcoming the drug-resistance are therefore urgently needed (10).

The blood-stage malaria parasites depend on a constant supply of glucose as their primary source of energy (11) (**Figure 1A**). Infected red blood cells show an approximately 100-fold increase in glucose consumption compared to uninfected erythrocytes (12). *P. falciparum* hexose transporter PfHT1 (13) (*PF3D7_0204700*) is transcribed from a single-copy gene with no close paralogue (14) and has been genetically validated as essential for the survival of the blood-stage parasite (15). A possible approach to kill the parasite is to “starve it out” by chemical intervention of the parasite hexose transporter (14, 16). The feasibility of this approach would depend on the successful development of selective PfHT1 inhibitors that do not affect the activities of human hexose transporter orthologs (e.g., hGLUT1). Previously, Compound 3361 (**C3361**) (16), a glucose analog, has been described to moderately inhibit PfHT1 and suppress the growth of the blood-stage parasites *in vitro* (17, 18). Nonetheless, modest potency and selectivity of **C3361** had limited its further development.

**Figure 1.**
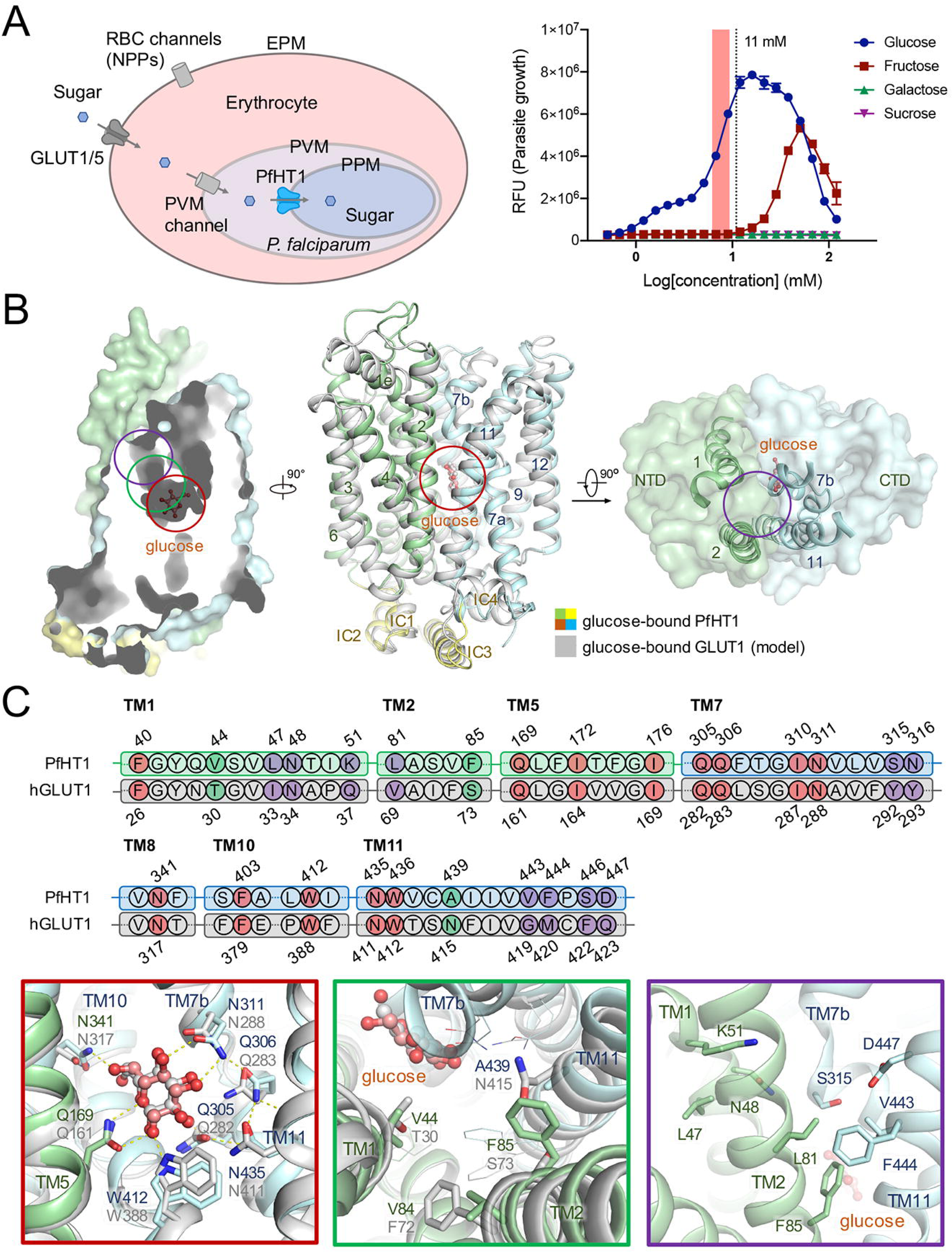
Structural comparison between PfHT1 and hGLUT1 reveals potential druggable site for PfHT1-specific inhibitors. **(A)** *Left:* Schematic representation of the transport processes involved in the uptake of hexoses in *P. falciparum-infected* erythrocytes. GLUT1/5, human glucose transporter 1/5; RBC, red blood cell; NPP, new permeability pathway; EPM, erythrocyte plasma membrane; PVM, parasitophorous vacuole membrane; PPM, parasite plasma membrane. *Right*: Glucose is essential for blood-stage parasite survival. The *P. falciparum* Dd2 grows only in the presence of glucose (peaking at 16 mM) or fructose (peaking at 50 mM). The physiological glucose concentration range in human blood was highlighted. The RPMI media used for parasite culture contains 11 mM of glucose (indicated by the vertical dotted line) in the absence of fructose. **(B)** Superimposition of structures between occluded glucose-PfHT1 complex (domain colored, PDB: 6M20) and a model of the outward-occluded glucose-hGLUT1 complex (gray). The protein structures are shown in cartoon representation. The amino-terminal (N), carboxy-terminal (C), and intracellular helical (ICH) domains of PfHT1 are colored in pale green, pale cyan, and yellow, respectively. **(C)** Sequence alignment of PfHT1 (green and cyan) and hGLUT1 (gray) highlights the portion that engages with glucose. The residues involved in the glucose binding site, the allosteric pocket, and the connecting channel are colored in red, purple, and green, respectively. Residue numbers for PfHT1 and hGLUT1 are shown above and below the alignment, respectively. Close-up views of the glucose binding site, connecting channel, and the extended pocket are presented below, with red, green, and purple box, respectively.

Comparing the structures of PfHT1 (18, 19) and hGLUT1 (20), we identified an additional pocket adjacent to the substrate-binding site. This discovery led to a hypothesis that another pharmacophore tethered to the carbohydrate core might render selective inhibitors for PfHT1. Based on this hypothesis, we designed a new class of small molecules containing a sugar moiety and an allosteric pocket-occupying motif connected by a flexible linker. Among them, **TH-PF01**, **TH-PF02**, and **TH-PF03** have exhibited selective biophysical and anti-plasmodial activities with moderate cytotoxicity. Furthermore, *in silico* computational simulations also confirmed their binding mode, lending further support to the dual inhibitor design. Taken together, our studies validated a new anti-malaria development strategy that simultaneously targets the orthosteric and allosteric sites of PfHT1.

## Results

### Inhibitor binding-induced pocket unique to PfHT1

Recently, we reported the structures of PfHT1 in complex with D-glucose and **C3361** at resolution of 2.6 Å and 3.7 Å, respectively (18). Structural comparison between PfHT1 and hGLUT1 shows that residues around their glucose binding site are nearly identical (**Figure 1B/1C/S1**). Interestingly, we discovered an additional pocket adjacent to the substrate-binding site, linked by a narrow channel which is highly hydrophobic in PfHT1 but more hydrophilic in hGLUT1 (**Figure 1B/1C**). Based on this observation, we hypothesized that extended carbohydrate derivatives might render selective inhibitors for PfHT1 by occupying the allosteric site.

To confirm the allosteric pocket as a potential selectivity-increasing site for PfHT1 inhibition, we set up to resolve structures of human glucose transporters in the presence of **C3361**. Eventually, the crystal structure of hGLUT3 in complex with **C3361** was resolved to 2.30 Å (**Figure 2/S2, Table S1**). The overall structure of hGLUT3 bound to **C3361** adopts a similar conformation with glucose-bound form (21) (**Figure 2A**). The sugar moiety of **C3361** is coordinated in an identical fashion to that of D-glucose by conserved residues in hGLUT3.

**Figure 2.**
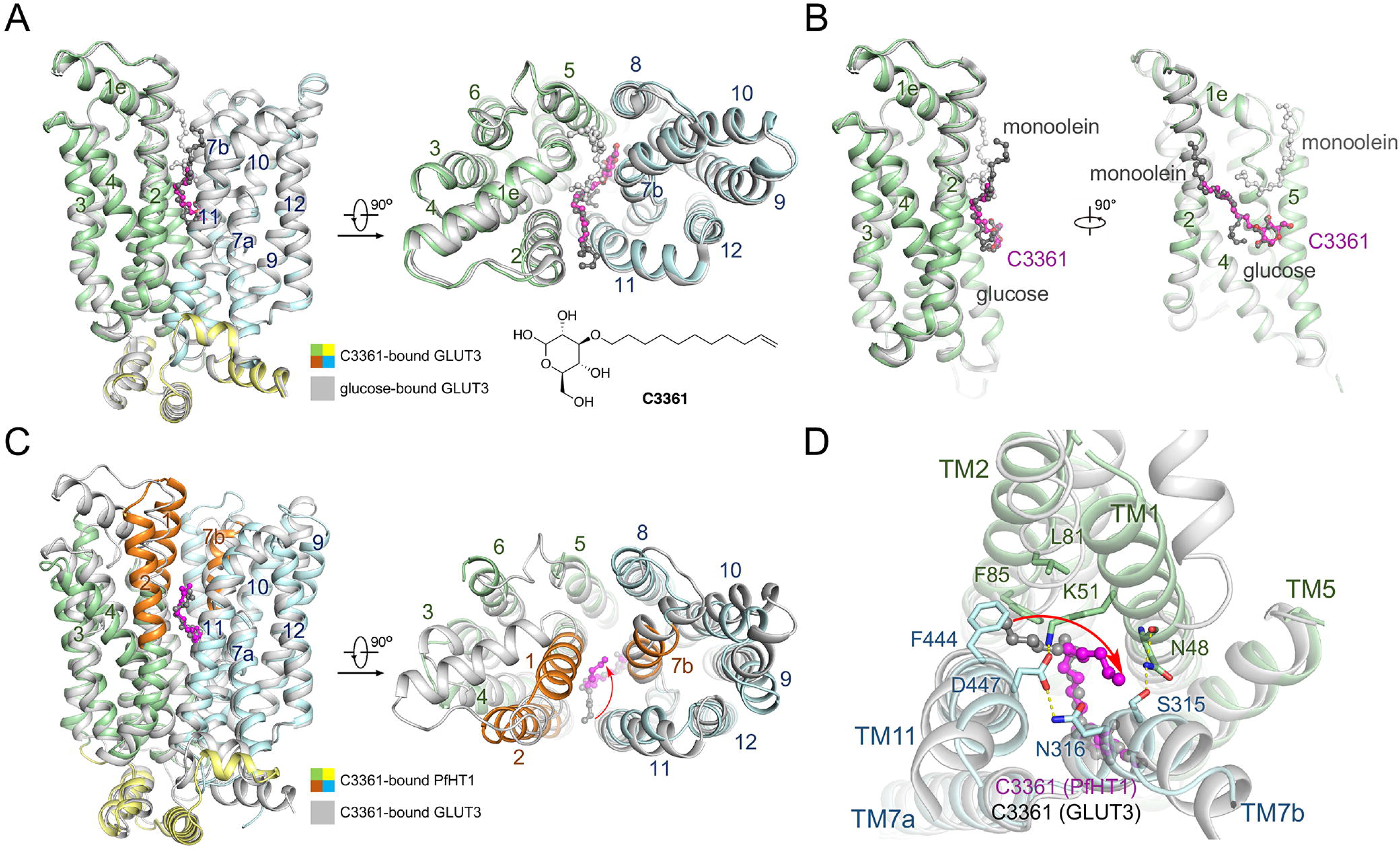
Crystal structure of GLUT3 bound to C3361 in an outward-occluded conformation. **(A)** Superimposed structures of glucose-GLUT3 complex (gray, PDB: 4ZW9) and **C3361**-GLUT3 complex (domains colored) presented in both side and intracellular views of overall structures. The protein structures are shown in cartoon representation, and the ligands are shown in ball-and-stick representation. The N, C, and ICH domains of **C3361**-bound GLUT3 are colored in pale green, pale cyan, and yellow, respectively. **(B)** The sugar moiety of **C3361** is coordinated by the same set of polar residues as for D-glucose, and the tail of the **C3361** occupied the position where a monoolein molecule located in the glucose-bound GLUT3 structure. **(C)** Superimposed structures of **C3361**-GLUT3 complex (gray) and **C3361**-PfHT1 complex (domains colored, PDB: 6M2L) presented in both side and intracellular views of overall structures. **(D) C3361** demonstrated distinct binding modes when complexed with GLUT3 or PfHT1, in particular, its tails pointed toward different directions.

The conformations of the aliphatic tail of **C3361** are different in hGLUT3 and in PfHT1. In hGLUT3, the tail of **C3361** occupies the pocket which accommodates monoolein, a lipid used in lipidic cubic phase crystallization, in the GLUT3-glucose complex (**Figure 2B**). The tail of **C3361** points to the interface between TM2 and TM11 (**Figure 2C**). In the occluded structure of **C3361**-bound PfHT1 (18), the tail projects into the central cavity (**Figure 2D**). These structural differences suggest that PfHT1 possesses unique intra-domain flexibility that may be exploited for designing selective inhibitors that target the allosteric site of PfHT1 without inhibiting GLUTs.

### Rational design of dual-pocket inhibitors of PfHT1

Inspired by the structural insights, we designed a series of substituted carbohydrate derivatives that consist of a sugar moiety, a tail group occupying the allosteric pocket, and an aliphatic linker (**Table 1/S2**). All these compounds were successfully prepared using a concise synthetic route (**Supplementary Information**) and tested by a previously established, proteoliposome-based counterflow assay against PfHT1 (**Figure S3**). In parallel, the cytotoxicity and blood-stage parasite growth inhibitory activities were also evaluated. (**Table 1/S2**).

**Table 1.**
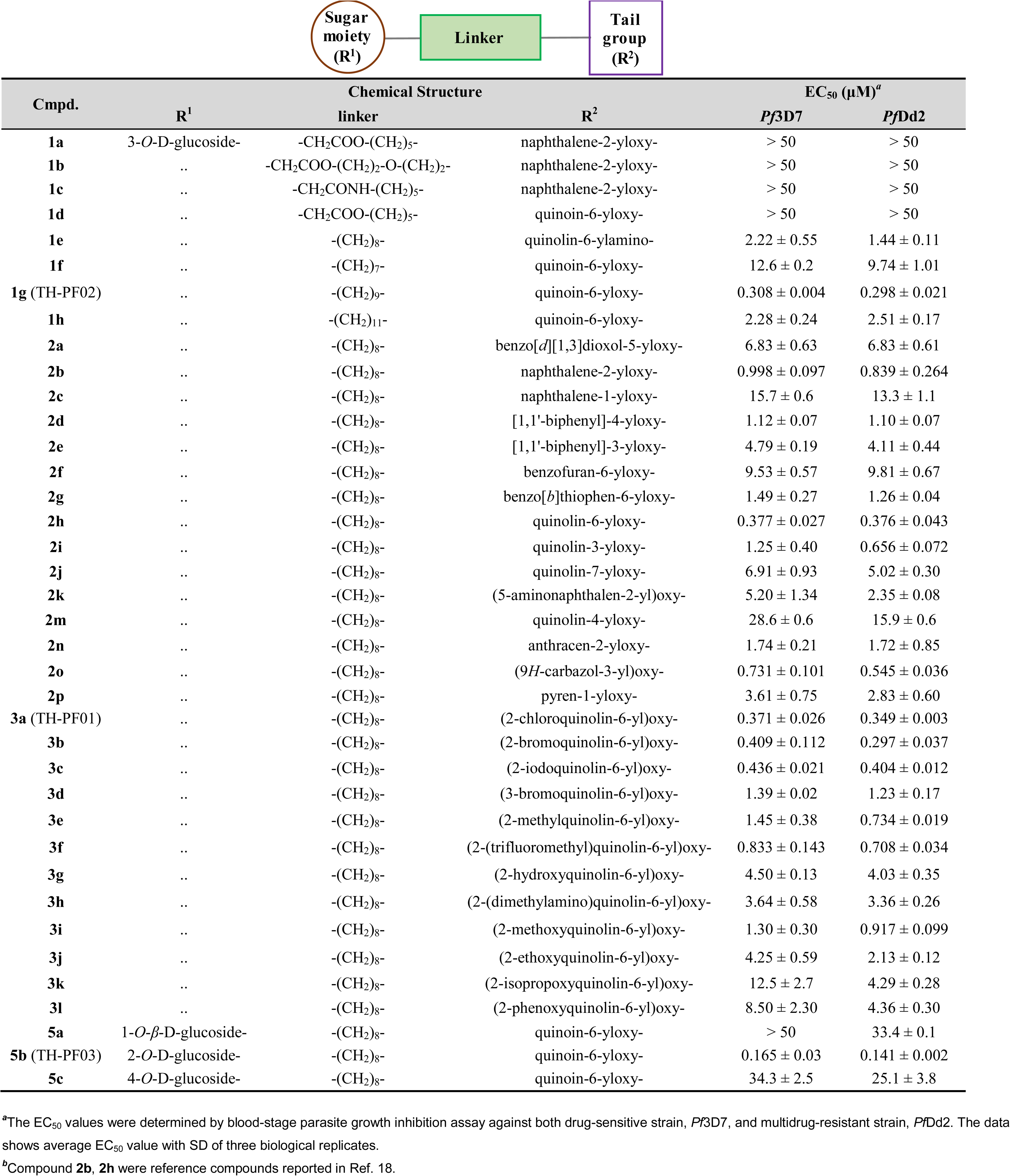
Structure-activity relationship (SAR) studies of the PfHT1 inhibitors.

Firstly, we examined different tail groups with various sizes to fit the allosteric pocket. Among them, the bicyclic rings, such as the naphthyl (**2b**) (18), quinolinyl (**2h**, **2i**) (18), and biphenyl (**2d**) rings, enhanced the potency, whereas smaller (**2a**, **2f**, **2g**) or larger (**2n**, **2p**) rings lowered the potency. Different positions of the linkers have been explored, suggesting that a particular orientation is desired presumably to fit well with the connecting channel (**2b** vs. **2c**, **2d** vs. **2e**, **2h** vs. **2m**). The introduction of nitrogen into the naphthyl ring significantly enhanced the inhibitory potency (**2b** vs. **2h)** suggesting potential polar interactions within the allosteric pocket. Such polar interaction was further supported by the fact that the inhibitory effect is sensitive to the positions of the nitrogen atom (**2i**, **2j**, **2k**). We also optimized the skeleton structure of **2h** to further explore the pharmacophore. We found that relatively small size of the substituent group was desired; nonetheless, neither electron-withdrawing (**3a**, **3b**, **3c**, **3f**) nor electron-donating (**3e**, **3i**) groups showed little impact on their potency.

Next, a C_9_ polymethylene linker has been shown to be optimal (**1g**), both shorter (**1f**) and longer chains (**1h**) decreased the activity significantly. The linkers with either ester (**1a**, **1b**, **1d**) or amido (**1c**) functionality lost their inhibitory activity almost completely, which might provide information on the structural constrains of the connecting channel. Installment of the tail group via amidogen ether bond (**1e**) rather than oxygen ether bond (**2h**) also decreased the potency. These structure-activity relationship (SAR) results are in good agreement with the hydrophobic channel that linking the orthosteric and allosteric pockets of PfHT1 (**Figure 1C**).

Finally, we explored the different substitution site on the carbohydrate core. Various derivatives of glucose, in which the substituents were introduced (**5a**, **5b**, **5c**), were successfully prepared. The *O*-1 or *O*-4 substituted derivatives showed almost no activity (**5a**, **5c**). By contrast, the *O*-2 and *O*-3 derivatives showed improved potency, demonstrating the necessity of the appropriate orientation (**5b**, **2h**).

From the extensive SAR studies, three glucose derivatives, designated **3a** (**TH-PF01**), **1g** (**TH-PF02**), and **5b** (**TH-PF03**), stood out as the lead compounds (**Figure 3A/3B/3C**). The half-maximal inhibitory concentration (IC_50_) values of the glucose transport activity for **TH-PF01**, **TH-PF02**, and **TH-PF03** were determined as 0.615 ± 0.046 μM, 0.329 ± 0.028 μM, and 1.22 ± 0.09 μM for PfHT1, and 111 ± 17 μM, 92.3 ± 7.3 μM, and 97.3 ± 5.9 μM for hGLUT1, respectively. In a similar trend, the half-maximal effective concentration (EC_50_) values of **TH-PF01**, **TH-PF02**, and **TH-PF03** for the parasite growth inhibition assay were shown to be 0.371 ± 0.026 μM, 0.308 ± 0.004 μM, and 0.165 ± 0.003 μM against the 3D7 strain, and 0.349 ± 0.003 μM, 0.298 ± 0.021 μM, and 0.141 ± 0.002 μM against the Dd2 strain of *P. falciparum*. Furthermore, these lead compounds showed reasonable therapeutic window indicated by their high CC_50_ / EC_50_ ratio values ranging 36.1 to 107.2 (**Figure 3C**). It was worth noting that all these rationally designed compounds showed equipotency to both the wild type strain (*Pf*3D7) and the multi-drug-resistant strain (*Pf*Dd2), starkly contrast to quinine that only showed activity against *Pf*3D7 (**Figure 3A**).

**Figure 3.**
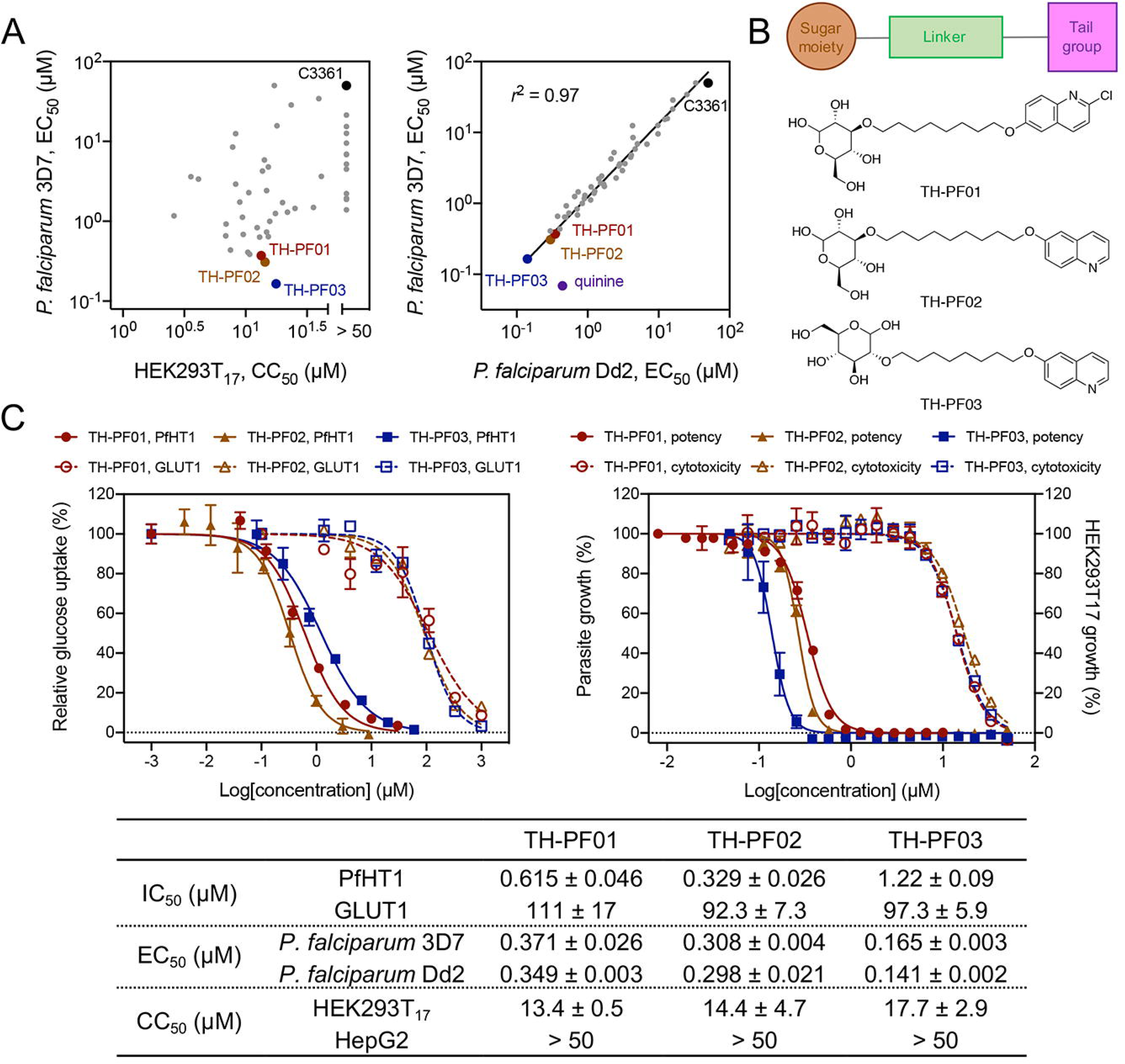
Rational design of selective inhibitors targeting PfHT1. **(A)** Potency against *P. falciparum* (3D7 and Dd2 strains) and cytotoxicity in HEK293T_17_. PfHT1 inhibitors showed equal potency to quinine-resistant strain, Dd2, as well as 3D7. All EC_50_ and CC50 values are average of 2 or 3 biological replicates. See the entire data set in Table S2. **C3361** was shown as reference compound reported in Ref. 18. **(B)** The generic chemical structures of **TH-PF01**, **TH-PF02**, and **TH-PF03** containing a sugar moiety, a substituted heteroaromatic tail, and a flexible linker. **(C) TH-PF01**, **TH-PF02**, and **TH-PF03** demonstrated good parasite growth inhibitory activities, selectivity, and cytotoxicity. The IC_50_ values were determined by proteoliposome-based counterflow assay. The blood-stage *P. falciparum* (3D7 and Dd2) and mammalian cells (HEK293T_7_ and HepG2) were incubated with the compounds for 72 hours, and the cell growth was quantified with SYBR green I and Cell Titer Glo correspondingly. The data is a representative of three bio-replicates and shown as average ± SD of three technical replicates.

### TH-PF series competitively inhibit PfHT1

To further elucidate the binding mode of **TH-PF** inhibitors against PfHT1, we next employed *in silico* molecular docking simulation. The results confirmed that the sugar moiety of **TH-PF01** could fit into the orthosteric glucose-binding pocket of PfHT1. On the other hand, the quinoline fragment was suggested to occupy the allosteric site, forming a hydrogen bond network between **TH-PF01**, Lys51, and Asp447 of the protein (**Figure 4A**). To validate these specific interactions observed, we performed an *in silico* alanine scan of residues within the **TH-PF01**-binding site. Reduced association energy predicted with mutants harboring K51A, Q169A, Q305A, and N341A agreed with the rational design that emphasizes the importance of the polar interactions (**Figure 4B**). To further test this hypothesis, several PfHT1 mutants that contained a single point mutation, Q169A, Q305A, N341A, K51A, F85S, F85Y, V44T, or a double mutation, K51A/D447A, were experimentally prepared and measured using the counterflow assay with 1 μM **TH-PF01** or isovolumetric DMSO as control. Compared with the wild type, **TH-PF01** showed reduced potency to all these mutants, further confirming the computational simulation results. More specifically, Q169A, Q305A, and N341A demonstrated the interaction between the PfHT1 and the sugar moiety of **TH-PF01**. And both K51A and K51A/D447A showed the additional hydrogen network formed between PfHT1 and the quinoline tail group. Finally, mutations of the residues within the connecting channel to their corresponding ones in hGLUT1(F85S and V44T) or merely enhancing the polarity (F85Y) decreased the inhibitory potency, indicating the hydrophobicity of the lead compounds was indeed critical to their selectivity (**Figure 4C**). Taken together, molecular simulation combined with the experimental mutagenesis strongly supported that **TH-PF** compounds recognize both the orthosteric and allosteric sites connected via the narrow channel as predicted (**Figure 4D**).

**Figure 4.**
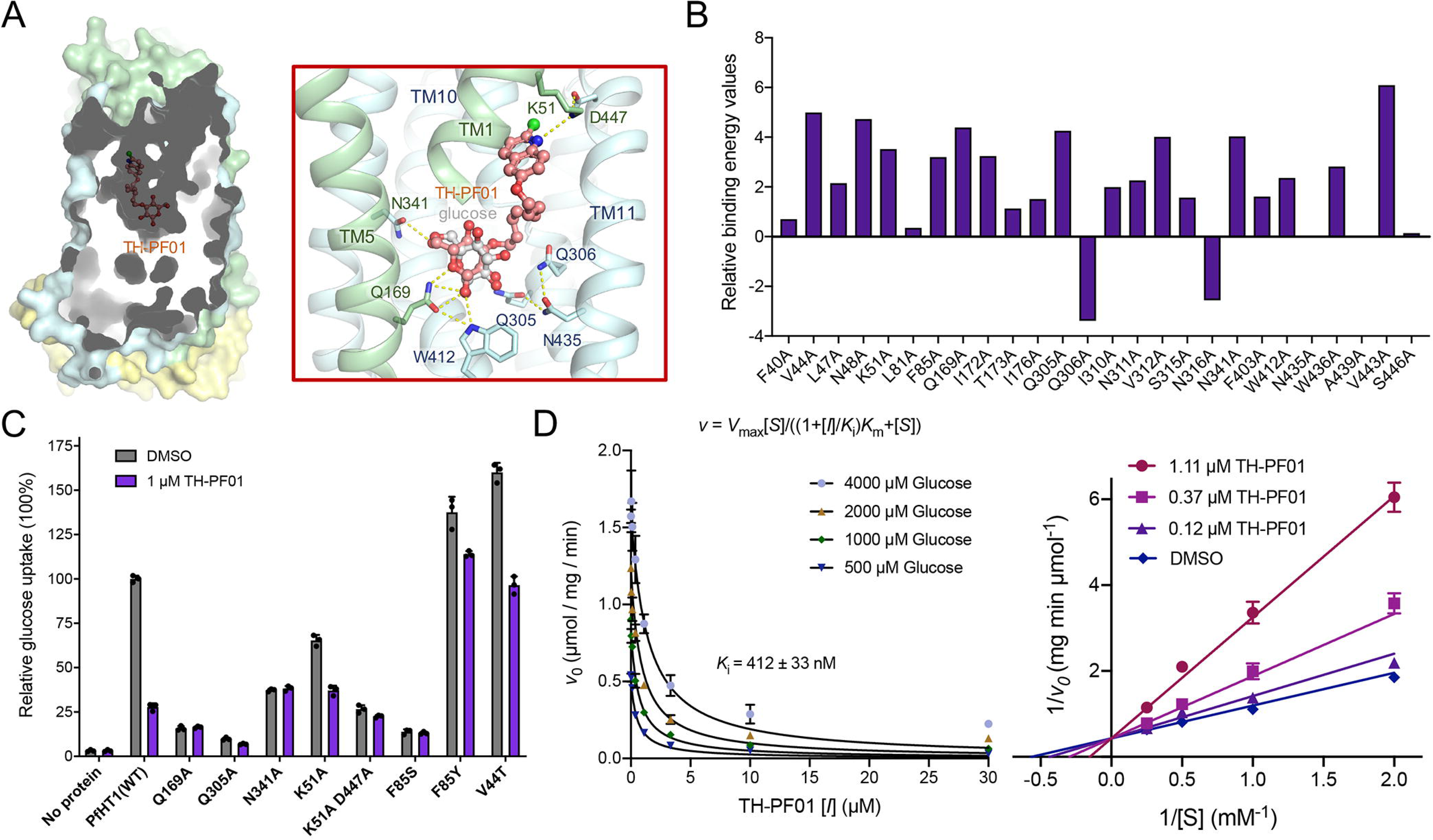
Biophysical characterizations of TH-PF01 binding to PfHT1. **(A)** The identical semi-transparent cut-open view of the protein surface is shown for PfHT1 with **TH-PF01** docked into the protein. Hydrogen bonds are shown as yellow, dashed lines. The N, C, and ICH domains of PfHT1 were colored in pale green, pale cyan, and yellow, respectively. **(B)** *In silico* alanine scanning results. Positive values indicate that the alanine substitution interacts less favorably with **TH-PF01** than the native residue. **(C)** Key residues involved in **TH-PF01** recognition were tested by protein mutagenesis. **(D)** *Left:* The curves represent the best fit of data to the competitive inhibition equation, *v* = *V*_max_[*S*]/((1+[*I*]*K*_i_)*K*_m_+[*S*]), where *K*_i_ is the apparent inhibition constant of TH-PF01. *Right*: Lineweaver-Burk plot of experimental kinetic data for inhibition of PfHT1 by **TH-PF01**, confirming a competitive inhibition mode. All experiments have been repeated three times and the data shown as mean ± SD.

### TH-PF series kill the blood-stage *P. falciparum* via PfHT1 inhibition

We further examined whether the disruption of the PfHT1 activity can explain the growth inhibition of the **TH-PF** compounds. We reasoned that if the primary anti-plasmodial mechanism of the **TH-PF** compounds was via inhibition of PfHT1, then IC_50_ values for the glucose transport activity of PfHT1 should correlate with EC_50_ values obtained in parasite growth inhibition assays. Indeed, a reasonable correlation between the two parameters was observed using 12 analogs from the **TH-PF** series covering a wide range of activities (**Figure 5A**). We also confirmed that the EC_50_ values of **TH-PF01** and **TH-PF03** improved in the culture media with lower glucose concentration while other anti-malarial drugs, quinine, mefloquine, and dihydroartemisinin, showed no effects by glucose concentration (**Figure 5B**). These results indicate that **TH-PF01** and **TH-PF03** are competing with glucose for the same substrate-binding site of PfHT1, confirming the on-target effects of the **TH-PF** series.

**Figure 5.**
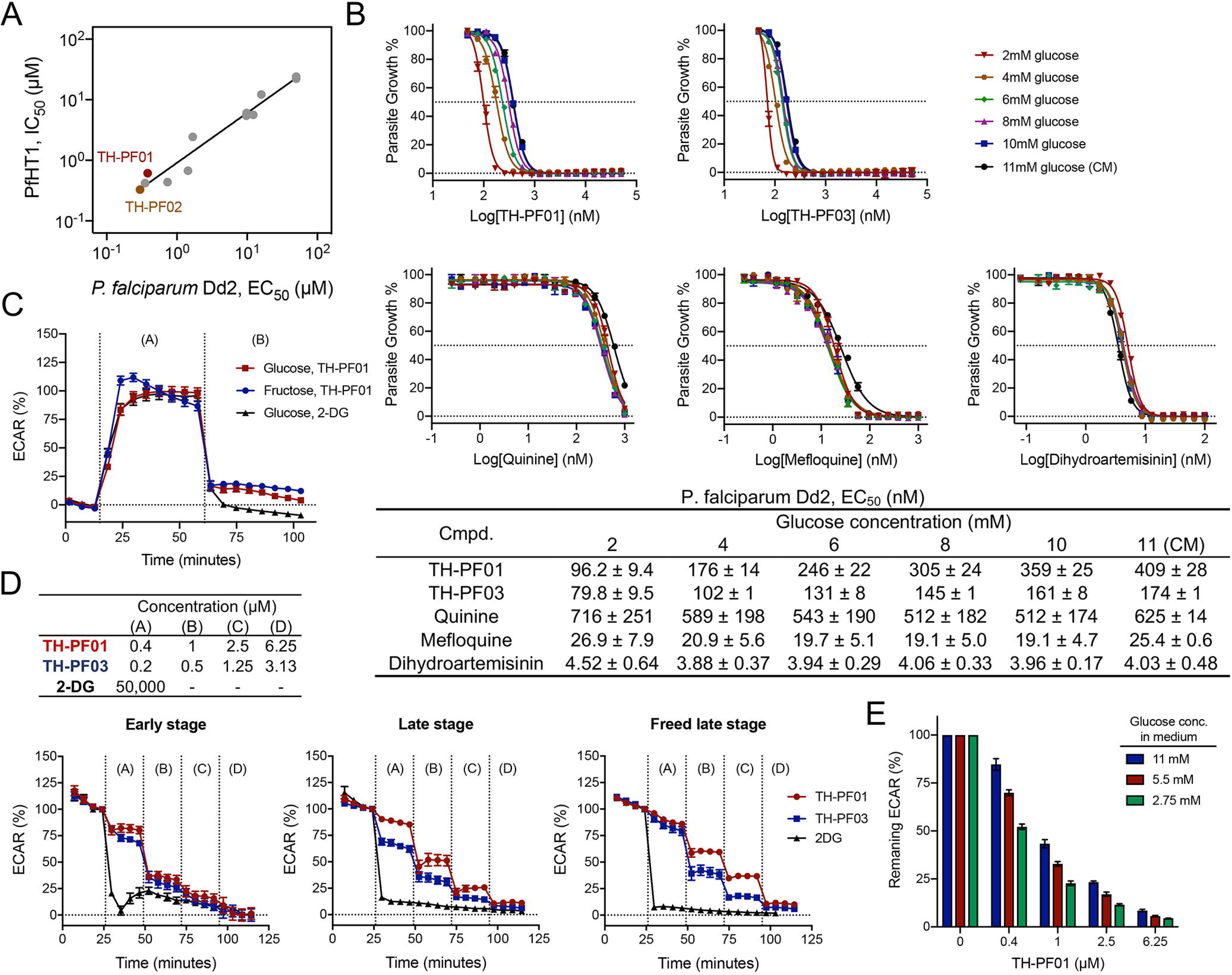
TH-PF01 derivatives selectively target PfHT1. **(A)** PfHT1 inhibitors showed a reasonable correlation between blood-stage parasite growth inhibition (EC_50_) and biochemical inhibition (IC_50_) of PfHT1 glucose transport activity. **(B)** Glucose concentration in culture media offsets EC_50_ values of **TH-PF01** and **TH-PF03** in a dose-response manner, but not common antimalarial drugs, quinine, mefloquine, and dihydroartemisinin (DHA). The blood-stage *P. falciparum* Dd2 was exposed to the test compounds in assay media containing glucose at different concentrations for 48 hours starting at the ring stage, and the parasite growth was determined by SYBR green I. The data represent four (**TH-PF01**) or two (**TH-PF03**, quinine, mefloquine, and DHA) independent experiments and shown as average ± SD of two technical replicates. **(C)** The inhibition of glycolytic activity by **TH-PF01** was observed by Seahorse extracellular flux analyzer. Dd2 schizont stage parasites in RBCs were seeded in medium without glucose and exposed to glucose (11 mM as final concentration) or fructose (40 mM as final concentration) at 15 min (the first vertical dotted line), resulting from robust increases of ECAR. The addition of **TH-PF01** (20 μM as final concentration) or 2-DG (50 mM as the final concentration) at 61 min (the second vertical line) lowered the ECAR, indicating glycolytic activity was inhibited. Data were normalized with ECAR values before and after the glucose additions as 0% and 100%. All data were average values pooled from two independent experiments with three technical replicates. Error bars represent SEM. **(D)** Extracellular flux analysis showed that **TH-PF01** and **TH-PF03** inhibit glycolytic activity in a dose-dependent manner in the early (Rings) and late stages (Trophozoites/Schizonts) of the parasites in RBCs as well as freed late stages from RBCs. The *Pf*Dd2 parasites were seeded in assay medium containing glucose (11 mM), and **TH-PF01** or **TH-PF03** was sequentially added to four times with the final concentrations of 0.4, 1, 2.5, and 6.25 μM or 0.2, 0.5, 1.25 and 3.13 μM. Glycolysis inhibitor, 2-DG, was added at 50 mM once. ECAR values were normalized with the values before the first compound addition as 100% and the values of background as 0%. All data were average values pooled from two independent experiments with two or three technical replicates. Error bars represent SEM. **(E)** The glucose concentration in the assay media and the potency of **TH-PF01** against the glycolysis activity show a negative correlation. *Pf*Dd2 schizont stage parasites in RBCs were seeded in assay medium supplemented with glucose at 11, 5.5 or 2.75 mM, and **TH-PF01** was sequentially added four times with the final concentrations of 0.4, 1, 2.5 and 6.25 μM. ECAR values after compound addition were normalized with the value before the first compound addition as 100% and the value of background as 0%. All data were average values pooled from two independent experiments with two technical replicates. Error bars represent SEM.

We further assessed whether the **TH-PF** series disrupts the glycolysis activity of the blood-stage *P. falciparum*. Seahorse extracellular flux analyzer has been used to simultaneously monitor glycolysis and mitochondrial respiration in live cells through extracellular acidification rate (ECAR) and oxygen consumption rate (OCR), respectively (22). Using magnetically-purified infected red blood cells, we observed robust initiation of glycolysis in late-stage parasites after the addition of glucose or fructose (**Figure 5C**). This increased ECAR was abolished by the addition of **TH-PF01** and 2-deoxyglucose, a glycolysis inhibitor, clearly demonstrating **TH-PF01**’s inhibitory activity of glycolysis. It should be noted that 2-deoxyglucose required a higher concentration (50 mM) than **TH-PF01** (20 μM) to diminish the increased ECAR. Furthermore, we confirmed that **TH-PF01** and **TH-PF03** reduced the ECAR at EC_50_ values and that the ECAR was decreased in a dose-dependent manner. This glycolysis inhibition was also observed with early-stage parasites in red blood cells and late-stage parasites extracted from red blood cells (**Figure 5D**). Lastly, we measured ECAR reduction by **TH-PF01** in media containing three different glucose concentrations (**Figure 5E**). Similar to the EC_50_ shift (**Figure 5B**), ECAR reduction was negatively correlated with glucose concentration. All of these findings provide strong evidence that the **TH-PF** compounds disrupt the glycolytic activity of the blood-stage parasites.

### Evaluation of glucose dependency of the blood-stage parasites

We further examined how the glucose dependency of *P. falciparum;* thus, the potency of the **TH-PF** series varies during the blood-stage of the parasite life-cycle. The blood-stage *P. falciparum* has a two-day life cycle comprising merozoite invasion, proliferation from ring stage to trophozoite and then multicellular schizont, and egress from red blood cells (23). We investigated the sub-stage specific activity of the **TH-PF** series using lactate dehydrogenase-based assay (24). We first treated the ring-stage parasites (*P. falciparum* 3D7) with **TH-PF01** or dihydroartemisinin (control) for 24, 36, 48, and 72 hours, and found that the EC_50_ values were very similar to the previously mentioned 72-hours growth inhibition assay [(i) in **Figure 6A/B**]. Next, we treated the early-ring, late-ring, trophozoite, or schizont stage with **TH-PF01** for 12 hours. Then, the parasites were washed with growth media and further incubated for an additional 36 hours without the compound. The ring-stage parasites were less sensitive to **TH-PF01** than the late-stage parasites (trophozoite and schizont) [(ii) in **Figure 6A/B**]. On the contrary, dihydroartemisinin was found to be less potent against the late stages. Lastly, we treated parasites with the compounds for 24, 36, 48, and 72 hours from the early-ring stage and then incubated them without the compounds for an additional 36 hours [(iii) in **Figure 6A/B**]. The obtained EC_50_ values also showed the ring-stage parasites were less sensitive to the PfHT1 inhibitor than late-stage parasites (24 hours treatment), but longer than 36 hours of treatment showed similar potency as 72 hours assay. Light microscopic observations of the compound treated parasites suggested that exposer to **TH-PF01** induced the ring-stage parasites to arrest its development but the arrested parasites restart growth after removal of **TH-PF01** (**Figure 6C/S4**).

**Figure 6.**
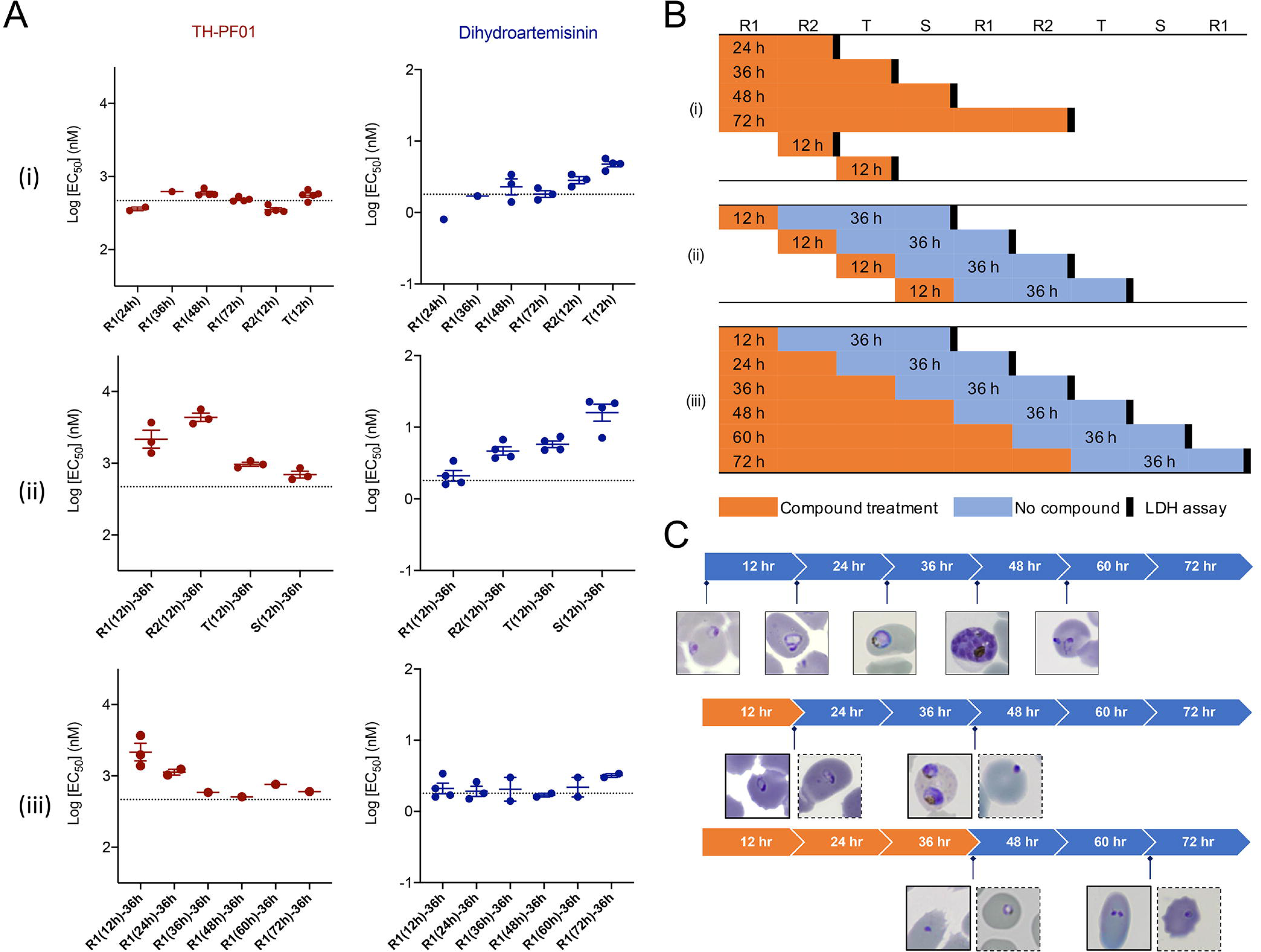
TH-PF01 suppressed parasite growth at different blood substages. **(A) TH-PF01** appeared as equipotent to all substages when the survival was assessed immediately after compound treatment; however, it required longer incubation time against ring-stage than late-stage parasites to show the same potency when incubated for additional 36 hours after washed. EC_50_s of **TH-PF01** and DHA against substages were determined by LDH assay in the time course experiments depicted in Figure 6B. The horizontal dashed line indicates EC_50_ value determined by a 72-hour SYBR assay. The data shows an average EC_50_ with SD of one to four bio-replicates. **(B)** A schematic representation of the substage assay. Tightly synchronized parasites (*Pf*3D7) were exposed to **TH-PF01** or DHA for various periods at the substages indicated. **(C)** Representative images of compound treated parasites. Solid outline: **TH-PF01** treated parasites; dotted outline: DHA treated parasites. See also Figure S5.

## Discussion

With the continuous emergence and spread of drug resistance, current anti-malaria chemotherapies are facing serous limitations (3). Innovative drug discovery strategies, novel targets, and therapeutic agents for malaria treatment are an urgent need. Given that proliferation of the malaria parasites depends on D-glucose and D-fructose, we have conceptualized a “selective starvation” strategy. Decreasing the uptake of D-glucose and D-fructose via PfHT1, the sole hexose importer in *P. falciparum*, could be a potential venue to kill drug-resistant malaria parasites. A comparison of the crystal structures of PfHT1 (18) and hGLUT1 led to the discovery of an allosteric site that further prompted us to design and develop selective PfHT1 inhibitors over its human orthologs. The **TH-PF** series demonstrated a strong correlation between blood-stage growth inhibition and the biophysical inhibition of glucose transporting activity. Moreover, the inhibitors quickly shut down the glycolysis of the blood-stage parasites and demonstrated even higher potency when glucose concentration in the growth media was lowered, confirming that PfHT1 is indeed the molecular target of the **TH-PF** series. Approximately 15-20% malaria patients suffer from life-threatening hypoglycemia and other complications, including irreversible brain damage and neurological sequelae (25). Thus, inhibiting PfHT1 cannot only starve and kill the blood-stage parasites but also might quickly relieve hypoglycemia symptoms. Lastly, these PfHT1 inhibitors may validate a novel approach for structure-based drug design. To the best of our knowledge, this is the first example of dual inhibition of the orthosteric and allosteric sites for a transmembrane protein. Our findings served as proof of the concept that PfHT1 is a druggable target for next-generation anti-malarials, laying a foundation for future therapeutic development.

## Materials and Methods

### Protein expression and purification

Protein over-expression and purification methods of hGLUT1 (N45T) and hGLUT3 (N43T) were described previously (20, 21). Recombinant PfHT1 and its mutants were expressed and purified with similar protocols. In brief, the protein with an N-terminal His_10_ tag was expressed in Sf9 cells following the Bac-to-Bac baculovirus system (Invitrogen). Cells were collected and solubilized to extract protein in the buffer containing 25 mM MES pH 6.0, 150 mM NaCl, 2% (w/v) *n*-dodecyl-β-D-maltopyranoside (DDM, Anatrace) and protease inhibitors cocktail (aprotinin at 5 μg/ml, pepstatin at 1 μg/ml, and leupeptin at 5 μg/ml; Amresco) at 4 °C for 2 h. After high-speed centrifugation (18,700 *g*), the supernatant was incubated with Ni-NTA resin (Qiagen) at 4 °C for 30 min. Then the resin was washed with the buffer, which contains 25 mM MES pH 6.0, 150 mM NaCl, 30 mM imidazole, and 0.02% (w/v) DDM. The target protein was eluted with wash buffer plus 270 mM imidazole. Eluted protein was then applied to size-exclusion chromatography (Superdex 200 Increase 10/300 GL column, GE Healthcare) with the buffer containing 25 mM MES pH 6.0, 150 mM NaCl, and 0.02% (w/v) DDM.

### Preparation of liposomes and proteoliposomes

Liposomes and proteoliposomes were prepared as previously described (21, 26, 27). *Escherichia coli* polar lipid extract (Avanti) were dissolved to 20 mg/ml in KPM 6.5 buffer (50 mM potassium phosphate pH 6.5, 2 mM MgSO_4_) plus 50 mM D-glucose. Liposomes were frozen and thawed in liquid nitrogen for 10 rounds and extruded through 0.4 μm polycarbonate membrane (Millipore). 200 μg/ml purified proteins were added into liposomes after incubation with 1% *n*-octyl-β-D-glucopyranoside (β-OG, Anatrace) at 4 °C for 30 min. Add 400 mg/ml Bio-Beads SM2 (Bio-Rad) into proteoliposomes to remove detergents one hour later. After another extrusion of proteoliposomes, the excessive glucose was removed by ultracentrifugation at 100,000 g for 1 h. Then the proteoliposomes were resuspended to 100 mg/ml in ice-cold KPM 6.5 buffer.

### Counterflow assay

To determine the IC_50_ values of the inhibitors, reconstituted proteoliposomes were pre-incubated with varying concentrations of inhibitors for 30 min on ice. Then 2 μl proteoliposomes were added to 98 μl KPM buffer, which contains inhibitors and 1 μCi D-[2-^3^H]-glucose (23.4 Ci/mmol, PerkinElmer). After 30 s’ reaction, the solution flowed through 0.22 μm membrane filter (Millipore) in vacuum filtration rapidly. 0.5 ml Optiphase HISAFE 3 (PerkinElmer) was incubated with the filter overnight before liquid scintillation counting with MicroBeta JET (PerkinElmer). IC_50_ values were determined by fitting the data to a nonlinear regression curve in Prism Software version 8 (GraphPad).

To determine the inhibition mechanism of inhibitors, the initial velocities were measured at 15 s. The non-radiolabeled glucose concentration in the external reaction solution was from 0.5 mM to 4 mM. The data were fitted to the competitive inhibition equation, *v* = *V*_max_[*S*]/((1+[*I*]/*K*_i_) *K*_m_+[*S*]), in Prism Software version 8 (GraphPad). All experiments were performed three times. Data were expressed as mean ± SD.

### Crystallization

For lipidic cubic phase crystallization of the GLUT3-**C3361** complex, the GLUT3 protein was purified in 0.06% (w/v) 6-cyclohexyl-1-hexyl-β-D-maltoside (Cymal-6, Anatrace) and concentrated to ~ 40 mg/ml. After pre-incubation with 50 mM **C3361** at 4 °C for 1 hour, the protein was mixed with monoolein (Sigma) in 2:3 protein to lipid ratio (w/w)(28). Then 75 nl mixture was loaded into the glass sandwich plate well with 1 μl precipitant solution by an LCP crystallization robot (Gryphon, Art Robbins Instruments). The best diffracting crystals were grown in the solution with 0.1 M NH_4_Cl, 0.1 M HEPES pH 7.0, and 40% PEG400 at 20 °C. Crystals appeared after 1 day and grew to full size within one week.

### Data collection and structural determination

The X-ray diffraction data were collected at the BL32XU beamline of SPring-8, Japan. Due to the small size of LCP crystals, a 10 × 15 μm micro-focus beam with 1.0 Å wavelength was applied for data collection. Wedges of 10° were collected for every single crystal with a 0.1° oscillation angle through the EIGER X 9M detector. Hundreds of data sets were screened and automatically collected by ZOO (29), followed by first-round automatic data processing through KAMO (30). Data sets with good diffraction and low R-merge factor were manually picked out and merged through XDS (31). Further processing was carried out using the CCP4 suite (32). The phase was solved by molecular replacement with a search model of GLUT3 (PDB code: 4ZW9) through PHASER (33). The structural model was adjusted through COOT (34) and refined by PHENIX (35). Supplementary information, Table S1 summarizes the statistics for data collection and structure refinement.

### Homology modeling and molecular docking

The outward-occluded models of hGLUT1 were built and refined based on the crystal structure of glucose-bound hGLUT3 (PDB: 4ZW9) as a template within Modeller-9.19 (36), and the best model was chosen by PROCHECK (37). Molecules (glucose and **TH-PF01**) were drawn in 2D sketcher in Schrödinger suite 2018-1 (38), and 3D structures were processed by default setting using the Ligprep program (39). The protein structures were processed by default setting using the Protein Preparation Wizard. Molecules were docked against PfHT1 (PDB: 6M2L) or hGLUT1 model using the extra-precision docking (Glide XP) method within the Glide program.

### *In silico* alanine scanning

The binding pocket was defined by identifying residues in direct contact with **TH-PF01**, including F40, V44, L47, N48, K51, L81, F85, Q169, I172, T173, I176, Q305, Q306, I310, N311, V312, S315, N316, N341, F403, W412, N435, W436, A439, V443, S446. To validate the role of these residues in the inhibitor binding, each pocket residue was mutated to alanine *in silico* using PyRosetta (40, 41). Then, the relative binding free energy change (ΔΔG) of each mutant over wild type was calculated using the Prime-GBSA method, keeping the ligand and other residues fixed.

### Chemical Synthesis and stock preparation

All the compounds were synthesized using a concise synthetic route. Please refer to the Supplementary Methods for details. All final products were validated with purity >95% purity by high-performance liquid chromatography (HPLC), NMR, and high-resolution mass spectrometric (HMS) analyses. The final products were reconstituted in 100% DMSO at 50 mM prior to dilutions in aqueous buffers or cell culture medium for the following assays.

### *P. falciparum in vitro* culture

Strains of *P. falciparum* (Dd2 and 3D7 strains) were gifts from the Institut Pasteur of Shanghai, Chinese Academy of Science. The erythrocytic stages of parasites were cultured following standard methods in RPMI 1640 medium (Thermo) supplemented with human blood, 5.94 g/L HEPES (Thermo Fisher Scientific), 0.5% Albumax II (Thermo Fisher Scientific), 50 mg/L hypoxanthine (Sigma-Aldrich), 2.1 g/L sodium bicarbonate (Sigma-Aldrich), and 25 mg/L gentamicin (Sangon Biotech). Cultures were grown at 37 °C in an atmosphere of 5% O_2_, 5% CO_2_, and 90% N_2_ and regularly synchronized by 5% sorbitol (Sigma-Aldrich) treatment.

### *In vitro* Drug Sensitivity and EC_50_ Determination

Drug susceptibility was measured by the growth assay. The DMSO diluted test compounds were printed in duplicate into 384-well black, clear-bottom plates by Tecan D300e Digital Dispenser along with dihydroartemisinin (5 μM) as killing control. Synchronized ring-stage parasites were cultured in the presence of the 18-point serial dilutions of the test compounds in culture medium (50 μL) at 1.0% parasitemia and 0.8% hematocrit for 72 h at 37 °C. SYBR Green I fluorescent dye (Invitrogen) in lysis buffer [20 mM Tris-HCl (Sangon Biotech), 5 mM EDTA (Thermo Scientific), 0.16% (w/v) Saponin (Sangon Biotech), 1.6% (v/v) Triton X-100 (Sangon Biotech)] was added, and after overnight incubation for optimal staining, fluorescence intensity was measured (excitation and emission wavelengths of 485 and 535 nm, respectively) by Envision (PerkinElmer). EC_50_ values were calculated using a nonlinear regression curve fit in Prism Software version 8 (GraphPad). The reported values were the results of two technical and at least two biological replicates.

### *In vitro* Drug Sensitivity and EC_50_ Determination with different glucose concentrations

Glucose (Sigma-Aldrich) was dissolved in glucose-free RPMI 1640 media (Thermo Fisher Scientific) to prepare glucose-containing culture media of the concentration of 2 mM, 4 mM, 6 mM, 8 mM, 10mM and 11mM, and adjusted to 7.4 pH and sterile-filtered. The drug susceptibility against Dd2 parasites was measured in the same way described above but extended incubation period to 48 hours.

### Substage selectivity and time course assay

Tightly synchronized 3D7 culture was obtained by a combination of sorbitol synchronization and heparin treatment. After sorbitol synchronization, the obtained ring-stage parasites were cultured with the presence of heparin sodium salt from Porcine Intestinal Mucosa (230 μg/mL; Sigma-Aldrich) until the majority of parasites developed to the late schizont stage. Then heparin was removed from culture to allow merozoites to invade erythrocytes. After 6 hours, the culture was sorbitol-synchronized, resulting in early ring-stage parasites (0~6 hpi), and resuspended in culture medium at 1.0% parasitemia and 0.8% hematocrit. The test compounds were prepared in the same way described above, and the obtained tightly synchronized culture was exposed to the tested compounds in the schedule schemed in Figure 6. After incubation with test compounds, the culture was freeze-thawed, and drug susceptibility was determined by LDH assay (42)

### Mammalian cell culture

HEK293T_17_ and HepG2 were purchased from ATCC then seeded in suitable plates filled with 89% DMEM (Gibco) supplemented with 10% inactivated FBS (Gibco) and 1% penicillin/ streptomycin (Gibco) maintained in an atmosphere of at 37°C with 5% CO2.

### *In vitro* cytotoxicity assay and CC_50_ Determination

Drug cytotoxicity was measured by cell viability assay. The test compounds in DMSO were printed into 384-well white, solid-bottom plates by Tecan D300e Digital Dispenser along with puromycin (5 μM) as killing control. HEK293T_17_ and HepG2 cells were seeded into the assay plates at approximately 2,000 cells/well and 2,500 cells/well respectively and incubated for 72 h. The viability was measured using Cell-Titer Glo (Promega) according to the manufacturers’ instructions, and luminescence signals were measured by Envision (PerkinElmer) with US LUM 384 setting. CC_50_ values were calculated using a nonlinear regression curve fit in Prism Software version 8 (GraphPad). The reported values were the results of two technical and at least two biological replicates.

### Extracellular flux analysis of *P. falciparum* using XFp analyzer

All assays were conducted according to the manufacturer’s manual with some modifications. A sensor cartridge was hydrated overnight in XF Calibrant Solution at 37 °C. Two assay media were employed for the analysis of parasites: RPMI 1640 medium ((Thermo Fisher Scientific, containing glucose 11 mM) and Seahorse XF RPMI Medium (Agilent Technologies). Injection solutions containing test compounds were prepared in assay medium at 10× of final concentration and loaded in the reagent delivery chambers of the sensor (20, 22, 24.5 and 27 μL for the first, second, third and fourth injections, respectively). Late-stage Dd2 parasites in RBCs were magnetically purified from 5% sorbitol-synchronized cultures using MACS LD columns (Miltenyi Biotec) and seeded at 0.8-million RBCs/well in a Seahorse miniplate which was precoated with Cell-Tak cell and tissue adhesive (Corning). Ring-stage Dd2 parasites in RBCs were obtained at roughly 20% parasitemia by culturing MACS-purified schizonts with a small amount of fresh blood overnight and seeded at 0.8-million RBCs/well. Freed-schizonts were prepared by saponin lysis (22) and seeded at 2.5-million RBCs/well. After centrifugation at 500 rpm for 5 min with slow acceleration and no braking, assay medium was added to all wells (180 μL as final volume), and the miniplate was loaded into the flux analyzer to start measurements (mix time: 30 sec; wait time: 1 min 30 sec; measure time: 3 min). In an assay plate, two wells were used for background correction.

### Extracellular flux analysis of HEK293T_17_ using XFp analyzer

For analysis of HEK293T_17_ cells, Seahorse XF DMEM Medium (Agilent Technologies) supplemented with glucose (25 mM), pyruvate (1mM), and glutamine (4mM) was used. HEK293T_17_ cells were pre-seeded at 60,000 cells/well in culture medium (80 μL) in a Seahorse miniplate precoated with Cell-Tak and adhesive in the same way described above. After culturing overnight, the medium was exchanged to the assay medium (180 μL/well), and the cells were placed in the non-CO_2_ incubator at 37 °C for 1h. The assay was conducted in the same way described above.

### Data Availability

The coordinates and structure factors for GLUT3 bound to C3361 have been deposited in the Protein Data Bank (PDB), http://www.wwpdb.org (PDB ID code 7CRZ).

## Author Contributions

H.Y. conceived the project. J.H., Y.Y., N.Z., X.J., N.K., N.Y., and H.Y. designed all experiments. H.Y. and J.H. are responsible for inhibitor development. J.H. and S.L. are responsible for computational simulations. J.H., D.P., Q.T., Y.X., T.Z., S.L., and Z.L. are responsible for chemical synthesis and characterizations. Y.Y., X.J., and S.Z. performed experiments related to protein generation, crystallization, and structural determination. Y.Y., J.H., X.J., S.Z., and N.W performed all biochemical assays. N.Z., T.S.K., and N.K. designed, executed, and analyzed parasitological and cytotoxicity experiments. All authors analyzed the data. H.Y., N.K, N.Y., J.H., X.J., Y.Y. and Y.X. prepared the manuscript.

## Acknowledgements

This work was supported by funds from the National Natural Science Foundation of China (Grant No. 21825702, 31630017, and 81861138009). H.Y. acknowledges the Beijing Outstanding Young Scientist program Grant No. BJJWZYJH01201910003013. We thank supports from the Beijing Municipal Government and BILL & MELINDA GATES foundation to GHDDI. We thank Kunio Hirata at Super Photon ring-8 (SPring-8, Japan) BL32XU beamline for the help of crystal screening and data collection. We thank the X-ray Crystallography Platform of the Tsinghua University Technology Center for Protein Research for the crystallization facility.

