## Supplemental Information for "Orthosteric-allosteric dual inhibitors of PfHT1 as selective anti-malarial agents"

**This PDF file includes:**

Supplementary text
Figures S1 to S5
Tables S1 to S2
SI References

### Supplementary Information Text

#### Chemical synthesis and characterizations

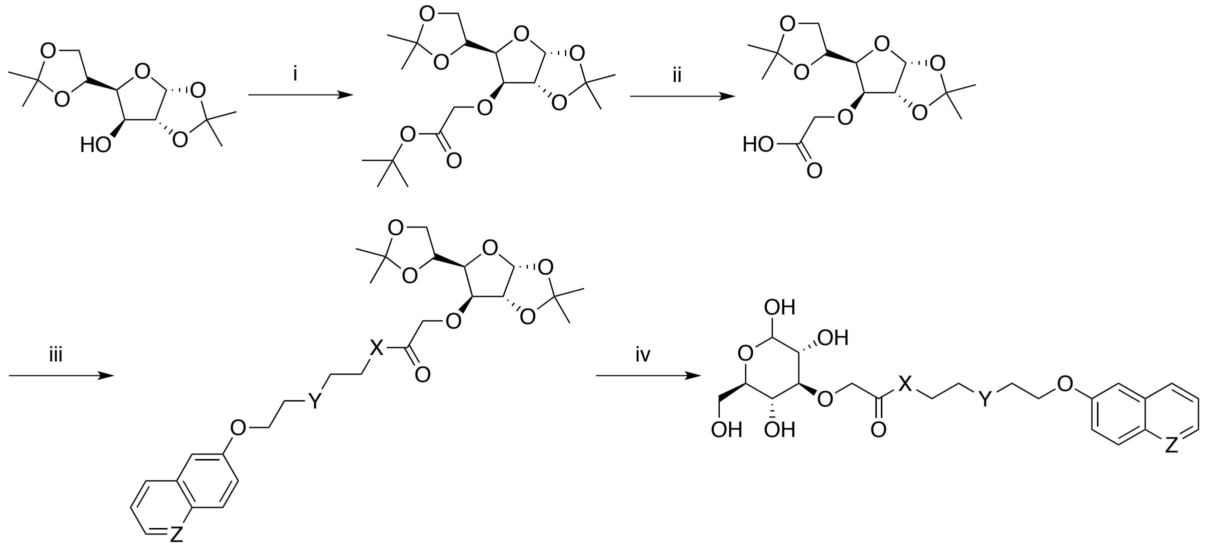

X is -O- or -NH-, Y is -O- or $-CH_2-$, Z is -CH= or -N=.

**Scheme S1: Synthesis of the D-glucose derivatives via ester or amido bond**

Reagents and conditions: (i) *tert*-butyl 2-bromoacetate, NaH, $Bu_4N^+I^-$, DMF, R.T., 24 h; (ii) 1M NaOH, reflux, 2 h, then acidification by 1M HCl; (iii) DAMP, EDCI, DCM, R.T., overnight; (iv) water, TFA, R.T., 40 min.

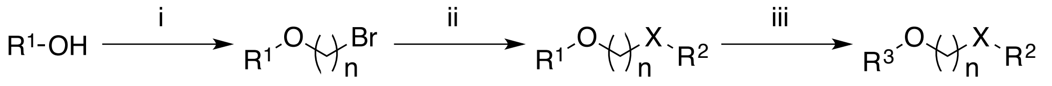

$R^1$ is a preprotected pyranose group, $R^2$ is partial substituted aryl or heteroaryl group, $R^3$ is deprotected pyranose group, X is -O- or -NH-, n is an integer from 7 to 11.

**Scheme S2: Synthesis of other pyranose derivatives via ether bond**

Reagents and conditions: (i) $Br-(CH_2)_n-Br$ ($n$ = 7 to 11), NaH, $Bu_4N^+I^-$, DMF, R.T., 24 h; (ii) $K_2CO_3$, DMF, 70 °C, overnight; (iii) water, TFA, R.T., 40 min. or $H_2$, Pb/C, methanol, R.T., 2-8 h.

#### General chemistry methods

NMR spectra were acquired on a Bruker AVANCE III HD 400 nuclear magnetic resonance spectrometer, running at 400 MHz for $^1H$ and 101 MHz for $^{13}C$, respectively. $^1H$ NMR spectra were recorded in $CHCl_3$-*d*, $(CH_3)_2SO$-$d_6$ and $(CH_3)_2CO$-$d_6$ using residual $CHCl_3$ (7.26 ppm), DMSO (2.50 ppm) and $(CH_3)_2CO$ (2.05 ppm) as the internal reference. $^{13}C$ NMR spectra were recorded in $CHCl_3$-*d*, $(CH_3)_2SO$-$d_6$ and $(CH_3)_2CO$-$d_6$ using residual $CHCl_3$ (77.16 ppm), DMSO (39.52 ppm), and $(CH_3)_2CO$ (29.84 ppm and 206.26 ppm) as the internal reference. Thin layer chromatography was performed on Merck Kieselgel 60 Å F254 plates eluting with the solvent described, visualized by a 254 nm UV lamp. Compounds were purified using flash chromatography (Innochem SilicaFlash P60, 230-400 mesh). Mass spectrometry was performed using a Thermo Scientific QExactive mass spectrometer (ESI). Unless otherwise noted, analytical grade solvents and commercially available reagents were used without further purification.

### Synthesis

#### Synthesis of *tert*-butyl 2-(((3a*R*,5*R*,6*S*,6a*R*)-5-((*R*)-2,2-dimethyl-1,3-dioxolan-4-yl)-2,2-dimethyltetrahydrofuro[2,3-*d*][1,3]dioxol-6-yl)oxy)acetate (1)

Diacetone-D-glucose (1.00 g, 3.84 mmol) was dissolved in anhydrous *N,N*-dimethylformamide (DMF) (25 ml) in a 100-ml round-bottomed flask. Sodium hydride (230 mg, 5.75 mmol) in mineral oil (60%) and tetra-*n*-butylammonium iodide (500 g, 1.35 mmol) were then added under argon in small portions over a period of 5 minutes. The mixture was stirred for 30 minutes in ice-water bath. Then *tert*-butyl 2-bromoacetate (2.24 g, 11.48 mmol) was added slowly. The mixture was allowed to warm to room temperature and stirred for 24 h. The progress of the reaction was monitored by TLC. Upon completion of the reaction, organic solvents were removed under vacuum, generating precipitants in water. The resulting residue was extracted with ethyl acetate, dried over anhydrous sodium sulfate, filtered, and concentrated under vacuum. The crude product was purified with flash column chromatography on silica gel (10 % ethyl acetate in petroleum ether) to yield 1.40 g (97.4%) *tert*-butyl 2-(((3a*R*,5*R*,6*S*,6a*R*)-5-((*R*)-2,2-dimethyl-1,3-dioxolan-4-yl)-2,2-dimethyltetrahydrofuro[2,3-*d*][1,3]dioxol-6-yl)oxy)acetate (**1**) as faint yellow oil. $^1$H NMR (400 MHz, Chloroform-*d*) δ 5.87 (d, *J* = 3.7 Hz, 1H), 4.69 (d, *J* = 3.7 Hz, 1H), 4.31 (q, *J* = 6.4 Hz, 1H), 4.16 – 4.03 (m, 4H), 4.01 – 3.94 (m, 1H), 3.92 (d, *J* = 2.8 Hz, 1H), 1.49 – 1.43 (m, 12H), 1.40 (s, 3H), 1.33 (s, 3H), 1.29 (s, 3H). $^{13}$C NMR (101 MHz, $CDCl_3$) δ 169.50, 111.90, 109.05, 105.30, 83.57, 83.40, 82.02, 81.20, 72.77, 68.98, 67.26, 28.21, 26.94, 26.89, 26.34, 25.49.

#### Synthesis of 2-(((3a*R*,5*R*,6*S*,6a*R*)-5-((*R*)-2,2-dimethyl-1,3-dioxolan-4-yl)-2,2-dimethyltetrahydro furo[2,3-*d*][1,3]dioxol-6-yl)oxy)acetic acid (2)

Compound **1** (1.40 g, 3.74 mmol) was dissolved in 20 ml 1M sodium hydroxide aqueous in a 100-ml round-bottomed flask. The mixture was heated to reflux for 2 h. Then the mixture was allowed to cool to room temperature and neutralized by 1M hydrochloric acid. The resulting residue was extracted with dichloromethane, dried over anhydrous sodium sulfate, filtered, and concentrated under vacuum to give the crude product 2-(((3a*R*,5*R*,6*S*,6a*R*)-5-((*R*)-2,2-dimethyl-1,3-dioxolan-4-yl)-2,2-dimethyltetrahydro furo[2,3-*d*][1,3]dioxol-6-yl)oxy)acetic acid (**2**) as white solid (1.09 g, 91.2%), which was used in the next step without further purification. $^1$H NMR (400 MHz, DMSO-$d_6$) δ 12.67 (s, 1H), 5.84 (d, *J* = 3.7 Hz, 1H), 4.70 (d, *J* = 3.7 Hz,

1H), 4.28 (q, *J* = 6.2 Hz, 1H), 4.22 – 4.05 (m, 3H), 4.00 (dd, *J* = 8.4, 6.4 Hz, 1H), 3.90 (d, *J* = 3.0 Hz, 1H), 3.80 (dd, *J* = 8.4, 6.3 Hz, 1H), 1.39 (s, 3H), 1.32 (s, 3H), 1.27 (s, 3H), 1.25 (s, 3H).

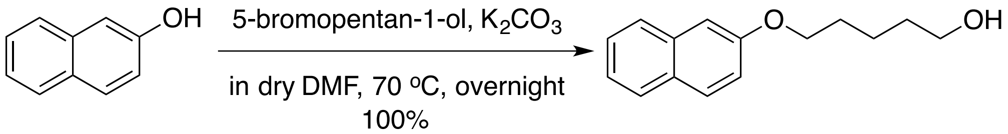

**Synthesis of 5-(naphthalen-2-yloxy)pentan-1-ol (3)**

Naphthalen-2-ol (500 mg, 3.47 mmol) was dissolved in 20 ml anhydrous DMF in a 100-ml round-bottomed flask. 5-bromopentan-1-ol (868 mg, 5.20 mmol) and potassium carbonate (960 mg, 6.94 mmol) were then added. The mixture was stirred at 70 °C for 24 h. The progress of the reaction was monitored by TLC. Upon completion of the reaction, the mixture was cooled to room temperature and organic solvents were removed under vacuum, generating precipitants in water. The resulting residue was extracted with ethyl acetate, dried over anhydrous sodium sulfate, filtered, and concentrated under vacuum. The crude product was purified by flash column chromatography on silica gel (25 % ethyl acetate in petroleum ether) to yield 781 mg (97.6%) 5-(naphthalen-2-yloxy)pentan-1-ol (**3**) as faint yellow solid. $^{1}$H NMR (400 MHz, Chloroform-*d*) δ 7.90 – 7.68 (m, 3H), 7.45 (ddd, *J* = 8.1, 6.8, 1.3 Hz, 1H), 7.35 (ddd, *J* = 8.1, 6.8, 1.2 Hz, 1H), 7.23 – 7.10 (m, 2H), 4.07 (t, *J* = 6.4 Hz, 2H), 3.68 (t, *J* = 6.3 Hz, 2H), 2.06 (s, 1H), 1.92 – 1.84 (m, 2H), 1.71 – 1.63 (m, 2H), 1.59 (dddd, *J* = 12.1, 7.3, 5.1, 1.6 Hz, 2H). $^{13}$C NMR (101 MHz, $CDCl_3$) δ 157.03, 134.64, 129.38, 128.95, 127.68, 126.74, 126.36, 123.56, 119.00, 106.62, 67.85, 62.71, 32.46, 29.06, 22.48.

**Synthesis of 5-(naphthalen-2-yloxy)pentyl 2-(((3a*R*,5*R*,6*S*,6a*R*)-5-((*R*)-2,2-dimethyl-1,3-dioxolan-4-yl)-2,2-dimethyltetrahydrofuro[2,3-*d*][1,3]dioxol-6-yl)oxy)acetate (4)**

Compound **3** (165 mg, 0.72 mmol), compound **2** (250 mg, 0.79 mmol), DAMP (20 mg, 0.16 mmol) and EDCI (165 mg, 0.86 mmol) were dissolved in 5 ml anhydrous dichloromethane (DCM). The mixture was stirred overnight at room temperature. The progress of the reaction was monitored by TLC. Upon completion of the reaction, organic solvents were removed under vacuum, generating precipitants in water. The resulting residue was extracted with ethyl acetate, dried over anhydrous sodium sulfate, filtered, and concentrated under vacuum. The crude product was purified with flash column chromatography on silica gel (20 % ethyl acetate in petroleum ether) to yield 329 mg (86.1%) 5-(naphthalen-2-yloxy)pentyl 2-(((3a*R*,5*R*,6*S*,6a*R*)-5-((*R*)-2,2-dimethyl-1,3-dioxolan-4-yl)-2,2-dimethyltetrahydrofuro[2,3-*d*][1,3]dioxol-6-yl)oxy)acetate (**4**) as faint yellow oil. $^{1}$H NMR (400 MHz, Chloroform-*d*) δ 7.73 (td, *J* = 8.9, 8.2, 6.2 Hz, 3H), 7.42 (ddd, *J* = 8.2, 6.8, 1.3 Hz, 1H), 7.32 (td, *J* = 7.4, 6.8, 1.2 Hz,

1H), 7.21 – 7.05 (m, 2H), 5.91 (d, *J* = 3.7 Hz, 1H), 4.73 (d, *J* = 3.7 Hz, 1H), 4.35 (dt, *J* = 7.7, 5.8 Hz, 1H), 4.26 (s, 2H), 4.23 – 3.97 (m, 8H), 1.86 (p, *J* = 6.6 Hz, 2H), 1.75 (p, *J* = 6.8 Hz, 2H), 1.61 – 1.53 (m, 2H), 1.49 (s, 3H), 1.43 (s, 3H), 1.36 (s, 3H), 1.30 (s, 3H). $^{13}$C NMR (101 MHz, $CDCl_3$) δ 170.32, 156.94, 134.60, 129.38, 128.94, 127.65, 126.70, 126.36, 123.56, 118.93, 111.83, 109.00, 106.53, 105.25, 83.66, 83.32, 81.10, 72.68, 68.37, 67.53, 67.22, 64.94, 28.86, 28.38, 26.85, 26.27, 25.41, 22.66.

**Synthesis of 5-(naphthalen-2-yloxy)pentyl 2-(((3*R*,4*S*,5*R*,6*R*)-2,3,5-trihydroxy-6-(hydroxyl methyl)tetrahydro-2*H*-pyran-4-yl)oxy)acetate (1a)**

Compound **4** (308 mg, 0.58 mmol) was dissolved in 2 ml trifluoroacetic acid in 25-ml round-bottomed flask, and water (0.2 ml) were then added. The mixture was stirred at room temperature for 40 minutes. The progress of the reaction was monitored by TLC. Upon completion of the reaction, organic solvents were directly evaporated under vacuum. The resulting residue was purified by flash column chromatography on silica gel (10 % methanol in dichloromethane) to yield 102 mg (39.0%) 5-(naphthalen-2-yloxy)pentyl 2-(((3*R*,4*S*,5*R*,6*R*)-2,3,5-trihydroxy-6-(hydroxyl methyl)tetrahydro-2*H*-pyran-4-yl)oxy)acetate (**1a**) as white solid. Yield: 39.0%. ESI-MS *m/z*: 473.1783 $[M+Na]^+$. $^1$H NMR (400 MHz, DMSO-$d_6$) δ 7.96 – 7.74 (m, 3H), 7.44 (ddd, *J* = 8.2, 6.9, 1.3 Hz, 1H), 7.40 – 7.27 (m, 2H), 7.15 (dd, *J* = 8.9, 2.5 Hz, 1H), 6.82 – 6.39 (m, 1H), 5.17 – 5.00 (m, 1H), 5.00 – 4.52 (m, 2H), 4.49 – 4.27 (m, 3H), 4.21 – 4.02 (m, 4H), 3.70 – 3.37 (m, 3H), 3.30 – 3.03 (m, 3H), 1.80 (p, *J* = 6.7 Hz, 2H), 1.68 (p, *J* = 6.8 Hz, 2H), 1.50 (tt, *J* = 10.0, 6.4 Hz, 2H). $^{13}$C NMR (101 MHz, DMSO) δ 172.12, 171.91, 156.55, 134.34, 129.26, 128.42, 127.51, 126.68, 126.36, 123.49, 118.78, 106.62, 96.72, 92.12, 86.14, 83.12, 76.49, 74.39, 72.00, 71.97, 69.44, 69.37, 68.87, 68.81, 67.36, 64.35, 60.81, 28.28, 27.86, 27.85, 22.10.

**Synthesis of 2-(2-(naphthalen-2-yloxy)ethoxy)ethyl 2-(((3*R*,4*S*,5*R*,6*R*)-2,3,5-trihydroxy-6-(hydroxymethyl)tetrahydro-2*H*-pyran-4-yl)oxy)acetate (1b)**

Compound **1b** was prepared in a similar manner described for **1a**. Yield: 23.1%. ESI-MS *m/z*: 475.1573 $[M+Na]^+$. $^1$H NMR (400 MHz, DMSO-$d_6$) δ 7.88 – 7.76 (m, 3H), 7.45 (ddd, *J* = 8.1, 6.7, 1.2 Hz, 1H), 7.39 – 7.30 (m, 2H), 7.18 (dd, *J* = 8.9, 2.5 Hz, 1H), 6.77 – 6.41 (m, 1H), 5.15 – 4.97 (m, 1H), 4.97 – 4.28 (m, 5H), 4.27 – 4.17 (m, 4H), 3.83 (dd, *J* = 5.7, 3.7 Hz, 2H), 3.73 (dd, *J* = 5.7, 3.7 Hz, 2H), 3.67 – 3.37 (m, 3H), 3.29 – 3.02 (m, 3H). $^{13}$C NMR (101 MHz, DMSO) δ 171.93, 171.74, 156.32, 134.27, 129.34, 128.50, 127.53, 126.72, 126.41, 123.61, 118.73, 106.72, 96.71, 92.12, 86.13, 83.09, 76.48, 74.38, 71.99, 71.95, 69.43, 69.34, 68.89, 68.85, 68.30, 67.11, 63.70, 60.80.

**Synthesis of 5-(quinolin-6-yloxy)pentyl 2-(((3*R*,4*S*,5*R*,6*R*)-2,3,5-trihydroxy-6-(hydroxymethyl)tetrahydro-2*H*-pyran-4-yl)oxy)acetate (1d)**

Compound **1d** was prepared in a similar manner described for **1a**. Yield: 26.2%. ESI-MS *m/z*: 452.1916 $[M+H]^+$. $^1$H NMR (400 MHz, DMSO-$d_6$) δ 8.73 (dd, *J* = 4.2, 1.7 Hz, 1H), 8.24 (dd, *J* = 8.4, 1.6 Hz, 1H), 7.91 (d, *J* = 9.0 Hz, 1H), 7.47 (dd, *J* = 8.3, 4.2 Hz, 1H), 7.43 – 7.34 (m, 2H), 6.88 – 6.31 (m, 1H), 5.03 (td, *J* = 32.1, 30.9, 4.0 Hz, 1H), 4.93 – 4.54 (m, 1H), 4.40 (s, 2H), 4.12 (tdd, *J* = 10.5, 6.5, 3.3 Hz, 4H), 3.48 – 3.02 (m, 8H), 1.82 (p, *J* = 6.7 Hz, 2H), 1.69 (p, *J* = 6.8 Hz, 2H), 1.55 – 1.46 (m, 2H). $^{13}$C NMR (101 MHz, DMSO) δ 172.11, 171.90, 156.55, 147.80, 143.60, 134.92, 130.24, 129.10, 122.31, 121.66, 106.38, 96.72, 92.12, 86.13, 83.11, 76.48, 74.39, 72.00, 71.96, 69.42, 69.35, 68.87, 68.81, 67.73, 66.37, 64.33, 60.80, 28.19, 27.84, 27.83, 22.06.

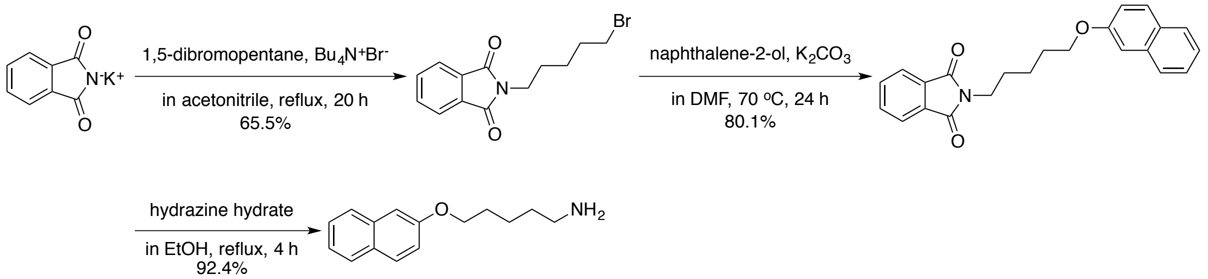

**Synthesis of 2-(5-bromopentyl)isoindoline-1,3-dione (5)**

A mixture of potassium phthalimide (500 mg, 2.70 mmol), 1,5-dibromopentane (1.55 g, 6.74 mmol) and catalytic amount of TBAB (174 mg, 0.54 mmol) were stirred in 20 ml acetonitrile under reflux for 20 h. Subsequently, the reaction mixture was filtered and the filtrate was concentrated under vacuum. The resulting residue was purified with flash column chromatography on silica gel (50 % ethyl acetate in petroleum ether) to yield 524 mg (65.5%) 2-(5-bromopentyl)isoindoline-1,3-dione (**5**) as faint yellow oil. $^1$H NMR (400 MHz, Chloroform-*d*) δ 7.83 (dt, *J* = 7.0, 3.5 Hz, 2H), 7.70 (dd, *J* = 5.5, 3.0 Hz, 2H), 3.69 (t, *J* = 7.2 Hz, 2H), 3.39 (t, *J* = 6.7 Hz, 2H), 1.90 (p, *J* = 6.9 Hz, 2H), 1.71 (p, *J* = 7.4 Hz, 2H), 1.54 – 1.44 (m, 2H).

**Synthesis of 2-(5-(naphthalen-2-yloxy)pentyl)isoindoline-1,3-dione (6)**

Compound **5** (500 mg, 1.69 mmol) was dissolved in 10 ml anhydrous DMF in a 50-ml round-bottomed flask. Naphthalen-2-ol (200 mg, 1.39 mmol) and potassium carbonate (383 mg, 2.77 mmol) were then added. The mixture was stirred at 70 °C for 24 h. The progress of the reaction was monitored by TLC. Upon completion of the reaction, the mixture was cooled to room temperature and organic solvents were removed under vacuum, generating precipitants in water. The resulting residue was extracted with ethyl acetate, dried over anhydrous sodium sulfate, filtered, and concentrated under vacuum. The crude product was purified by flash column chromatography on silica gel (50 % ethyl acetate in petroleum ether) to yield 486 mg

(80.1%) 2-(5-(naphthalen-2-yloxy)pentyl) isoindoline-1,3-dione (**6**) as faint yellow oil. $^1$H NMR (400 MHz, Chloroform-*d*) δ 7.84 (dt, *J* = 7.6, 3.7 Hz, 2H), 7.79 – 7.65 (m, 5H), 7.42 (ddd, *J* = 8.2, 6.8, 1.3 Hz, 1H), 7.31 (ddd, *J* = 8.1, 6.8, 1.2 Hz, 1H), 7.17 – 7.05 (m, 2H), 4.07 (t, *J* = 6.4 Hz, 2H), 3.74 (t, *J* = 7.2 Hz, 2H), 1.95 – 1.85 (m, 2H), 1.80 (p, *J* = 7.4 Hz, 2H), 1.58 (qd, *J* = 9.6, 9.1, 5.9 Hz, 2H).

**Synthesis of 5-(naphthalen-2-yloxy)pentan-1-amine (7)**

A solution of 2-(5-(naphthalen-2-yloxy)pentyl) isoindoline-1,3-dione (486 mg, 1.4 mmol) and hydrazine hydrate (80%, 1.6 mL, 25 mmol) in 10 ml ethanol was reflux for 4 h, and then cooled to room temperature. The mixture was filtered and the solid was washed with 95% ethanol. The whole filtrate was concentrated and the resulting residue was dissolved in DCM. The mixture was dried over anhydrous sodium sulfate, filtered, and concentrated to give the crude product, which was used in the next step without further purification. $^1$H NMR (400 MHz, DMSO-$d_6$) δ 7.89 – 7.70 (m, 3H), 7.45 (t, *J* = 7.6 Hz, 1H), 7.37 – 7.25 (m, 2H), 7.15 (dd, *J* = 8.9, 2.5 Hz, 1H), 4.07 (t, *J* = 6.5 Hz, 2H), 2.56 (t, *J* = 6.4 Hz, 2H), 2.21 (d, *J* = 97.3 Hz, 0H), 1.78 (p, *J* = 6.7 Hz, 2H), 1.46 (qd, *J* = 13.5, 11.8, 6.6 Hz, 4H).

**Synthesis of 2-(((3a*R*,5*R*,6*S*,6a*R*)-5-((*R*)-2,2-dimethyl-1,3-dioxolan-4-yl)-2,2-dimethyltetra hydrofuro[2,3-*d*][1,3]dioxol-6-yl)oxy)-*N*-(5-(naphthalen-2-yloxy)pentyl)acetamide (8)**

Compound **7** (303 mg, 1.32 mmol), compound **2** (465 mg, 1.46 mmol), DAMP (33 mg, 0.27 mmol) and EDCI (315 mg, 1.65 mmol) were dissolved in 15 ml anhydrous DCM. The mixture was stirred at room temperature overnight. The progress of the reaction was monitored by TLC. Upon completion of the reaction, organic solvents were removed under vacuum, generating precipitants in water. The resulting residue was extracted with ethyl acetate, dried over anhydrous sodium sulfate, filtered, and concentrated under vacuum. The crude product was purified with flash column chromatography on silica gel (25 % ethyl acetate in petroleum ether) to yield 238 mg (34.0%) 2-(((3a*R*,5*R*,6*S*,6a*R*)-5-((*R*)-2,2-dimethyl-1,3-dioxolan-4-yl)-2,2-dimethyltetrahydrofuro[2,3-*d*][1,3]dioxol-6-yl)oxy)-*N*-(5-(naphthalen-2-yloxy)pentyl)acetamide (**8**) as faint yellow oil. $^1$H NMR (400 MHz, Chloroform-*d*) δ 7.85 – 7.63 (m, 3H), 7.48 (t, *J* = 6.0 Hz, 1H), 7.43 (t, *J* = 7.5 Hz, 1H), 7.32 (t, *J* = 7.5 Hz, 1H), 7.20 – 7.04 (m, 2H), 5.90 (d, *J* = 3.7 Hz, 1H), 4.54 (d, *J* = 3.7 Hz, 1H), 4.35 (ddd, *J* = 9.8, 6.1, 4.3 Hz, 1H), 4.23 – 3.94 (m, 8H), 3.38 (dq, *J* = 13.8, 6.8 Hz, 1H), 3.26 (dq, *J* = 13.5, 6.7 Hz, 1H), 1.88 (p, *J* = 6.4 Hz, 2H), 1.59 (dp, *J* = 22.3, 7.9 Hz, 4H), 1.49 (s, 3H), 1.43 (s, 3H), 1.37 (s, 3H), 1.32 (s, 3H). $^{13}$C NMR (101 MHz, $CDCl_3$) δ 169.09, 156.95, 134.58, 129.34, 128.91, 127.63, 126.68, 126.32, 123.53, 118.91, 112.13, 109.58, 106.53, 105.35, 82.79, 82.03, 80.87, 72.51, 68.40, 67.90, 67.57, 38.93, 29.75, 28.94, 27.10, 26.79, 26.18, 25.40, 23.66.

**Synthesis of *N*-(5-(naphthalen-2-yloxy)pentyl)-2-(((3*R*,4*S*,5*R*,6*R*)-2,3,5-trihydroxy-6-(hydroxymethyl)tetrahydro-2*H*-pyran-4-yl)oxy)acetamide (1c)**

Compound **8** (238 mg, 0.45 mmol) was dissolved in 3.6 ml trifluoroacetic acid in 25-ml round-bottomed flask, and water (0.4 ml) were then added. The mixture was stirred at room temperature for 40 minutes. The progress of the reaction was monitored by TLC. Upon completion of the reaction, organic solvents were directly evaporated under vacuum. The resulting residue was purified by flash column chromatography on silica gel (10 % methanol in dichloromethane) to yield 202 mg (99.9%) *N*-(5-(naphthalen-2-yloxy)pentyl)-2-(((3*R*,4*S*,5*R*,6*R*)-2,3,5-trihydroxy-6-(hydroxymethyl)tetrahydro-2*H*-pyran-4-yl)oxy)acetamide (**1c**) as faint yellow solid. Yield: 99.9%. ESI-MS *m/z*: 450.2131 $[M+H]^+$. $^1$H NMR (400 MHz, DMSO-$d_6$) δ 8.33 (dt, *J* = 16.6, 5.7 Hz, 1H), 7.80 (dt, *J* = 7.7, 3.1 Hz, 3H), 7.44 (t, *J* = 7.5 Hz, 1H), 7.38 – 7.25 (m, 2H), 7.15 (dd, *J* = 9.0, 2.5 Hz, 1H), 6.90 – 6.42 (m, 1H), 5.75 – 5.19 (m, 2H), 4.77 (dt, *J* = 142.1, 4.7 Hz, 1H), 4.42 (dt, *J* = 59.2, 6.1 Hz, 1H), 4.27 – 4.13 (m, 2H), 4.06 (t, *J* = 6.5 Hz, 2H), 3.73 – 3.02 (m, 8H), 1.78 (p, *J* = 6.7 Hz, 2H), 1.49 (dq, *J* = 21.7, 8.4, 7.9 Hz, 4H). $^{13}$C NMR (101 MHz, DMSO) δ 171.06, 170.94, 157.01, 134.78, 129.69, 128.85, 127.94, 127.12, 126.79, 123.92, 119.23, 107.03, 97.07, 92.65, 86.17, 83.35, 76.83, 74.80, 72.58, 72.44, 71.29, 71.19, 70.23, 70.08, 67.87, 61.20, 38.52, 38.50, 29.27, 29.20, 28.81, 23.49.

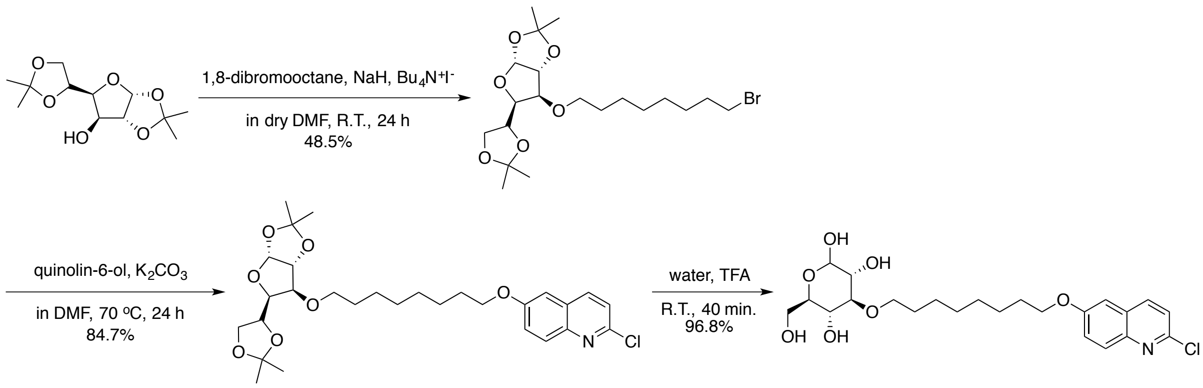

**Synthesis of (3a*R*,5*R*,6*S*,6a*R*)-6-((8-bromooctyl)oxy)-5-((*R*)-2,2-dimethyl-1,3-dioxolan-4-yl)-2,2-dimethyltetrahydrofuro[2,3-*d*][1,3]dioxole (9)**

Diacetone-D-glucose (5.00 g, 19.18 mmol) was dissolved in 25 ml anhydrous DMF in a 100-ml round-bottomed flask. Sodium hydride (1.20 g, 30.01 mmol) in mineral oil (60%) and tetra-*n*-butylammonium iodide (1.00 g, 2.71 mmol) were then added under argon in small portions over a period of 5 minutes. The mixture was stirred for 30 minutes in ice-water bath. Then 1,8-dibromooctane (5.3 ml, 28.78 mmol) was added slowly. The mixture was allowed to warm to room temperature and stirred for 24 h. The progress of the reaction was monitored by TLC. Upon completion of the reaction, organic solvents were removed under vacuum, generating precipitants in water. The resulting residue was extracted with ethyl acetate, dried over anhydrous sodium sulfate, filtered, and concentrated under vacuum. The crude product was purified with flash column chromatography on silica gel (8 % ethyl acetate in petroleum ether) to yield 4.20 g (48.51%) (3a*R*,5*R*,6*S*,6a*R*)-6-((8-bromooctyl)oxy)-5-((*R*)-2,2-dimethyl-1,3-dioxolan-4-yl)-2,2-dimethyltetrahydrofuro[2,3-*d*][1,3]dioxole (**9**) as faint yellow oil. ESI-MS *m/z*: 451.43 $[M+H]^+$. $^1$H NMR (400 MHz, Chloroform-*d*) δ 5.78 (d, *J* = 3.7 Hz, 1H), 4.44 (d, *J* = 3.7 Hz, 1H), 4.22 (q, *J* = 6.4 Hz, 1H), 4.04 (dd, *J* = 7.6, 2.8 Hz, 1H), 3.99 (dd, *J* = 8.5, 6.2 Hz, 1H), 3.89 (dd, *J* = 8.5, 6.0 Hz, 1H), 3.77 (d, *J* = 3.0 Hz, 1H), 3.53 (dt, *J* = 9.4, 6.4 Hz, 1H), 3.43 (dt, *J* = 9.3, 6.5 Hz, 1H), 3.32 (t, *J* = 6.8 Hz, 2H), 1.77 (p, *J* = 6.9 Hz, 2H), 1.53 – 1.44 (m, 2H), 1.41 (s, 3H), 1.35 (d, *J* = 4.7 Hz, 5H), 1.31 – 1.19 (m, 12H). $^{13}$C NMR (101 MHz, $CDCl_3$) δ 111.82, 108.98, 105.38, 82.68, 82.23, 81.31, 72.65, 70.74, 67.35, 34.03, 32.89, 29.80, 29.31, 28.81, 28.21, 26.97, 26.90, 26.38, 26.07, 25.55.

**Synthesis of 2-chloro-6-((8-(((3a*R*,5*R*,6*S*,6a*R*)-5-((*R*)-2,2-dimethyl-1,3-dioxolan-4-yl)-2,2-dimethyltetrahydrofuro[2,3-*d*][1,3]dioxol-6-yl)oxy)octyl)oxy)quinoline (10)**

Compound **9** (400 mg, 0.89 mmol) was dissolved in 10 ml anhydrous DMF in a 100-ml round-bottomed flask. 2-chloroquinolin-6-ol (160 mg, 0.89 mmol) and potassium carbonate (184 mg, 1.33 mmol) were then added into the solution. The mixture was stirred at 70℃ for 10 h. The progress of the reaction was monitored by TLC. Upon completion of the reaction, the mixture was cooled to room temperature and organic solvents were removed under vacuum. The resulting precipitant was extracted with ethyl acetate, dried over anhydrous sodium sulfate, filtered, and concentrated under vacuum. The crude product was purified using flash column chromatography on silica gel (10 % ethyl acetate in petroleum ether) to yield 477 mg (97.9%) 2-chloro-6-((8-(((3a*R*,5*R*,6*S*,6a*R*)-5-((*R*)-2,2-dimethyl-1,3-dioxolan-4-yl)-2,2-dimethyltetrahydrofuro[2,3-*d*][1,3]dioxol-6-yl)oxy)octyl)oxy)quinoline (**10**) as faint yellow oil. $^1$H NMR (400 MHz, Chloroform-*d*) δ 7.97 (dd, *J* = 8.7, 0.7 Hz, 1H), 7.90 (d, *J* = 9.2 Hz, 1H), 7.37 (dd, *J* = 9.2, 2.7 Hz, 1H), 7.32 (d, *J* = 8.6 Hz, 1H), 7.05 (d, *J* = 2.7 Hz, 1H), 5.87 (d, *J* = 3.7 Hz, 1H), 4.52 (d, *J* = 3.7 Hz, 1H), 4.30 (dt, *J* = 7.5, 6.0 Hz, 1H), 4.12 (dd, *J* = 7.5, 3.1 Hz, 1H), 4.09 – 4.01 (m, 3H), 3.98 (dd, *J* = 8.6, 6.0 Hz, 1H), 3.84 (d, *J* = 3.0 Hz, 1H), 3.60 (dt, *J* = 9.3, 6.4 Hz, 1H), 3.50 (dt, *J* = 9.2, 6.4 Hz, 1H), 1.84 (dt, *J* = 14.6, 6.5 Hz, 2H), 1.62 – 1.43 (m, 7H), 1.41 (s, 3H), 1.40 – 1.32 (m, 9H), 1.31 (s, 3H). $^{13}$C NMR (101 MHz, $CDCl_3$) δ 157.71, 148.02, 143.81, 137.72, 129.98, 128.07, 123.46, 122.56, 111.85, 108.99, 106.04, 105.38, 82.63, 82.22, 81.29, 72.67, 70.75, 68.47, 67.34, 29.85, 29.46, 29.25, 26.97, 26.90, 26.38, 26.17, 26.15, 25.56.

**Synthesis of (3*R*,4*S*,5*R*,6*R*)-4-((8-((2-chloroquinolin-6-yl)oxy)octyl)oxy)-6-(hydroxymethyl)tetrahydro-2*H*-pyran-2,3,5-triol (3a)**

Compound **10** (1.26 g, 2.29 mmol) was dissolved in 7.2 ml trifluoroacetic acid in 50-ml round-bottomed flask, and water (0.8 ml) were then added. The mixture was stirred at room temperature for 40 minutes. The progress of the reaction was monitored by TLC. Upon completion of the reaction, organic solvents were directly evaporated under vacuum. The resulting residue was purified by flash column chromatography on silica gel (10 % methanol in dichloromethane) to yield 878 mg (81.6%) (3*R*,4*S*,5*R*,6*R*)-4-((8-((2-chloroquinolin-6-yl)oxy)octyl)oxy)-6-(hydroxymethyl)tetrahydro-2*H*-pyran-2,3,5-triol (**3a**) as white solid. ESI-MS *m/z*: 470.1950 $[M+H]^+$. $^1H$ NMR (400 MHz, DMSO-$d_6$) δ 8.31 (dd, *J* = 8.7, 0.7 Hz, 1H), 7.91 – 7.80 (m, 1H), 7.51 (d, *J* = 8.6 Hz, 1H), 7.48 – 7.37 (m, 2H), 6.74 – 6.25 (m, 1H), 5.01 – 4.73 (m, 2H), 4.61 – 4.46 (m, 1H), 4.45 – 4.23 (m, 1H), 4.08 (t, *J* = 6.5 Hz, 2H), 3.73 – 2.89 (m, 8H), 1.77 (p, *J* = 6.7 Hz, 2H), 1.46 (dq, *J* = 26.6, 6.6 Hz, 4H), 1.38 – 1.22 (m, 6H). $^{13}C$ NMR (101 MHz, DMSO) δ 157.08, 147.02, 142.99, 138.75, 129.23, 128.05, 123.36, 122.48, 106.69, 96.95, 92.36, 85.20, 81.93, 76.75, 74.61, 72.12, 71.99, 71.94, 69.93, 69.76, 68.04, 61.11, 29.99, 29.96, 29.03, 28.85, 28.58, 25.61, 25.58, 25.55.

**Synthesis of (3*R*,4*S*,5*R*,6*R*)-6-(hydroxymethyl)-4-((8-(quinolin-6-ylamino)octyl)oxy)tetrahydro-2*H*-pyran-2,3,5-triol (1e)**

Compound **1e** was prepared in a similar manner as described for **3a**. Yield: 78.2%; ESI-MS *m/z*: 435.2489 $[M+H]^+$. $^1H$ NMR (400 MHz, DMSO-$d_6$) δ 8.45 (dd, *J* = 4.2, 1.7 Hz, 1H), 7.97 (dd, *J* = 8.3, 1.6 Hz, 1H), 7.68 (d, *J* = 9.1 Hz, 1H), 7.27 (dd, *J* = 8.3, 4.2 Hz, 1H), 7.19 (dd, *J* = 9.1, 2.6 Hz, 1H), 6.80 – 6.23 (m, 2H), 6.10 (t, *J* = 5.3 Hz, 1H), 4.97 – 4.73 (m, 2H), 4.50 (dd, *J* = 6.7, 4.3 Hz, 1H), 4.41 – 4.24 (m, 1H), 3.69 – 2.92 (m, 10H), 1.66 – 1.56 (m, 2H), 1.50 (p, *J* = 6.7 Hz, 2H), 1.40 (dq, *J* = 11.6, 6.8 Hz, 2H), 1.31 (d, *J* = 5.0 Hz, 6H). $^{13}C$ NMR (101 MHz, DMSO) δ 146.99, 144.79, 142.16, 133.03, 130.09, 129.30, 121.64, 121.24, 100.77, 96.94, 92.33, 85.17, 81.90, 76.72, 74.57, 72.09, 71.93, 71.89, 69.91, 69.74, 61.09, 42.92, 29.98, 29.95, 29.07, 28.96, 28.41, 26.80, 25.60, 25.57.

**Synthesis of (3*R*,4*S*,5*R*,6*R*)-6-(hydroxymethyl)-4-((7-(quinolin-6-yloxy)heptyl)oxy)tetrahydro-2*H*-pyran-2,3,5-triol (1f)**

Compound **1f** was prepared in a similar manner described for **3a**. Yield: 89.0%; ESI-MS *m/z*: 422.2166 $[M+H]^+$. $^1H$ NMR (400 MHz, DMSO-$d_6$) δ 8.72 (dd, *J* = 4.2, 1.7 Hz, 1H), 8.23 (dd, *J* = 8.4, 1.6 Hz, 1H), 7.90 (d, *J* = 9.0 Hz, 1H), 7.46 (dd, *J* = 8.3, 4.2 Hz, 1H), 7.41 – 7.33 (m, 2H), 6.78 – 6.25 (m, 1H), 5.02 – 4.74 (m, 2H), 4.61 – 4.47 (m, 1H), 4.43 – 4.24 (m, 1H), 4.09 (t, *J* = 6.5 Hz, 2H), 3.75 – 2.88 (m, 8H), 1.86 – 1.70 (m, 2H), 1.58 – 1.48 (m, 2H), 1.44 (q, *J* = 6.9 Hz, 2H), 1.40 – 1.26 (m, 4H). $^{13}C$ NMR (101 MHz, DMSO) δ 156.57, 147.82, 143.71, 134.76,

130.32, 129.08, 122.22, 121.61, 106.32, 96.96, 92.35, 85.19, 81.92, 76.74, 74.59, 72.11, 71.92, 71.88, 69.92, 69.75, 67.89, 61.10, 29.94, 29.90, 28.77, 28.59, 25.60, 25.57.

**Synthesis of (3*R*,4*S*,5*R*,6*R*)-6-(hydroxymethyl)-4-((9-(quinolin-6-yloxy)nonyl)oxy)tetrahydro-2*H*-pyran-2,3,5-triol (1g)**

Compound **1g** was prepared in a similar manner described for **3a**. Yield: 86.2%; ESI-MS *m/z*: 450.2483 $[M+H]^+$. $^1H$ NMR (400 MHz, DMSO-$d_6$) δ 8.72 (dd, *J* = 4.3, 1.7 Hz, 1H), 8.23 (dd, *J* = 8.4, 1.6 Hz, 1H), 7.90 (d, *J* = 9.0 Hz, 1H), 7.46 (dd, *J* = 8.3, 4.2 Hz, 1H), 7.42 – 7.31 (m, 2H), 6.71 – 6.27 (m, 1H), 4.97 – 4.77 (m, 2H), 4.58 – 4.44 (m, 1H), 4.43 – 4.23 (m, 1H), 4.09 (t, *J* = 6.5 Hz, 2H), 3.71 – 2.91 (m, 8H), 1.78 (p, *J* = 6.7 Hz, 2H), 1.47 (dt, *J* = 18.8, 7.1 Hz, 4H), 1.38 – 1.25 (m, 8H). $^{13}C$ NMR (101 MHz, DMSO) δ 156.56, 147.82, 143.71, 134.75, 130.32, 129.08, 122.22, 121.61, 106.33, 96.96, 92.35, 85.18, 81.91, 76.74, 74.58, 72.11, 71.96, 71.92, 69.91, 69.74, 67.86, 61.09, 29.99, 29.95, 29.07, 29.02, 28.81, 28.61, 25.58.

**Synthesis of (3*R*,4*S*,5*R*,6*R*)-6-(hydroxymethyl)-4-((11-(quinolin-6-yloxy)undecyl)oxy)tetrahydro-2*H*-pyran-2,3,5-triol (1h)**

Compound **1h** was prepared in a similar manner described for **3a**. Yield: 92.4%; ESI-MS *m/z*: 478.2798 $[M+H]^+$. $^1H$ NMR (400 MHz, DMSO-$d_6$) δ 8.72 (dd, *J* = 4.2, 1.7 Hz, 1H), 8.23 (ddd, *J* = 8.5, 1.8, 0.8 Hz, 1H), 7.90 (d, *J* = 9.0 Hz, 1H), 7.46 (dd, *J* = 8.3, 4.2 Hz, 1H), 7.43 – 7.31 (m, 2H), 6.76 – 6.26 (m, 1H), 5.00 – 4.75 (m, 2H), 4.58 – 4.45 (m, 1H), 4.43 – 4.22 (m, 1H), 4.08 (t, *J* = 6.5 Hz, 2H), 3.72 – 2.89 (m, 8H), 1.83 – 1.69 (m, 2H), 1.55 – 1.39 (m, 4H), 1.36 – 1.21 (m, 12H). $^{13}C$ NMR (101 MHz, DMSO) δ 156.57, 147.84, 143.72, 134.78, 130.33, 129.10, 122.24, 121.64, 106.33, 96.94, 92.36, 85.20, 76.75, 74.60, 72.12, 72.01, 69.92, 69.76, 67.86, 61.11, 29.98, 29.13, 29.10, 29.08, 29.06, 28.83, 28.62, 25.62, 25.59.

**Synthesis of (3*S*,4*R*,5*S*,6*S*)-4-((9-((2-chloroquinolin-6-yl)oxy)nonyl)oxy)-6-(hydroxymethyl)tetrahydro-2*H*-pyran-2,3,5-triol (1i)**

Compound **1i** was prepared in a similar manner described for **3a**. Yield: 78.2%; ESI-MS *m/z*: 506.1917 $[M+Na]^+$. $^1H$ NMR (400 MHz, DMSO-$d_6$) δ 8.31 (d, *J* = 8.6 Hz, 1H), 7.89 – 7.79 (m, 1H), 7.52 (d, *J* = 8.6 Hz, 1H), 7.48 – 7.37 (m, 2H), 6.77 – 6.25 (m, 1H), 4.98 – 4.77 (m, 2H), 4.56 – 4.46 (m, 1H), 4.38 – 4.25 (m, 1H), 4.13 – 4.04 (m, 2H), 3.67 – 2.90 (m, 8H), 1.77 (p, *J* = 6.7 Hz, 2H), 1.46 (dp, *J* = 22.6, 6.9 Hz, 4H), 1.38 – 1.19 (m, 8H). $^{13}C$ NMR (101 MHz, DMSO) δ 157.08, 147.01, 142.99, 138.74, 129.23, 128.04, 123.35, 122.48, 106.69, 96.94, 85.20, 76.75, 74.60, 71.99, 69.76, 68.03, 61.11, 29.97, 29.09, 29.03, 28.82, 28.58, 25.60, 25.57.

**Synthesis of (3*R*,4*S*,5*R*,6*R*)-4-((8-(benzo[*d*][1,3]dioxol-5-yloxy)octyl)oxy)-6-(hydroxymethyl)tetrahydro-2*H*-pyran-2,3,5-triol (2a)**

Compound **2a** was prepared in a similar manner described for **3a**. Yield: 91.8%; ESI-MS *m/z*: 451.1942 $[M+Na]^+$. $^1H$ NMR (400 MHz, DMSO-$d_6$) δ 6.78 (d, *J* = 8.5 Hz, 1H), 6.70 – 6.38 (m, 2H), 6.33 (dd, *J* = 8.5, 2.5 Hz, 1H), 5.94 (s, 2H), 4.95 – 4.79 (m, 2H), 4.57 – 4.47 (m, 1H), 4.42 – 4.23 (m, 1H), 3.85 (t, *J* = 6.5 Hz, 2H), 3.72 – 2.90 (m, 8H), 1.71 – 1.59 (m, 2H), 1.55 – 1.43 (m, 2H), 1.42 – 1.24 (m, 8H). $^{13}C$ NMR (101 MHz, DMSO) δ 154.15, 147.91, 140.94, 108.04, 105.61, 100.93, 97.74, 96.94, 85.20, 76.75, 74.60, 71.99, 69.76, 68.19, 61.11, 29.95, 29.03, 28.87, 28.77, 25.57, 25.56.

**Synthesis of (3*R*,4*S*,5*R*,6*R*)-6-(hydroxymethyl)-4-((8-(naphthalen-2-yloxy)octyl)oxy)tetrahydro-2*H*-pyran-2,3,5-triol (2b)**

Compound **2b** was prepared in a similar manner described for **3a**. Yield: 83.2%; ESI-MS *m/z*: 457.2202 $[M+Na]^+$. $^1H$ NMR (400 MHz, DMSO-$d_6$) δ 7.80 (dt, *J* = 8.4, 3.9 Hz, 3H), 7.44 (ddd, *J* = 8.1, 6.8, 1.3 Hz, 1H), 7.39 – 7.27 (m, 2H), 7.15 (dd, *J* = 9.0, 2.5 Hz, 1H), 6.64 – 6.24 (m, 1H), 4.93 – 4.73 (m, 2H), 4.56 – 4.41 (m, 1H), 4.40 – 4.22 (m, 1H), 4.07 (t, *J* = 6.5 Hz, 2H), 3.70 – 2.91 (m, 8H), 1.77 (p, *J* = 6.8 Hz, 2H), 1.48 (dp, *J* = 22.6, 6.9 Hz, 4H), 1.32 (q, *J* = 7.3, 6.3 Hz, 6H). $^{13}C$ NMR (101 MHz, DMSO) δ 156.57, 134.31, 129.19, 128.37, 127.45, 126.62, 126.29, 123.41, 118.73, 106.59, 96.91, 92.32, 85.17, 81.89, 76.71, 74.58, 72.07, 71.90, 71.87, 69.92, 69.75, 67.50, 61.10, 29.95, 29.91, 28.99, 28.83, 28.67, 25.57, 25.54.

**Synthesis of (3*R*,4*S*,5*R*,6*R*)-6-(hydroxymethyl)-4-((8-(naphthalen-1-yloxy)octyl)oxy)tetrahydro-2*H*-pyran-2,3,5-triol (2c)**

Compound **2c** was prepared in a similar manner described for **3a**. Yield: 94.2%; ESI-MS *m/z*: 457.2205 $[M+Na]^+$. $^1H$ NMR (400 MHz, DMSO-$d_6$) δ 8.20 – 8.11 (m, 1H), 7.90 – 7.81 (m, 1H), 7.55 – 7.36 (m, 4H), 6.94 (dd, *J* = 7.5, 1.2 Hz, 1H), 6.71 – 6.27 (m, 1H), 5.00 – 4.77 (m, 2H), 4.57 – 4.46 (m, 1H), 4.43 – 4.22 (m, 1H), 4.13 (t, *J* = 6.3 Hz, 2H), 3.74 – 2.91 (m, 8H), 1.93 – 1.77 (m, 2H), 1.51 (h, *J* = 6.5 Hz, 4H), 1.43 – 1.27 (m, 6H). $^{13}C$ NMR (101 MHz, DMSO) δ 154.13, 134.03, 127.47, 126.42, 126.30, 125.26, 124.98, 121.48, 119.73, 105.08, 96.95, 92.36, 85.21, 81.94, 76.76, 74.61, 72.13, 72.12, 72.00, 71.96, 69.93, 69.76, 67.74, 61.12, 30.02, 29.98, 29.06, 28.91, 28.75, 25.79, 25.63, 25.60.

**Synthesis of (3*R*,4*S*,5*R*,6*R*)-4-((8-([1,1'-biphenyl]-4-yloxy)octyl)oxy)-6-(hydroxymethyl)tetrahydro-2*H*-pyran-2,3,5-triol (2d)**

Compound **2d** was prepared in a similar manner described for **3a**. Yield: 93.6%; ESI-MS *m/z*: 483.2362 $[M+Na]^+$. $^1H$ NMR (400 MHz, DMSO-$d_6$) δ 7.65 – 7.55 (m, 4H), 7.42 (t, *J* = 7.7 Hz, 2H), 7.35 – 7.26 (m, 1H), 7.06 – 6.96 (m, 2H), 6.68 – 6.27 (m, 1H), 4.97 – 4.75 (m, 2H), 4.56 – 4.45 (m, 1H), 4.41 – 4.22 (m, 1H), 3.99 (t, *J* = 6.5 Hz, 2H), 3.74 – 2.87 (m, 8H), 1.78 – 1.64 (m, 2H), 1.50 (t, *J* = 6.5 Hz, 2H), 1.45 – 1.37 (m, 2H), 1.36 – 1.25 (m, 6H). $^{13}C$ NMR (101 MHz,

DMSO) δ 158.79, 140.32, 132.82, 129.31, 128.19, 127.12, 126.60, 115.32, 97.39, 92.80, 85.65, 82.37, 77.20, 75.05, 72.56, 72.42, 72.37, 70.38, 70.21, 67.96, 61.56, 30.43, 30.40, 29.47, 29.31, 29.18, 26.04, 26.01.

**Synthesis of (3*R*,4*S*,5*R*,6*R*)-4-((8-([1,1'-biphenyl]-3-yloxy)octyl)oxy)-6-(hydroxymethyl)tetrahydro-2*H*-pyran-2,3,5-triol (2e)**

Compound **2e** was prepared in a similar manner described for **3a**. Yield: 82.0%; ESI-MS *m/z*: 483.2349 $[M+Na]^+$. $^1$H NMR (400 MHz, DMSO-$d_6$) δ 7.70 – 7.61 (m, 2H), 7.50 – 7.41 (m, 2H), 7.40 – 7.31 (m, 2H), 7.20 (dt, *J* = 7.6, 1.2 Hz, 1H), 7.16 (t, *J* = 2.1 Hz, 1H), 6.92 (ddd, *J* = 8.2, 2.6, 0.9 Hz, 1H), 6.71 – 6.27 (m, 1H), 4.95 – 4.79 (m, 2H), 4.56 – 4.49 (m, 1H), 4.44 – 4.23 (m, 1H), 4.02 (t, *J* = 6.5 Hz, 2H), 3.70 – 2.91 (m, 8H), 1.73 (p, *J* = 6.6 Hz, 2H), 1.45 (dq, *J* = 30.8, 6.6 Hz, 4H), 1.37 – 1.26 (m, 6H). $^{13}$C NMR (101 MHz, DMSO) δ 159.20, 141.69, 140.11, 129.99, 128.89, 127.55, 126.80, 118.89, 113.55, 112.67, 96.95, 92.35, 85.15, 81.89, 76.75, 74.60, 72.12, 71.97, 71.93, 69.89, 69.73, 67.46, 61.07, 30.00, 29.96, 29.05, 28.88, 28.79, 25.59.

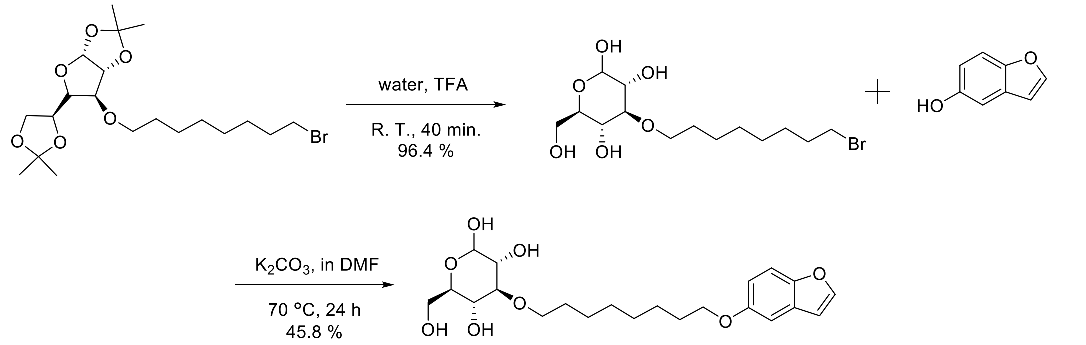

**Synthesis of (3R,4S,5R,6R)-4-((8-bromooctyl)oxy)-6-(hydroxymethyl)tetrahydro-2H-pyran-2,3,5-triol (11)**

Compound **9** (3.5 g, 7.78 mmol) was dissolved in 14.4 ml trifluoroacetic acid in 50-ml round-bottomed flask, and water (1.6 ml) were then added. The mixture was stirred at room temperature for 40 minutes. The progress of the reaction was monitored by TLC. Upon completion of the reaction, organic solvents were directly evaporated under vacuum. The resulting residue was purified by flash column chromatography on silica gel (10 % methanol in dichloromethane) to yield 2.6 g (91.5%) (3R,4S,5R,6R)-4-((8-bromooctyl)oxy)-6-(hydroxymethyl)tetrahydro-2H-pyran-2,3,5-triol (**11**) as white solid. $^1$H NMR (400 MHz, DMSO-$d_6$) δ 6.71 – 6.22 (m, 1H), 4.99 – 4.71 (m, 2H), 4.59 – 4.41 (m, 1H), 4.41 – 4.20 (m, 1H), 3.66 – 2.92 (m, 10H), 1.78 (p, *J* = 7.1 Hz, 2H), 1.48 (q, *J* = 6.9 Hz, 2H), 1.41 – 1.20 (m, 8H). $^{13}$C NMR (101 MHz, DMSO) δ 96.93, 92.34, 85.17, 81.90, 76.72, 74.59, 72.09, 71.92, 71.88, 69.92, 69.76, 61.12, 35.23, 32.29, 29.94, 29.90, 28.88, 28.18, 27.55, 25.53, 25.50.

**Synthesis of (3*R*,4*S*,5*R*,6*R*)-4-((8-(benzofuran-6-yloxy)octyl)oxy)-6-(hydroxymethyl)tetrahydro-2*H*-pyran-2,3,5-triol (2f)**

Compound **11** (185 mg, 1.0 mmol) was dissolved in 10 ml anhydrous DMF in a 100-ml round-bottomed flask. Benzofuran-5-ol (134 mg, 1.0 mmol) and potassium carbonate (184 mg, 1.33 mmol) were then added into the solution. The mixture was stirred at 50℃ for 12 h. The progress of the reaction was monitored by TLC. Upon completion of the reaction, the mixture was cooled to room temperature and organic solvents were removed under vacuum. The resulting precipitant was extracted with ethyl acetate, dried over anhydrous sodium sulfate, filtered, and concentrated under vacuum. The crude product was purified using flash column chromatography on silica gel (10 % methanol in dichloromethane) to yield 194 mg (45.8%) (3*R*,4*S*,5*R*,6*R*)-4-((8-(benzofuran-6-yloxy)octyl)oxy)-6-(hydroxymethyl)tetrahydro-2*H*-pyran-2,3,5-triol (**2f**) as colorless solid. ESI-MS *m/z*: 447.1988 $[M+Na]^+$. $^1$H NMR (400 MHz, DMSO-$d_6$) δ 7.92 (t, *J* = 1.7 Hz, 1H), 7.46 (d, *J* = 8.9 Hz, 1H), 7.18 – 7.11 (m, 1H), 6.91 – 6.83 (m, 2H), 6.66 – 6.27 (m, 1H), 4.92 – 4.76 (m, 2H), 4.53 – 4.45 (m, 1H), 4.41 – 4.24 (m, 1H), 3.96 (t, *J* = 6.5 Hz, 2H), 3.69 – 2.91 (m, 8H), 1.72 (p, *J* = 6.8 Hz, 2H), 1.50 (p, *J* = 6.8 Hz, 2H), 1.41 (q, *J* = 7.1 Hz, 2H), 1.36 – 1.27 (m, 6H). $^{13}$C NMR (101 MHz, DMSO) δ 154.95, 149.11, 146.56, 127.82, 113.35, 111.64, 106.87, 104.40, 96.92, 92.33, 85.17, 81.90, 76.72, 74.58, 72.09, 71.93, 71.89, 69.92, 69.76, 68.05, 61.11, 29.96, 29.92, 29.00, 28.84, 28.80, 25.58, 25.54.

**Synthesis of (3*R*,4*S*,5*R*,6*R*)-4-((8-(benzo[*b*]thiophen-6-yloxy)octyl)oxy)-6-(hydroxymethyl)tetrahydro-2*H*-pyran-2,3,5-triol (2g)**

Compound **2g** was prepared in a similar manner described for **2f**. Yield: 52.6%; ESI-MS *m/z*: 463.1766 $[M+Na]^+$. $^1$H NMR (400 MHz, DMSO-$d_6$) δ 7.83 (d, *J* = 8.8 Hz, 1H), 7.72 (d, *J* = 5.4 Hz, 1H), 7.40 (d, *J* = 2.3 Hz, 1H), 7.35 (d, *J* = 5.4 Hz, 1H), 6.98 (dd, *J* = 8.7, 2.2 Hz, 1H), 6.66 – 6.27 (m, 1H), 4.95 – 4.74 (m, 2H), 4.54 – 4.44 (m, 1H), 4.41 – 4.23 (m, 1H), 4.00 (t, *J* = 6.5 Hz, 2H), 3.73 – 2.89 (m, 8H), 1.73 (p, *J* = 6.5 Hz, 2H), 1.45 (dq, *J* = 31.3, 6.8 Hz, 4H), 1.38 – 1.26 (m, 6H). $^{13}$C NMR (101 MHz, DMSO) δ 156.46, 140.66, 131.21, 128.31, 123.80, 123.16, 114.81, 106.51, 96.93, 92.34, 85.18, 81.90, 76.73, 74.59, 72.09, 71.93, 71.90, 69.92, 69.76, 67.69, 61.11, 29.97, 29.93, 29.01, 28.85, 28.74, 25.57, 25.55.

**Synthesis of (3*R*,4*S*,5*R*,6*R*)-6-(hydroxymethyl)-4-((8-(quinolin-6-yloxy)octyl)oxy)tetrahydro-2*H*-pyran-2,3,5-triol (2h)**

Compound **2h** was prepared in a similar manner described for **3a**. Yield: 96.8%; ESI-MS *m/z*: 436.2328 $[M+H]^+$ and 458.2145 $[M+Na]^+$. $^1$H NMR (400 MHz, DMSO-$d_6$) δ 8.72 (dd, *J* = 4.2, 1.7 Hz, 1H), 8.23 (dd, *J* = 8.3, 1.6 Hz, 1H), 7.90 (d, *J* = 9.1 Hz, 1H), 7.45 (dd, *J* = 8.3, 4.2 Hz,

1H), 7.42 – 7.30 (m, 2H), 6.83 – 6.14 (m, 1H), 5.03 – 4.74 (m, 2H), 4.62 – 4.45 (m, 1H), 4.44 – 4.22 (m, 1H), 4.08 (t, *J* = 6.5 Hz, 2H), 3.75 – 2.89 (m, 8H), 1.77 (p, *J* = 6.8 Hz, 2H), 1.47 (dp, *J* = 22.2, 6.8 Hz, 4H), 1.31 (d, *J* = 8.8 Hz, 6H). $^{13}$C NMR (101 MHz, DMSO) δ 156.58, 147.83, 143.72, 134.78, 130.32, 129.10, 122.24, 121.62, 106.33, 96.97, 92.37, 85.20, 81.94, 76.76, 74.61, 72.13, 71.98, 71.93, 69.94, 69.77, 67.89, 61.12, 29.99, 29.96, 29.03, 28.86, 28.63, 25.58.

**Synthesis of (3*R*,4*S*,5*R*,6*R*)-6-(hydroxymethyl)-4-((8-(quinolin-3-yloxy)octyl)oxy)tetrahydro-2*H*-pyran-2,3,5-triol (2i)**

Compound **2i** was prepared in a similar manner described for **3a**. Yield: 86.0%; ESI-MS *m/z*: 436.2323 $[M+H]^+$. $^1$H NMR (400 MHz, DMSO-$d_6$) δ 8.62 (d, *J* = 2.9 Hz, 1H), 8.00 – 7.91 (m, 1H), 7.91 – 7.84 (m, 1H), 7.75 (d, *J* = 2.9 Hz, 1H), 7.60 – 7.52 (m, 2H), 6.71 – 6.21 (m, 1H), 4.95 – 4.74 (m, 2H), 4.55 – 4.43 (m, 1H), 4.41 – 4.23 (m, 1H), 4.13 (t, *J* = 6.5 Hz, 2H), 3.74 – 2.90 (m, 8H), 1.80 (p, *J* = 6.8 Hz, 2H), 1.59 – 1.40 (m, 4H), 1.40 – 1.28 (m, 6H). $^{13}$C NMR (101 MHz, DMSO) δ 152.17, 144.26, 142.80, 128.69, 128.57, 127.00, 126.97, 126.49, 113.20, 96.92, 92.33, 85.17, 81.90, 76.72, 74.58, 72.08, 71.92, 71.88, 69.91, 69.75, 67.99, 61.10, 29.95, 29.92, 28.98, 28.80, 28.47, 25.56, 25.54, 25.48.

**Synthesis of (3*R*,4*S*,5*R*,6*R*)-6-(hydroxymethyl)-4-((8-(quinolin-7-yloxy)octyl)oxy)tetrahydro-2*H*-pyran-2,3,5-triol (2j)**

Compound **2j** was prepared in a similar manner described for **3a**. Yield: 87.6%; ESI-MS *m/z*: 436.2331 $[M+H]^+$. $^1$H NMR (400 MHz, DMSO-$d_6$) δ 8.80 (dd, *J* = 4.3, 1.8 Hz, 1H), 8.26 (dd, *J* = 8.2, 1.7 Hz, 1H), 7.87 (d, *J* = 9.0 Hz, 1H), 7.41 – 7.32 (m, 2H), 7.24 (dd, *J* = 8.9, 2.5 Hz, 1H), 6.68 – 6.28 (m, 1H), 5.01 – 4.78 (m, 2H), 4.57 – 4.46 (m, 1H), 4.43 – 4.22 (m, 1H), 4.12 (t, *J* = 6.5 Hz, 2H), 3.72 – 2.91 (m, 8H), 1.78 (p, *J* = 6.6 Hz, 2H), 1.48 (dp, *J* = 22.2, 7.2, 6.7 Hz, 4H), 1.40 – 1.28 (m, 6H). $^{13}$C NMR (101 MHz, DMSO) δ 159.54, 150.61, 149.46, 135.64, 129.21, 123.00, 119.53, 119.17, 107.78, 96.94, 85.19, 76.75, 74.60, 71.98, 69.75, 67.79, 61.10, 29.95, 29.02, 28.86, 28.59, 25.58, 25.57.

**Synthesis of (3*R*,4*S*,5*R*,6*R*)-4-((8-((5-aminonaphthalen-2-yl)oxy)octyl)oxy)-6-(hydroxymethyl)tetrahydro-2*H*-pyran-2,3,5-triol (2k)**

Compound **2k** was prepared in a similar manner as described for **3a**. Yield: 64.8%; ESI-MS *m/z*: 472.2299 $[M+Na]^+$. $^1$H NMR (400 MHz, DMSO-$d_6$) δ 7.95 (d, *J* = 9.2 Hz, 1H), 7.16 – 7.06 (m, 2H), 7.01 – 6.91 (m, 2H), 6.68 – 6.29 (m, 2H), 5.61 (s, 2H), 4.95 – 4.78 (m, 2H), 4.55 – 4.47 (m, 1H), 4.43 – 4.24 (m, 1H), 4.03 (t, *J* = 6.5 Hz, 2H), 3.73 – 2.90 (m, 8H), 1.75 (p, *J* = 6.7 Hz, 2H), 1.56 – 1.39 (m, 4H), 1.39 – 1.23 (m, 6H). $^{13}$C NMR (101 MHz, DMSO) δ 156.79, 145.23, 136.12, 127.77, 124.48, 118.35, 116.39, 114.97, 107.22, 106.12, 97.38, 92.79, 85.64,

82.36, 77.19, 75.04, 72.57, 72.54, 72.43, 72.39, 70.36, 70.19, 67.75, 61.55, 30.44, 30.41, 29.50, 29.36, 29.23, 26.09, 26.06, 26.03.

**Synthesis of 6-((8-(((3*R*,4*S*,5*R*,6*R*)-2,3,5-trihydroxy-6-(hydroxymethyl)tetrahydro-2*H*-pyran-4-yl)oxy)octyl)oxy)-2*H*-chromen-2-one (2l)**

Compound **2l** was prepared in a similar manner described for **3a**. Yield: 89.4%; ESI-MS *m/z*: 475.1937 $[M+Na]^+$. $^1$H NMR (400 MHz, DMSO-$d_6$) δ 8.00 (d, *J* = 9.6 Hz, 1H), 7.32 (d, *J* = 9.0 Hz, 1H), 7.27 (d, *J* = 3.0 Hz, 1H), 7.18 (dd, *J* = 9.0, 3.0 Hz, 1H), 6.71 – 6.28 (m, 2H), 4.93 – 4.78 (m, 2H), 4.55 – 4.47 (m, 1H), 4.41 – 4.25 (m, 1H), 3.98 (t, *J* = 6.5 Hz, 2H), 3.69 – 2.91 (m, 8H), 1.72 (p, *J* = 6.6 Hz, 2H), 1.48 (q, *J* = 6.5 Hz, 2H), 1.45 – 1.36 (m, 2H), 1.36 – 1.25 (m, 7H). $^{13}$C NMR (101 MHz, DMSO) δ 160.21, 155.04, 147.76, 144.17, 119.88, 119.24, 117.36, 116.55, 111.32, 96.95, 92.37, 85.20, 81.93, 76.75, 74.62, 72.13, 71.99, 71.94, 69.93, 69.77, 68.16, 61.11, 29.99, 29.96, 29.03, 28.84, 28.65, 25.60, 25.57, 25.52.

**Synthesis of (3*R*,4*S*,5*R*,6*R*)-6-(hydroxymethyl)-4-((8-(quinolin-4-yloxy)octyl)oxy)tetrahydro-2*H*-pyran-2,3,5-triol (2m)**

Compound **2m** was prepared in a similar manner described for **3a**. Yield: 78.2%; ESI-MS *m/z*: 436.2327 $[M+H]^+$. $^1$H NMR (400 MHz, DMSO-$d_6$) δ 8.71 (d, *J* = 5.2 Hz, 1H), 8.14 (dd, *J* = 8.4, 1.4 Hz, 1H), 7.94 (d, *J* = 8.3 Hz, 1H), 7.74 (ddd, *J* = 8.5, 6.8, 1.5 Hz, 1H), 7.57 (dd, *J* = 8.2, 6.8 Hz, 1H), 7.02 (d, *J* = 5.3 Hz, 1H), 6.77 – 6.25 (m, 1H), 5.06 – 4.20 (m, 6H), 3.81 – 2.89 (m, 8H), 1.86 (p, *J* = 6.6 Hz, 2H), 1.50 (p, *J* = 7.2 Hz, 4H), 1.42 – 1.26 (m, 6H). $^{13}$C NMR (101 MHz, DMSO) δ 161.34, 151.98, 148.90, 130.24, 128.89, 126.20, 121.96, 121.23, 119.24, 116.26, 102.01, 97.38, 92.79, 85.59, 82.34, 77.17, 75.03, 72.54, 72.40, 72.36, 70.34, 70.19, 68.85, 63.23, 61.51, 30.42, 30.38, 29.43, 29.25, 28.77, 26.01.

**Synthesis of (3*R*,4*S*,5*R*,6*R*)-4-((8-(anthracen-2-yloxy)octyl)oxy)-6-(hydroxymethyl)tetrahydro-2*H*-pyran-2,3,5-triol (2n)**

Compound **2n** was prepared in a similar manner described for **3a**. Yield: 85.3%; ESI-MS *m/z*: 507.2361 $[M+Na]^+$. $^1$H NMR (400 MHz, DMSO-$d_6$) δ 8.48 (s, 1H), 8.37 (s, 1H), 8.09 – 7.92 (m, 3H), 7.45 (dddd, *J* = 18.8, 8.0, 6.5, 1.3 Hz, 2H), 7.36 (d, *J* = 2.4 Hz, 1H), 7.18 (dd, *J* = 9.1, 2.4 Hz, 1H), 6.70 – 6.28 (m, 1H), 4.99 – 4.77 (m, 2H), 4.51 (t, *J* = 5.9 Hz, 1H), 4.44 – 4.23 (m, 1H), 4.32 (s, 1H), 4.11 (t, *J* = 6.5 Hz, 2H), 3.75 – 2.90 (m, 8H), 1.80 (p, *J* = 6.8 Hz, 2H), 1.58 – 1.41 (m, 4H), 1.33 (q, *J* = 10.4, 6.5 Hz, 6H). $^{13}$C NMR (101 MHz, DMSO) δ 156.18, 132.47, 131.69, 129.77, 128.13, 127.75, 127.45, 126.02, 125.66, 124.51, 123.90, 120.72, 104.33, 96.95, 85.21, 76.76, 74.61, 71.99, 69.77, 67.54, 61.11, 29.97, 29.05, 28.91, 28.68, 25.66, 25.59.

**Synthesis of (3*R*,4*S*,5*R*,6*R*)-4-((8-((9*H*-carbazol-3-yl)oxy)octyl)oxy)-6-(hydroxymethyl)tetrahydro-2*H*-pyran-2,3,5-triol (2o)**

Compound **2o** was prepared in a similar manner described for **2f**. Yield: 42.5%; ESI-MS *m/z*: 496.2317 $[M+Na]^+$. $^1H$ NMR (400 MHz, DMSO-$d_6$) δ 11.24 (s, 1H), 8.14 (d, *J* = 7.8 Hz, 1H), 7.48 – 7.41 (m, 1H), 7.37 – 7.25 (m, 2H), 7.18 – 7.11 (m, 1H), 7.05 (d, *J* = 8.0 Hz, 1H), 6.68 (d, *J* = 8.0 Hz, 1H), 6.65 – 6.29 (m, 1H), 4.91 – 4.78 (m, 2H), 4.51 (t, *J* = 5.9 Hz, 1H), 4.42 – 4.25 (m, 1H), 4.19 (t, *J* = 6.3 Hz, 2H), 3.70 – 2.92 (m, 9H), 1.90 (p, *J* = 6.5 Hz, 2H), 1.64 – 1.46 (m, 4H), 1.45 – 1.30 (m, 6H). $^{13}C$ NMR (101 MHz, DMSO) δ 155.03, 141.09, 138.91, 126.54, 124.52, 122.10, 121.77, 118.63, 111.44, 110.43, 103.72, 100.40, 96.94, 85.21, 81.93, 76.76, 74.60, 72.13, 72.00, 69.76, 67.35, 61.11, 29.98, 29.08, 28.94, 28.88, 25.79, 25.58.

**Synthesis of (3*R*,4*S*,5*R*,6*R*)-6-(hydroxymethyl)-4-((8-(pyren-1-yloxy)octyl)oxy)tetrahydro-2*H*-pyran-2,3,5-triol (2p)**

Compound **2p** was prepared in a similar manner described for **2f**. Yield: 54.2%; ESI-MS *m/z*: 531.2360 $[M+Na]^+$. $^1H$ NMR (400 MHz, DMSO-$d_6$) δ 8.37 (d, *J* = 9.2 Hz, 1H), 8.24 (d, *J* = 8.5 Hz, 1H), 8.19 (ddd, *J* = 7.6, 6.2, 1.1 Hz, 2H), 8.13 (d, *J* = 9.2 Hz, 1H), 8.09 – 7.93 (m, 3H), 7.75 (d, *J* = 8.5 Hz, 1H), 6.71 – 6.27 (m, 1H), 5.00 – 4.77 (m, 2H), 4.55 – 4.48 (m, 1H), 4.41 – 4.25 (m, 3H), 3.74 – 2.92 (m, 8H), 1.93 (p, *J* = 6.5 Hz, 2H), 1.62 – 1.47 (m, 4H), 1.45 – 1.30 (m, 6H). $^{13}C$ NMR (101 MHz, DMSO) δ 153.17, 131.71, 131.58, 127.78, 126.87, 126.77, 126.50, 125.42, 125.11, 124.93, 124.80, 124.63, 124.59, 121.29, 119.81, 110.27, 97.39, 92.81, 85.66, 82.39, 77.21, 75.05, 72.57, 72.45, 72.40, 70.38, 70.22, 68.98, 61.56, 30.46, 30.42, 29.51, 29.36, 26.22, 26.08, 26.05.

**Synthesis of (3*R*,4*S*,5*R*,6*R*)-4-((8-((2-bromoquinolin-6-yl)oxy)octyl)oxy)-6-(hydroxymethyl)tetrahydro-2*H*-pyran-2,3,5-triol (3b)**

Compound **3b** was prepared in a similar manner described for **3a**. Yield: 93.2%; ESI-MS *m/z*: 536.1258 $[M+Na]^+$. $^1H$ NMR (400 MHz, DMSO-$d_6$) δ 8.20 (d, *J* = 8.6 Hz, 1H), 7.91 – 7.80 (m, 1H), 7.62 (d, *J* = 8.6 Hz, 1H), 7.44 (dd, *J* = 7.0, 2.9 Hz, 2H), 6.73 – 6.24 (m, 1H), 4.93 – 4.74 (m, 2H), 4.56 – 4.43 (m, 1H), 4.41 – 4.23 (m, 1H), 4.09 (t, *J* = 6.5 Hz, 2H), 3.72 – 2.91 (m, 8H), 1.78 (p, *J* = 6.7 Hz, 2H), 1.47 (dp, *J* = 22.6, 6.8 Hz, 4H), 1.38 – 1.27 (m, 6H). $^{13}C$ NMR (101 MHz, DMSO) δ 157.10, 143.79, 138.26, 138.24, 129.26, 128.19, 125.81, 123.29, 106.79, 96.92, 92.32, 85.17, 81.89, 76.71, 74.57, 72.08, 71.91, 71.87, 69.91, 69.74, 68.04, 61.09, 29.94, 29.91, 28.98, 28.79, 28.53, 25.56, 25.53, 25.50.

**Synthesis of (3*R*,4*S*,5*R*,6*R*)-6-(hydroxymethyl)-4-((8-((2-iodoquinolin-6-yl)oxy)octyl)oxy)tetrahydro-2*H*-pyran-2,3,5-triol (3c)**

Compound **3c** was prepared in a similar manner described for **3a**. Yield: 84.7%; ESI-MS *m/z*: 562.1279 $[M+H]^+$. $^1$H NMR (400 MHz, DMSO-$d_6$) δ 7.97 – 7.92 (m, 1H), 7.84 (d, *J* = 8.7 Hz, 1H), 7.79 (d, *J* = 8.5 Hz, 1H), 7.43 – 7.35 (m, 2H), 6.69 – 6.25 (m, 1H), 4.98 – 4.73 (m, 2H), 4.58 – 4.22 (m, 2H), 4.08 (t, *J* = 6.5 Hz, 2H), 3.73 – 2.90 (m, 8H), 1.77 (p, *J* = 6.7 Hz, 2H), 1.56 – 1.47 (m, 2H), 1.43 (q, *J* = 7.3, 6.8 Hz, 2H), 1.39 – 1.24 (m, 6H). $^{13}$C NMR (101 MHz, DMSO) δ 157.03, 145.02, 136.79, 131.90, 129.43, 128.26, 123.02, 116.19, 106.74, 96.92, 92.34, 85.18, 81.90, 76.73, 74.58, 72.09, 71.94, 71.90, 69.91, 69.74, 68.02, 61.09, 29.96, 29.92, 28.99, 28.82, 28.56, 25.57, 25.54, 25.53.

**Synthesis of (3*R*,4*S*,5*R*,6*R*)-4-((8-((3-bromoquinolin-6-yl)oxy)octyl)oxy)-6-(hydroxymethyl)tetrahydro-2*H*-pyran-2,3,5-triol (3d)**

Compound **3d** was prepared in a similar manner described for **3a**. Yield: 76.4%; ESI-MS *m/z*: 514.1438 $[M+H]^+$. $^1$H NMR (400 MHz, DMSO-$d_6$) δ 8.75 (d, *J* = 2.3 Hz, 1H), 8.55 (d, *J* = 2.2 Hz, 1H), 7.91 (d, *J* = 9.1 Hz, 1H), 7.43 (dd, *J* = 9.2, 2.7 Hz, 1H), 7.35 (d, *J* = 2.8 Hz, 1H), 6.72 – 6.27 (m, 1H), 5.00 – 4.75 (m, 2H), 4.54 – 4.47 (m, 1H), 4.33 (dt, *J* = 43.3, 6.2 Hz, 1H), 4.08 (t, *J* = 6.5 Hz, 2H), 3.74 – 2.86 (m, 8H), 1.78 (p, *J* = 6.7 Hz, 2H), 1.54 – 1.47 (m, 2H), 1.47 – 1.39 (m, 2H), 1.38 – 1.27 (m, 6H). $^{13}$C NMR (101 MHz, DMSO) δ 157.43, 148.11, 141.83, 136.01, 130.33, 130.31, 122.86, 117.18, 105.74, 96.93, 92.34, 85.18, 81.91, 76.73, 74.59, 72.10, 71.94, 71.90, 69.92, 69.75, 68.03, 61.10, 29.96, 29.93, 29.00, 28.84, 28.53, 25.58, 25.55, 25.53.

**Synthesis of (3*R*,4*S*,5*R*,6*R*)-6-(hydroxymethyl)-4-((8-((2-methylquinolin-6-yl)oxy)octyl)oxy)tetrahydro-2*H*-pyran-2,3,5-triol (3e)**

Compound **3e** was prepared in a similar manner described for **3a**. Yield: 91.2%; ESI-MS *m/z*: 450.2491 $[M+H]^+$. $^1$H NMR (400 MHz, DMSO-$d_6$) δ 8.11 (d, *J* = 8.4 Hz, 1H), 7.80 (d, *J* = 9.0 Hz, 1H), 7.32 (ddd, *J* = 15.0, 7.2, 3.4 Hz, 3H), 6.79 – 6.23 (m, 1H), 5.00 – 4.73 (m, 2H), 4.59 – 4.44 (m, 1H), 4.42 – 4.19 (m, 1H), 4.07 (t, *J* = 6.5 Hz, 2H), 3.77 – 2.89 (m, 8H), 2.60 (s, 3H), 1.77 (p, *J* = 6.7 Hz, 2H), 1.47 (dp, *J* = 22.8, 7.5, 6.8 Hz, 4H), 1.40 – 1.25 (m, 6H). $^{13}$C NMR (101 MHz, DMSO) δ 155.96, 155.79, 143.16, 134.96, 129.54, 127.13, 122.21, 121.77, 106.33, 96.94, 92.33, 85.17, 81.90, 76.72, 74.57, 72.09, 71.92, 71.88, 69.91, 69.74, 67.78, 61.09, 29.96, 29.92, 28.99, 28.83, 28.64, 25.56, 24.51.

**Synthesis of (3*R*,4*S*,5*R*,6*R*)-6-(hydroxymethyl)-4-((8-((2-(trifluoromethyl)quinolin-6-yl)oxy)octyl)oxy)tetrahydro-2*H*-pyran-2,3,5-triol (3f)**

Compound **3f** was prepared in a similar manner described for **3a**. Yield: 68.7%; ESI-MS *m/z*: 504.2196 $[M+H]^+$. $^1$H NMR (400 MHz, DMSO-$d_6$) δ 8.54 (d, *J* = 8.6 Hz, 1H), 8.05 (d, *J* = 9.2 Hz, 1H), 7.89 (dd, *J* = 8.6, 1.3 Hz, 1H), 7.54 (d, *J* = 8.4 Hz, 2H), 6.71 – 6.21 (m, 1H), 4.95 –

4.75 (m, 2H), 4.55 – 4.44 (m, 1H), 4.41 – 4.22 (m, 1H), 4.14 (t, $J$ = 6.6 Hz, 2H), 3.73 – 2.91 (m, 8H), 1.80 (p, $J$ = 6.8 Hz, 2H), 1.47 (dq, $J$ = 21.4, 7.0 Hz, 4H), 1.40 – 1.26 (m, 6H). $^{13}$C NMR (101 MHz, DMSO) δ 158.33, 144.02, 143.69, 142.39, 137.47, 130.74, 130.38, 124.41, 123.19, 117.17, 117.16, 106.18, 96.93, 92.34, 85.18, 81.91, 76.72, 74.59, 72.09, 71.93, 71.89, 69.92, 69.76, 68.20, 61.10, 29.96, 29.92, 28.99, 28.81, 28.49, 25.57, 25.54, 25.51.

**Synthesis of (3*R*,4*S*,5*R*,6*R*)-6-(hydroxymethyl)-4-((8-((2-hydroxyquinolin-6-yl)oxy)octyl)oxy)tetrahydro-2*H*-pyran-2,3,5-triol (3g)**

Compound **3g** was prepared in a similar manner described for **3a**. Yield: 43.6%; ESI-MS *m/z*: 452.2281 [M+H]$^+$. $^1$H NMR (400 MHz, DMSO-$d_6$) δ 11.62 (s, 1H), 7.83 (d, $J$ = 9.5 Hz, 1H), 7.28 – 7.17 (m, 2H), 7.13 (dd, $J$ = 8.9, 2.7 Hz, 1H), 6.69 – 6.26 (m, 2H), 4.95 – 4.75 (m, 2H), 4.57 – 4.45 (m, 1H), 4.33 (dt, $J$ = 44.4, 6.1 Hz, 1H), 3.96 (t, $J$ = 6.5 Hz, 2H), 3.73 – 2.86 (m, 8H), 1.71 (p, $J$ = 6.7 Hz, 2H), 1.49 (q, $J$ = 6.1 Hz, 2H), 1.44 – 1.36 (m, 2H), 1.36 – 1.23 (m, 6H). $^{13}$C NMR (101 MHz, DMSO) δ 161.50, 153.47, 139.81, 133.24, 122.24, 119.90, 119.69, 116.33, 110.04, 96.93, 92.34, 85.18, 81.91, 76.73, 74.59, 72.10, 71.95, 71.91, 69.91, 69.75, 67.86, 61.10, 29.97, 29.93, 29.02, 28.84, 28.72, 25.58, 25.55.

**Synthesis of (3*R*,4*S*,5*R*,6*R*)-4-((8-((2-(dimethylamino)quinolin-6-yl)oxy)octyl)oxy)-6-(hydroxymethyl)tetrahydro-2*H*-pyran-2,3,5-triol (3h)**

Compound **3h** was prepared in a similar manner described for **3a**. Yield: 34.2%; ESI-MS *m/z*: 475.1935 [M+Na]$^+$. $^1$H NMR (400 MHz, DMSO-$d_6$) δ 7.92 (d, $J$ = 9.3 Hz, 1H), 7.47 (d, $J$ = 8.7 Hz, 1H), 7.15 (d, $J$ = 10.1 Hz, 2H), 7.04 (dd, $J$ = 9.1, 1.3 Hz, 1H), 6.67 – 6.26 (m, 1H), 4.95 – 4.74 (m, 2H), 4.55 – 4.43 (m, 1H), 4.42 – 4.21 (m, 1H), 3.99 (t, $J$ = 6.5 Hz, 2H), 3.72 – 2.91 (m, 14H), 1.74 (p, $J$ = 6.8 Hz, 2H), 1.47 (dh, $J$ = 29.9, 6.6 Hz, 4H), 1.38 – 1.24 (m, 6H). $^{13}$C NMR (101 MHz, DMSO) δ 156.27, 153.27, 142.86, 136.22, 127.13, 122.48, 120.95, 109.68, 107.21, 96.92, 92.33, 85.17, 81.90, 76.72, 74.58, 72.09, 71.93, 71.89, 69.92, 69.75, 67.64, 61.10, 37.76, 29.97, 29.93, 29.01, 28.86, 28.78, 25.60, 25.55.

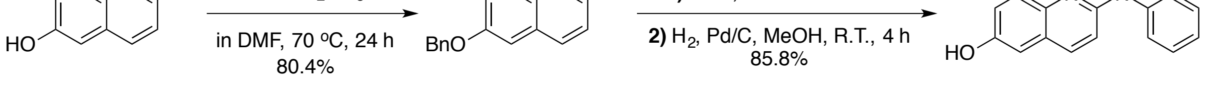

**Synthesis of 6-(benzyloxy)-2-chloroquinoline (12)**

2-chloroquinolin-6-ol (1013 mg, 5.64 mmol) was dissolved in 10 ml anhydrous DMF in a 50-ml round-bottomed flask. Benzyl bromide (1.34 ml, 11.28 mmol) and potassium carbonate (1.17 g, 8.47 mmol) were then added into the solution. The mixture was stirred at 70℃ for 24 h. The progress of the reaction was monitored by TLC. Upon completion of the reaction, the mixture was cooled to room temperature and organic solvents were removed under vacuum.

The resulting precipitant was extracted with ethyl acetate, dried over anhydrous sodium sulfate, filtered, and concentrated under vacuum. The crude product was purified using flash column chromatography on silica gel (9 % ethyl acetate in petroleum ether) to yield 1.40 g (91.6%) 6-(benzyloxy)-2-chloroquinoline (**12**) as white solid. $^{1}$H NMR (400 MHz, Chloroform-*d*) δ 7.96 (dd, *J* = 15.9, 8.9 Hz, 2H), 7.51 – 7.31 (m, 7H), 7.15 (d, *J* = 2.8 Hz, 1H), 5.18 (s, 2H). $^{13}$C NMR (101 MHz, $CDCl_3$) δ 157.20, 148.20, 143.89, 137.67, 136.26, 130.06, 128.72, 128.25, 127.86, 127.52, 123.36, 122.53, 106.71, 70.41.

**Synthesis of 2-(phenylamino)quinolin-6-ol (13)**

Compound **12** (200 mg, 0.74 mmol) was dissolved in 3 ml NMP in a 20-ml microwave reaction vial, and aniline (345 mg, 3.70 mmol) was then added. The mixture was reacted in a microwave oven at 150 $^{o}$C for 30 minutes. The temperature was maintained by modulating the power level. The progress of the reaction was monitored by TLC. Upon completion of the reaction, the mixture was cooled to room temperature and the portion of solute was precipitated in water. The resulting residue was extracted with ethyl acetate, dried over anhydrous sodium sulfate, filtered, and concentrated under vacuum. The crude product was purified by flash column chromatography on silica gel (16 % ethyl acetate in petroleum ether) to afford 6-(benzyloxy)-*N*-phenylquinolin-2-amine as white solid. Then 6-(benzyloxy)-*N*-phenylquinolin-2-amine was dissolved in 6 ml methanol and 2 ml ethyl acetate in a 25-ml round bottomed flask. Palladium (5 %) on carbon (120 mg, 0.05 mmol) was then added, and the flask was evacuated carefully and flushed with hydrogen gas for three times using a balloon. The mixture was stirred at room temperature for 1 h under a hydrogen atmosphere (using a balloon). Upon completion of the reaction, the Pd/C was filtered off using a pad of celite and the filtrate was concentrated. The crude product was purified using flash column chromatography on silica gel (50 % ethyl acetate in petroleum ether) to yield 150 g (85.8%) 2-(phenylamino)quinolin-6-ol (**13**) as faint yellow oil. $^{1}$H NMR (400 MHz, Chloroform-*d*) δ 7.82 (d, *J* = 9.0 Hz, 1H), 7.72 (d, *J* = 9.0 Hz, 1H), 7.56 – 7.50 (m, 2H), 7.41 – 7.35 (m, 2H), 7.23 (dd, *J* = 9.0, 2.8 Hz, 1H), 7.09 (tt, *J* = 7.3, 1.1 Hz, 1H), 7.04 (d, *J* = 2.8 Hz, 1H), 7.02 (d, *J* = 8.9 Hz, 1H).

**Synthesis of (3*R*,4*S*,5*R*,6*R*)-6-(hydroxymethyl)-4-((8-((2-(phenylamino)quinolin-6-yl)oxy)octyl)oxy)tetrahydro-2*H*-pyran-2,3,5-triol (3m)**

Compound **3m** was synthesized starting from compound **13** and compound **9** following a similar manner described for **3a**. Yield: 78.6%; ESI-MS *m/z*: 527.2756 $[M+H]^{+}$. $^{1}$H NMR (400 MHz, DMSO-$d_6$) δ 9.23 (s, 1H), 8.07 – 7.87 (m, 3H), 7.60 (d, *J* = 8.9 Hz, 1H), 7.38 – 7.26 (m, 2H), 7.26 – 7.15 (m, 2H), 7.03 (d, *J* = 8.9 Hz, 1H), 6.91 (tt, *J* = 7.3, 1.2 Hz, 1H), 6.72 – 6.26 (m, 1H), 4.97 – 4.74 (m, 2H), 4.55 – 4.43 (m, 1H), 4.39 – 4.25 (m, 1H), 4.03 (t, *J* = 6.5 Hz, 2H),

3.75 – 2.88 (m, 8H), 1.76 (p, *J* = 6.7 Hz, 2H), 1.51 (p, *J* = 6.5 Hz, 2H), 1.47 – 1.39 (m, 2H), 1.39 – 1.24 (m, 6H). $^{13}$C NMR (101 MHz, DMSO) δ 154.19, 152.76, 142.06, 141.73, 136.05, 128.57, 127.61, 124.04, 120.82, 120.48, 117.91, 114.21, 107.56, 96.92, 92.33, 85.18, 81.90, 76.72, 74.58, 72.09, 71.93, 71.89, 69.92, 69.75, 67.72, 61.11, 29.96, 29.93, 29.01, 28.85, 28.75, 25.58, 25.55.

**Synthesis of (3*R*,4*S*,5*R*,6*R*)-6-(hydroxymethyl)-4-((8-((2-phenoxyquinolin-6-yl)oxy)octyl)oxy)tetrahydro-2*H*-pyran-2,3,5-triol (3l)**

Compound **3l** was prepared in a similar manner as described for **3m**. Yield: 60.2%; ESI-MS *m/z*: 528.2602 [M+H]$^+$. $^1$H NMR (400 MHz, DMSO-$d_6$) δ 8.29 (d, *J* = 8.8 Hz, 1H), 7.54 (d, *J* = 9.1 Hz, 1H), 7.48 – 7.41 (m, 2H), 7.38 (d, *J* = 2.8 Hz, 1H), 7.28 (dd, *J* = 9.1, 2.8 Hz, 1H), 7.26 – 7.16 (m, 4H), 6.70 – 6.25 (m, 1H), 4.98 – 4.75 (m, 2H), 4.57 – 4.47 (m, 1H), 4.43 – 4.23 (m, 1H), 4.05 (t, *J* = 6.5 Hz, 2H), 3.70 – 2.89 (m, 8H), 1.76 (p, *J* = 6.7 Hz, 2H), 1.55 – 1.39 (m, 4H), 1.37 – 1.25 (m, 6H). $^{13}$C NMR (101 MHz, DMSO) δ 159.79, 155.61, 153.72, 140.91, 139.43, 129.66, 128.32, 126.42, 124.57, 121.95, 121.36, 113.05, 107.19, 96.94, 92.35, 85.19, 81.92, 76.75, 74.60, 72.11, 71.97, 71.92, 69.92, 69.76, 67.85, 61.10, 29.98, 29.95, 29.03, 28.86, 28.66, 25.59, 25.57.

**Synthesis of (3*R*,4*S*,5*R*,6*R*)-6-(hydroxymethyl)-4-((8-((2-((3,4,5-trimethoxyphenyl)amino)quinolin-6-yl)oxy)octyl)oxy)tetrahydro-2*H*-pyran-2,3,5-triol (3n)**

Compound **3n** was prepared in a similar manner as described for **3m**. Yield: 69.0%; ESI-MS *m/z*: 617.3066 [M+H]$^+$. $^1$H NMR (400 MHz, DMSO-$d_6$) δ 9.22 (s, 1H), 7.95 (d, *J* = 8.9 Hz, 1H), 7.59 (d, *J* = 8.9 Hz, 1H), 7.42 (s, 2H), 7.22 (dd, *J* = 8.9, 2.8 Hz, 1H), 7.18 (d, *J* = 2.8 Hz, 1H), 6.98 (d, *J* = 8.9 Hz, 1H), 6.78 – 6.22 (m, 1H), 4.99 – 4.74 (m, 2H), 4.50 (dd, *J* = 6.8, 4.2 Hz, 1H), 4.41 – 4.23 (m, 1H), 4.02 (t, *J* = 6.5 Hz, 2H), 3.82 (s, 6H), 3.73 – 2.87 (m, 11H), 1.82 – 1.69 (m, 2H), 1.57 – 1.48 (m, 2H), 1.48 – 1.39 (m, 2H), 1.39 – 1.26 (m, 6H). $^{13}$C NMR (101 MHz, DMSO) δ 154.20, 152.81, 152.77, 152.66, 141.91, 137.94, 135.96, 131.41, 127.72, 123.94, 120.89, 114.22, 107.47, 96.95, 95.62, 92.33, 85.17, 81.90, 76.72, 74.57, 72.09, 71.92, 71.88, 69.91, 69.74, 67.72, 61.09, 60.13, 55.55, 29.96, 29.93, 29.02, 29.01, 28.86, 28.75, 25.59, 25.55.

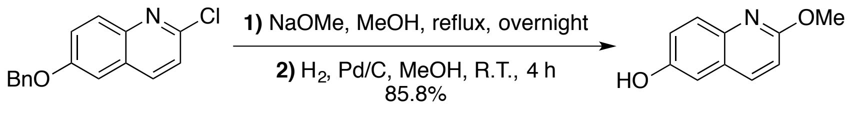

**Synthesis of 2-methoxyquinolin-6-ol (14)**

Compound **12** (200 mg, 0.74 mmol) was dissolved in 10 ml anhydrous methanol in a 50-ml round-bottomed flask, and sodium methoxide (1.2 g, 22.2 mmol) was then added. The mixture was stirred at 70 °C for 24 h. The progress of the reaction was monitored by TLC. Upon completion of the reaction, the mixture was cooled to room temperature and the portion of solvent was removed under vacuum. The crude product was purified by flash column chromatography on silica gel (9 % ethyl acetate in petroleum ether) to yield 180 mg (92 %) 6-(benzyloxy)-2-methoxyquinoline as white solid. Then 6-(benzyloxy)-2-methoxyquinoline (180 mg, 0.62 mmol) was dissolved in 6 ml methanol and 2 ml ethyl acetate in a 25-ml round bottomed flask. Palladium (5 %) on carbon (200 mg, 0.09 mmol) was then added, and the flask was evacuated carefully and flushed with hydrogen gas for three times using a balloon. The mixture was stirred at room temperature for 1 h under a hydrogen atmosphere (using a balloon). Upon completion of the reaction, the Pd/C was filtered off using a pad of celite and the filtrate was concentrated. The resulting residue was used in the next step without further purification.

**Synthesis of (3*R*,4*S*,5*R*,6*R*)-6-(hydroxymethyl)-4-((8-((2-methoxyquinolin-6-yl)oxy)octyl)oxy)tetrahydro-2*H*-pyran-2,3,5-triol (3i)**

Compound **3i** was synthesized starting from compound **14** and compound **9** following a similar manner described for **3a**. Yield: 88.0%; ESI-MS *m/z*: 466.2432 $[M+H]^+$. $^1$H NMR (400 MHz, DMSO-$d_6$) δ 8.12 (dd, *J* = 9.0, 0.7 Hz, 1H), 7.67 (d, *J* = 8.9 Hz, 1H), 7.32 (d, *J* = 2.8 Hz, 1H), 7.29 (dd, *J* = 8.9, 2.8 Hz, 1H), 6.97 (d, *J* = 8.8 Hz, 1H), 6.66 – 6.25 (m, 1H), 4.93 – 4.74 (m, 2H), 4.54 – 4.44 (m, 1H), 4.39 – 4.24 (m, 1H), 4.04 (t, *J* = 6.5 Hz, 2H), 3.94 (s, 3H), 3.73 – 2.88 (m, 8H), 1.84 – 1.70 (m, 2H), 1.50 (t, *J* = 6.8 Hz, 2H), 1.47 – 1.39 (m, 2H), 1.39 – 1.25 (m, 6H). $^{13}$C NMR (101 MHz, DMSO) δ 160.48, 155.02, 141.04, 138.32, 127.94, 125.54, 121.28, 112.90, 107.49, 96.92, 92.33, 85.17, 81.90, 76.72, 74.58, 72.08, 71.92, 71.88, 69.91, 69.75, 67.77, 61.09, 52.92, 29.95, 29.92, 28.99, 28.83, 28.68, 25.56, 25.54.

**Synthesis of (3*R*,4*S*,5*R*,6*R*)-4-((8-((2-ethoxyquinolin-6-yl)oxy)octyl)oxy)-6-(hydroxymethyl)tetrahydro-2*H*-pyran-2,3,5-triol (3j)**

Compound **3j** was prepared in a similar manner described for **3i**. Yield: 84.0%; ESI-MS *m/z*: 480.2594 $[M+H]^+$. $^1$H NMR (400 MHz, DMSO-$d_6$) δ 8.11 (d, *J* = 8.9 Hz, 1H), 7.64 (d, *J* = 8.9 Hz, 1H), 7.37 – 7.21 (m, 2H), 6.94 (d, *J* = 8.8 Hz, 1H), 6.68 – 6.28 (m, 1H), 4.91 – 4.79 (m, 2H), 4.56 – 4.25 (m, 4H), 4.03 (t, *J* = 6.5 Hz, 2H), 3.67 – 2.93 (m, 8H), 1.75 (p, *J* = 6.6 Hz, 2H), 1.55 – 1.39 (m, 4H), 1.38 – 1.26 (m, 9H). $^{13}$C NMR (101 MHz, DMSO) δ 160.18, 154.98, 141.13, 138.30, 127.96, 125.48, 121.27, 113.06, 107.44, 96.95, 92.36, 85.19, 81.92, 76.75, 74.61, 72.12, 71.98, 71.93, 69.93, 69.76, 67.78, 61.11, 61.01, 29.99, 29.96, 29.04, 28.88, 28.72, 25.60, 14.52.

**Synthesis of (3*R*,4*S*,5*R*,6*R*)-6-(hydroxymethyl)-4-((8-((2-isopropoxyquinolin-6-yl)oxy)octyl)oxy)tetrahydro-2*H*-pyran-2,3,5-triol (3k)**

Compound **3k** was prepared in a similar manner as described for **3i**. Yield: 65.0%; ESI-MS *m/z*: 494.2742 $[M+H]^+$. $^1$H NMR (400 MHz, DMSO-$d_6$) δ 8.09 (d, *J* = 8.9 Hz, 1H), 7.63 (d, *J* = 8.9 Hz, 1H), 7.34 – 7.21 (m, 2H), 6.88 (d, *J* = 8.8 Hz, 1H), 6.68 – 6.28 (m, 1H), 5.50 – 5.34 (m, 1H), 4.96 – 4.75 (m, 2H), 4.57 – 4.44 (m, 1H), 4.42 – 4.23 (m, 1H), 4.03 (t, *J* = 6.5 Hz, 2H), 3.71 – 2.91 (m, 9H), 1.75 (p, *J* = 6.5 Hz, 2H), 1.54 – 1.38 (m, 4H), 1.38 – 1.26 (m, 12H). $^{13}$C NMR (101 MHz, DMSO) δ 159.69, 154.92, 141.17, 138.28, 127.93, 125.32, 121.22, 113.51, 107.42, 96.94, 92.36, 85.19, 81.92, 76.75, 74.60, 72.11, 71.97, 71.93, 69.93, 69.76, 67.77, 67.14, 61.11, 29.99, 29.95, 29.03, 28.87, 28.72, 25.59, 21.92.

**Synthesis of (3*R*,4*S*,5*R*,6*R*)-4-((8-((7-chloroquinolin-4-yl)oxy)octyl)oxy)-6-(hydroxymethyl)tetrahydro-2*H*-pyran-2,3,5-triol (4a)**

Compound **4a** was prepared in a similar manner described for **3a**. Yield: 71.2%; ESI-MS *m/z*: 470.1939 $[M+H]^+$. $^1$H NMR (400 MHz, DMSO-$d_6$) δ 8.74 (d, *J* = 5.2 Hz, 1H), 8.15 (d, *J* = 8.9 Hz, 1H), 7.98 (d, *J* = 2.1 Hz, 1H), 7.59 (dd, *J* = 8.9, 2.2 Hz, 1H), 7.05 (d, *J* = 5.3 Hz, 1H), 6.72 – 6.25 (m, 1H), 4.95 – 4.75 (m, 2H), 4.52 – 4.34 (m, 1H), 4.26 (dt, *J* = 9.7, 6.5 Hz, 3H), 3.73 – 2.90 (m, 8H), 1.90 – 1.77 (m, 2H), 1.55 – 1.42 (m, 4H), 1.42 – 1.26 (m, 6H). $^{13}$C NMR (101 MHz, DMSO) δ 160.86, 153.11, 149.16, 134.34, 127.27, 126.24, 123.75, 119.43, 102.10, 96.93, 85.18, 81.90, 76.72, 74.57, 72.08, 71.92, 71.87, 69.92, 69.75, 68.60, 61.09, 29.94, 29.90, 28.94, 28.76, 28.26, 25.55, 25.52, 25.50.

**Synthesis of (3*R*,4*S*,5*R*,6*R*)-4-((8-((7-bromoquinolin-4-yl)oxy)octyl)oxy)-6-(hydroxymethyl)tetrahydro-2*H*-pyran-2,3,5-triol (4b)**

Compound **4b** was prepared in a similar manner described for **3a**. Yield: 69.6%; ESI-MS *m/z*: 514.1440 $[M+H]^+$. $^1$H NMR (400 MHz, DMSO-$d_6$) δ 8.73 (d, *J* = 5.2 Hz, 1H), 8.14 (d, *J* = 2.0 Hz, 1H), 8.07 (d, *J* = 8.9 Hz, 1H), 7.70 (dd, *J* = 8.9, 2.0 Hz, 1H), 7.06 (d, *J* = 5.3 Hz, 1H), 6.74 – 6.24 (m, 1H), 4.98 – 4.74 (m, 2H), 4.58 – 4.16 (m, 4H), 3.75 – 2.89 (m, 8H), 1.92 – 1.78 (m, 2H), 1.49 (qt, *J* = 7.5, 4.7 Hz, 4H), 1.43 – 1.26 (m, 6H). $^{13}$C NMR (101 MHz, DMSO) δ 160.92, 153.02, 149.41, 130.50, 128.77, 123.77, 123.06, 119.67, 102.17, 96.92, 92.32, 85.18, 81.90, 76.71, 74.57, 72.08, 71.91, 71.87, 69.92, 69.75, 68.61, 61.09, 29.94, 29.90, 28.94, 28.76, 28.25, 25.55, 25.52, 25.50.

**Synthesis of (3*R*,4*S*,5*R*,6*R*)-6-(hydroxymethyl)-4-((8-((7-iodoquinolin-4-yl)oxy)octyl)oxy)tetrahydro-2*H*-pyran-2,3,5-triol (4c)**

Compound **4c** was prepared in a similar manner described for **3a**. Yield: 29.4%; ESI-MS *m/z*: 562.1308 $[M+H]^+$. $^1H$ NMR (400 MHz, DMSO-$d_6$) δ 8.70 (d, *J* = 5.2 Hz, 1H), 8.34 (d, *J* = 1.6 Hz, 1H), 7.90 (d, *J* = 8.7 Hz, 1H), 7.84 (dd, *J* = 8.7, 1.7 Hz, 1H), 7.05 (d, *J* = 5.3 Hz, 1H), 6.71 – 6.27 (m, 1H), 4.96 – 4.76 (m, 2H), 4.58 – 4.16 (m, 4H), 3.73 – 2.89 (m, 8H), 1.85 (p, *J* = 6.6 Hz, 2H), 1.48 (p, *J* = 7.0 Hz, 4H), 1.40 – 1.26 (m, 6H). $^{13}C$ NMR (101 MHz, DMSO) δ 160.94, 152.63, 149.51, 136.95, 134.04, 123.35, 119.95, 102.19, 96.93, 96.72, 92.33, 85.18, 81.91, 76.73, 74.58, 72.10, 71.95, 71.90, 69.91, 69.74, 68.57, 61.09, 29.96, 29.92, 28.97, 28.78, 28.28, 25.57, 25.54, 25.52.

**Synthesis of (3*R*,4*S*,5*R*,6*R*)-4-((8-((6-bromoquinolin-4-yl)oxy)octyl)oxy)-6-(hydroxymethyl)tetrahydro-2*H*-pyran-2,3,5-triol (4d)**

Compound **4d** was prepared in a similar manner described for **3a**. Yield: 82.4%; ESI-MS *m/z*: 514.1441 $[M+H]^+$. $^1H$ NMR (400 MHz, DMSO-$d_6$) δ 8.74 (d, *J* = 5.2 Hz, 1H), 8.24 (d, *J* = 2.1 Hz, 1H), 7.92 – 7.81 (m, 2H), 7.07 (d, *J* = 5.3 Hz, 1H), 6.79 – 6.23 (m, 1H), 5.01 – 4.73 (m, 2H), 4.60 – 4.14 (m, 4H), 3.75 – 2.84 (m, 8H), 1.86 (p, *J* = 6.7 Hz, 2H), 1.48 (p, *J* = 6.8 Hz, 4H), 1.42 – 1.24 (m, 6H). $^{13}C$ NMR (101 MHz, DMSO) δ 159.83, 152.32, 147.27, 132.77, 131.06, 123.58, 122.05, 118.67, 102.41, 96.94, 92.34, 85.18, 81.91, 76.74, 74.58, 72.10, 71.95, 71.90, 69.91, 69.74, 68.73, 61.10, 29.98, 29.94, 28.97, 28.80, 28.19, 25.60, 25.57, 25.53.

**Synthesis of (3*R*,4*S*,5*R*,6*R*)-4-((8-((8-bromoquinolin-4-yl)oxy)octyl)oxy)-6-(hydroxymethyl)tetrahydro-2*H*-pyran-2,3,5-triol (4e)**

Compound **4e** was prepared in a similar manner described for **3a**. Yield: 78.0%; ESI-MS *m/z*: 514.1446 $[M+H]^+$. $^1H$ NMR (400 MHz, DMSO-$d_6$) δ 8.82 (d, *J* = 5.2 Hz, 1H), 8.17 (dd, *J* = 8.3, 1.3 Hz, 1H), 8.12 (dd, *J* = 7.5, 1.4 Hz, 1H), 7.47 (dd, *J* = 8.3, 7.5 Hz, 1H), 7.13 (d, *J* = 5.3 Hz, 1H), 6.69 – 6.27 (m, 1H), 4.94 – 4.76 (m, 2H), 4.55 – 4.17 (m, 4H), 3.72 – 2.87 (m, 8H), 1.93 – 1.78 (m, 2H), 1.50 (p, *J* = 8.5, 7.5 Hz, 4H), 1.42 – 1.24 (m, 6H). $^{13}C$ NMR (101 MHz, DMSO) δ 161.06, 152.54, 145.34, 133.43, 126.39, 123.96, 122.20, 121.66, 102.46, 96.93, 92.34, 85.18, 81.91, 76.73, 74.58, 72.10, 71.94, 71.90, 69.91, 69.74, 68.85, 61.09, 29.96, 29.93, 28.97, 28.78, 28.28, 25.58, 25.54, 25.53.

**Synthesis of (3*R*,4*S*,5*R*,6*R*)-6-(hydroxymethyl)-4-((8-((7-methoxyquinolin-4-yl)oxy)octyl)oxy)tetrahydro-2*H*-pyran-2,3,5-triol (4f)**

Compound **4f** was prepared in a similar manner described for **3a**. Yield: 89.2%; ESI-MS *m/z*: 466.2442 $[M+H]^+$. $^1H$ NMR (400 MHz, DMSO-$d_6$) δ 8.62 (d, *J* = 5.3 Hz, 1H), 8.02 (d, *J* = 9.1 Hz, 1H), 7.31 (d, *J* = 2.5 Hz, 1H), 7.18 (dd, *J* = 9.1, 2.6 Hz, 1H), 6.87 (d, *J* = 5.3 Hz, 1H), 6.72 – 6.21 (m, 1H), 4.97 – 4.73 (m, 2H), 4.57 – 4.12 (m, 4H), 3.90 (s, 3H), 3.74 – 2.89 (m, 8H),

1.85 (p, $J$ = 6.6 Hz, 2H), 1.57 – 1.42 (m, 4H), 1.41 – 1.26 (m, 6H). $^{13}$C NMR (101 MHz, DMSO) δ 160.81, 160.31, 151.97, 150.63, 122.77, 117.82, 115.30, 107.31, 100.08, 96.94, 92.33, 85.18, 81.91, 76.72, 74.57, 72.09, 71.92, 71.88, 69.91, 69.75, 68.18, 61.09, 55.36, 29.95, 29.91, 28.96, 28.78, 28.35, 25.56, 25.53.

**Synthesis of (3*R*,4*S*,5*R*,6*R*)-6-(hydroxymethyl)-4-((8-((6-methoxyquinolin-4-yl)oxy)octyl)oxy)tetrahydro-2*H*-pyran-2,3,5-triol (4g)**

Compound **4g** was prepared in a similar manner as described for **3a**. Yield: 87.5%; ESI-MS *m/z*: 466.2429 $[M+H]^+$. $^1$H NMR (400 MHz, DMSO-$d_6$) δ 8.85 (d, $J$ = 2.2 Hz, 1H), 8.76 (s, 1H), 7.98 (dd, $J$ = 9.3, 1.9 Hz, 1H), 7.49 (dd, $J$ = 9.1, 2.5 Hz, 1H), 7.43 (s, 1H), 6.66 – 6.23 (m, 1H), 4.93 – 4.71 (m, 2H), 4.51 – 4.09 (m, 4H), 3.74 – 2.91 (m, 8H), 2.50 (s, 3H), 1.80 (p, $J$ = 6.2 Hz, 2H), 1.48 (dt, $J$ = 16.3, 7.2 Hz, 4H), 1.40 – 1.28 (m, 6H). $^{13}$C NMR (101 MHz, DMSO) δ 159.65, 145.55, 143.98, 142.90, 138.25, 130.20, 123.26, 107.41, 96.93, 92.34, 85.18, 81.91, 76.73, 74.59, 72.10, 71.95, 71.91, 69.91, 69.75, 68.28, 61.10, 29.97, 29.93, 29.00, 28.82, 28.50, 25.58, 25.54.

**Synthesis of (3*R*,4*S*,5*R*,6*R*)-4-((8-((6,7-dimethoxyquinolin-4-yl)oxy)octyl)oxy)-6-(hydroxymethyl)tetrahydro-2*H*-pyran-2,3,5-triol (4h)**

Compound **4h** was prepared in a similar manner described for **3a**. Yield: 89.2%; ESI-MS *m/z*: 496.2547 $[M+H]^+$. $^1$H NMR (400 MHz, DMSO-$d_6$) δ 7.86 (d, $J$ = 7.6 Hz, 1H), 7.54 (s, 1H), 7.02 (s, 1H), 6.76 – 6.27 (m, 1H), 5.96 (d, $J$ = 7.6 Hz, 1H), 5.00 – 4.76 (m, 2H), 4.58 – 4.17 (m, 4H), 3.93 (s, 3H), 3.83 (s, 3H), 3.70 – 2.88 (m, 8H), 1.73 (t, $J$ = 7.5 Hz, 2H), 1.45 (q, $J$ = 6.8 Hz, 2H), 1.35 – 1.16 (m, 8H). $^{13}$C NMR (101 MHz, DMSO) δ 175.07, 152.85, 146.35, 143.03, 135.16, 120.80, 107.84, 105.23, 98.31, 96.95, 92.34, 85.17, 81.90, 76.73, 74.58, 72.10, 71.91, 71.86, 69.90, 69.74, 61.08, 56.00, 55.48, 51.93, 29.91, 29.88, 28.92, 28.59, 28.39, 25.93, 25.48, 25.45.

**Synthesis of (3*R*,4*S*,5*R*,6*R*)-6-(hydroxymethyl)-4-((8-((7-(trifluoromethyl)quinolin-4-yl)oxy)octyl)oxy)tetrahydro-2*H*-pyran-2,3,5-triol (4i)**

Compound **4i** was prepared in a similar manner as described for **3a**. Yield: 87.0%; ESI-MS *m/z*: 504.2193 $[M+H]^+$. $^1$H NMR (400 MHz, DMSO-$d_6$) δ 8.86 (d, $J$ = 5.2 Hz, 1H), 8.35 (d, $J$ = 8.7 Hz, 1H), 8.27 (d, $J$ = 1.8 Hz, 1H), 7.83 (dd, $J$ = 8.8, 1.8 Hz, 1H), 7.18 (d, $J$ = 5.3 Hz, 1H), 6.72 – 6.28 (m, 1H), 4.98 – 4.76 (m, 2H), 4.56 – 4.19 (m, 4H), 3.73 – 2.89 (m, 8H), 1.94 – 1.81 (m, 2H), 1.50 (p, $J$ = 7.2 Hz, 4H), 1.43 – 1.24 (m, 6H). $^{13}$C NMR (101 MHz, DMSO) δ 160.69, 153.54, 147.58, 129.87 ($^2J$ = 32.3 Hz), 126.09 ($^3J$ = 5.1 Hz), 124.03 ($^1J$ = 273.7 Hz), 123.75, 122.85, 121.02, 120.99, 103.47, 96.93, 92.34, 85.19, 81.92, 76.73, 74.58, 72.10, 71.95, 71.90, 69.92, 69.75, 68.82, 61.09, 39.52, 29.96, 29.92, 28.97, 28.78, 28.25, 25.57, 25.54, 25.51.

**Synthesis of (3*R*,4*S*,5*R*,6*R*)-6-(hydroxymethyl)-4-((8-((7-hydroxyquinolin-4-yl)oxy)octyl)oxy)tetrahydro-2*H*-pyran-2,3,5-triol (4j)**

Compound **4j** was prepared in a similar manner described for **3a**. Yield: 46.8%; ESI-MS *m/z*: 452.2287 $[M+H]^+$. $^1$H NMR (400 MHz, DMSO-$d_6$) δ 10.07 (s, 1H), 8.54 (d, *J* = 5.2 Hz, 1H), 7.97 (d, *J* = 9.0 Hz, 1H), 7.16 (d, *J* = 2.4 Hz, 1H), 7.08 (dd, *J* = 9.0, 2.4 Hz, 1H), 6.78 (d, *J* = 5.3 Hz, 1H), 6.73 – 6.17 (m, 1H), 5.05 – 4.70 (m, 2H), 4.63 – 4.08 (m, 4H), 3.72 – 2.88 (m, 8H), 1.83 (h, *J* = 8.5, 7.5 Hz, 2H), 1.49 (h, *J* = 7.3 Hz, 4H), 1.41 – 1.27 (m, 6H). $^{13}$C NMR (101 MHz, DMSO) δ 160.83, 158.61, 151.76, 150.70, 122.85, 117.81, 114.48, 110.03, 99.27, 96.92, 92.33, 85.18, 81.90, 76.72, 74.58, 72.08, 71.92, 71.88, 69.92, 69.75, 68.04, 61.10, 29.96, 29.92, 28.98, 28.79, 28.38, 25.55.

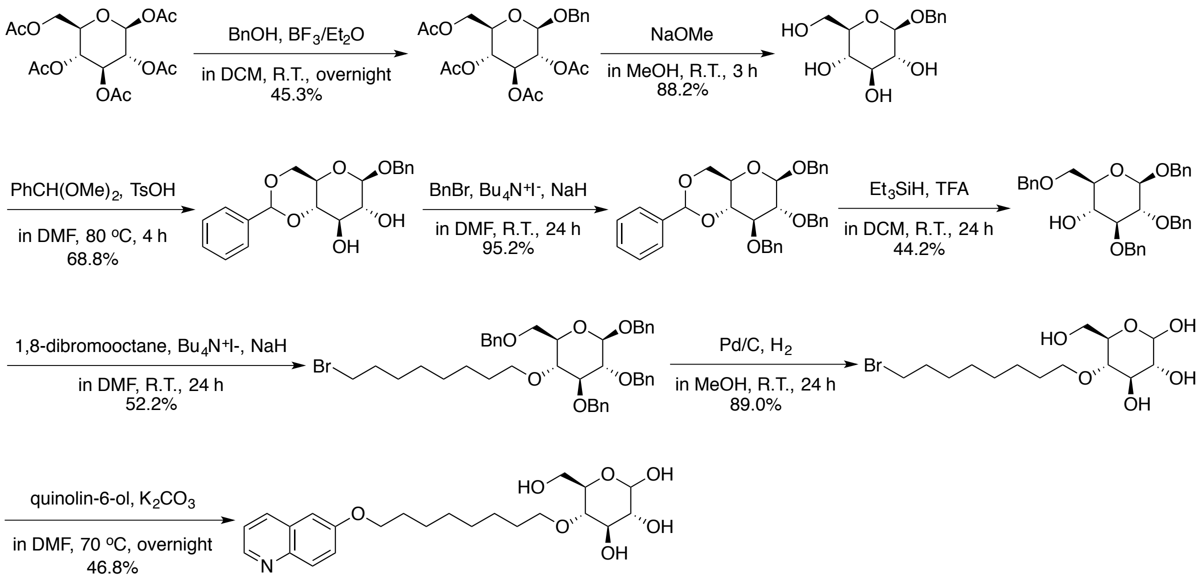

**Synthesis of (2*R*,3*R*,4*S*,5*R*,6*R*)-4,5,6-tris(benzyloxy)-2-((benzyloxy)methyl)tetrahydro-2*H*-pyran-3-ol (15)**

This compound was synthesized starting from *β*-D-glucose-penta-acetate following a modified procedure of previously established method (1). Yield: 44.2 %. $^1$H NMR (400 MHz, Chloroform-*d*) δ 7.44-7.27 (m, 20H), 4.95 (t, J = 11.4 Hz, 3H), 4.80-4.59 (m, 5H), 4.53 (d, J = 7.2 Hz, 1H), 3.80 (dd, J = 10.5, 3.8 Hz, 1H), 3.73 (dd, J = 10.4, 5.3 Hz, 1H), 3.65-3.58 (m, 1H), 3.53-3.42 (m, 3H), 2.44 (s, 1H). $^{13}$C NMR (101 MHz, $CDCl_3$) δ 138.73, 138.47, 138.09, 137.47, 128.64, 128.53, 128.45, 128.28, 128.09, 128.06, 127.93, 127.91, 127.84, 127.80, 102.71, 84.21, 81.89, 75.39, 74.90, 74.27, 73.78, 71.63, 71.28, 70.36.

**Synthesis of (2*R*,3*R*,4*S*,5*R*,6*R*)-2,3,4-tris(benzyloxy)-6-((benzyloxy)methyl)-5-((8-bromooctyl)oxy)tetrahydro-2*H*-pyran (16)**

Compound **15** (720 mg, 1.33 mmol) was dissolved in 10 ml anhydrous *N,N*-dimethylformamide (DMF) in a 50-ml round-bottomed flask. Sodium hydride (80 mg, 2 mmol) in mineral oil (60%) and tetra-*n*-butylammonium iodide (246 mg, 0.67 mmol) were then added under argon in small portions over a period of 5 minutes. The mixture was stirred for 30 minutes in ice-water bath. Then 1,8-dibromooctane (490 µl, 2.66 mmol) was added slowly. The mixture was allowed to warm to room temperature and stirred for 24 h. The progress of the reaction was monitored by TLC. Upon completion of the reaction, organic solvents were removed under vacuum generating precipitants in water. The resulting residue was extracted with ethyl acetate, dried over anhydrous sodium sulfate, filtered, and concentrated under vacuum. The crude product was purified with flash column chromatography on silica gel (8 % ethyl acetate in petroleum ether) to yield 509 mg (52.2%) (2*R*,3*R*,4*S*,5*R*,6*R*)-2,3,4-tris(benzyloxy)-6-((benzyloxy)methyl)-5-((8-bromooctyl)oxy)tetrahydro-2*H*-pyran (**16**) as faint yellow oil. $^{1}$H NMR (400 MHz, Chloroform-*d*) δ 7.79 – 6.99 (m, 20H), 5.03 – 4.53 (m, 8H), 4.49 (d, *J* = 7.5 Hz, 1H), 3.81 – 3.61 (m, 3H), 3.57 – 3.33 (m, 7H), 1.48 – 1.17 (m, 12H). $^{13}$C NMR (101 MHz, $CDCl_3$) δ 139.04, 138.75, 138.50, 138.33, 137.54, 128.39, 128.34, 128.31, 128.16, 127.95, 127.78, 127.73, 127.69, 127.59, 127.58, 127.54, 102.59, 84.70, 82.23, 78.20, 75.61, 75.16, 74.89, 73.53, 73.13, 71.12, 69.11, 33.94, 32.78, 30.33, 29.31, 28.68, 28.11, 26.03.

**Synthesis of (2*R*,3*R*,4*R*,5*S*,6*R*)-5-((8-bromooctyl)oxy)-6-(hydroxymethyl)tetrahydro-2*H*-pyran-2,3,4-triol (17)**

Compound **16** (509 mg, 0.70 mmol) was dissolved in 5 ml methanol in a 25-ml round bottomed flask. Palladium (10%) on carbon (172 mg, 0.16 mmol) was then added, and the flask was evacuated carefully and flushed with hydrogen gas for three times using a balloon. The mixture was stirred at room temperature for 24 h under a hydrogen atmosphere (using a balloon). Upon completion of the reaction, the Pd/C was filtered off using a pad of celite and the filtrate was concentrated. The crude product was purified with flash column chromatography on silica gel (8% methanol in dichloromethane) to yield 230 mg (89.0%) (2*R*,3*R*,4*R*,5*S*,6*R*)-5-((8-bromooctyl)oxy)-6-(hydroxymethyl)tetrahydro-2*H*-pyran-2,3,4-triol (**17**) as white solid. $^{1}$H NMR (400 MHz, DMSO-$d_6$) δ 6.71 – 6.16 (m, 1H), 4.92 – 4.46 (m, 3H), 4.46 – 4.22 (m, 1H), 3.75 (dt, *J* = 9.3, 6.4 Hz, 1H), 3.66 – 3.38 (m, 6H), 3.25 – 2.85 (m, 3H), 1.78 (p, *J* = 6.8 Hz, 2H), 1.55 – 1.32 (m, 4H), 1.30 – 1.23 (m, 6H). $^{13}$C NMR (101 MHz, DMSO) δ 96.74, 92.10, 78.33, 78.09, 76.66, 75.52, 75.06, 73.01, 72.57, 71.52, 70.77, 60.68, 35.17, 32.22, 31.24, 29.83, 29.81, 28.88, 28.75, 28.73, 28.69, 28.08, 27.48, 25.51, 25.49, 22.06, 13.93.

**Synthesis of (2*R*,3*R*,4*R*,5*S*,6*R*)-6-(hydroxymethyl)-5-((8-(quinolin-6-yloxy)octyl)oxy)tetrahydro-2*H*-pyran-2,3,4-triol (5c)**

Compound **17** (216 mg, 0.58 mmol) was dissolved in 10 ml anhydrous DMF in a 50-ml round-bottomed flask, and quinolin-6-ol (488 mg, 3.36 mmol), potassium carbonate (512 mg, 3.70 mmol) were then added into the solution. The mixture was stirred at 70℃ for 10 h. The progress of the reaction was monitored by TLC. Upon completion of the reaction, the mixture was cooled to room temperature and organic solvents were removed under vacuum. The resulting precipitant was extracted with ethyl acetate, dried over anhydrous sodium sulfate, filtered, and concentrated under vacuum. The crude product was purified using flash column chromatography on silica gel (10 % ethyl acetate in petroleum ether) to yield 118 mg (46.8%) (2*R*,3*R*,4*R*,5*S*,6*R*)-6-(hydroxymethyl)-5-((8-(quinolin-6-yloxy)octyl)oxy)tetrahydro-2*H*-pyran-2,3,4-triol (**5c**) as white solid. Yield: 46.8%; ESI-MS *m/z*: 436.2331 $[M+H]^+$. $^1H$ NMR (400 MHz, DMSO-$d_6$) δ 8.72 (dd, *J* = 4.2, 1.7 Hz, 1H), 8.23 (dd, *J* = 8.4, 1.6 Hz, 1H), 7.90 (d, *J* = 9.0 Hz, 1H), 7.46 (dd, *J* = 8.3, 4.2 Hz, 1H), 7.42 – 7.32 (m, 2H), 6.72 – 6.18 (m, 1H), 5.02 – 4.67 (m, 2H), 4.65 – 4.52 (m, 1H), 4.52 – 4.22 (m, 1H), 4.08 (t, *J* = 6.5 Hz, 2H), 3.82 – 3.40 (m, 5H), 3.27 – 2.86 (m, 3H), 1.77 (p, *J* = 6.8 Hz, 2H), 1.44 (p, *J* = 6.8 Hz, 4H), 1.39 – 1.24 (m, 6H). $^{13}C$ NMR (101 MHz, DMSO) δ 156.58, 147.85, 143.73, 134.79, 130.33, 129.11, 122.25, 121.64, 106.33, 96.80, 92.17, 78.38, 78.13, 76.71, 75.57, 75.10, 73.07, 72.62, 71.64, 70.84, 67.88, 60.72, 29.93, 29.92, 28.95, 28.84, 28.62, 25.64, 25.63, 25.57.

**Synthesis of (2*R*,3*S*,4*S*,5*R*)-2-(hydroxymethyl)-6-((8-(quinolin-6-yloxy)octyl)oxy)tetrahydro-2*H*-pyran-3,4,5-triol (5a)**

This compound was synthesized from (3*R*,4*S*,5*R*,6*R*)-3,4,5-tris(benzyloxy)-6-((benzyloxy)methyl)tetrahydro-2*H*-pyran-2-ol following a similar procedure described for **5c**. Yield: 52.4%; ESI-MS *m/z*: 436.2326 $[M+H]^+$. $^1H$ NMR (400 MHz, Methanol-$d_4$) δ 8.66 (d, *J* = 4.1 Hz, 1H), 8.26 (d, *J* = 8.3 Hz, 1H), 7.91 (d, *J* = 9.2 Hz, 1H), 7.47 (dd, *J* = 8.4, 4.3 Hz, 1H), 7.41 (dd, *J* = 9.2, 2.8 Hz, 1H), 7.28 (d, *J* = 2.7 Hz, 1H), 4.25 (d, *J* = 7.8 Hz, 1H), 4.12 (t, *J* = 6.4 Hz, 2H), 3.94 – 3.84 (m, 2H), 3.67 (dd, *J* = 12.1, 4.9 Hz, 1H), 3.54 (dt, *J* = 9.6, 6.8 Hz, 1H), 3.35 (t, *J* = 8.0 Hz, 1H), 3.27 (t, *J* = 5.9 Hz, 2H), 3.17 (t, *J* = 8.4 Hz, 1H), 1.86 (p, *J* = 6.7 Hz, 2H), 1.63 (q, *J* = 6.9 Hz, 2H), 1.53 (q, *J* = 7.1 Hz, 2H), 1.42 (dd, *J* = 11.0, 5.3 Hz, 6H). $^{13}C$ NMR (101 MHz, MeOD) δ 158.97, 148.39, 144.66, 137.17, 131.22, 130.31, 124.23, 122.68, 107.19, 104.37, 78.14, 77.92, 75.13, 71.68, 70.86, 69.46, 62.79, 30.77, 30.52, 30.46, 30.27, 27.15, 27.05.

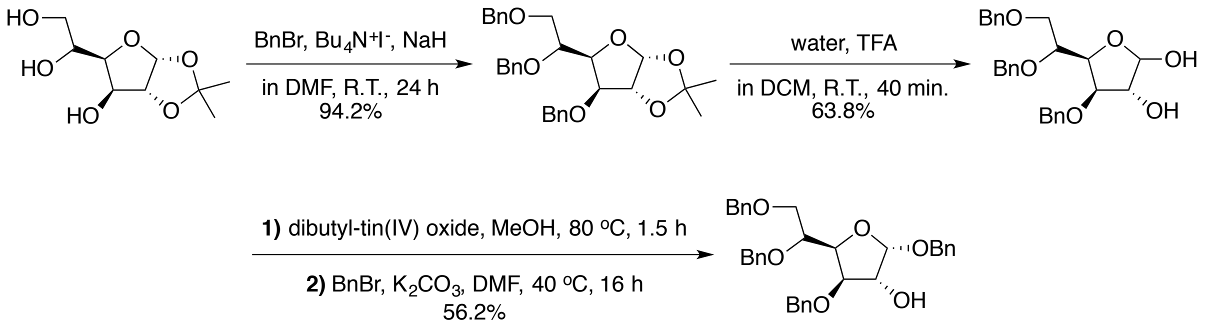

**Synthesis of (2*S*,3*R*,4*R*,5*R*)-2,4-bis(benzyloxy)-5-((*R*)-1,2-bis(benzyloxy)ethyl)tetrahydrofuran-3-ol (18)**

This compound was synthesized starting from 1,2-O-isopropylidene-α-D-glucofuranose following a modified procedure of previously established method, and the characterization data matched well to those reported previously (1). Yield: 56.2 %. $^1$H NMR (400 MHz, Chloroform-*d*) δ 7.43 – 7.18 (m, 20H), 5.22 (d, *J* = 4.6 Hz, 1H), 4.79 (dd, *J* = 13.0, 11.6 Hz, 2H), 4.70 (d, *J* = 11.7 Hz, 1H), 4.62 – 4.46 (m, 5H), 4.36 (dd, *J* = 8.4, 4.4 Hz, 1H), 4.25 (ddd, *J* = 6.2, 4.6, 1.9 Hz, 1H), 4.09 – 3.99 (m, 2H), 3.84 (dd, *J* = 10.6, 2.1 Hz, 1H), 3.68 (dd, *J* = 10.6, 5.7 Hz, 1H), 2.96 (d, *J* = 5.8 Hz, 1H). $^{13}$C NMR (101 MHz, $CDCl_3$) δ 139.00, 138.71, 138.02, 137.18, 128.67, 128.61, 128.46, 128.42, 128.34, 128.27, 128.13, 127.75, 127.72, 127.66, 127.65, 127.55, 127.48, 100.37, 84.08, 77.96, 76.48, 76.22, 73.56, 72.74, 71.76, 71.25, 70.22.

**Synthesis of (2*R*,3*R*,4*S*,5*S*,6*R*)-6-(hydroxymethyl)-3-((8-(quinolin-6-yloxy)octyl)oxy)tetrahydro-2*H*-pyran-2,4,5-triol (5b)**

This compound was synthesized from (2*S*,3*R*,4*R*,5*R*)-2,4-bis(benzyloxy)-5-((*R*)-1,2-bis(benzyloxy)ethyl)tetrahydrofuran-3-ol (**18**) following a similar procedure described for **5c**. Yield: 51.0%; ESI-MS *m/z*: 436.2328 [M+H]$^+$. $^1$H NMR (400 MHz, DMSO-$d_6$) δ 8.72 (dd, *J* = 4.2, 1.7 Hz, 1H), 8.23 (dd, *J* = 8.4, 1.6 Hz, 1H), 7.90 (d, *J* = 9.1 Hz, 1H), 7.46 (dd, *J* = 8.3, 4.2 Hz, 1H), 7.38 (dd, *J* = 9.1, 2.8 Hz, 1H), 7.35 (d, *J* = 2.8 Hz, 1H), 6.75 – 6.15 (m, 1H), 5.14 – 4.79 (m, 2H), 4.77 – 4.44 (m, 1H), 4.44 – 4.30 (m, 1H), 4.08 (t, *J* = 6.5 Hz, 2H), 3.74 – 3.37 (m, 5H), 3.19 – 2.69 (m, 3H), 1.78 (p, *J* = 6.7 Hz, 2H), 1.53 – 1.40 (m, 4H), 1.38 – 1.25 (m, 6H). $^{13}$C NMR (101 MHz, DMSO) δ 156.57, 147.84, 143.72, 134.77, 130.33, 129.09, 122.24, 121.63, 106.33, 96.66, 89.98, 83.23, 80.41, 76.59, 76.11, 72.04, 71.79, 71.71, 70.74, 70.53, 69.68, 67.88, 61.19, 61.18, 29.84, 29.76, 28.98, 28.96, 28.84, 28.62, 25.56.

**Synthesis of (3*R*,4*S*,5*S*,6*R*)-3-((8-((2-chloroquinolin-6-yl)oxy)octyl)oxy)-6-(hydroxymethyl)tetrahydro-2*H*-pyran-2,4,5-triol (5d)**

Compound **5d** was prepared in a similar manner described for **5b**. Yield: 68.2%; ESI-MS *m/z*: 492.1754 [M+Na]$^+$. $^1$H NMR (400 MHz, DMSO-$d_6$) δ 8.31 (d, *J* = 8.5 Hz, 1H), 7.93 – 7.78 (m,

1H), 7.52 (d, *J* = 8.6 Hz, 1H), 7.48 – 7.36 (m, 2H), 6.69 – 6.15 (m, 1H), 5.13 – 4.78 (m, 2H), 4.75 – 4.34 (m, 2H), 4.08 (t, *J* = 6.5 Hz, 2H), 3.73 – 3.38 (m, 6H), 3.15 – 2.90 (m, 2H), 1.77 (p, *J* = 6.7 Hz, 2H), 1.52 – 1.39 (m, 4H), 1.37 – 1.26 (m, 6H). $^{13}$C NMR (101 MHz, DMSO) δ 157.07, 147.01, 142.99, 138.74, 129.23, 128.04, 123.34, 122.47, 106.69, 96.66, 89.97, 83.23, 80.40, 76.59, 76.11, 72.04, 71.79, 71.70, 70.74, 70.53, 69.68, 68.03, 61.19, 29.75, 28.96, 28.82, 28.57, 25.56, 25.53.

**Synthesis of (3*R*,4*S*,5*S*,6*R*)-6-(hydroxymethyl)-3-((9-(quinolin-6-yloxy)nonyl)oxy)tetrahydro-2*H*-pyran-2,4,5-triol (5e)**

Compound **5e** was prepared in a similar manner described for **5b**. Yield: 56.0%; ESI-MS *m/z*: 450.2510 [M+H]$^{+}$. $^{1}$H NMR (400 MHz, DMSO-$d_6$) δ 8.72 (dd, *J* = 4.2, 1.7 Hz, 1H), 8.23 (ddd, *J* = 8.4, 1.7, 0.8 Hz, 1H), 7.90 (d, *J* = 9.0 Hz, 1H), 7.46 (dd, *J* = 8.3, 4.2 Hz, 1H), 7.42 – 7.33 (m, 2H), 6.71 – 6.13 (m, 1H), 5.11 – 4.77 (m, 2H), 4.73 – 4.26 (m, 2H), 4.09 (t, *J* = 6.5 Hz, 2H), 3.74 – 3.38 (m, 5H), 3.20 – 2.67 (m, 3H), 1.78 (p, *J* = 6.7 Hz, 2H), 1.51 – 1.40 (m, 4H), 1.38 – 1.26 (m, 8H). $^{13}$C NMR (101 MHz, DMSO) δ 156.56, 147.83, 143.71, 134.76, 130.32, 129.08, 122.22, 121.62, 106.33, 96.64, 89.96, 83.22, 80.40, 76.58, 76.10, 72.02, 71.77, 71.70, 70.72, 70.52, 69.68, 67.87, 61.17, 29.84, 29.76, 29.06, 28.98, 28.95, 28.80, 28.62, 25.58.

### Supplementary Figures and Tables

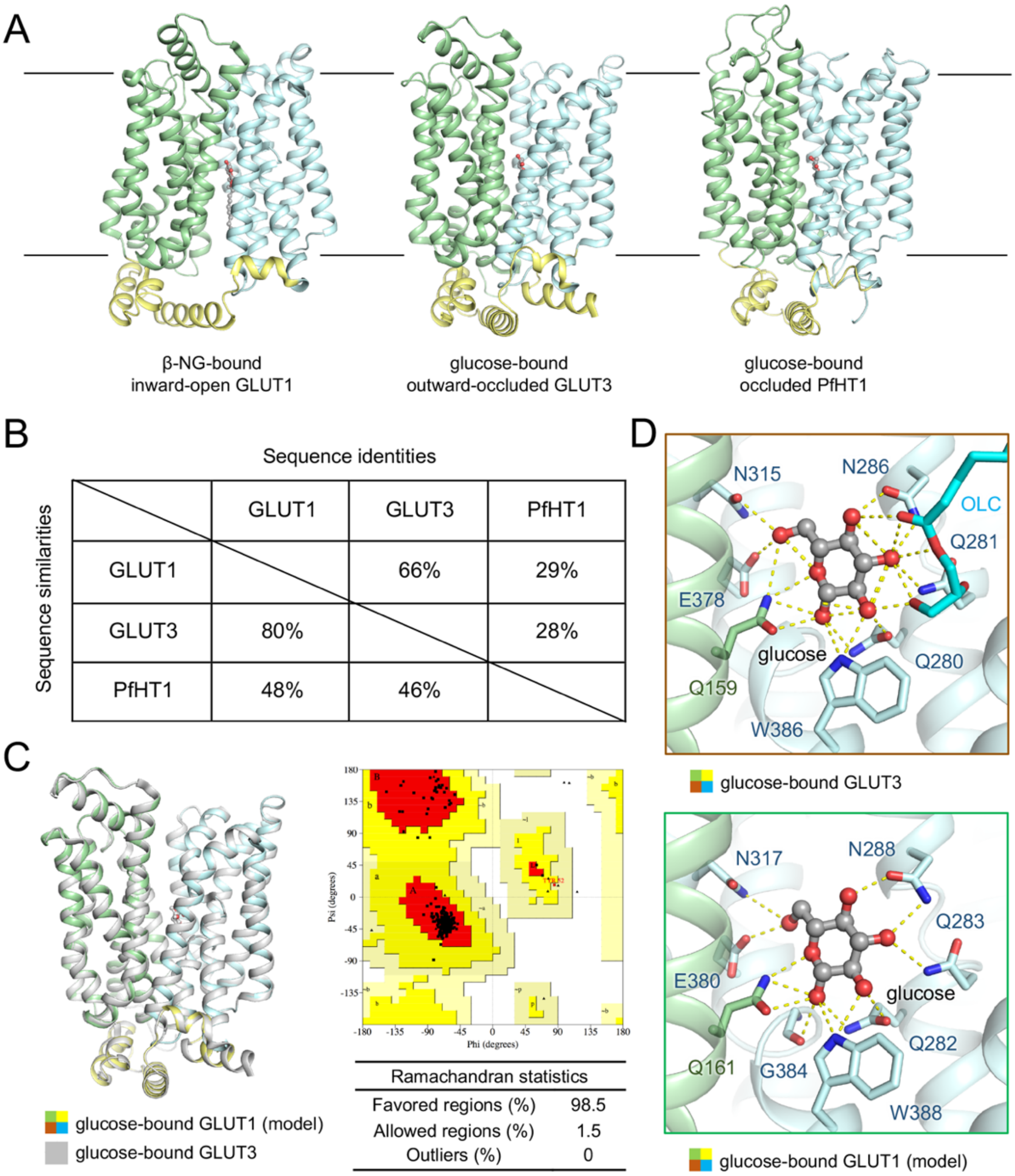

**Figure S1. Homolog modelling of human glucose transporter in outward-occluded conformation. (A)** An overview of the resolved structures of human hGLUT1, hGLUT3 and PfHT1. **(B)** Sequence conservation of the indicated sugar transporters. Pairwise sequence comparison was performed in Blastp (NCBI. Basic Local Alignment Search Tool). The Uniprot (https://www.uniprot.org/) entry numbers of aligned proteins are: PfHT1 (Q7KWJ5), hGLUT1 (P11166), and hGLUT3 (P11169). **(C)** Homolog model of hGLUT1 in outward-occluded conformation were built and refined using Modeller-9.19, and the best model was chosen by PROCHECK. Glucose was redocked into the hGLUT1 model by XP glide docking program in Schrodinger suite. Shown here is an overlay of the crystal structure of glucose-bound hGLUT3 (gray) and the predicted model of glucose-bound PfHT1 complex (domain colored). **(D)** The predicted binding mode of glucose in hGLUT1 model compared to the coordination of glucose in the outward-occluded hGLUT3 structure. Details of the polar interactions of glucose with residues from the N-terminal and C-terminal domains are shown by yellow dashed lines.

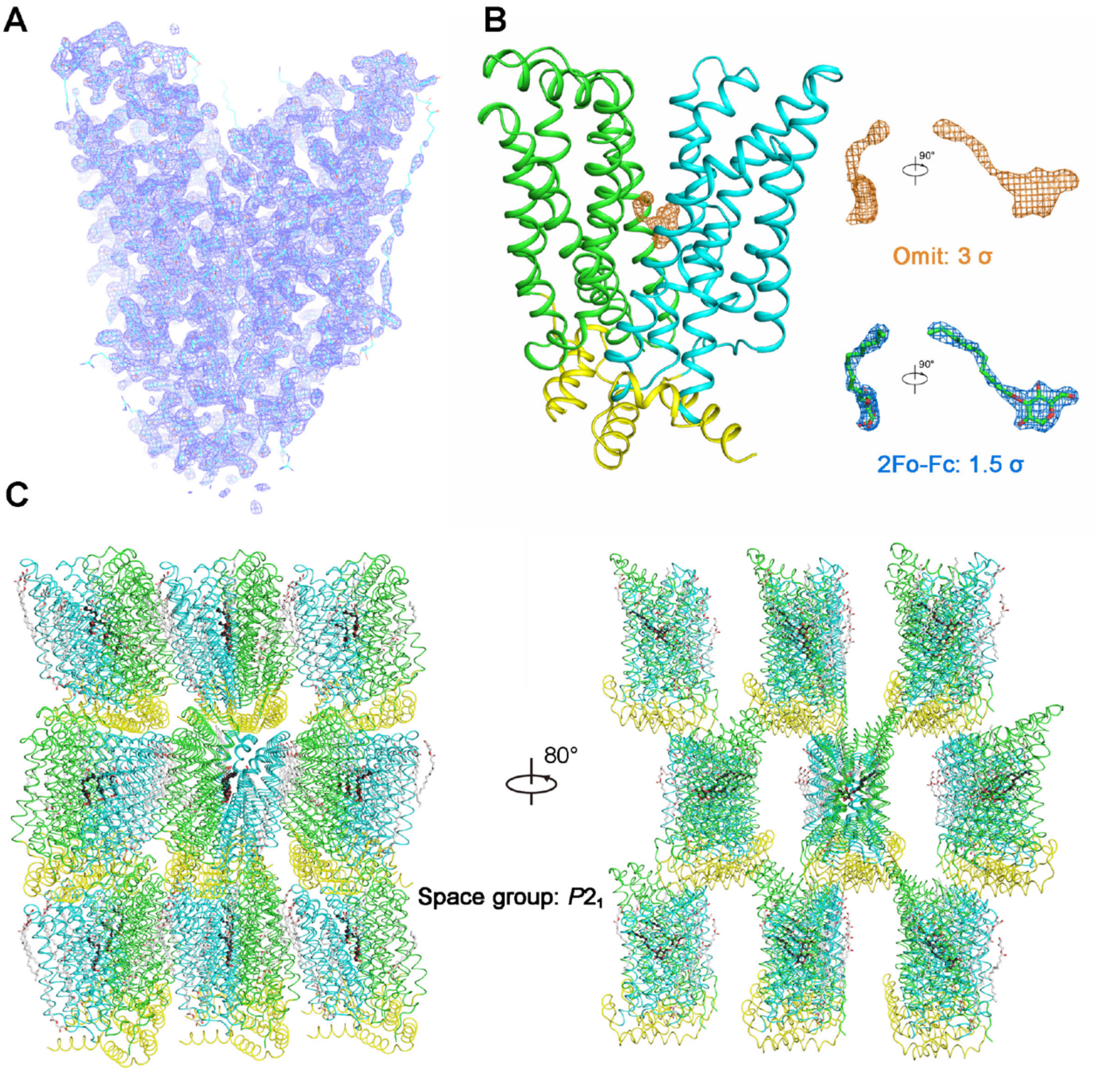

**Figure S2. Structural determination of GLUT3 bound to C3361. (A)** Electron density map of the GLUT3-**C3361** complex at 2.30 Å resolution. **(B)** Omit and 2Fo-Fc maps for the bound inhibitor, **C3361**. **(C)** Crystal packing of GLUT3-**C3361** in the space group $P2_1$.

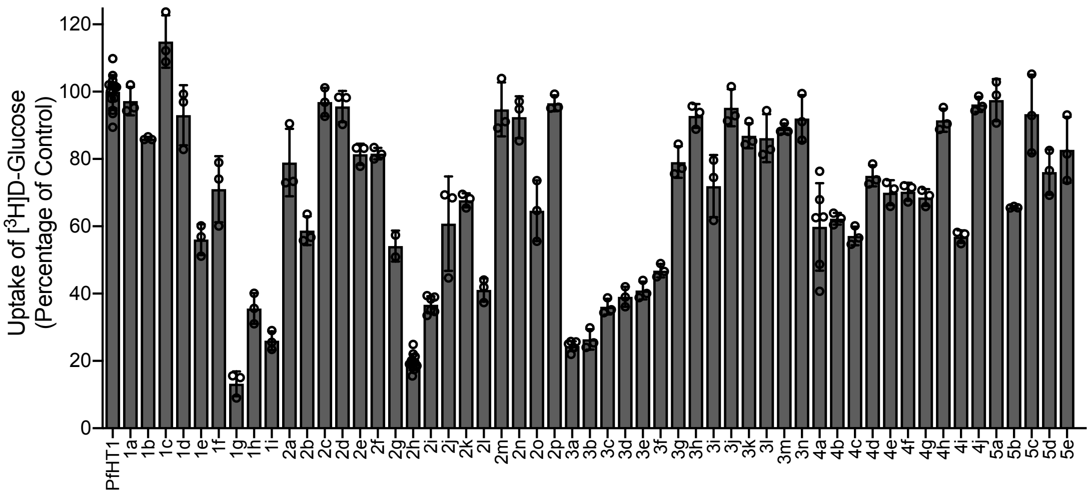

**Figure S3. The inhibitory activity of synthetic small molecules against PfHT1 using proteoliposome-based counterflow assay.** All of the compounds were tested at 1 μM.

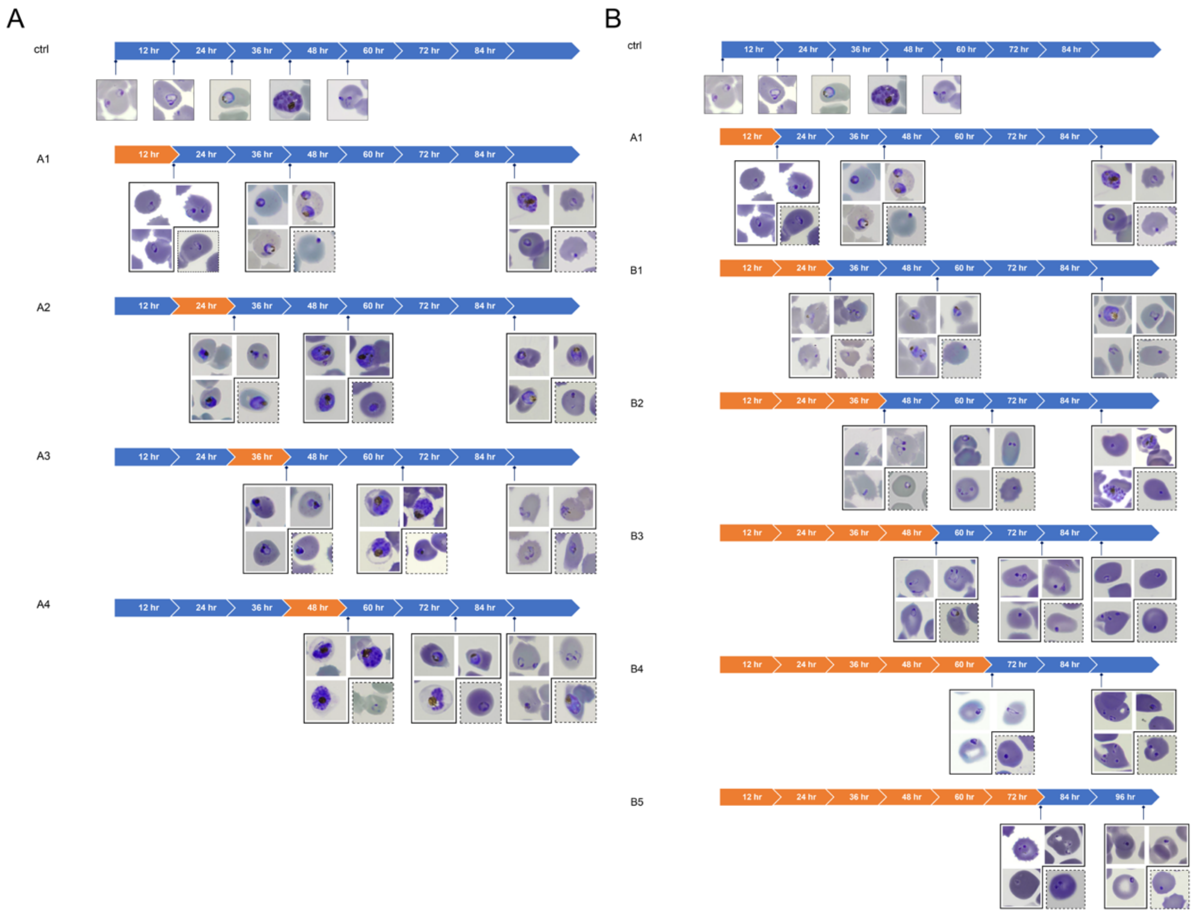

**Figure S4. Representative images of compound treated parasites in the substage assay described in Figure 5.** Solid outline: **TH-PF01** treated parasites; dotted outline: DHA treated parasites.

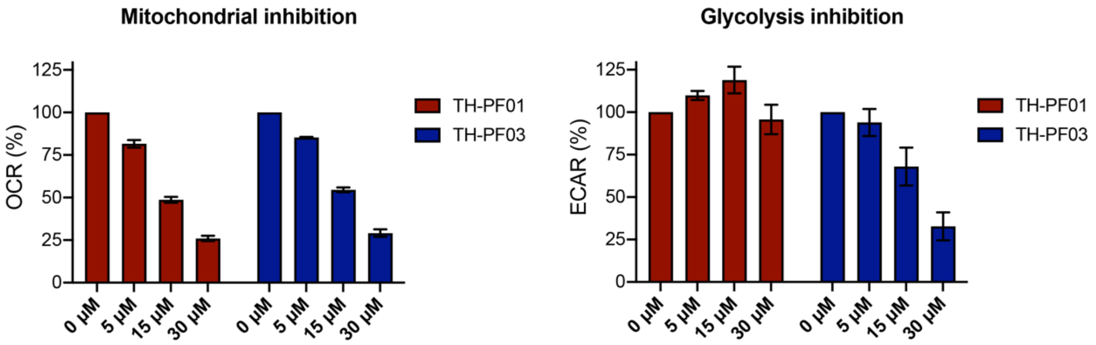

**Figure S5. Cytotoxicity of TH-PF01 and TH-PF03 are likely due to mitochondrial toxicity.** Extracellular flux analysis revealed minor mitochondrial inhibition of **TH-PF01** and **TH-PF03** and glycolysis inhibition of **TH-PF03** at higher doses in HEK293T$_{17}$ cells. HEK293T$_{17}$ cells were seeded in assay medium supplemented with glucose (25 mM), pyruvate (1 mM) and glutamine (4 mM), and **TH-PF01** or **TH-PF03** was sequentially added four times with the final concentrations of 5, 15, and 30 μM. OCR and ECAR values detected after each compound injection were normalized with negative control (media) as 100% and with positive controls (antimycin A/rotenone for OCR, 2-DG for ECAR) as 0%. All data were average values pooled from two independent experiments with two technical replicates. Error bars represent SEM.

**Table S1. Data collection and refinement statistics**

| | GLUT3-C3361 |
|---|---|
| **Data collection** | |
| Light Sources | BL32XU, SPring-8, Japan |
| Number of crystals | 247 |
| Space group | $P2_1$ |
| Cell dimensions | |
| *a*, *b*, *c* (Å) | 48.329, 118.248, 54.252 |
| α, β, γ (°) | 90.00, 103.54, 90.00 |
| Resolution (Å) | 50 ~ 2.30 (2.40 ~ 2.30) |
| $R_{sym}$ or $R_{merge}$ | 0.51 (2.41) |
| $I / \sigma I$ | 8.57 (2.10) |
| $CC_{1/2}$ (%) | 99.9 (100) |
| Completeness (%) | 39.63 (34.67) |
| Redundancy | 99.2 (87.8) |
| **Refinement** | |
| Resolution (Å) | 19.81 ~ 2.30 |
| No. reflections (test set) | 26, 263 (1,320) |
| $R_{work}$ / $R_{free}$ | 0.1657/0.2120 |
| No. atoms | |
| Protein | 3,608 |
| Ligand (Cpd3361) | 23 |
| Others (Lipids, Waters) | 285 |
| *B*-factors | |
| Protein | 34.76 |
| Ligand (Cpd3361) | 34.39 |
| Others (Lipids, Waters) | 51.56 |
| R.m.s. deviations | |
| Bond lengths (Å) | 0.006 |
| Bond angles (°) | 0.970 |
| Ramachandran statistics | |
| Favored regions (%) | 98.5 |
| Allowed regions (%) | 1.3 |
| Outliers (%) | 0.2 |

*Values in parentheses are for highest-resolution shell.

**Table S2. Characterization of *in vitro* parasite inhibition and cytotoxicity for each compound.**

| R[1]-linker-R[2] | | | | | | | |
|---|---|---|---|---|---|---|---|
| **Cmpd.** | **Chemical Structure** | | | **EC$_{50}$ (μM)** | | **CC$_{50}$ (μM)** | |
| | **R[1]** | **linker** | **R[2]** | ***Pf*3D7** | ***Pf*Dd2** | **HEK293T$_{17}$** | **HepG2** |
| **1a** | 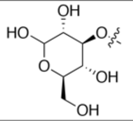 | -CH$_2$COO-(CH$_2$)$_5$- | 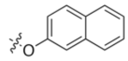 | > 50 | > 50 | > 50 | > 50 |
| **1b** | .. | -CH$_2$COO-(CH$_2$)$_2$-O-(CH$_2$)$_2$- | 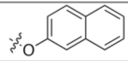 | > 50 | > 50 | > 50 | > 50 |
| **1c** | .. | -CH$_2$CONH-(CH$_2$)$_5$- | 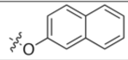 | > 50 | > 50 | > 50 | > 50 |
| **1d** | .. | -CH$_2$COO-(CH$_2$)$_5$- | 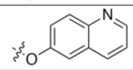 | > 50 | > 50 | > 50 | > 50 |
| **1e** | .. | -(CH$_2$)$_8$- | 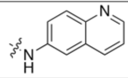 | 2.22 ± 0.55 | 1.44 ± 0.11 | > 50 | > 50 |
| **1f** | .. | -(CH$_2$)$_7$- | 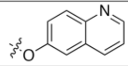 | 12.6 ± 0.2 | 9.74 ± 1.01 | > 50 | > 50 |
| **1g** (TH-PF02) | .. | -(CH$_2$)$_9$- | 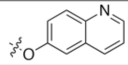 | 0.308 ± 0.004 | 0.298 ± 0.021 | 14.4 ± 4.7 | 34.5 ± 4.1 |
| **1h** | .. | -(CH$_2$)$_{11}$- | 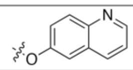 | 2.28 ± 0.24 | 2.51 ± 0.17 | 10.6 ± 0.4 | > 50 |
| **1i** | .. | -(CH$_2$)$_9$- | 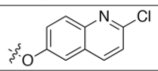 | 0.385 ± 0.033 | 0.351 ± 0.034 | 10.8 ± 1.3 | > 50 |
| **2a** | .. | -(CH$_2$)$_8$- | 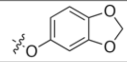 | 6.83 ± 0.63 | 6.83 ± 0.61 | > 50 | > 50 |
| **2b** | .. | -(CH$_2$)$_8$- | 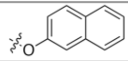 | 0.998 ± 0.097 | 0.839 ± 0.264 | 15.7 ± 5.9 | 27.5 ± 2.3 |
| **2c** | .. | -(CH$_2$)$_8$- | 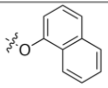 | 15.7 ± 0.6 | 13.3 ± 1.1 | 18.0 ± 2.5 | 29.1 ± 8.6 |
| **2d** | .. | -(CH$_2$)$_8$- | 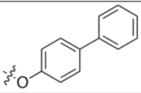 | 1.12 ± 0.07 | 1.10 ± 0.07 | 9.42 ± 3.55 | 18.1 ± 2.0 |
| **2e** | .. | -(CH$_2$)$_8$- | 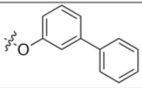 | 4.79 ± 0.19 | 4.11 ± 0.44 | 15.3 ± 1.7 | 28.3 ± 6.5 |
| **2f** | .. | -(CH$_2$)$_8$- | 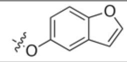 | 9.53 ± 0.57 | 9.81 ± 0.67 | > 50 | > 50 |
| **2g** | .. | -(CH$_2$)$_8$- | 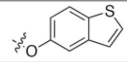 | 1.49 ± 0.27 | 1.26 ± 0.04 | 35.7 ± 4.1 | > 50 |
| **2h** | .. | -(CH$_2$)$_8$- | 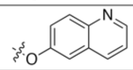 | 0.377 ± 0.027 | 0.376 ± 0.043 | 42.7 ± 2.0 | > 50 |
| **2i** | .. | -(CH$_2$)$_8$- | 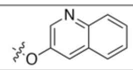 | 1.25 ± 0.40 | 0.656 ± 0.072 | 17.5 ± 4.0 | > 33 |
| **2j** | .. | -(CH$_2$)$_8$- | 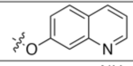 | 6.91 ± 0.93 | 5.02 ± 0.30 | > 50 | > 50 |
| **2k** | .. | -(CH$_2$)$_8$- | 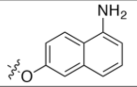 | 5.20 ± 1.34 | 2.35 ± 0.08 | > 50 | > 50 |
| **2l** | .. | -(CH$_2$)$_8$- | 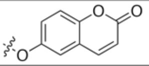 | 1.94 ± 0.34 | 1.52 ± 0.09 | > 50 | > 50 |

| | | | | | | | |
|---|---|---|---|---|---|---|---|
| **2m** | .. | -(CH$_2$)$_8$- | 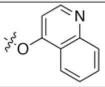 | 28.6 ± 0.6 | 15.9 ± 0.6 | 23.5 ± 2.7 | > 50 |
| **2n** | .. | -(CH$_2$)$_8$- | 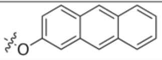 | 1.74 ± 0.21 | 1.72 ± 0.85 | 8.95 ± 2.77 | 36.3 ± 9.5 |
| **2o** | .. | -(CH$_2$)$_8$- | 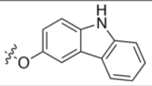 | 0.731 ± 0.101 | 0.545 ± 0.036 | 14.8 ± 4.2 | 21.6 ± 2.2 |
| **2p** | .. | -(CH$_2$)$_8$- | 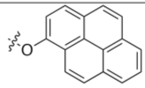 | 3.61 ± 0.75 | 2.83 ± 0.60 | 3.58 ± 2.01 | > 50 |
| **3a** (TH-PF01) | .. | -(CH$_2$)$_8$- | 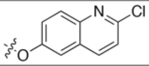 | 0.371 ± 0.026 | 0.349 ± 0.003 | 13.4 ± 0.5 | > 50 |
| **3b** | .. | -(CH$_2$)$_8$- | 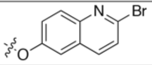 | 0.409 ± 0.112 | 0.297 ± 0.037 | 10.4 ± 2.5 | 28.4 ± 4.7 |
| **3c** | .. | -(CH$_2$)$_8$- | 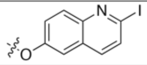 | 0.436 ± 0.021 | 0.404 ± 0.012 | 7.52 ± 1.24 | 23.4 ± 3.2 |
| **3d** | .. | -(CH$_2$)$_8$- | 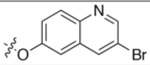 | 1.39 ± 0.02 | 1.23 ± 0.17 | > 50 | > 50 |
| **3e** | .. | -(CH$_2$)$_8$- | 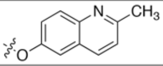 | 1.45 ± 0.38 | 0.734 ± 0.019 | 25.2 ± 4.3 | > 50 |
| **3f** | .. | -(CH$_2$)$_8$- | 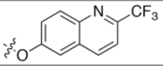 | 0.833 ± 0.143 | 0.708 ± 0.034 | 12.4 ± 2.5 | 34.7 ± 4.2 |
| **3g** | .. | -(CH$_2$)$_8$- | 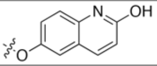 | 4.50 ± 0.13 | 4.03 ± 0.35 | > 50 | > 50 |
| **3h** | .. | -(CH$_2$)$_8$- | 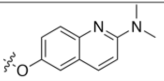 | 3.64 ± 0.58 | 3.36 ± 0.26 | 41.4 ± 2.5 | > 50 |
| **3i** | .. | -(CH$_2$)$_8$- | 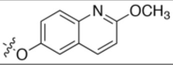 | 1.30 ± 0.30 | 0.917 ± 0.099 | 21.9 ± 6.2 | > 50 |
| **3j** | .. | -(CH$_2$)$_8$- | 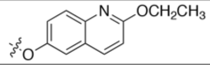 | 4.25 ± 0.59 | 2.13 ± 0.12 | 13.7 ± 0.2 | > 50 |
| **3k** | .. | -(CH$_2$)$_8$- | 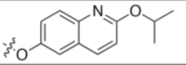 | 12.5 ± 2.7 | 4.29 ± 0.28 | 8.30 ± 0.07 | 19.0 ± 2.2 |
| **3l** | .. | -(CH$_2$)$_8$- | 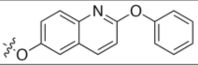 | 8.50 ± 2.30 | 4.36 ± 0.30 | 7.84 ± 0.06 | 21.1 ± 0.4 |
| **3m** | .. | -(CH$_2$)$_8$- | 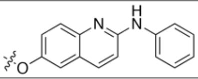 | 1.17 ± 0.26 | 0.894 ± 0.166 | 2.61 ± 0.05 | 10.8 ± 1.7 |
| **3n** | .. | -(CH$_2$)$_8$- | 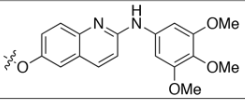 | 1.71 ± 0.38 | 1.12 ± 0.01 | > 20 | > 50 |
| **4a** | .. | -(CH$_2$)$_8$- | 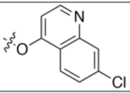 | 0.924 ± 0.310 | 0.499 ± 0.039 | 6.93 ± 0.69 | 17.5 ± 2.4 |
| **4b** | .. | -(CH$_2$)$_8$- | 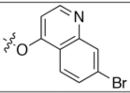 | 0.657 ± 0.217 | 0.376 ± 0.007 | 6.89 ± 0.82 | > 20 |
| **4c** | .. | -(CH$_2$)$_8$- | 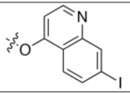 | 0.643 ± 0.031 | 0.473 ± 0.114 | 15.3 ± 9.7 | 19.8 ± 0.9 |
| **4d** | .. | -(CH$_2$)$_8$- | 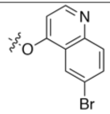 | 3.37 ± 0.10 | 2.75 ± 0.23 | 4.13 ± 0.87 | 7.96 ± 0.57 |

| | | | | | | | |
|---|---|---|---|---|---|---|---|
| **4e** | .. | -(CH2)8- | 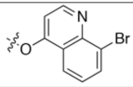 | 5.84 ± 0.30 | 4.66 ± 0.61 | 14.2 ± 2.7 | 34.7 ± 3.7 |
| **4f** | .. | -(CH2)8- | 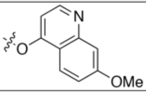 | 2.42 ± 0.52 | 1.58 ± 0.10 | 14.8 ± 2.0 | > 50 |
| **4g** | .. | -(CH2)8- | 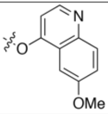 | 1.88 ± 0.04 | 1.68 ± 0.02 | > 50 | > 50 |
| **4h** | .. | -(CH2)8- | 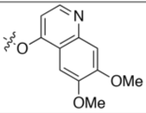 | 15.3 ± 0.1 | 12.2 ± 2.2 | > 50 | > 50 |
| **4i** | .. | -(CH2)8- | 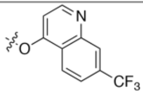 | 0.634 ± 0.062 | 0.491 ± 0.106 | 13.0 ± 2.2 | 27.8 ± 1.4 |
| **4j** | .. | -(CH2)8- | 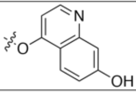 | 21.4 ± 3.8 | 15.8 ± 1.1 | > 50 | > 50 |
| **5a** | 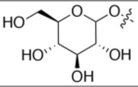 | -(CH2)8- | 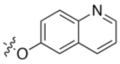 | > 50 | 33.4 ± 0.1 | 17.1 ± 1.5 | > 50 |
| **5b** (TH-PF03) | 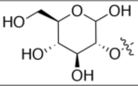 | -(CH2)8- | 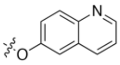 | 0.165 ± 0.03 | 0.141 ± 0.002 | 17.7 ± 2.9 | > 50 |
| **5c** | 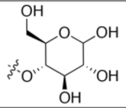 | -(CH2)8- | 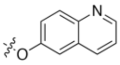 | 34.3 ± 2.5 | 25.1 ± 3.8 | 39.5 ± 2.1 | > 50 |
| **5d** | 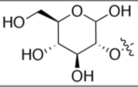 | -(CH2)8- | 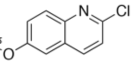 | 0.685 ± 0.077 | 0.718 ± 0.078 | 9.56 ± 0.39 | > 50 |
| **5e** | 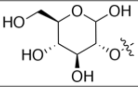 | -(CH2)9- | 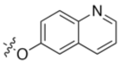 | 2.91 ± 0.40 | 3.08 ± 0.30 | 8.29 ± 0.77 | > 50 |
| **DHA** | 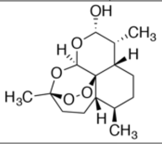 | | | 0.00480 ± 0.00075 | 0.00377 ± 0.00106 | - | - |
| **Mefloquine** | 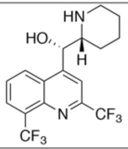 | | | 0.0179 ± 0.0036 | 0.0194 ± 0.0040 | - | - |
| **Quinine** | 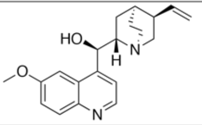 | | | 0.0690 ± 0.0017 | 0.441 ± 0.038 | - | - |

### NMR Spectra

$^{1}$H NMR of Compound **1** in $CDCl_3$

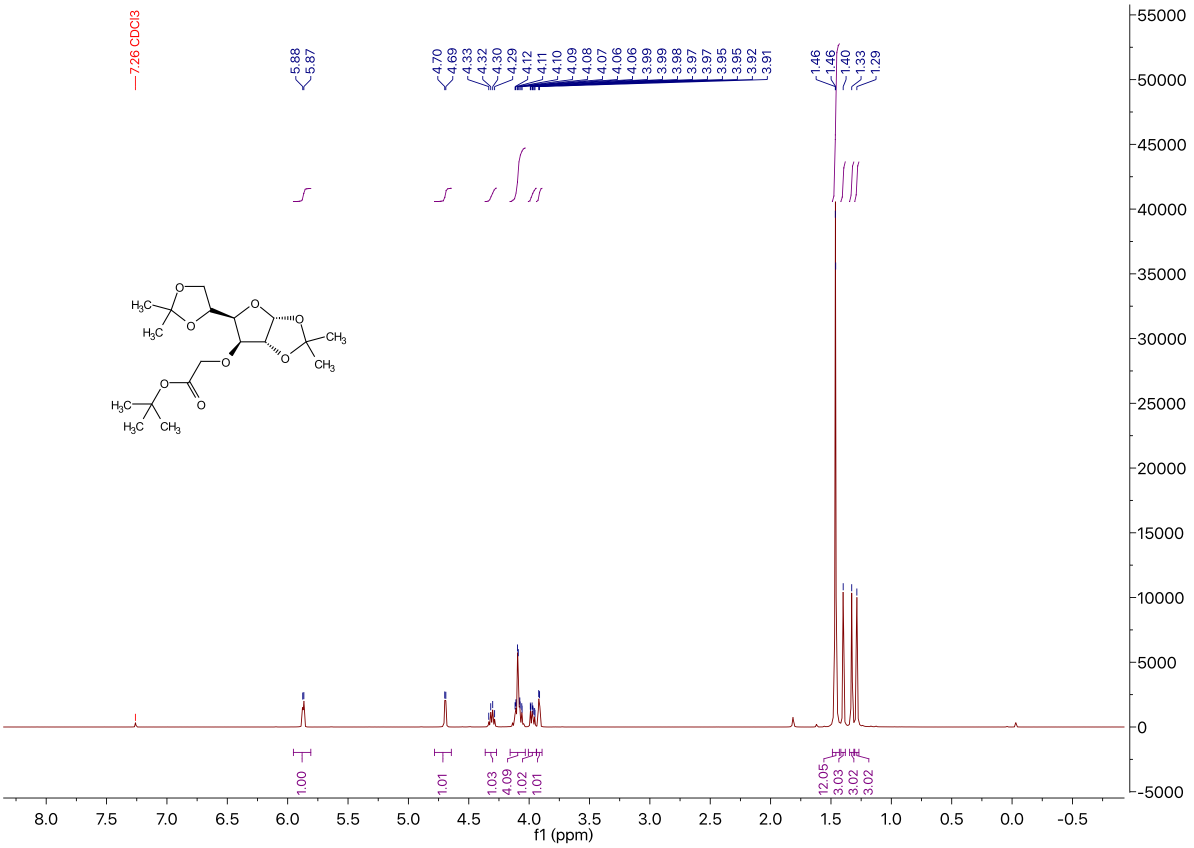

### $^{13}C$ NMR of Compound **1** in $CDCl_3$

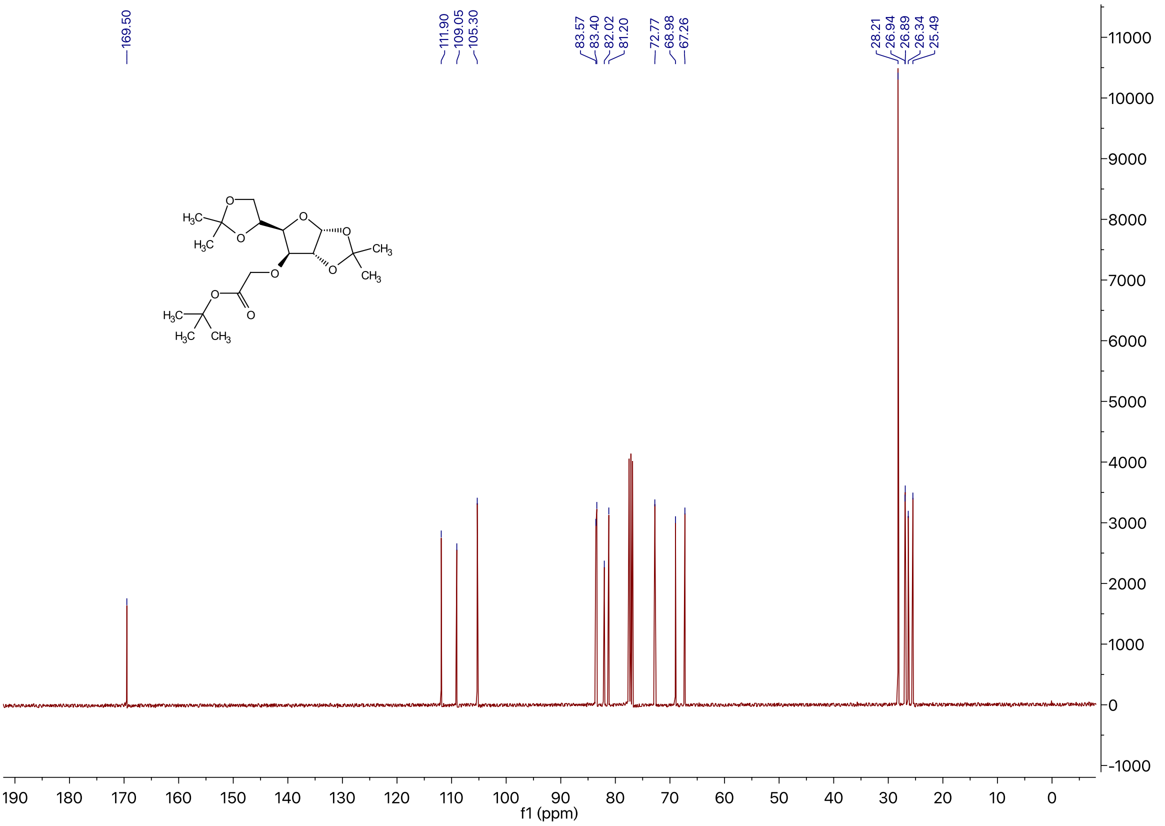

### $^{1}H$ NMR of Compound **2** in DMSO-$d_6$

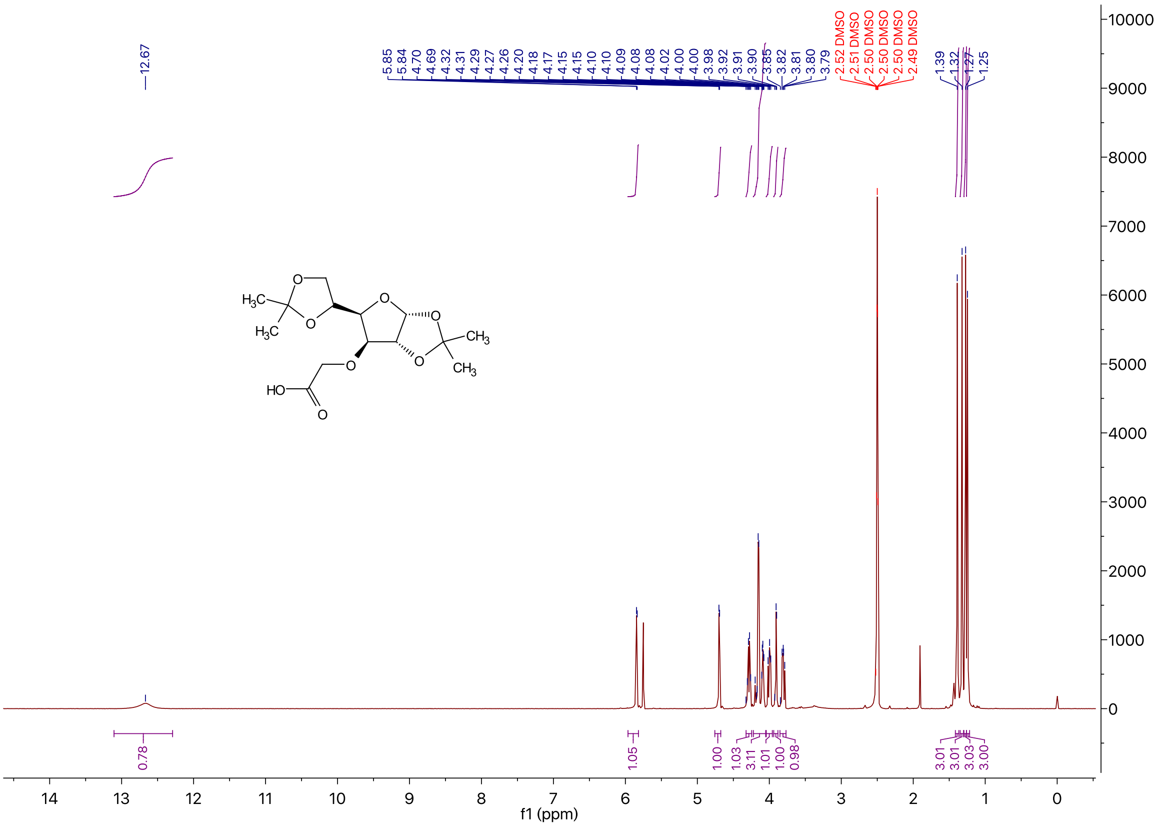

$^{1}H$ NMR of Compound **3** in $CDCl_3$

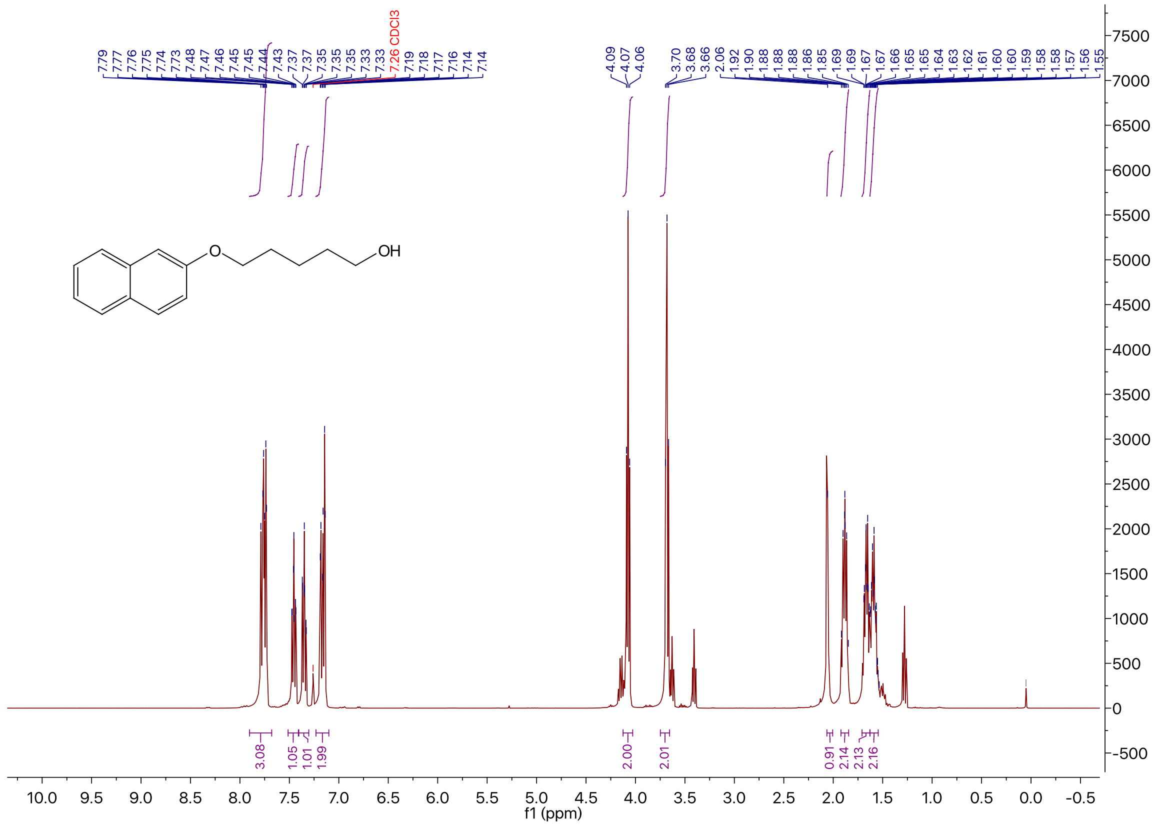

$^{13}C$ NMR of Compound **3** in $CDCl_3$

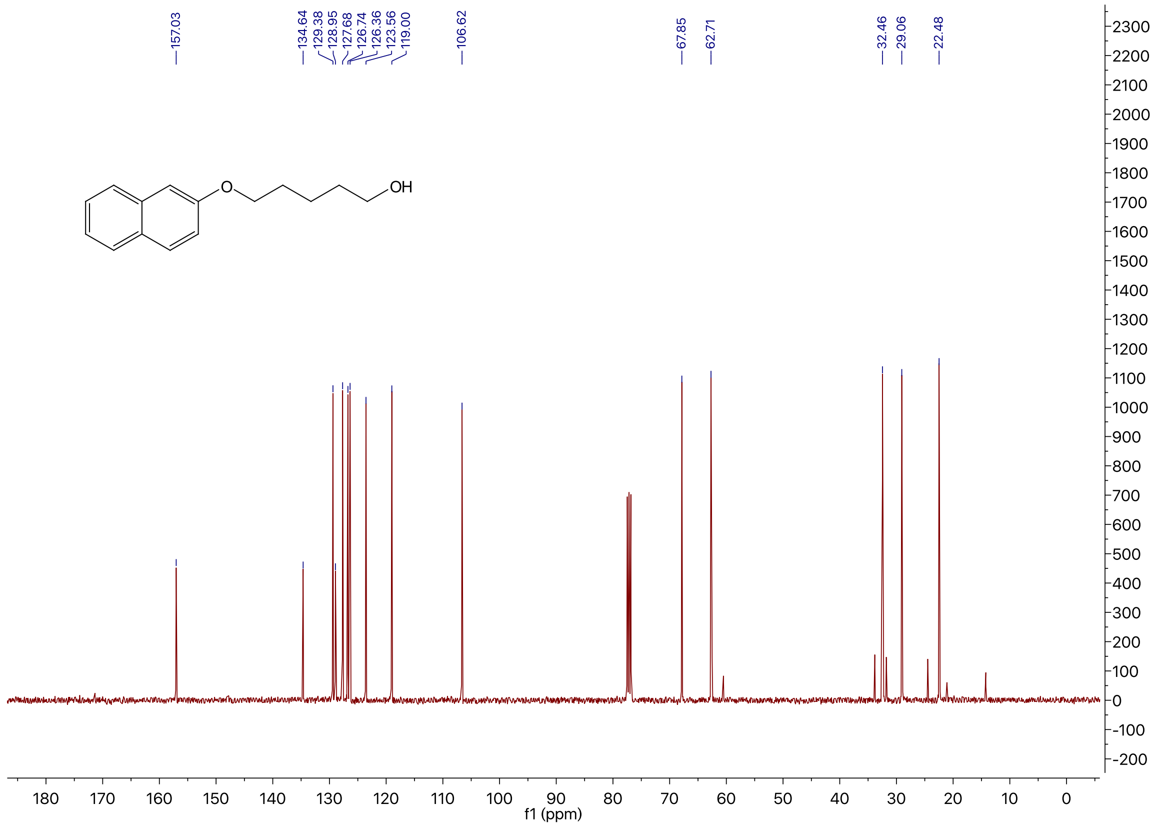

[1]H NMR of Compound **4** in $CDCl_3$

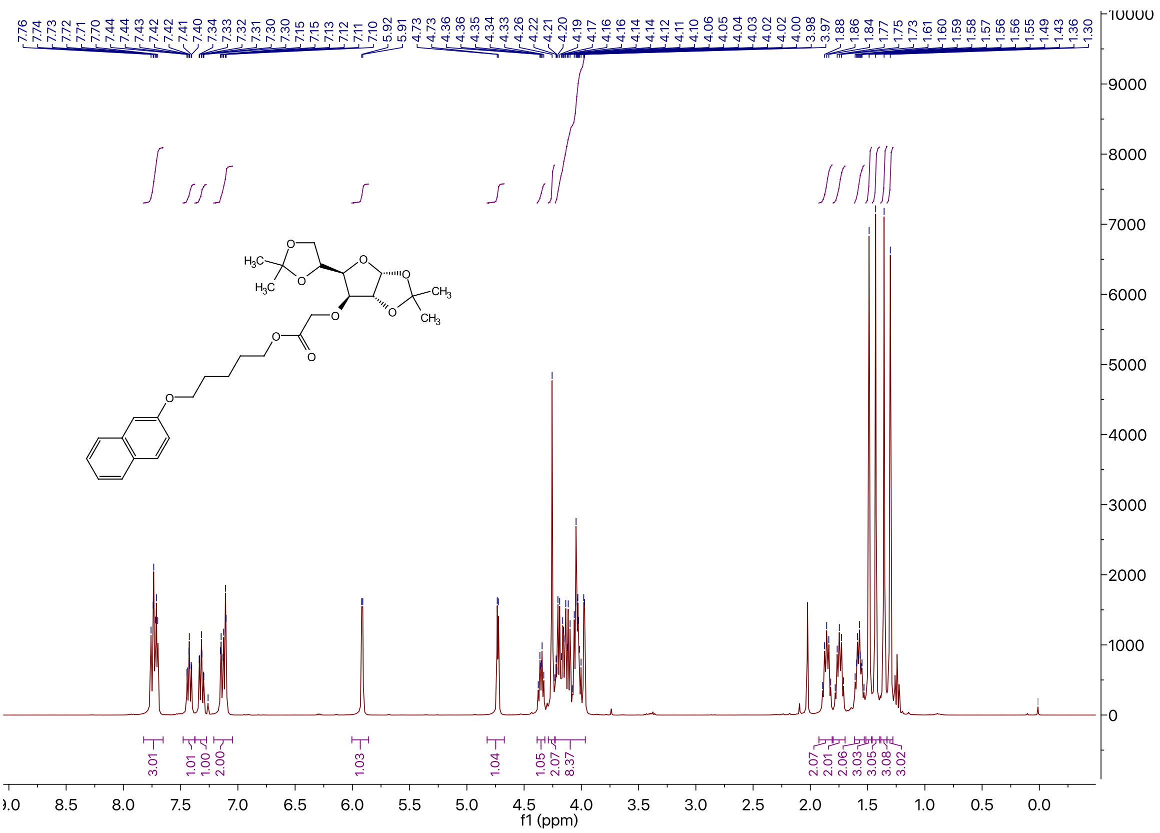

[13]C NMR of Compound **4** in $CDCl_3$

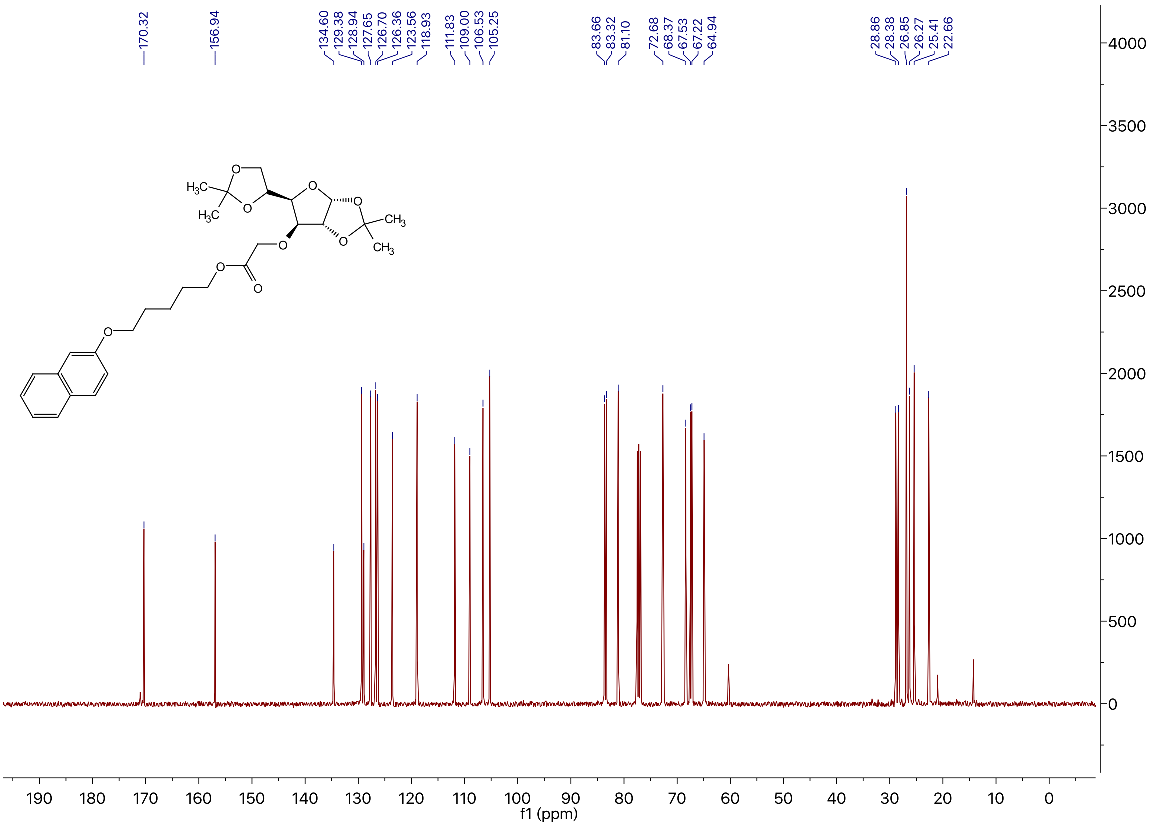

[1]H NMR of Compound **1a** in DMSO-*d*$_6$

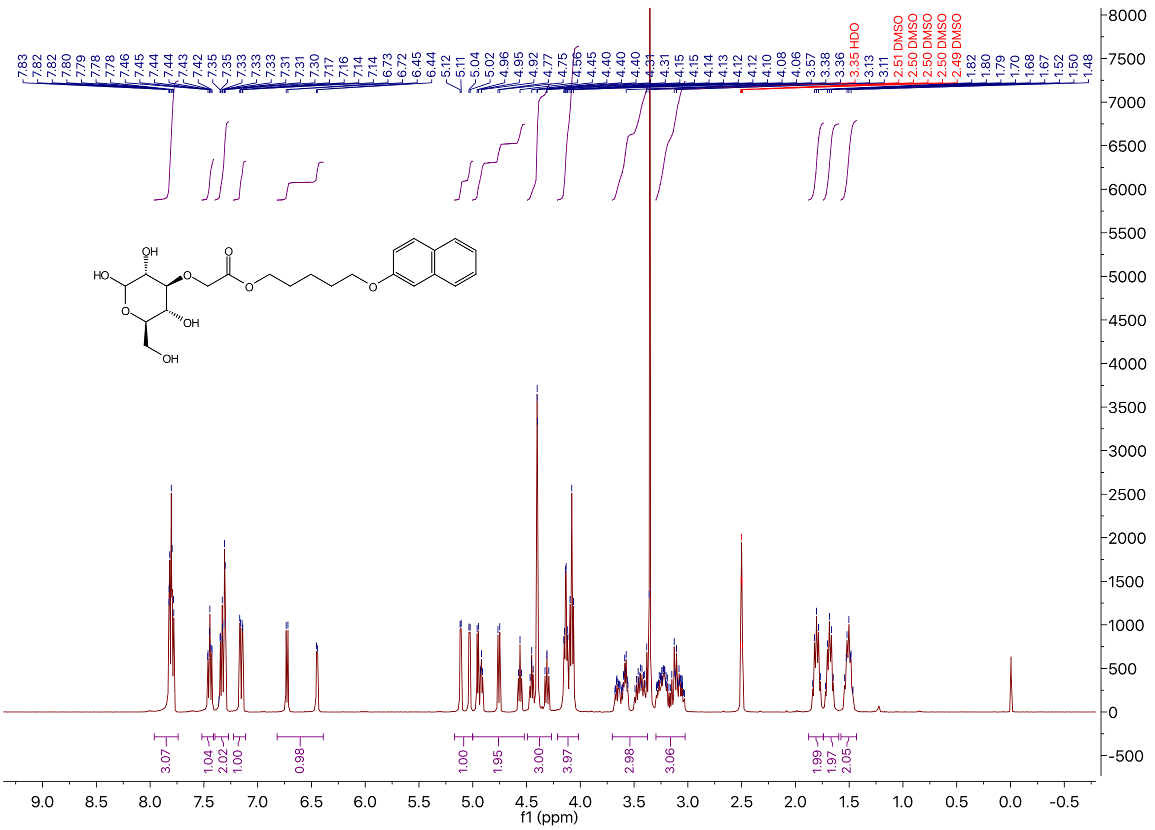

[13]C NMR of Compound **1a** in DMSO-*d*$_6$

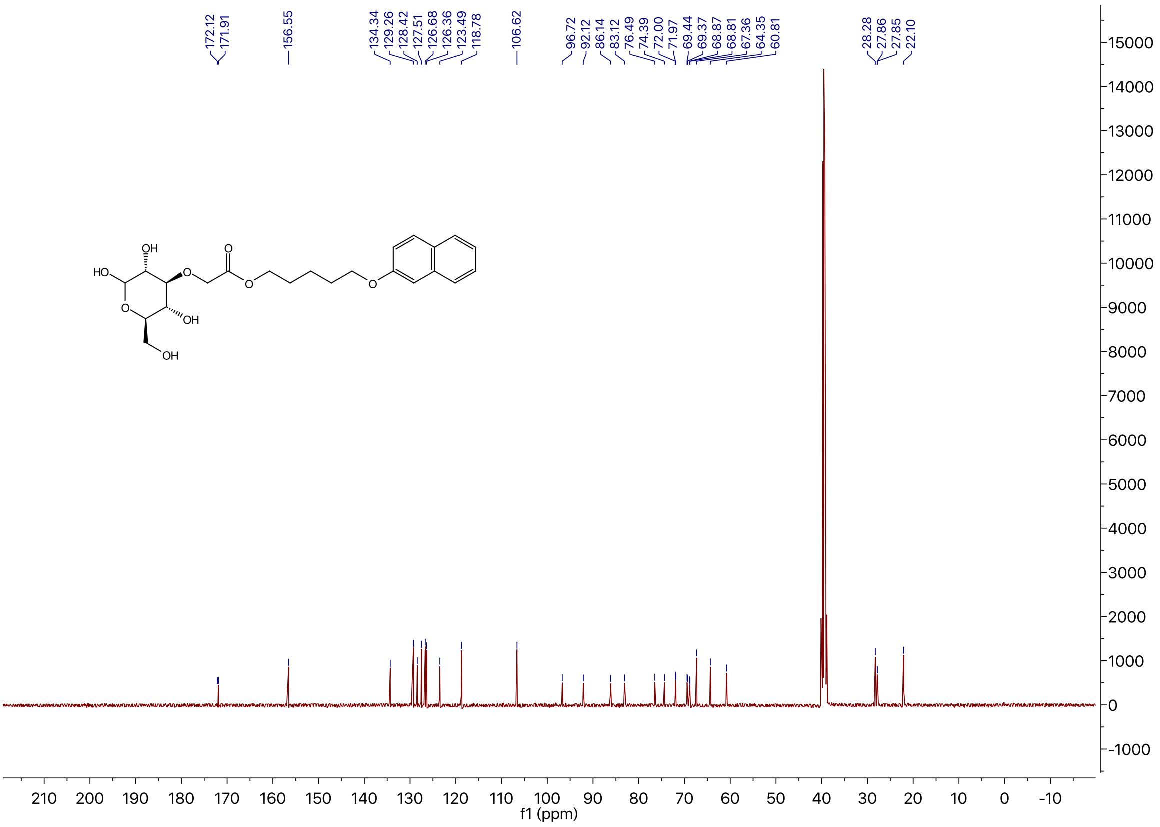

$^{1}$H NMR of Compound **1b** in DMSO-$d_6$

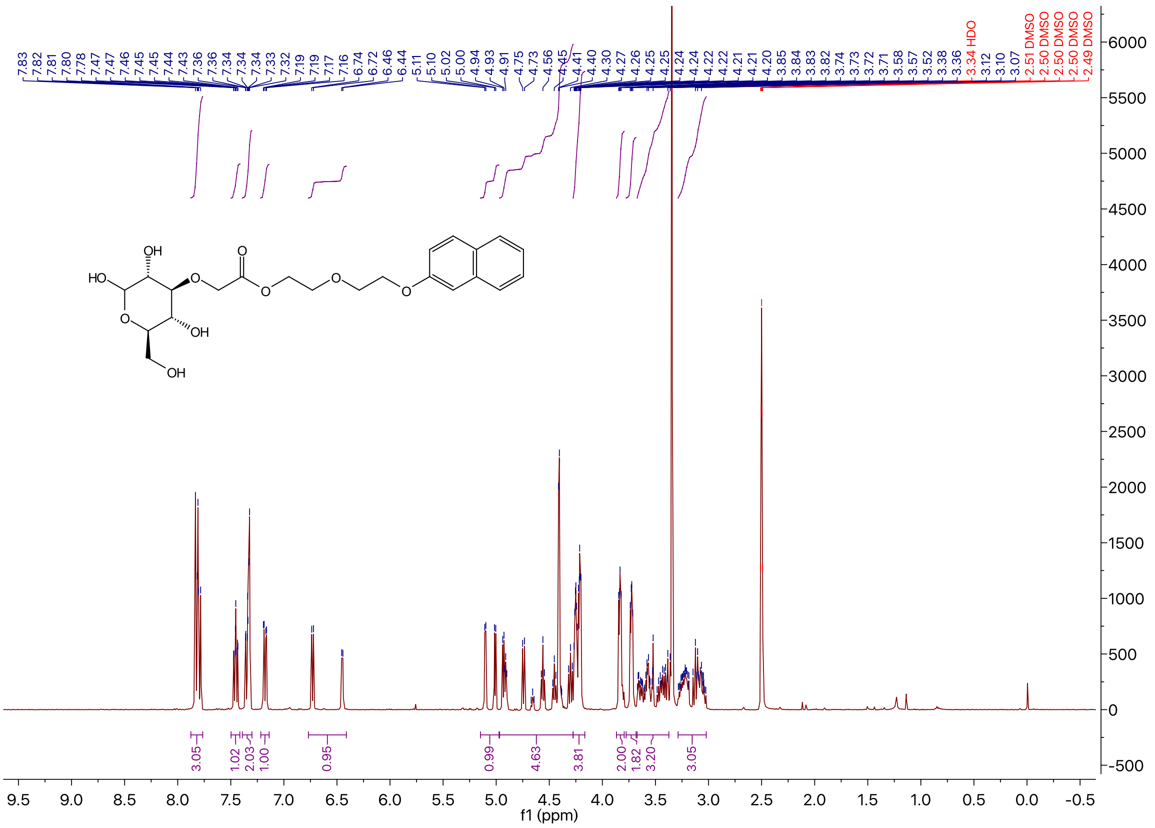

$^{13}$C NMR of Compound **1b** in DMSO-$d_6$

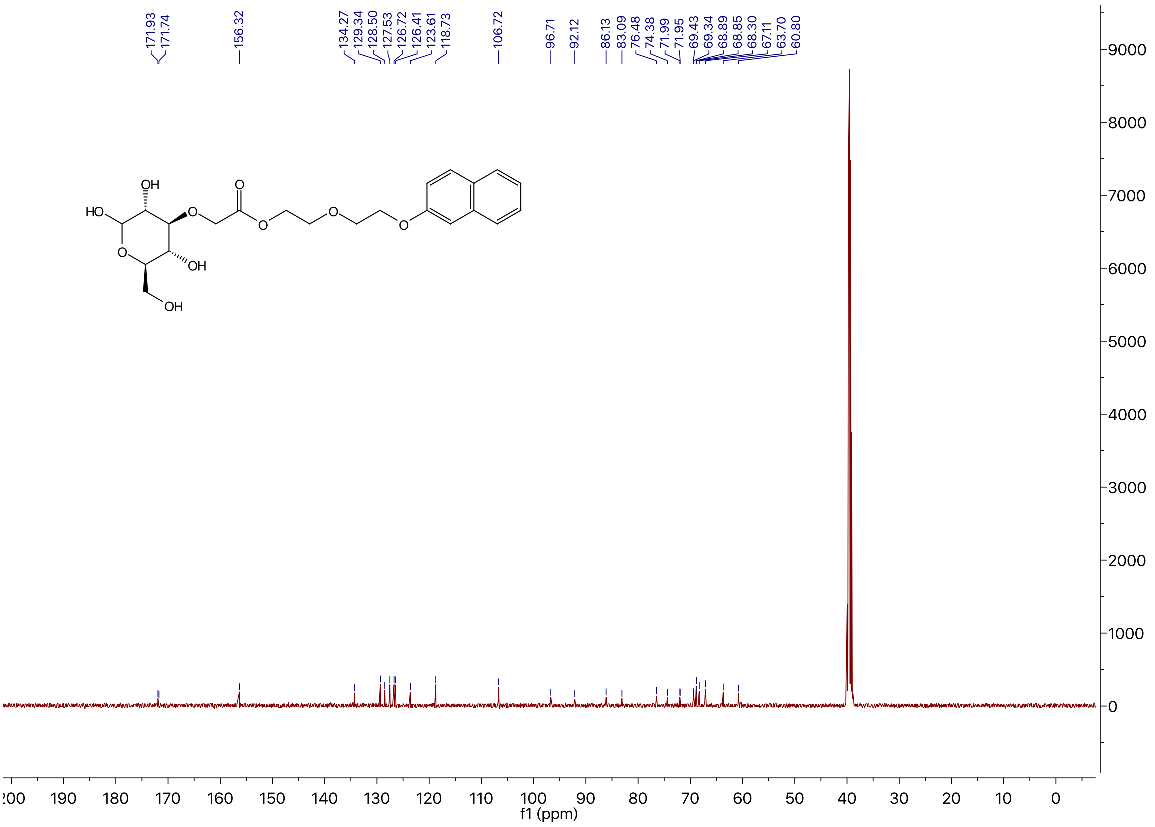

[1]H NMR of Compound **1d** in DMSO-$d_6$

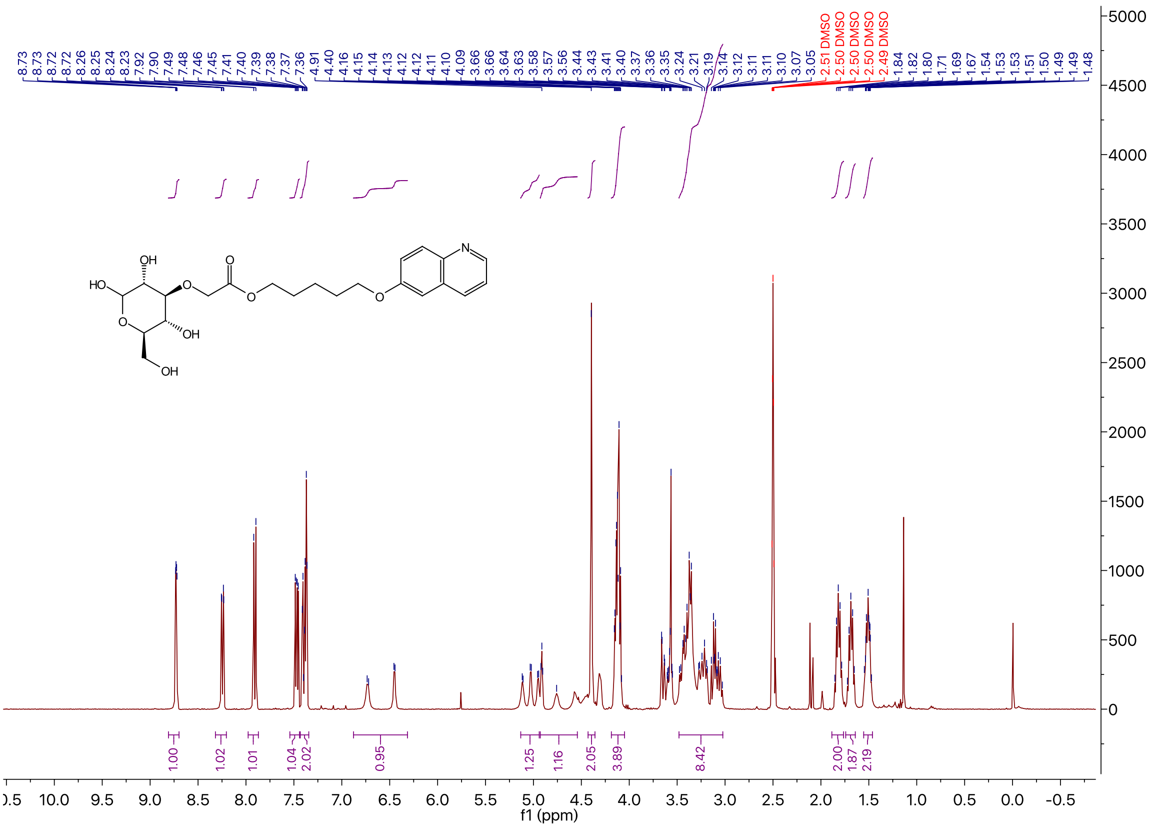

[13]C NMR of Compound **1d** in DMSO-$d_6$

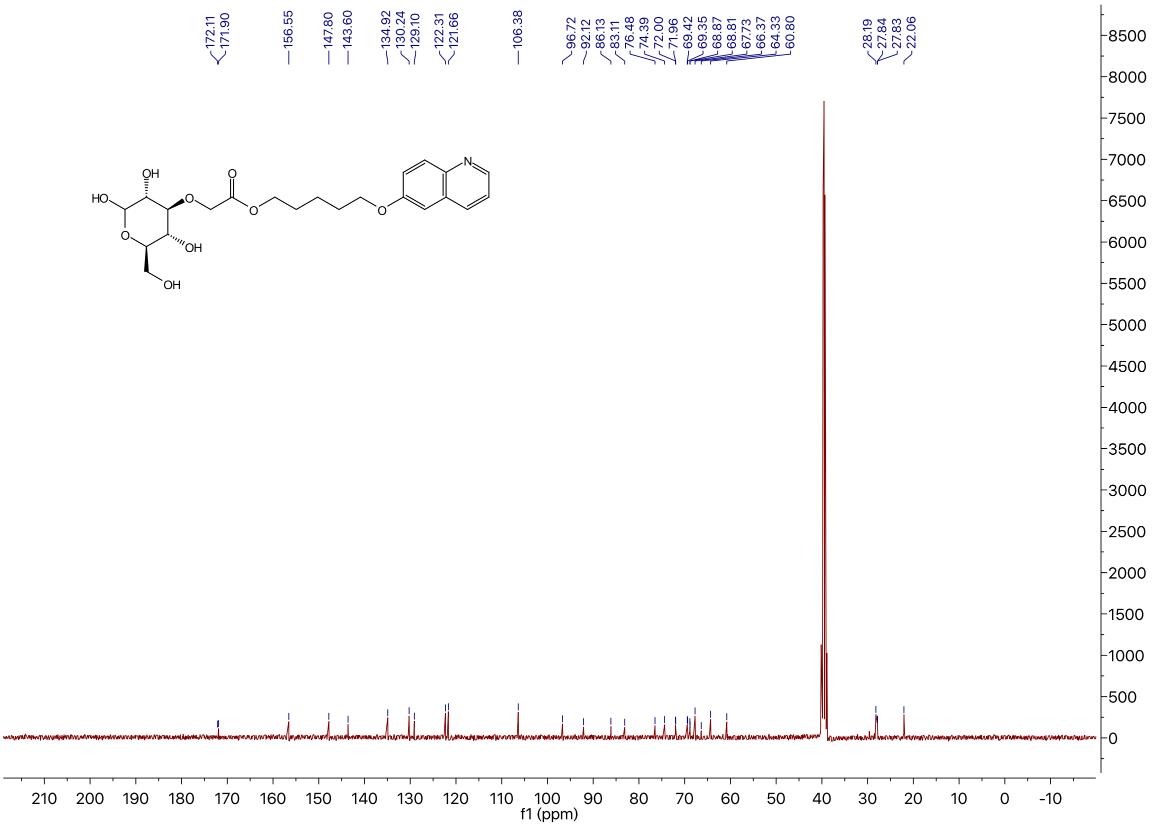

$^1$H NMR of Compound **5** in $CDCl_3$

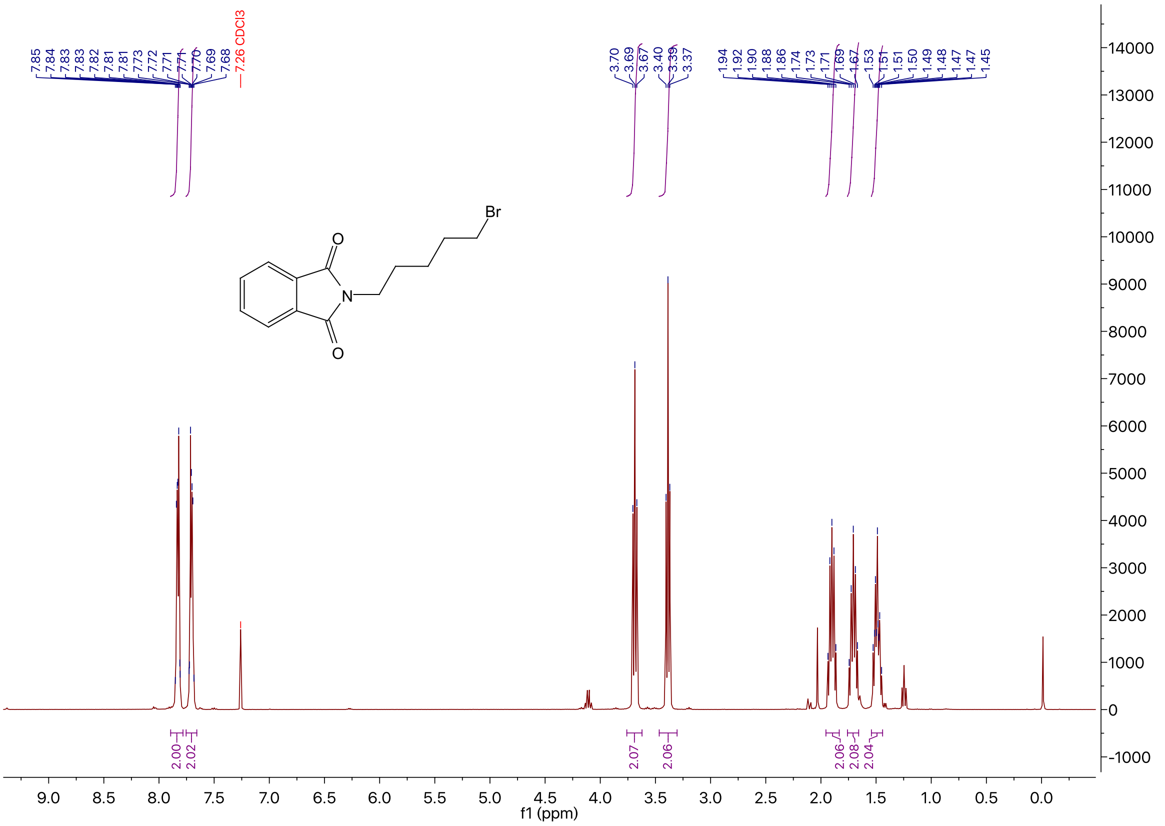

$^1$H NMR of Compound **6** in $CDCl_3$

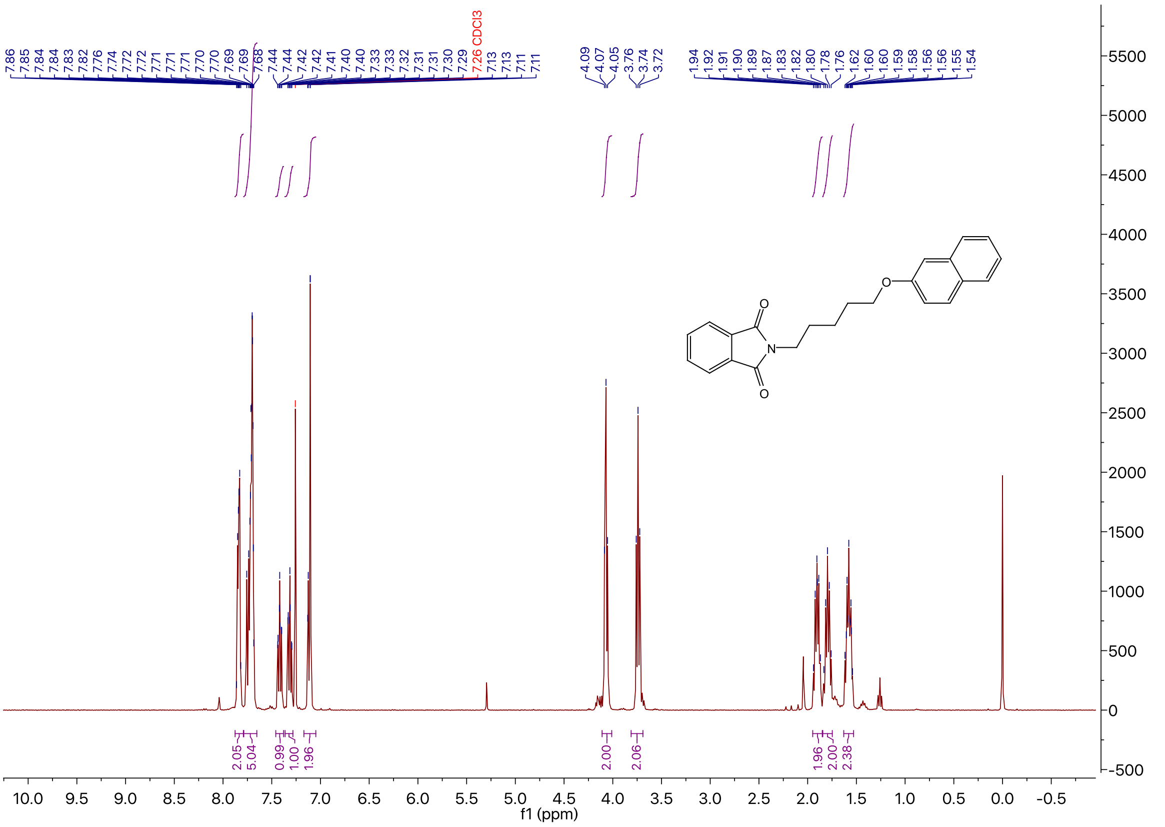

$^1$H NMR of Compound **7** in DMSO-$d_6$

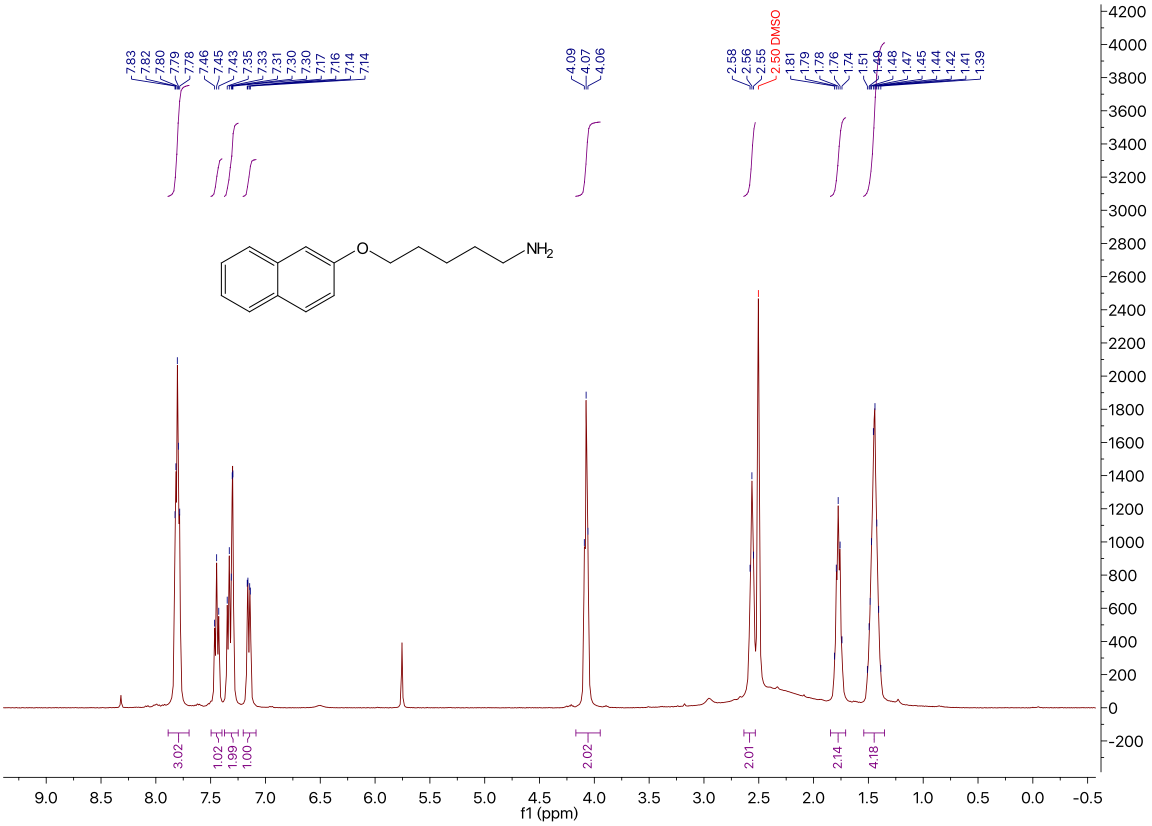

$^1$H NMR of Compound **8** in $CDCl_3$

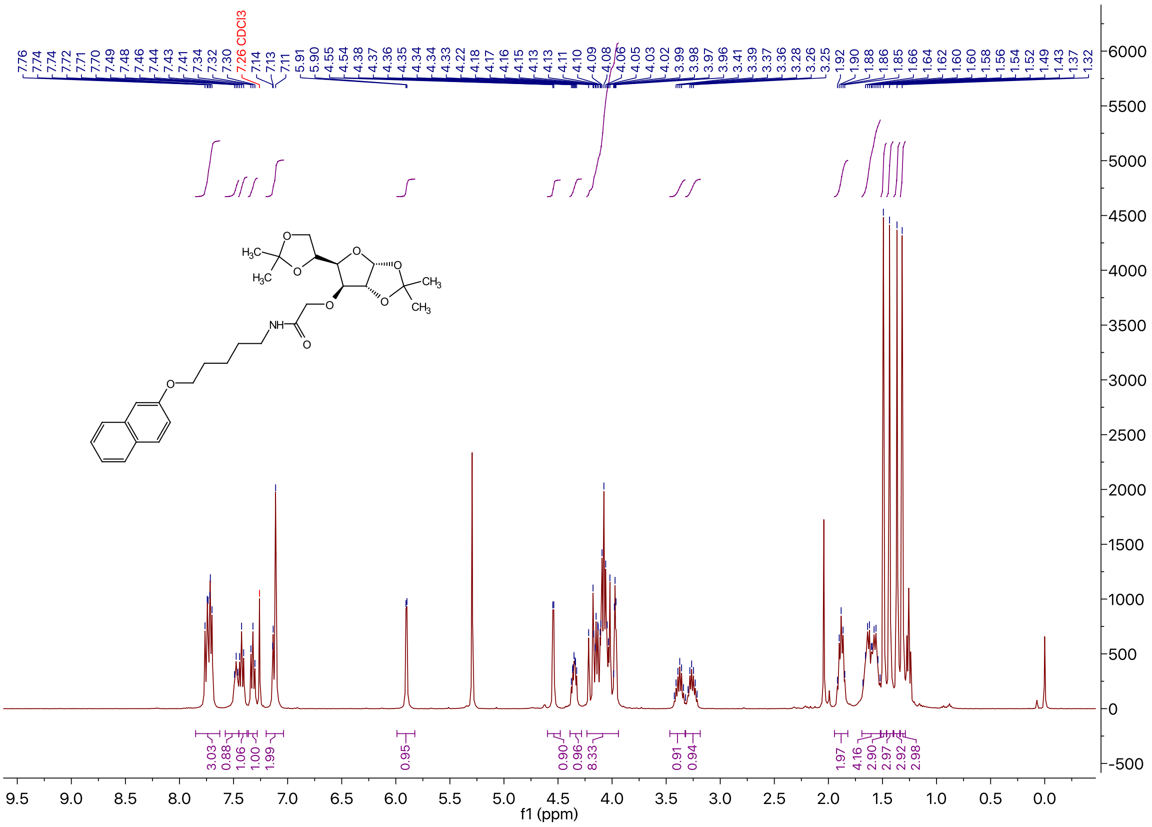

### $^{13}$C NMR of Compound **8** in $CDCl_3$

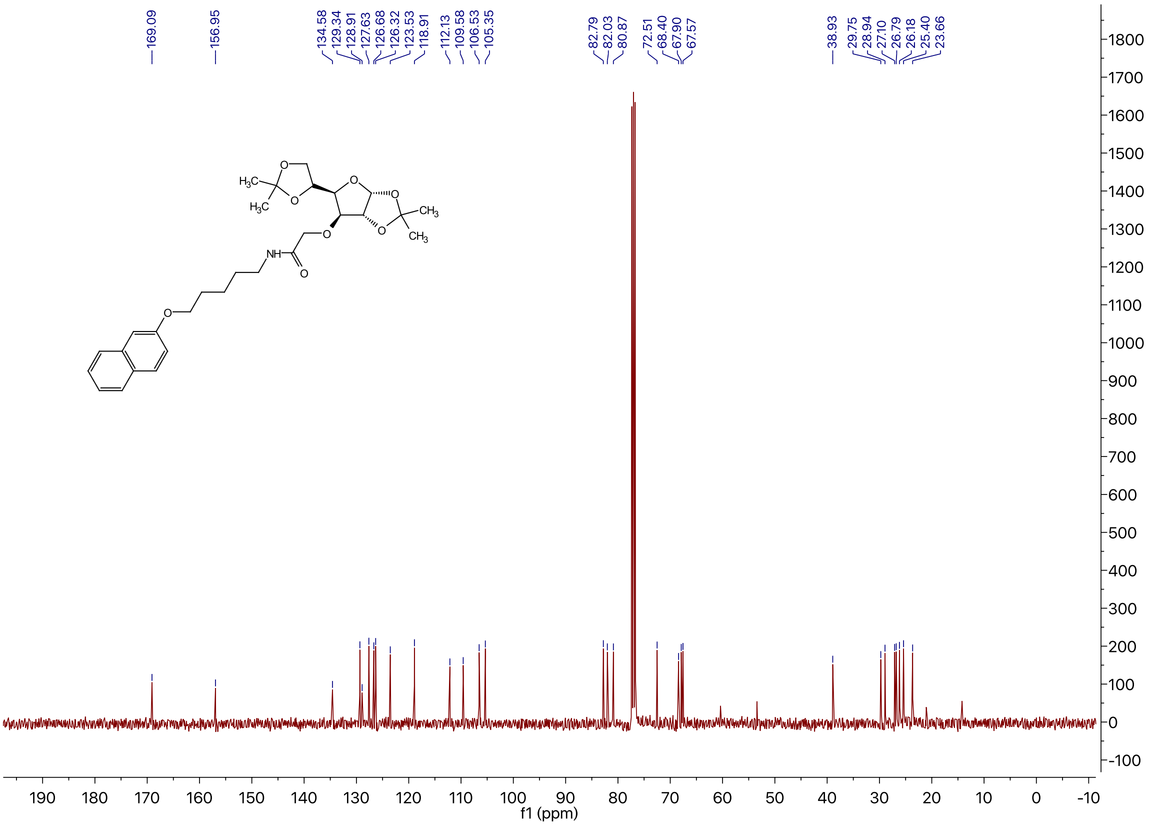

### $^{1}$H NMR of Compound **1c** in DMSO-$d_6$

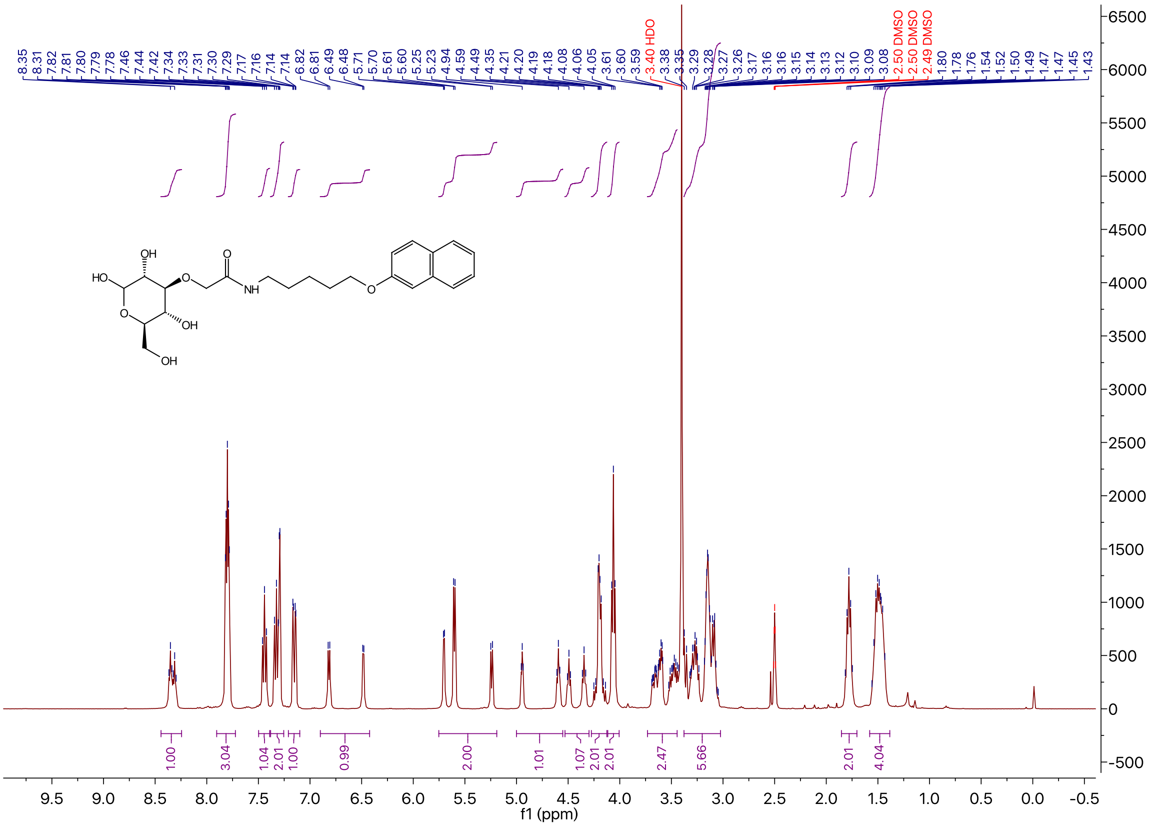

$^{13}C$ NMR of Compound **1c** in DMSO-$d_6$

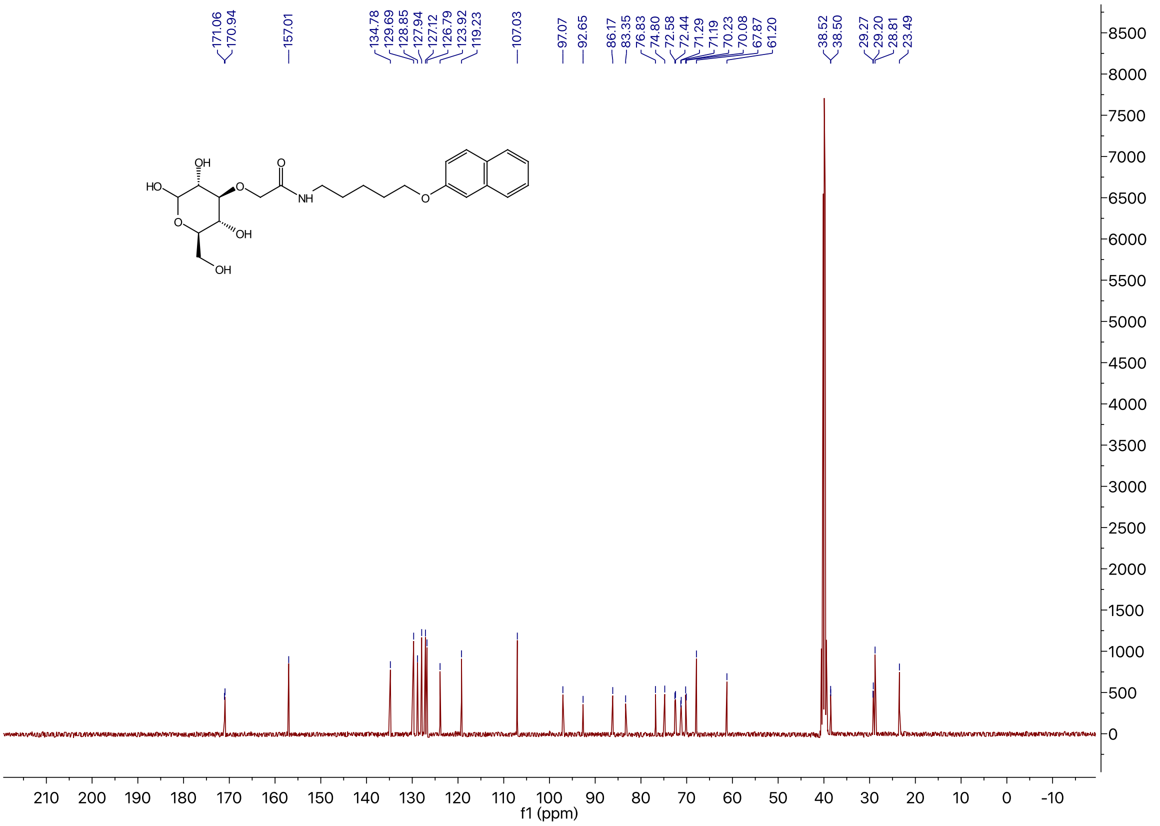

$^{1}H$ NMR of Compound **9** in $CDCl_3$

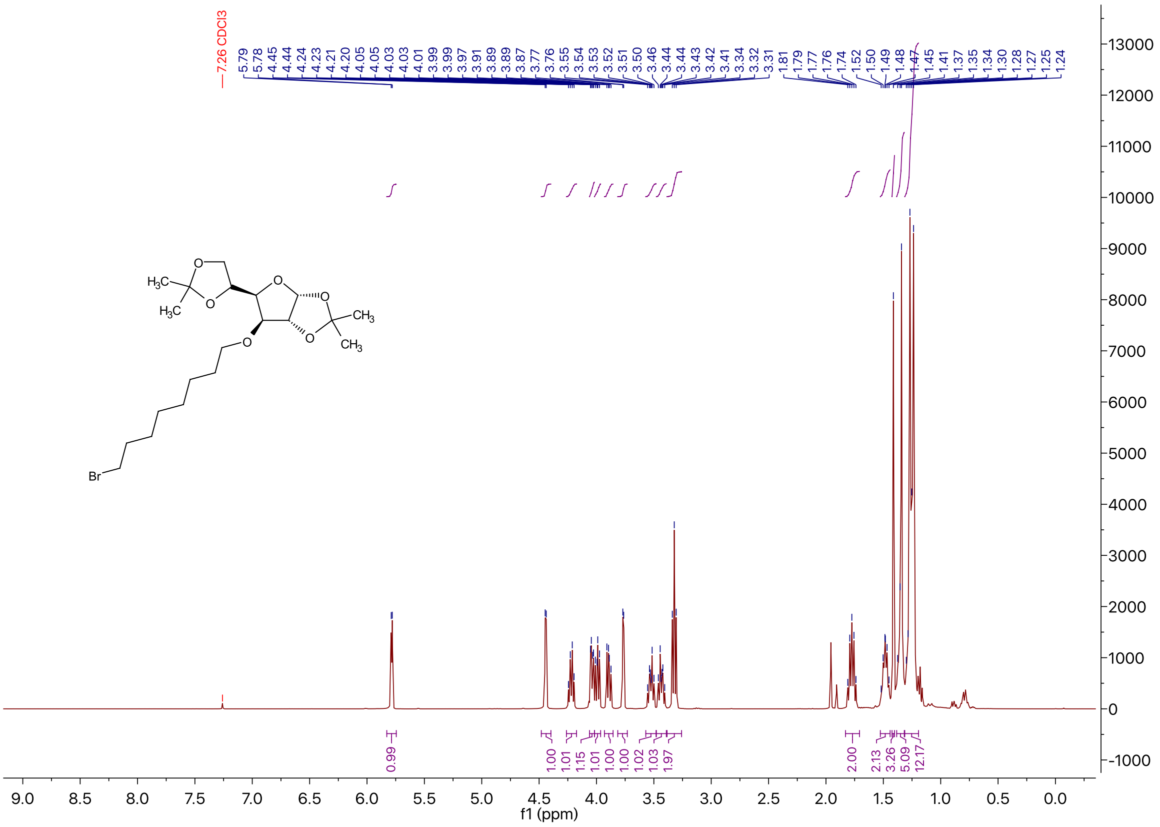

$^{13}C$ NMR of Compound **9** in $CDCl_3$

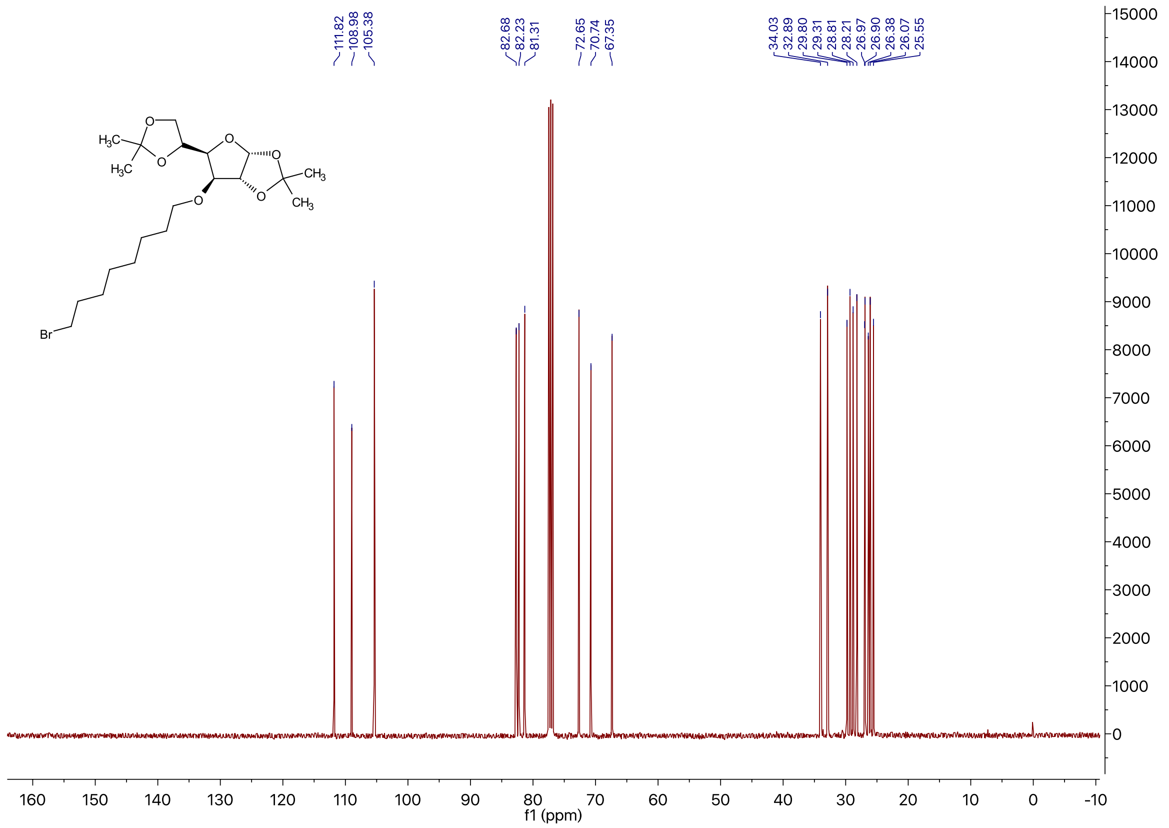

$^{1}H$ NMR of Compound **10** in $CDCl_3$

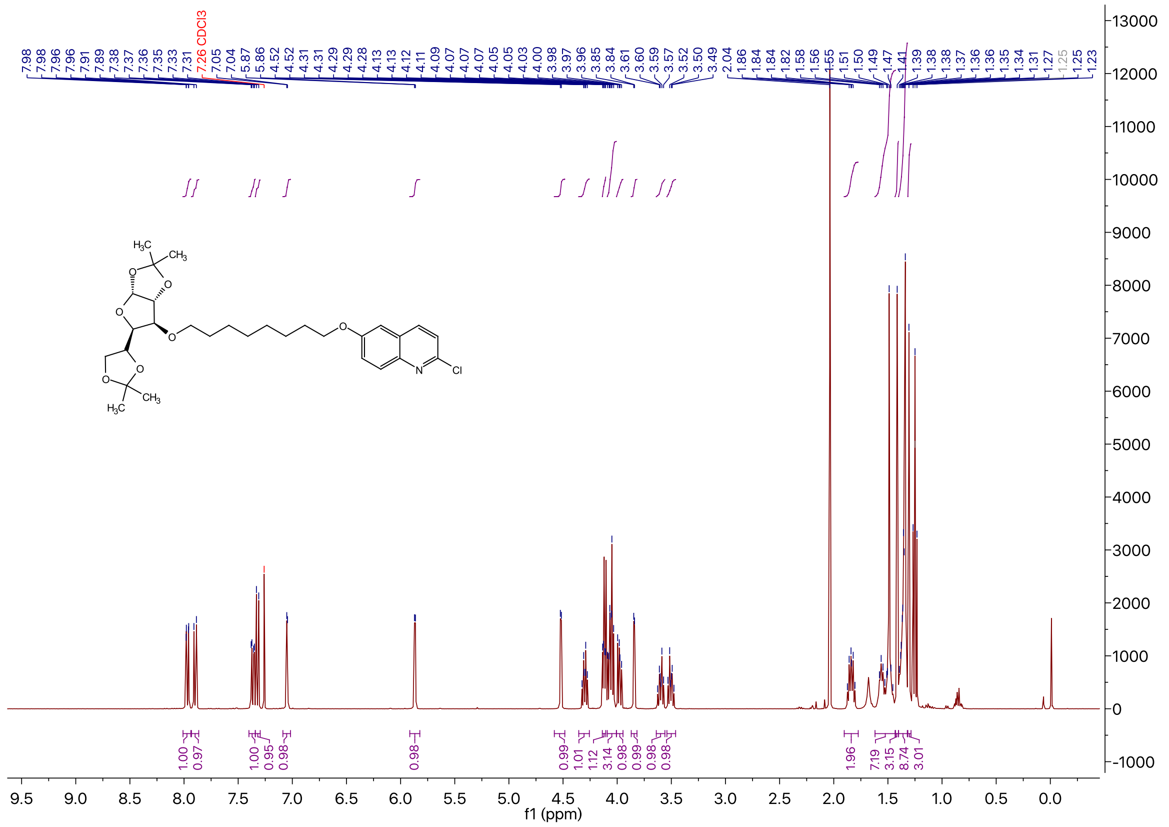

<sup></sup>

### $^{13}$C NMR of Compound **10** in $CDCl_3$

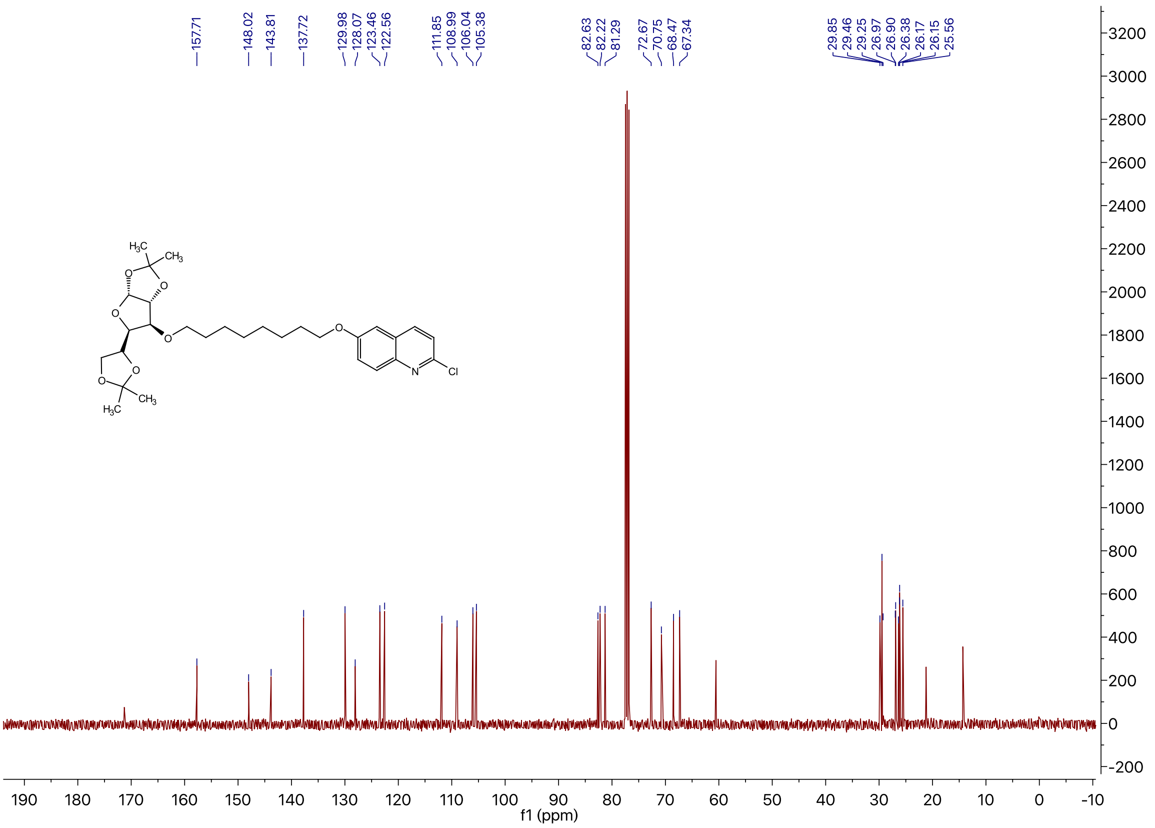

### $^{1}$H NMR of Compound **3a** in DMSO-$d_6$

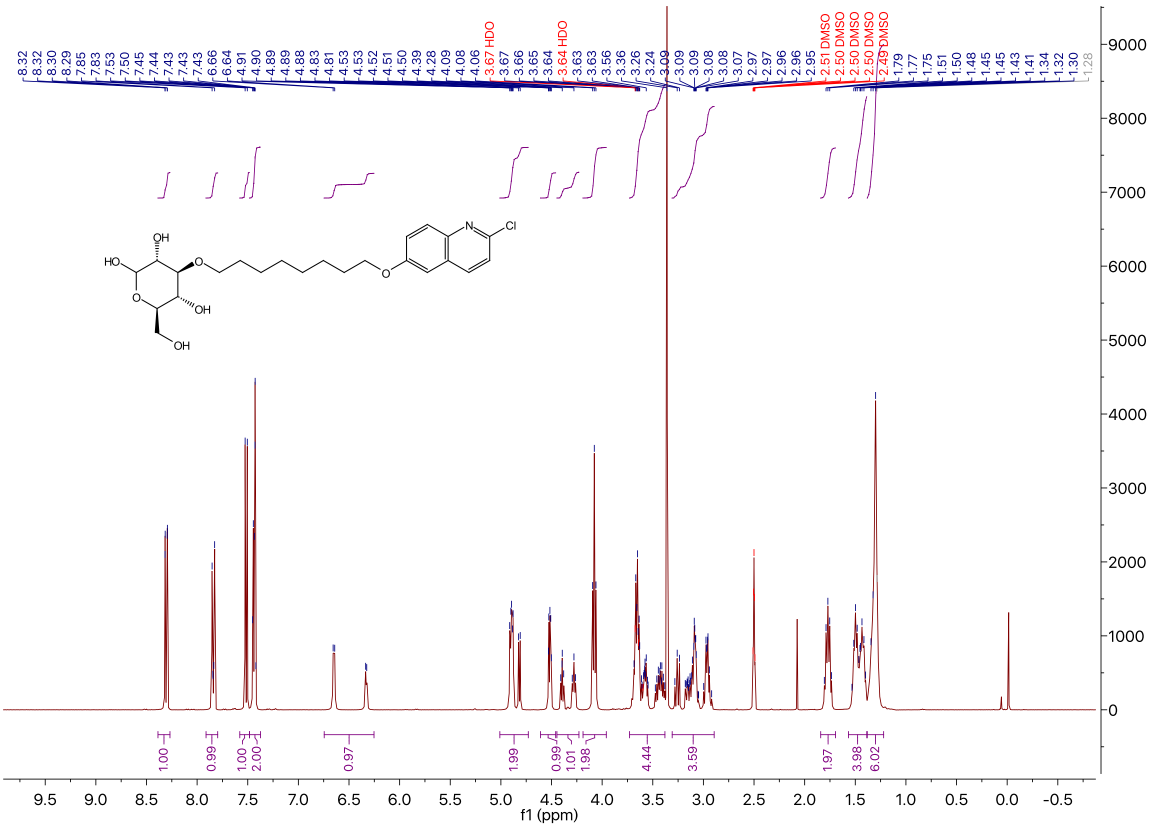

$^{13}$C NMR of Compound **3a** in DMSO-$d_6$

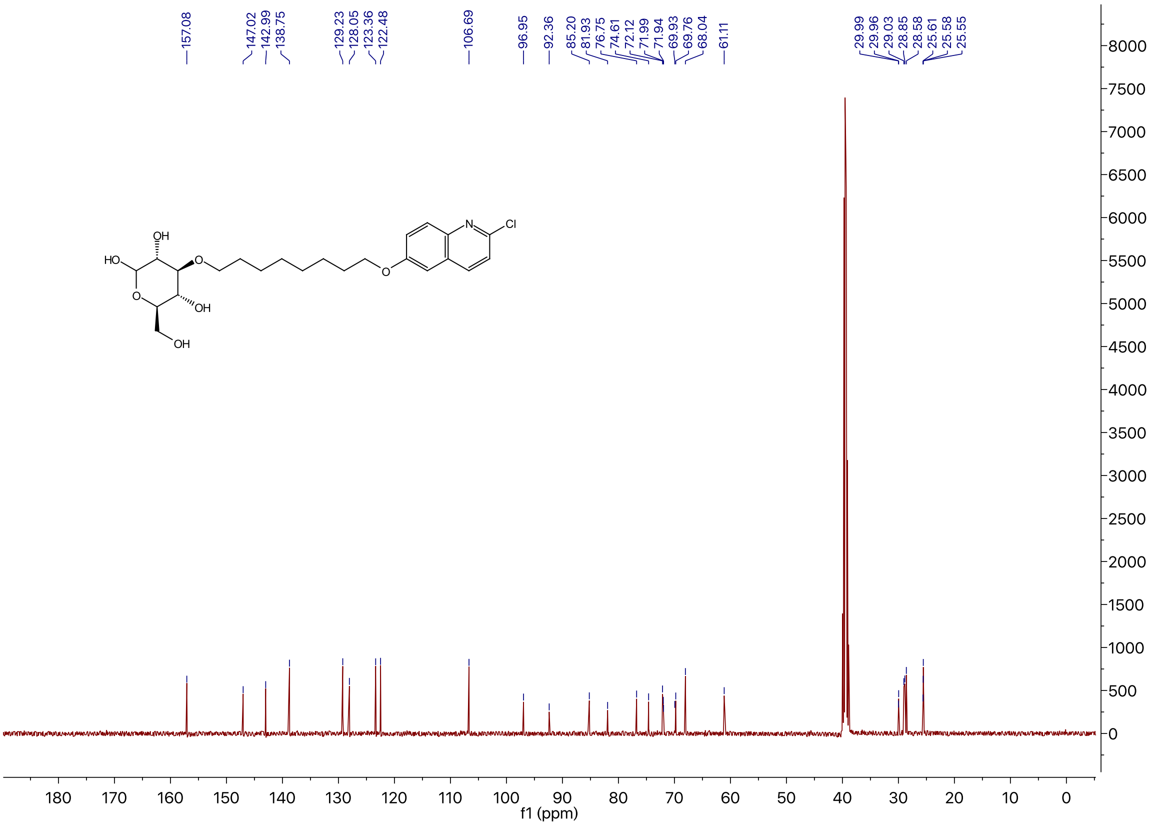

$^{1}$H NMR of Compound **1e** in DMSO-$d_6$

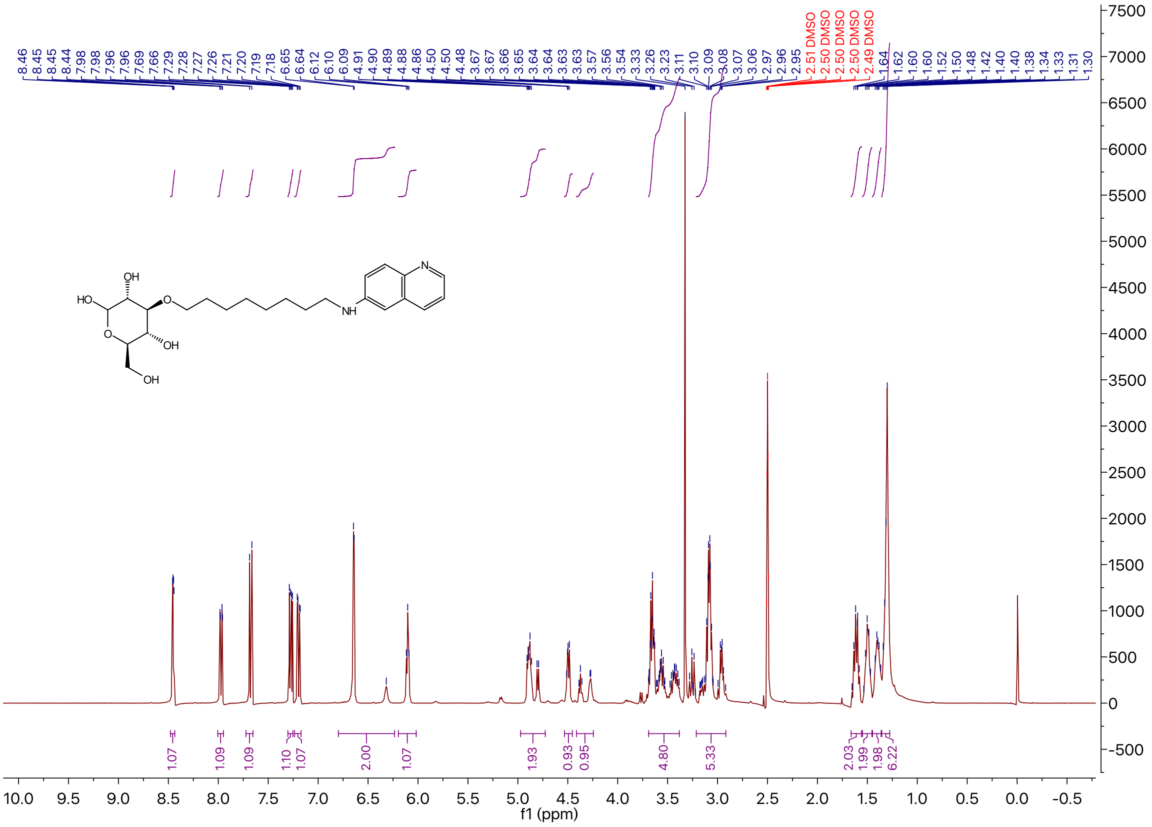

$^{13}C$ NMR of Compound **1e** in DMSO-$d_6$

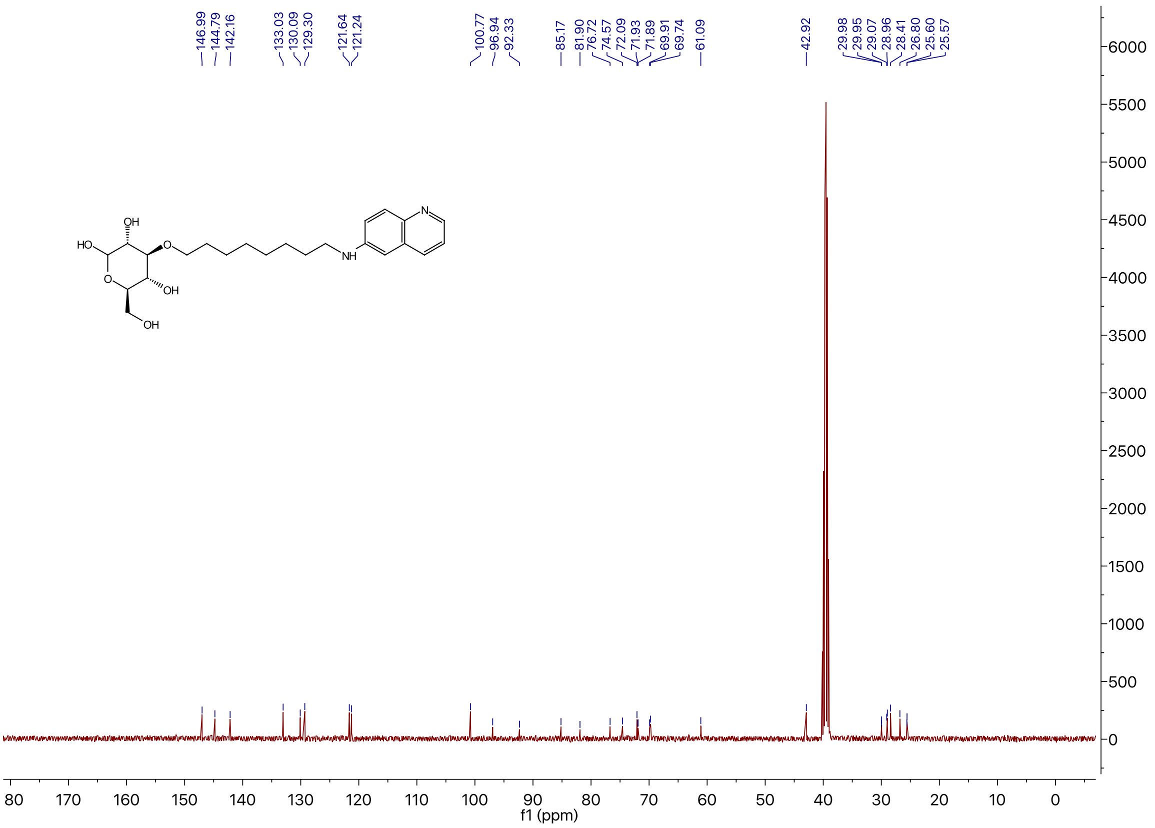

$^{1}H$ NMR of Compound **1f** in DMSO-$d_6$

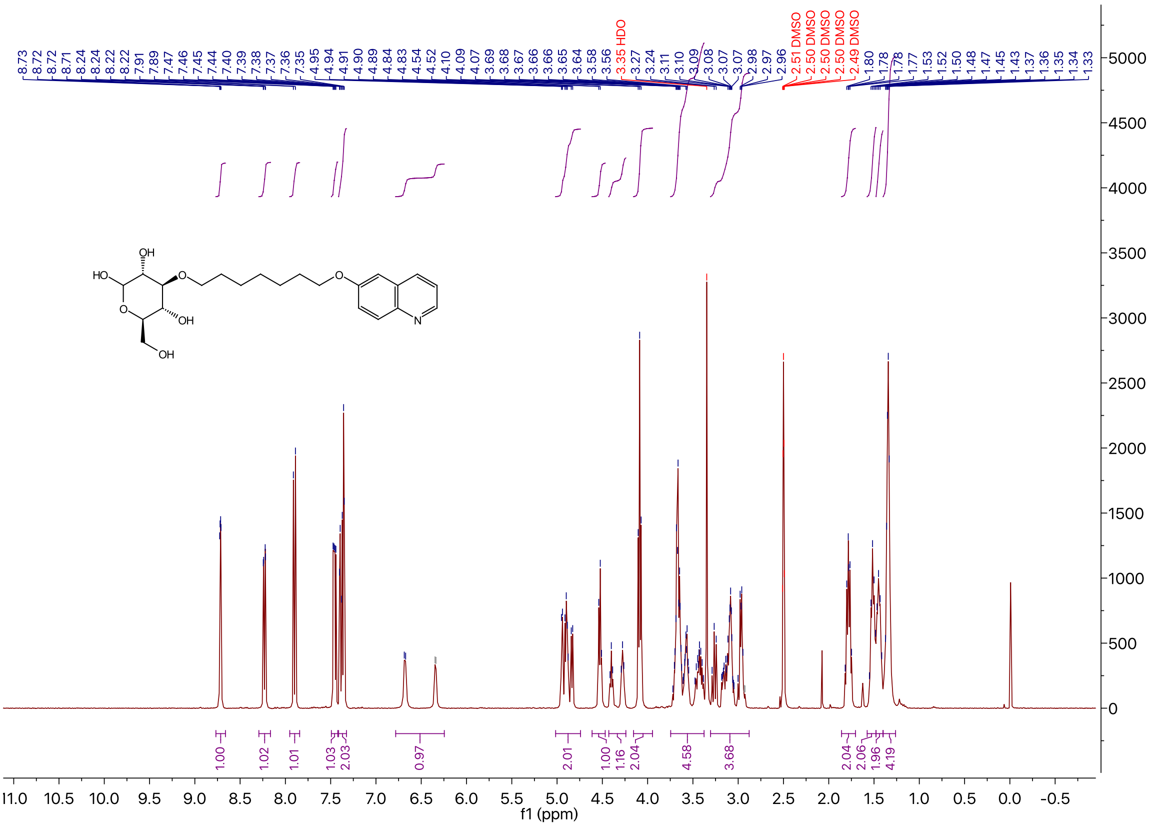

$^{13}$C NMR of Compound **1f** in DMSO-$d_6$

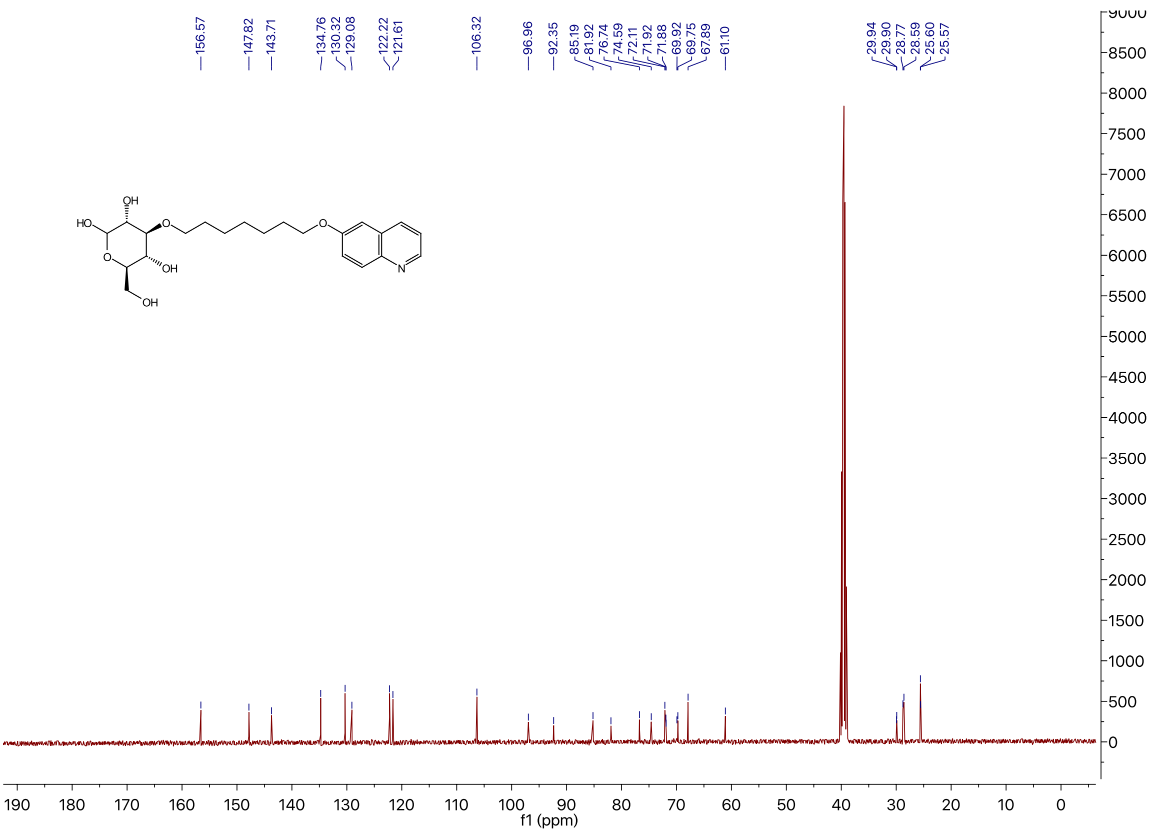

$^{1}$H NMR of Compound **1g** in DMSO-$d_6$

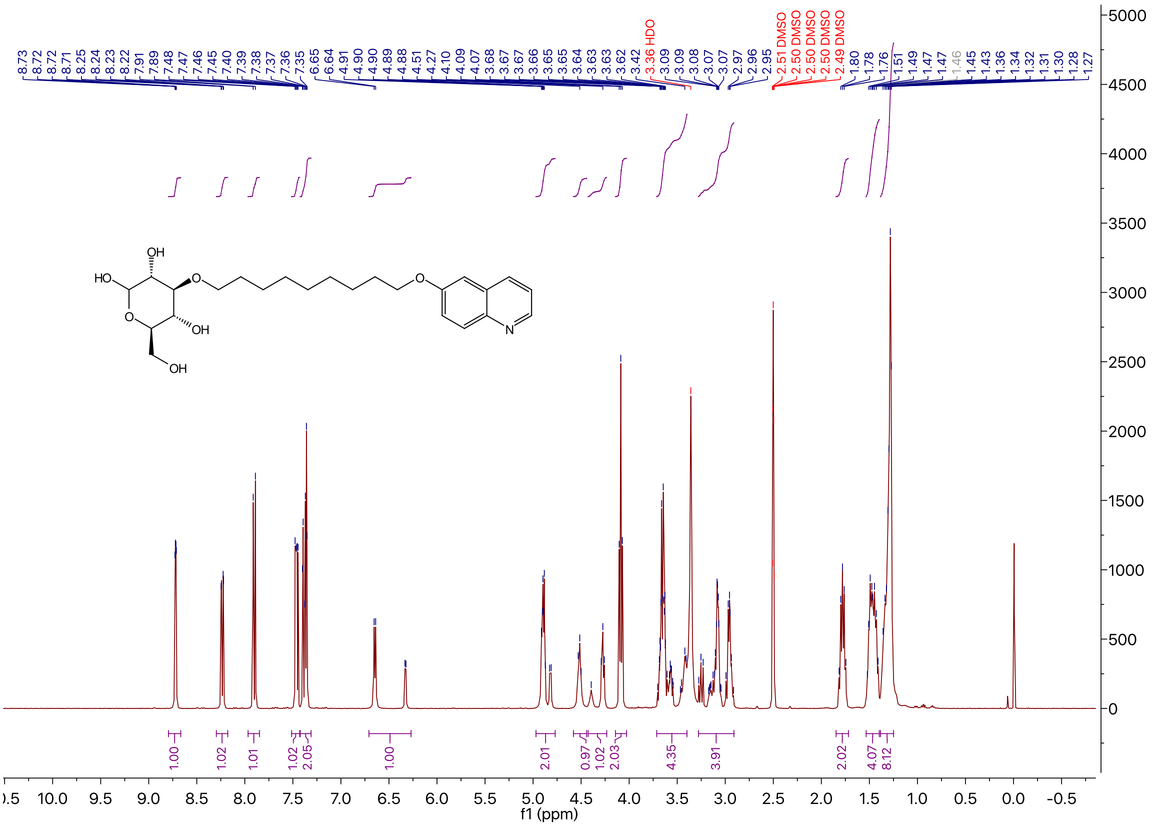

$^{13}C$ NMR of Compound **1g** in DMSO-$d_6$

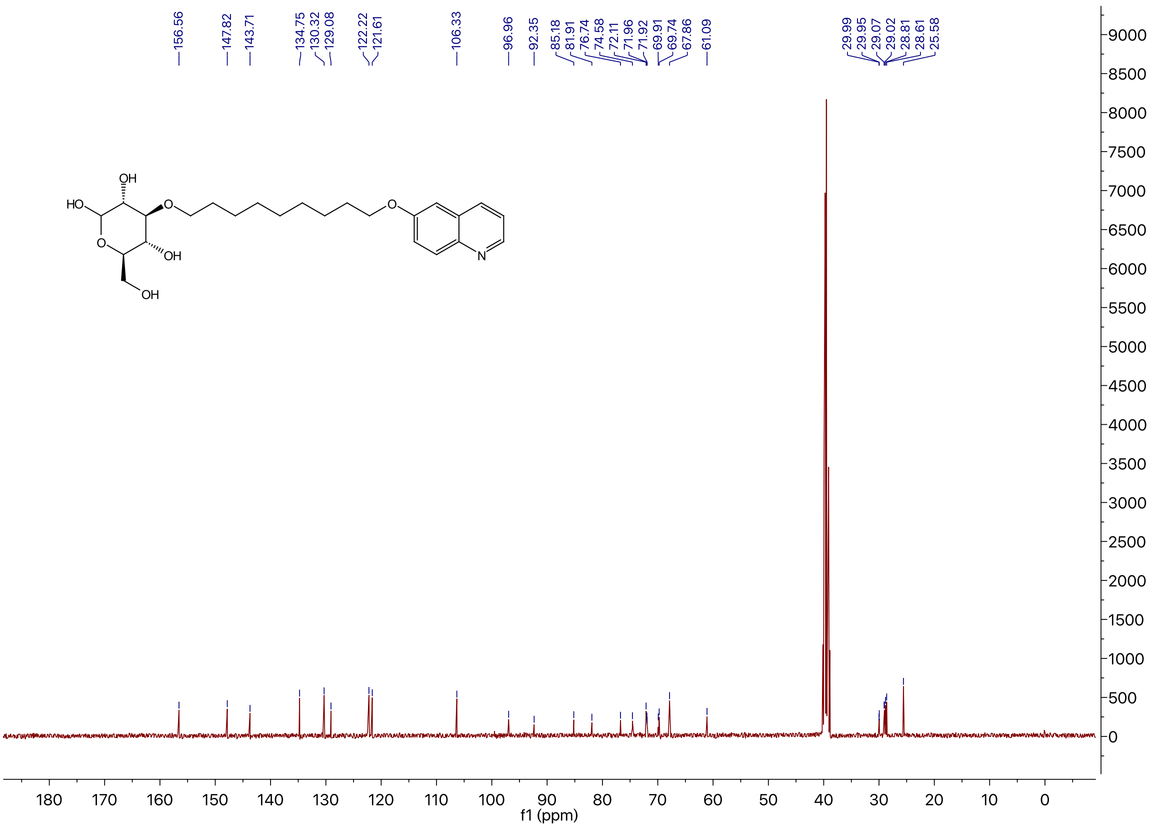

$^{1}H$ NMR of Compound **1h** in DMSO-$d_6$

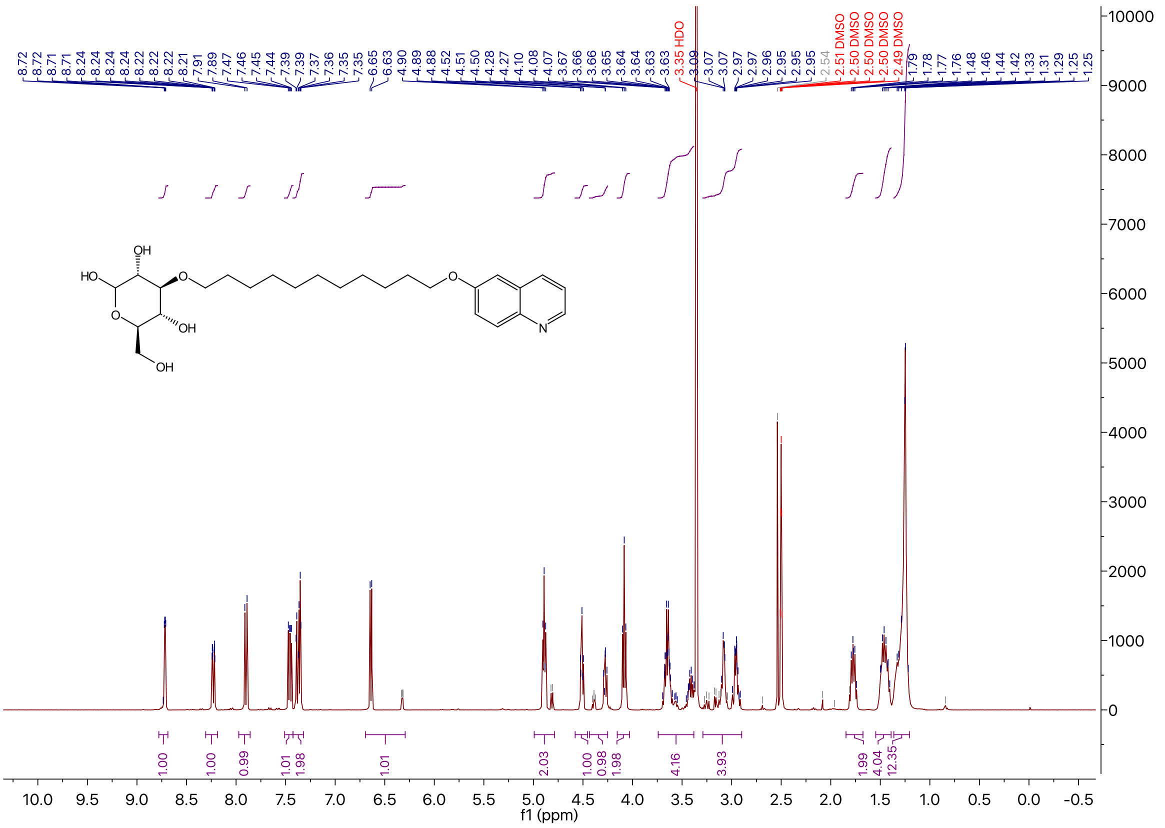

### $^{13}$C NMR of Compound **1h** in DMSO-$d_6$

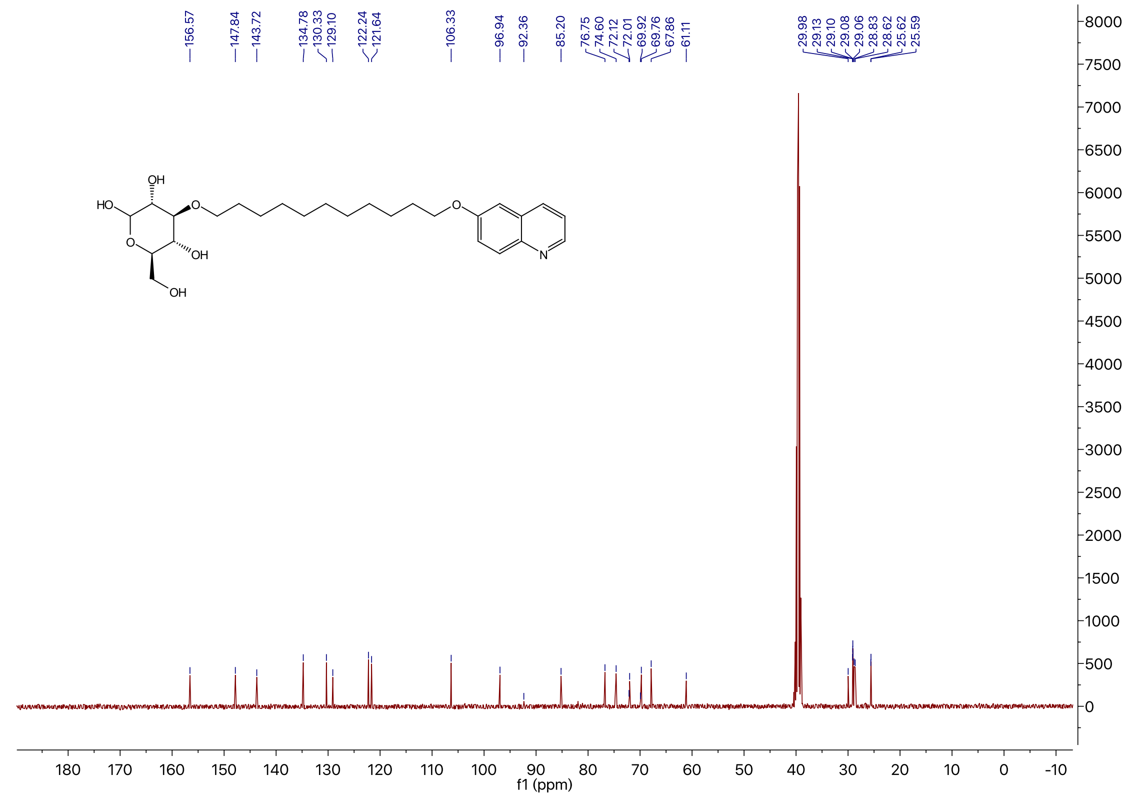

### $^{1}$H NMR of Compound **1i** in DMSO-$d_6$

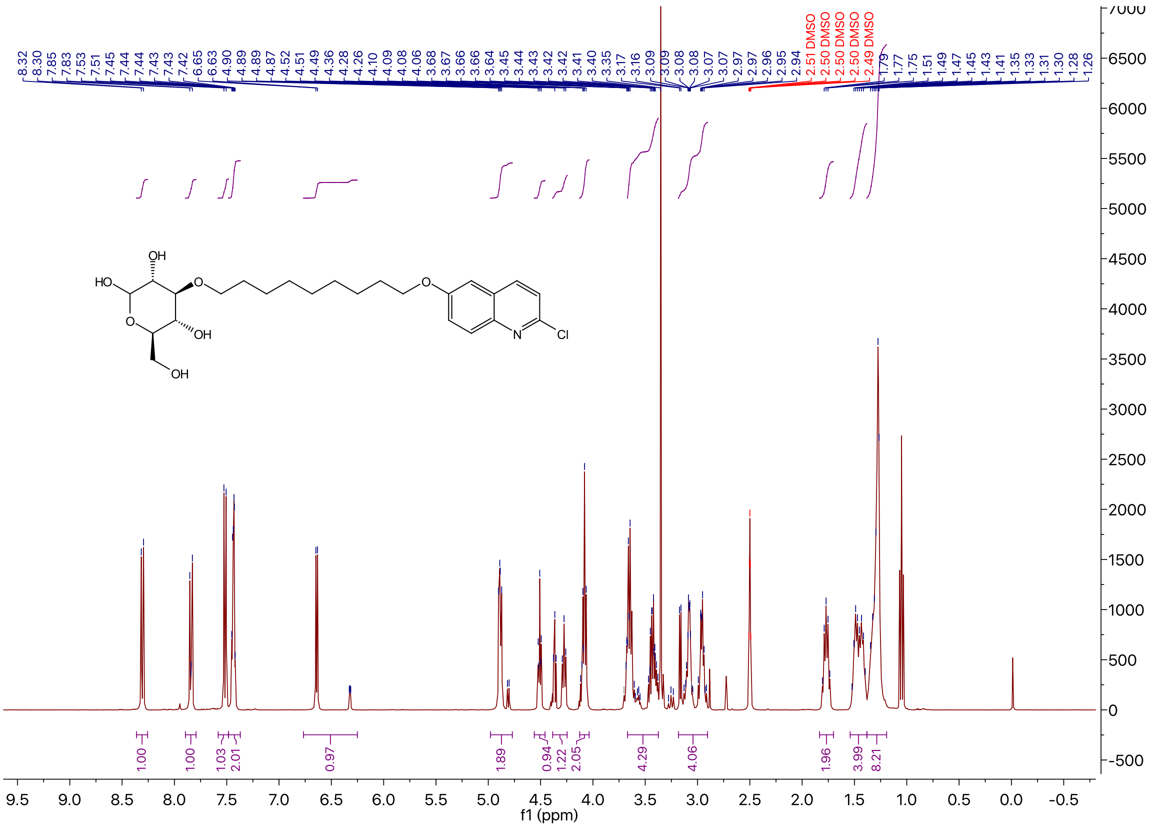

### $^{13}$C NMR of Compound **1i** in DMSO-$d_6$

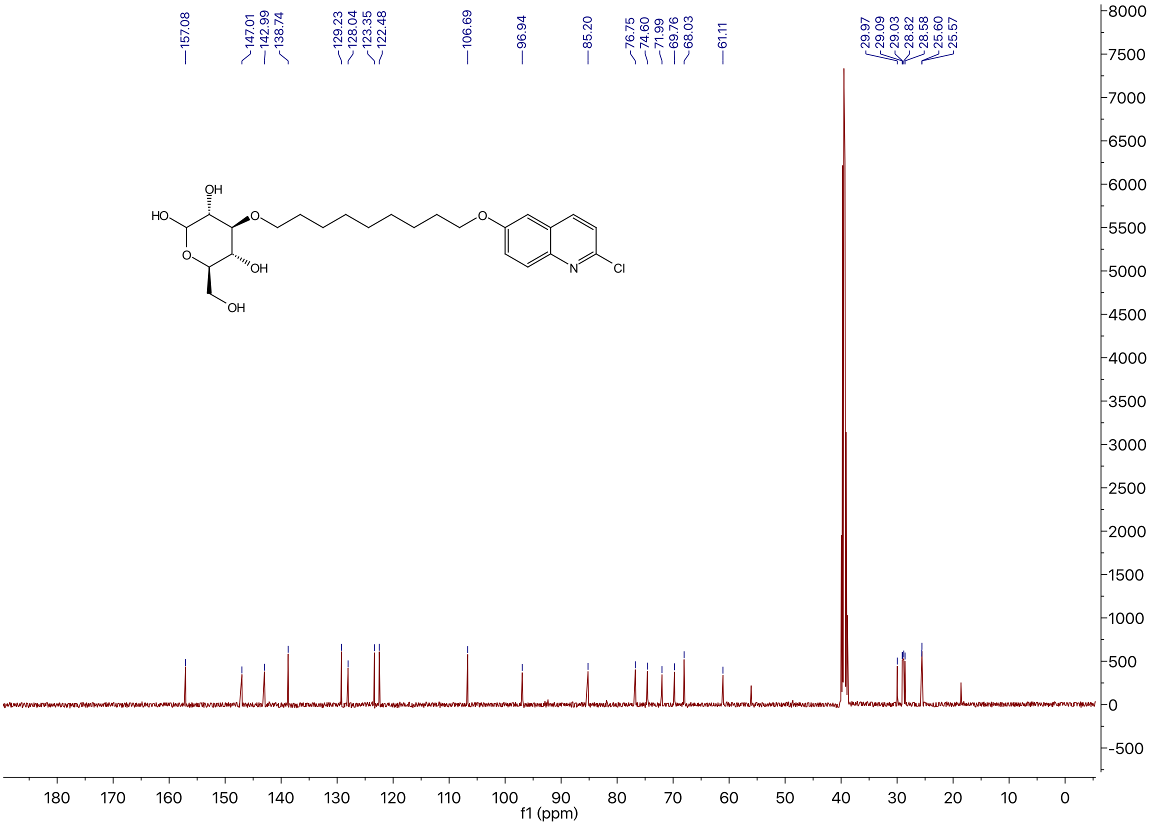

### $^{1}$H NMR of Compound **2a** in DMSO-$d_6$

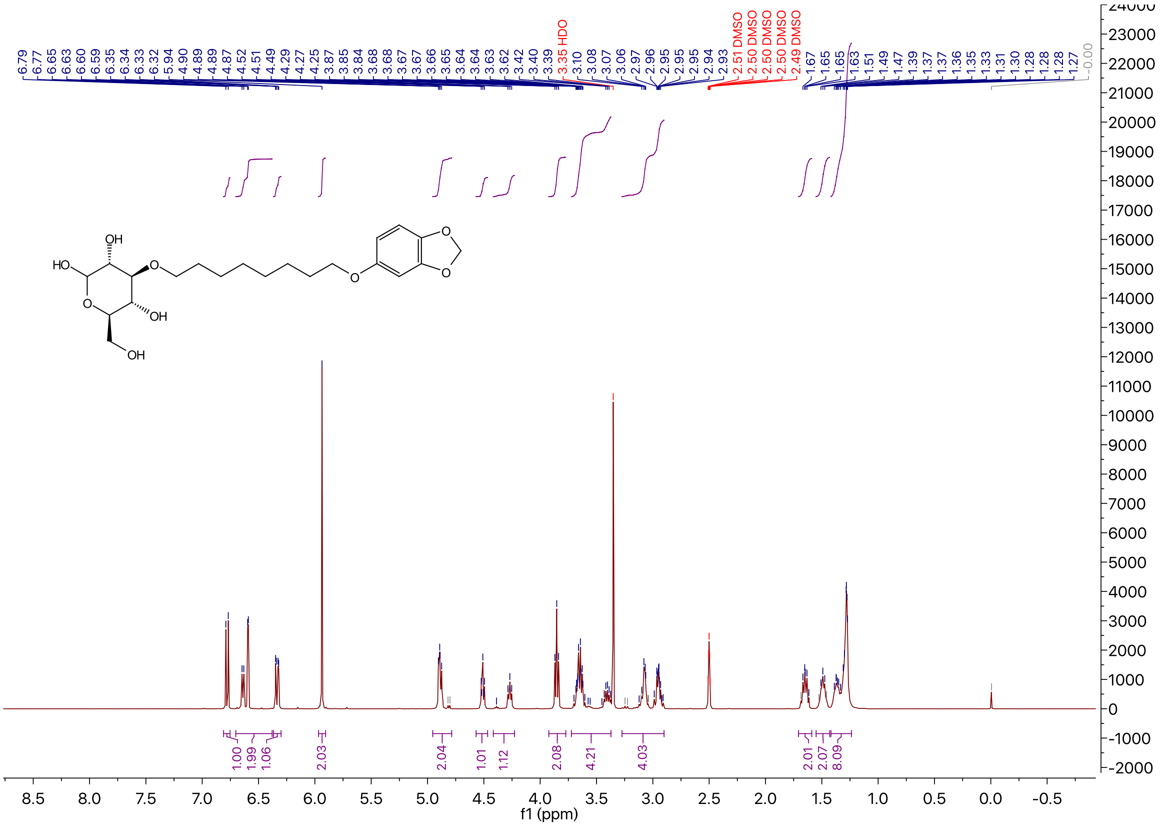

### $^{13}$C NMR of Compound **2a** in DMSO-$d_6$

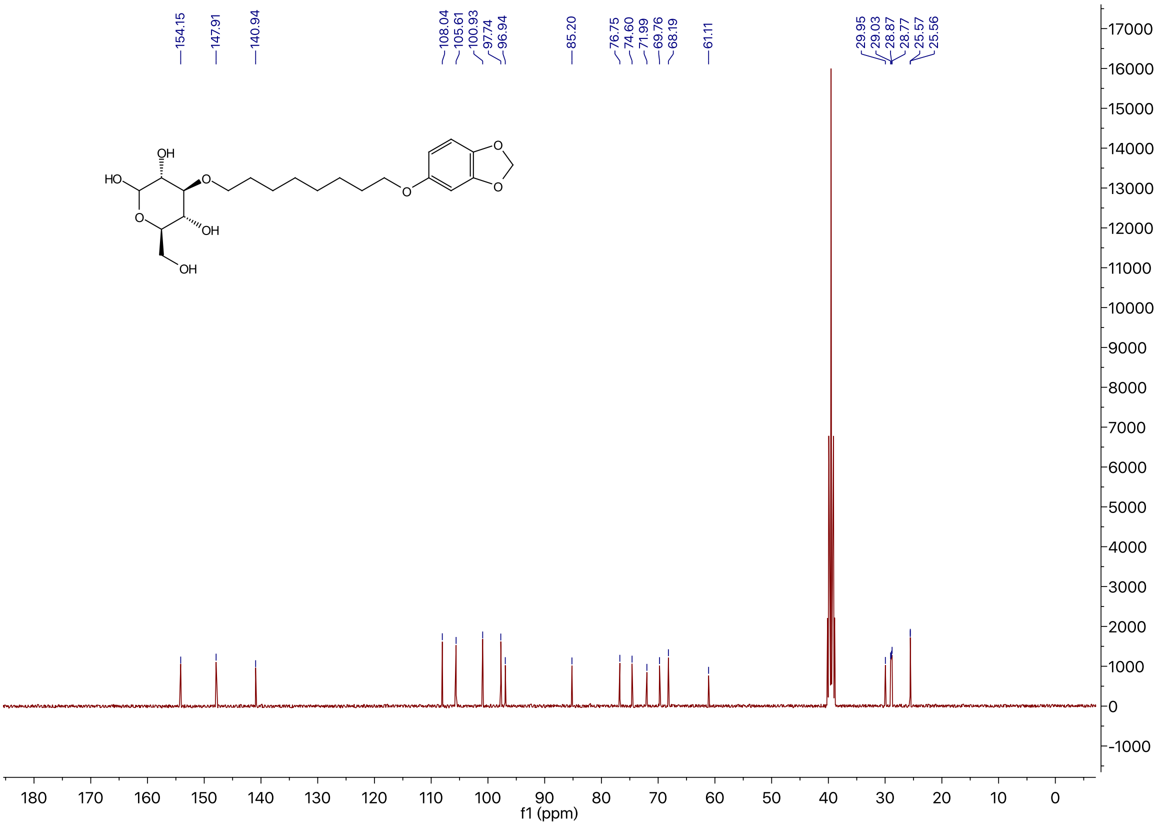

### $^1$H NMR of Compound **2b** in DMSO-$d_6$

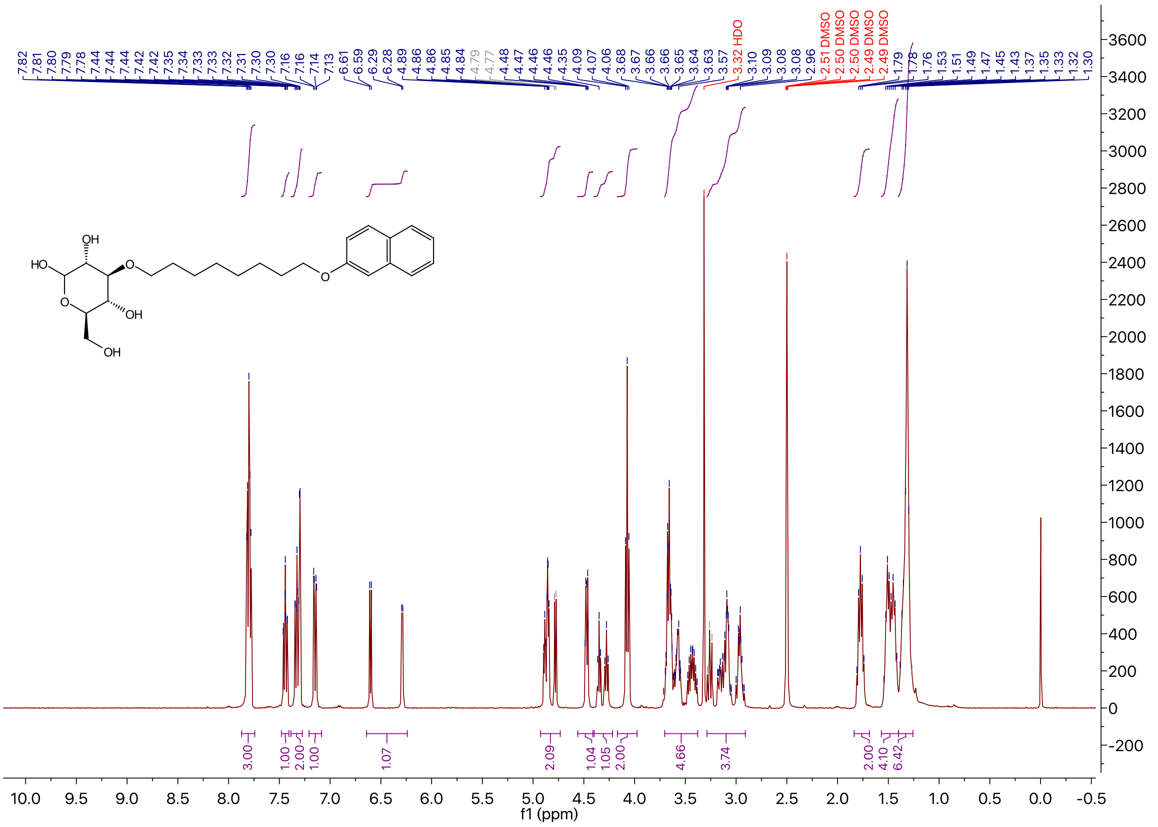

### $^{13}C$ NMR of Compound **2b** in DMSO-$d_6$

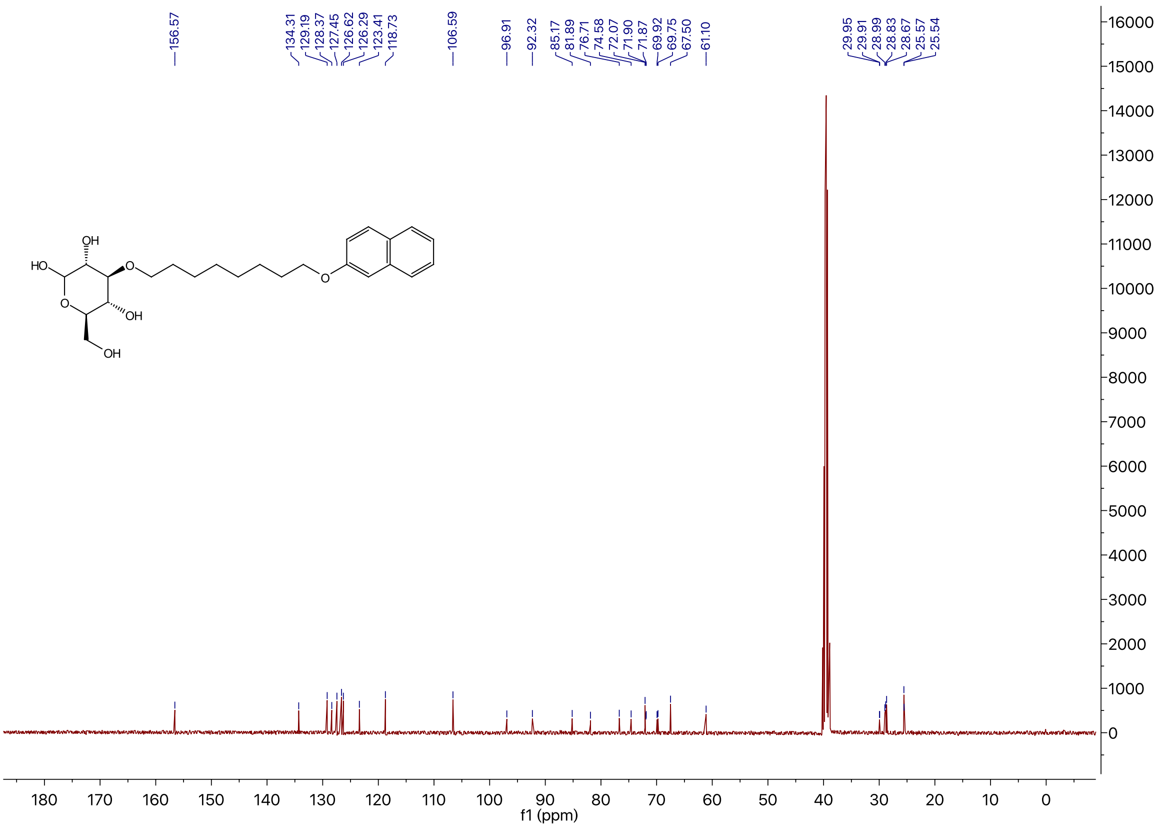

### $^{1}H$ NMR of Compound **2c** in DMSO-$d_6$

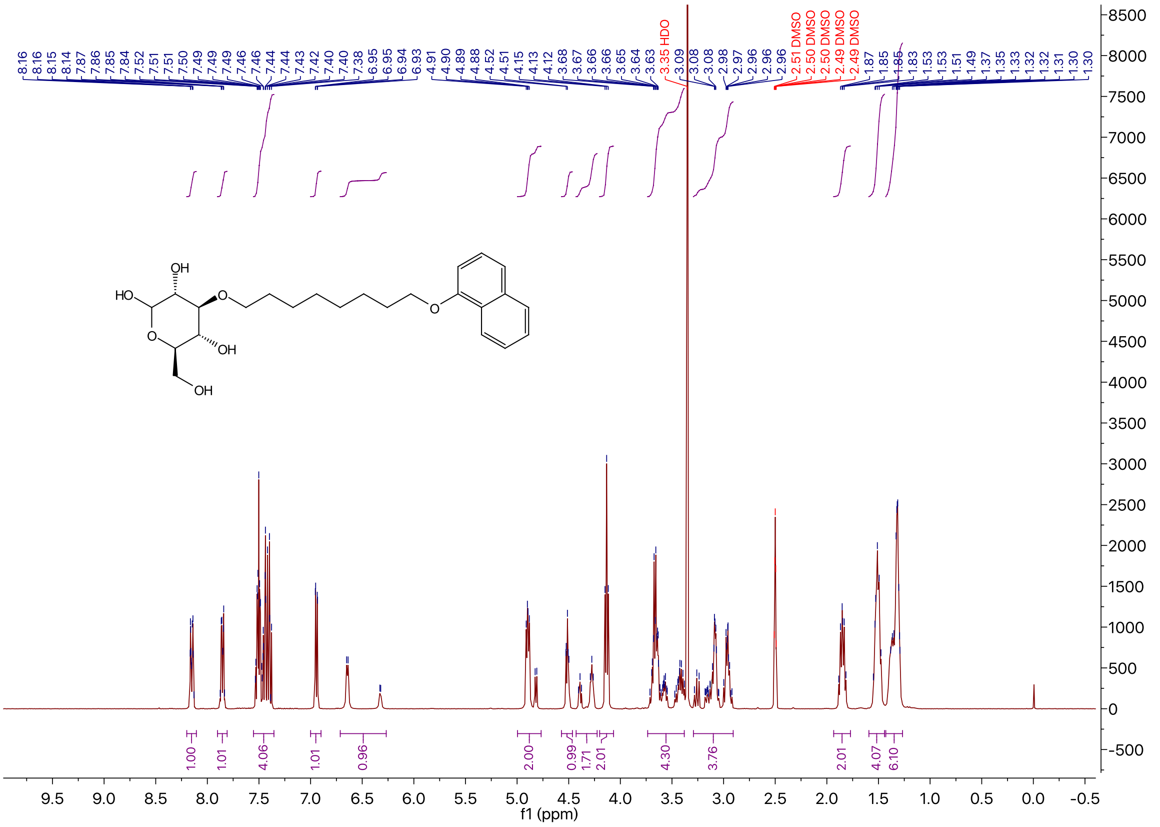

[13]C NMR of Compound **2c** in DMSO-$d_6$

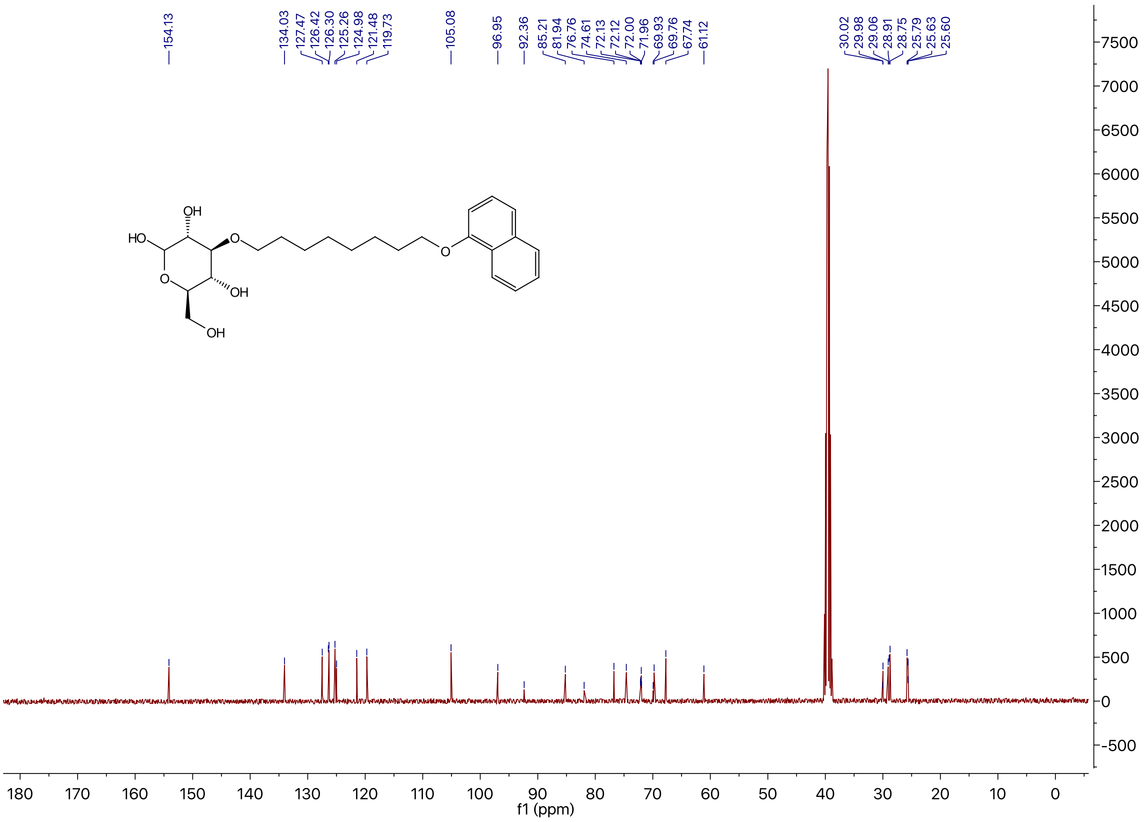

[1]H NMR of Compound **2d** in DMSO-$d_6$

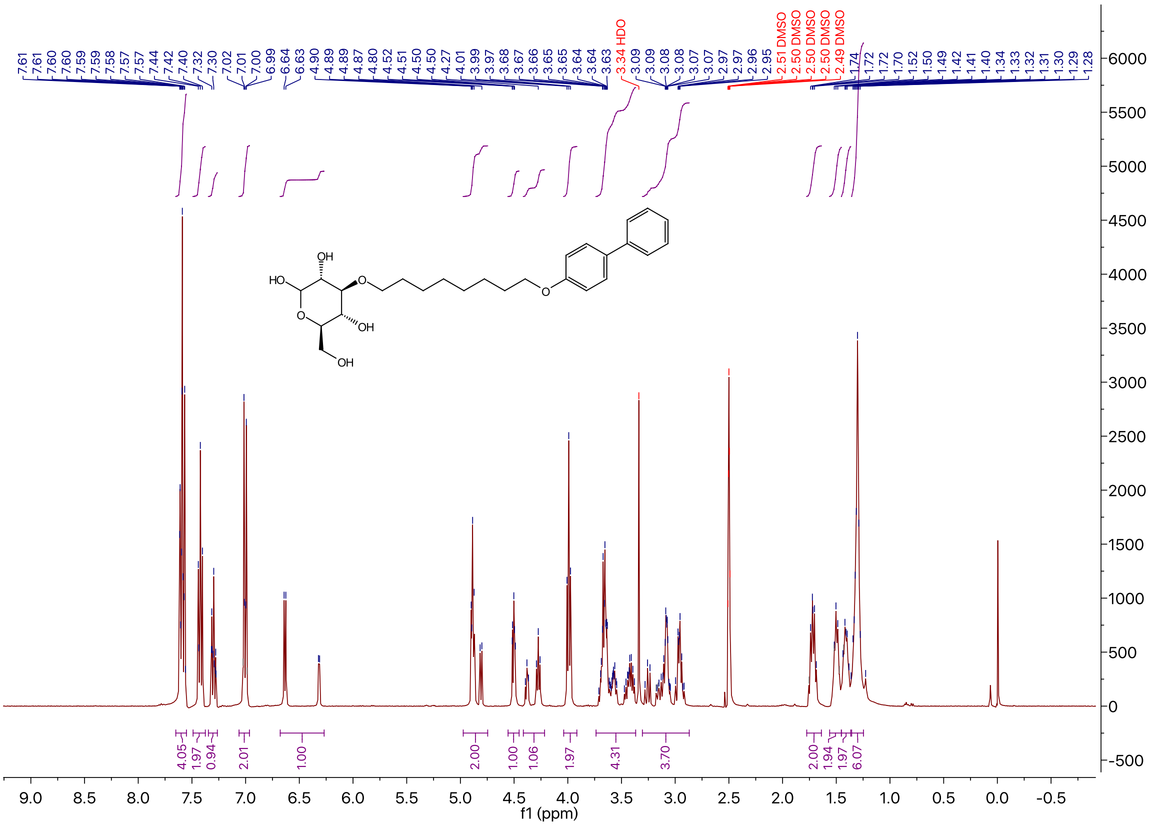

### $^{13}$C NMR of Compound **2d** in DMSO-$d_6$

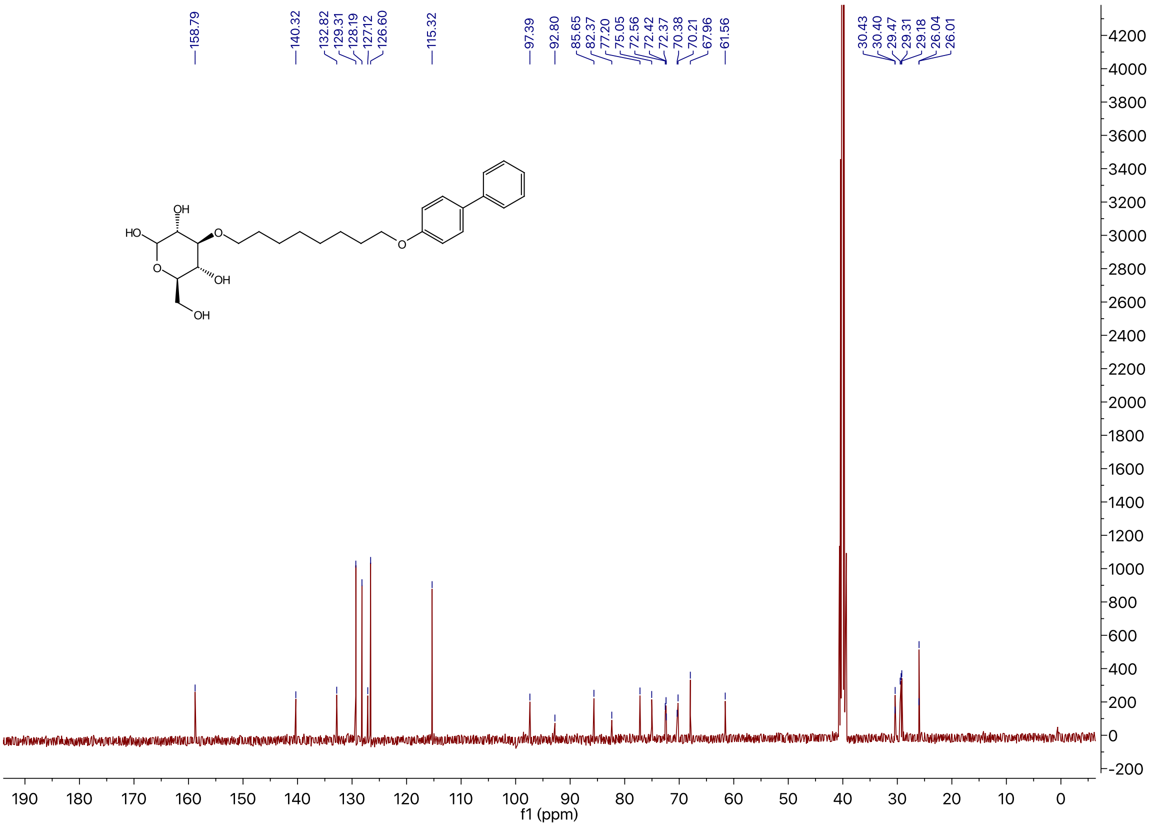

### $^{1}$H NMR of Compound **2e** in DMSO-$d_6$

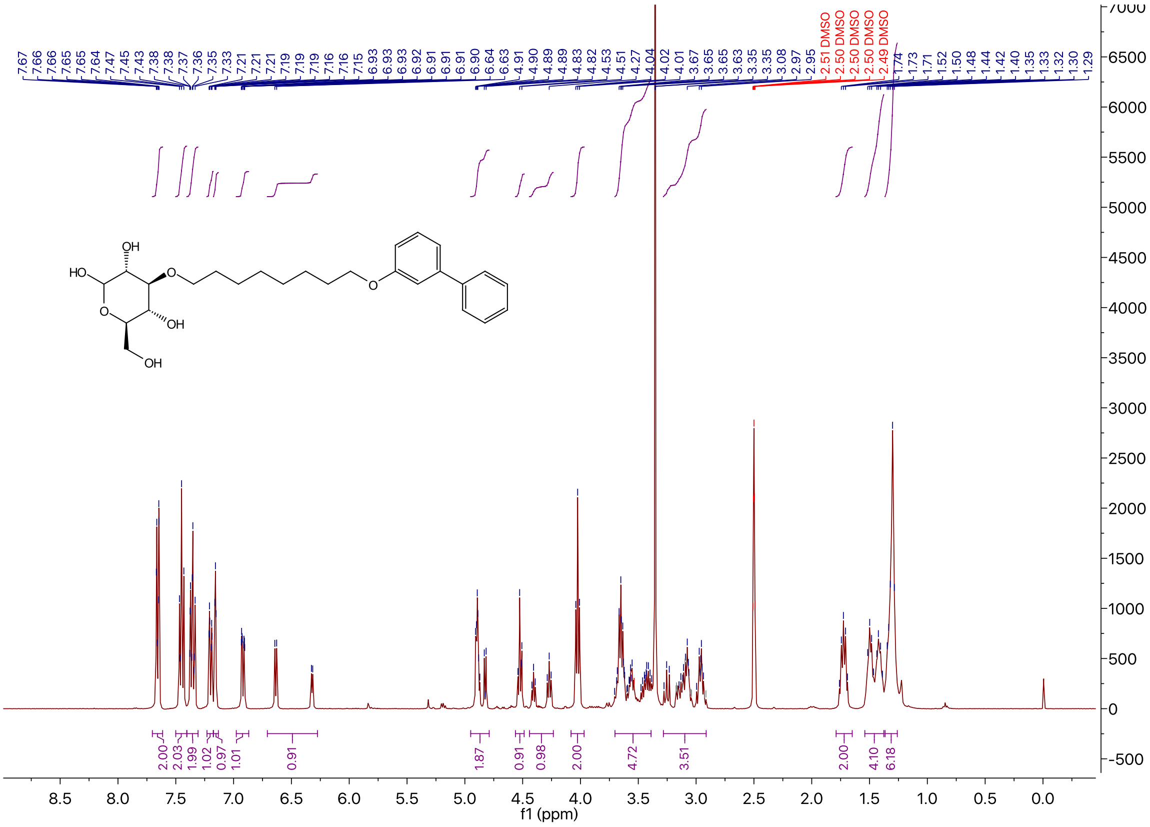

$^{13}$C NMR of Compound **2e** in DMSO-$d_6$

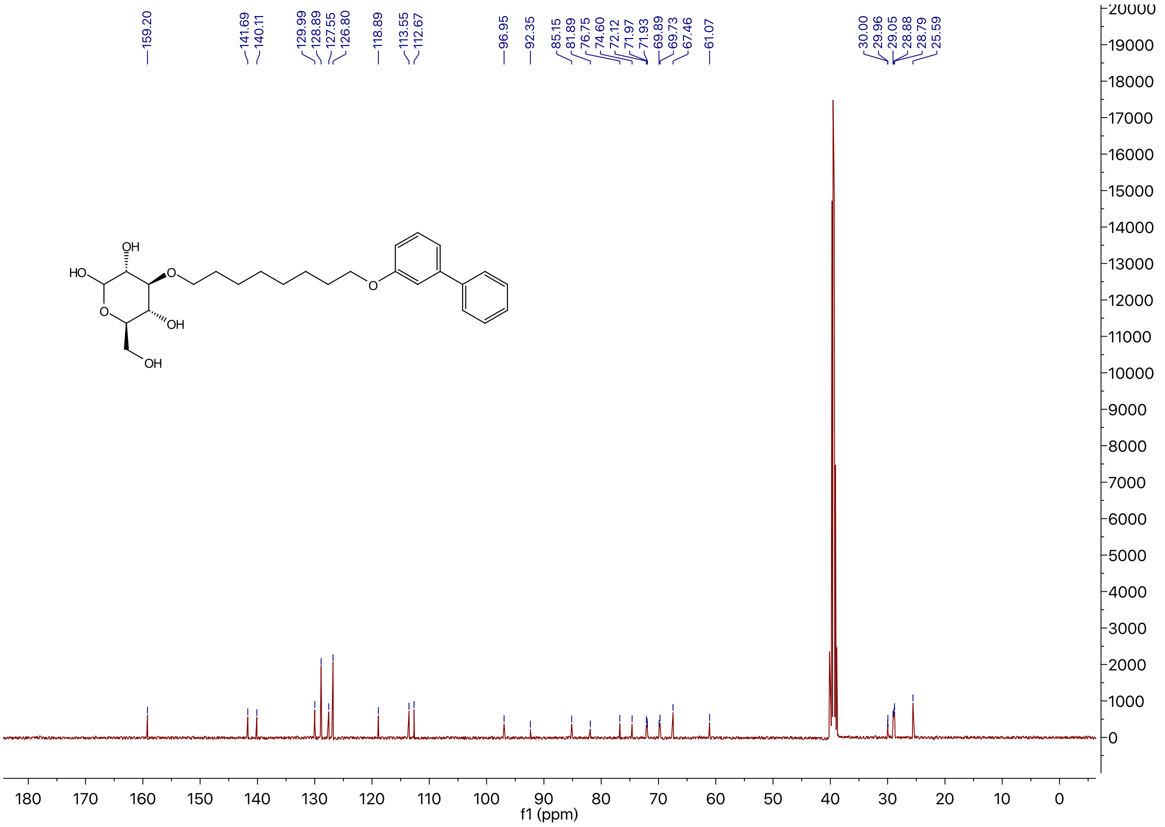

$^{1}$H NMR of Compound **11** in DMSO-$d_6$

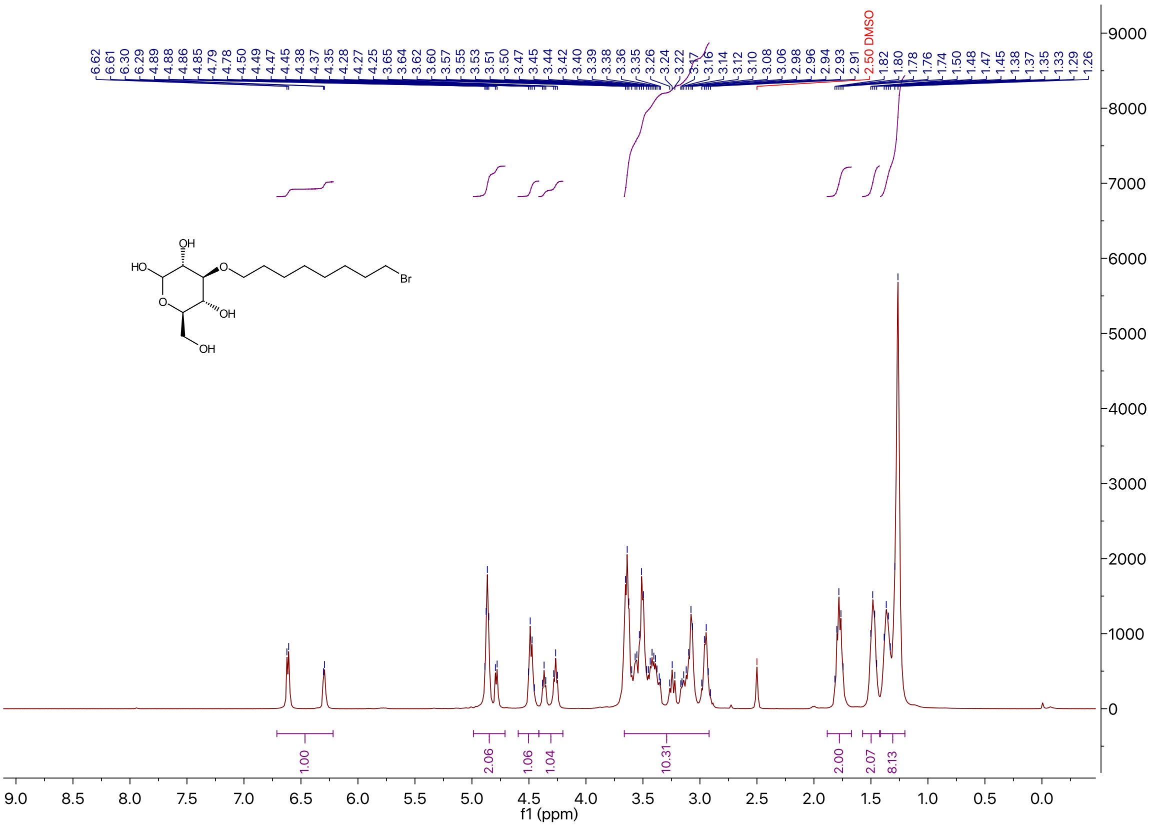

$^{13}C$ NMR of Compound **11** in DMSO-$d_6$

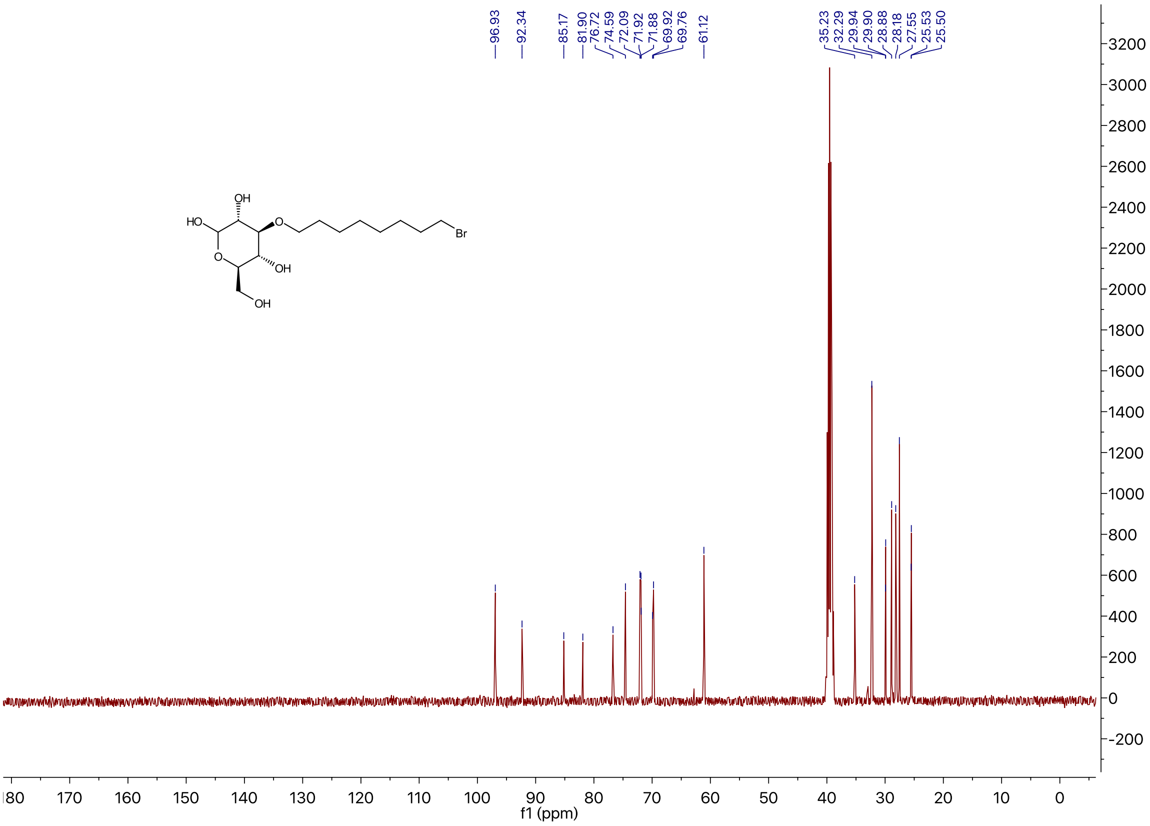

$^{1}H$ NMR of Compound **2f** in DMSO-$d_6$

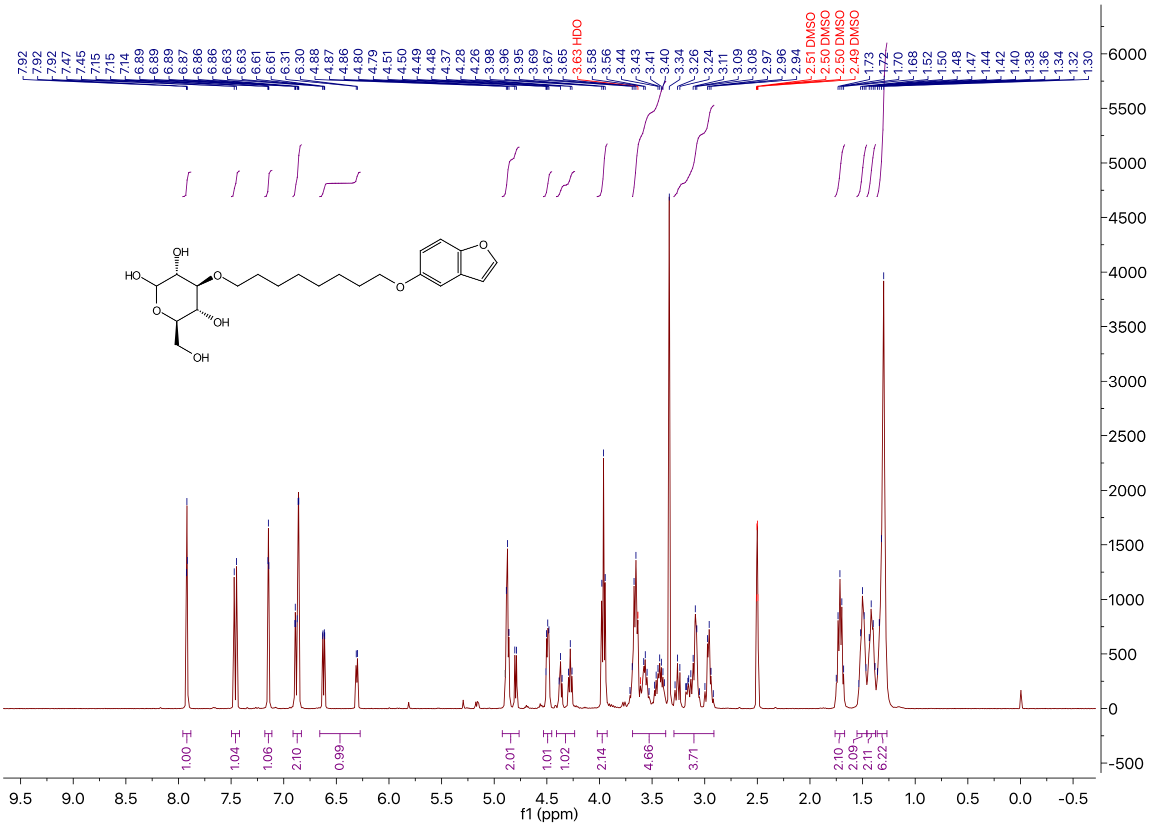

### $^{13}$C NMR of Compound **2f** in DMSO-$d_6$

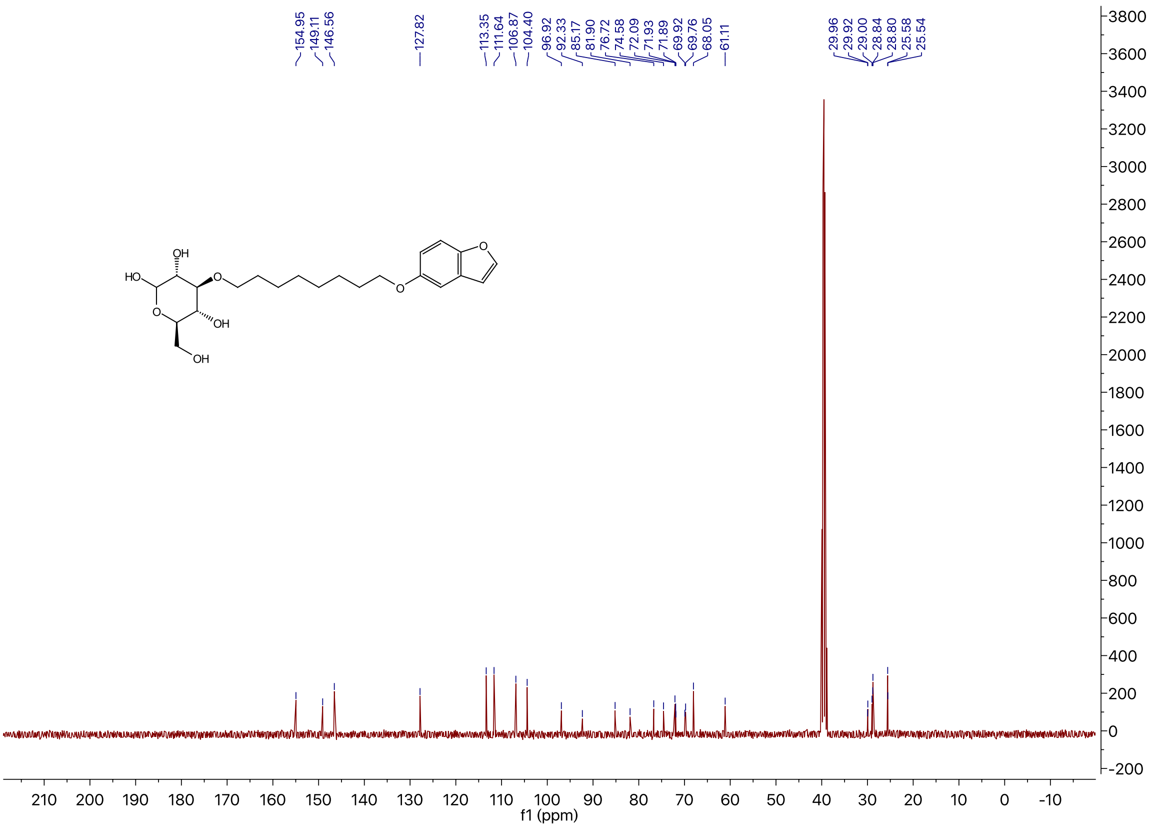

### $^{1}$H NMR of Compound **2g** in DMSO-$d_6$

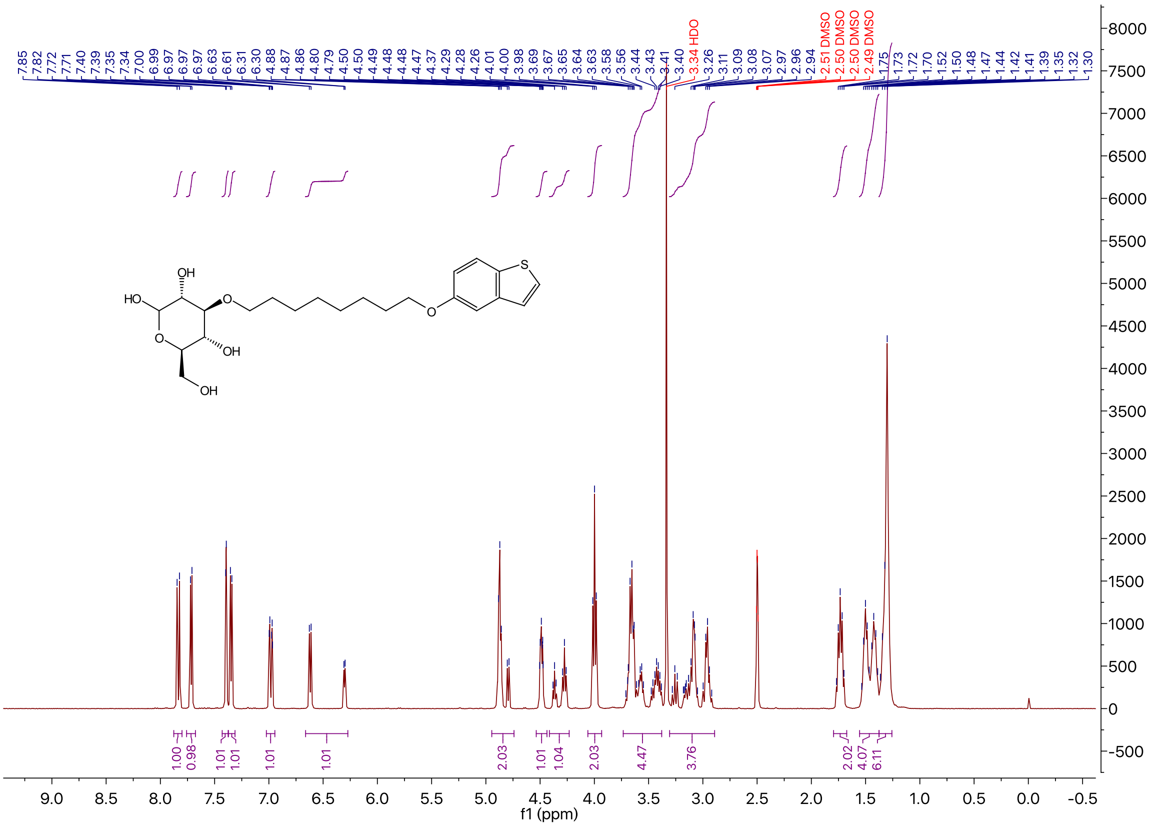

### $^{13}$C NMR of Compound **2g** in DMSO-$d_6$

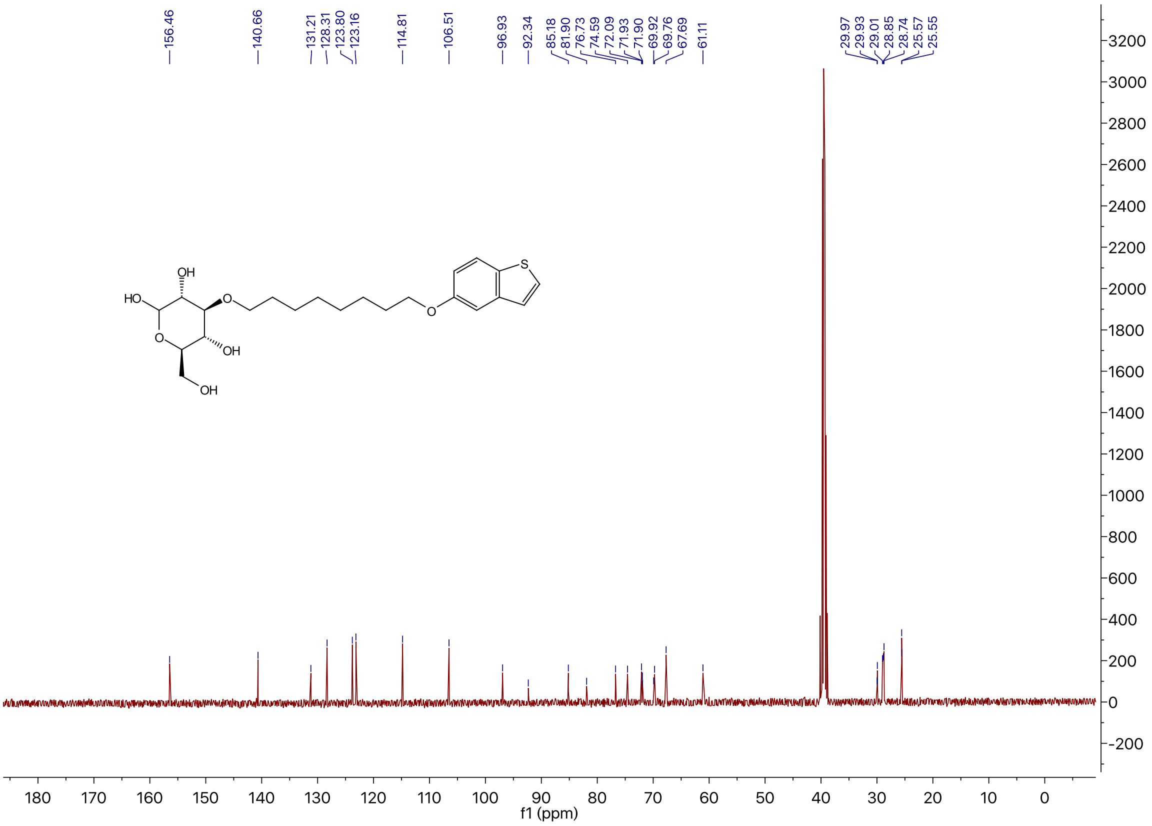

### $^{1}$H NMR of Compound **2h** in DMSO-$d_6$

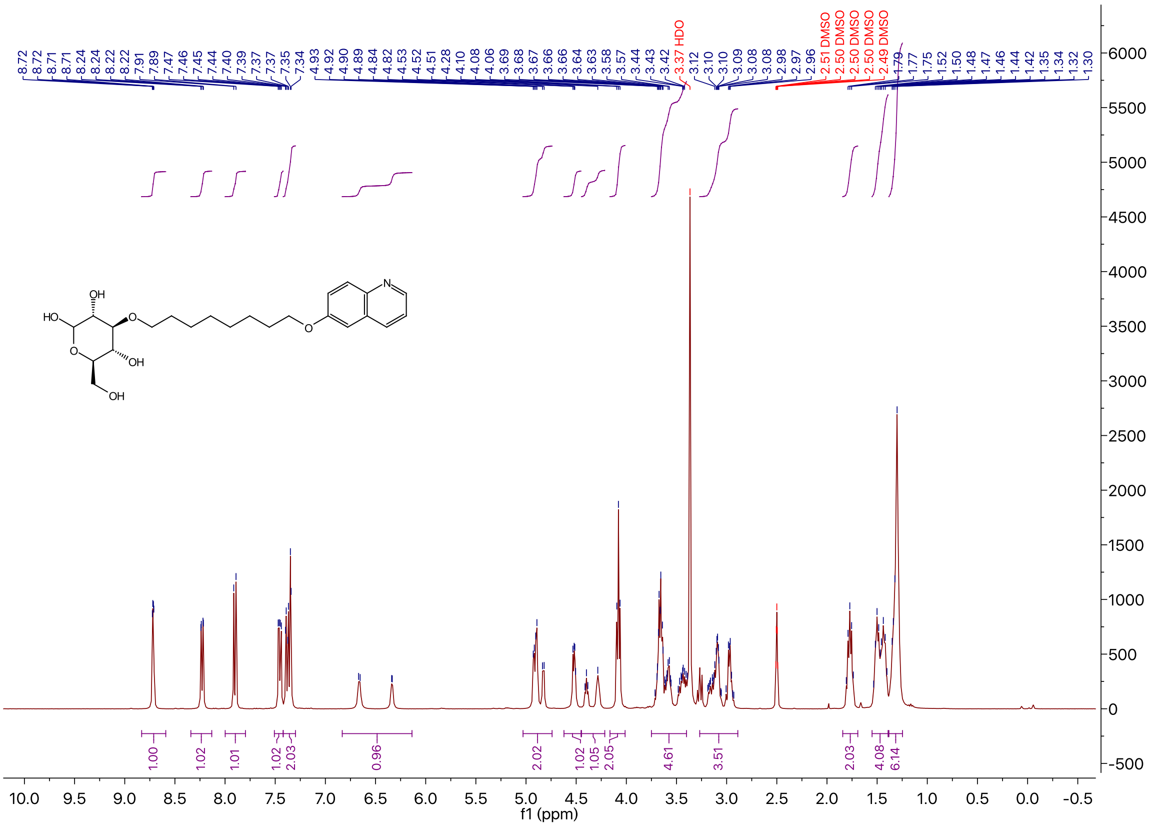

### $^{13}C$ NMR of Compound **2h** in DMSO-$d_6$

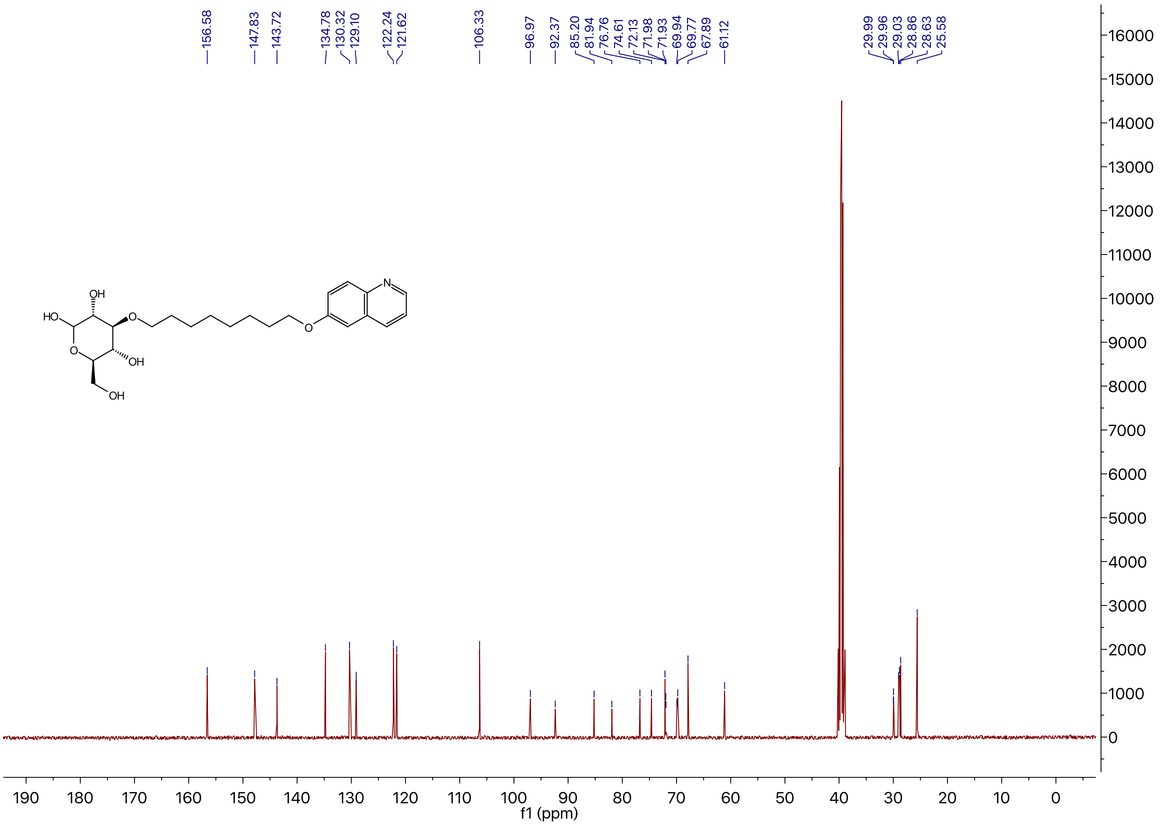

### $^{1}H$ NMR of Compound **2i** in DMSO-$d_6$

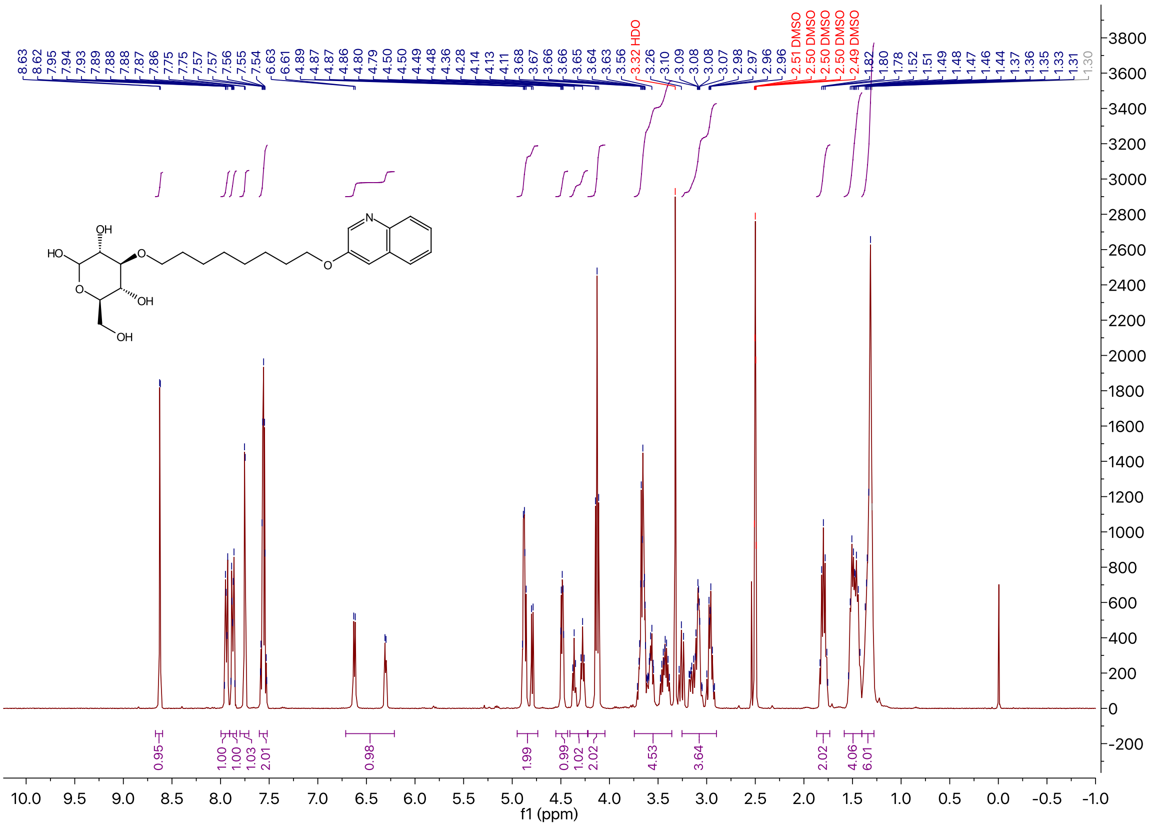

### [13]C NMR of Compound **2i** in DMSO-$d_6$

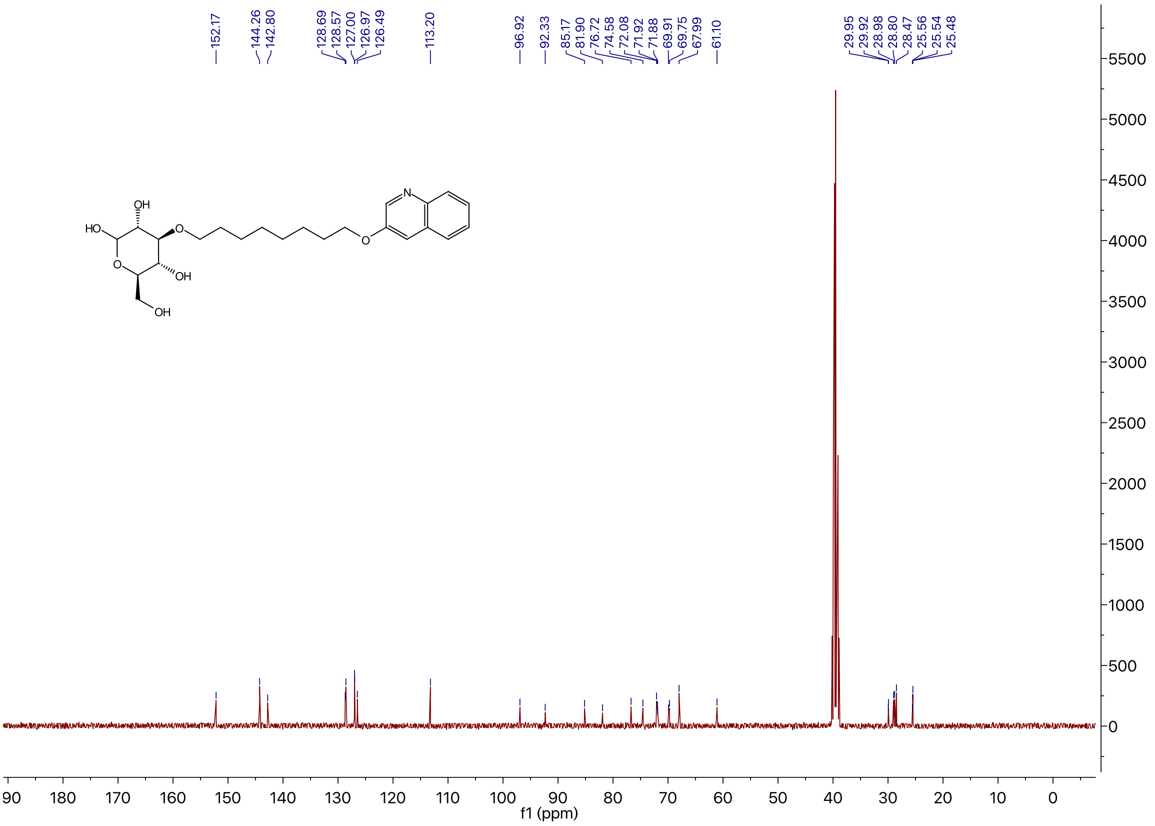

### [1]H NMR of Compound **2j** in DMSO-$d_6$

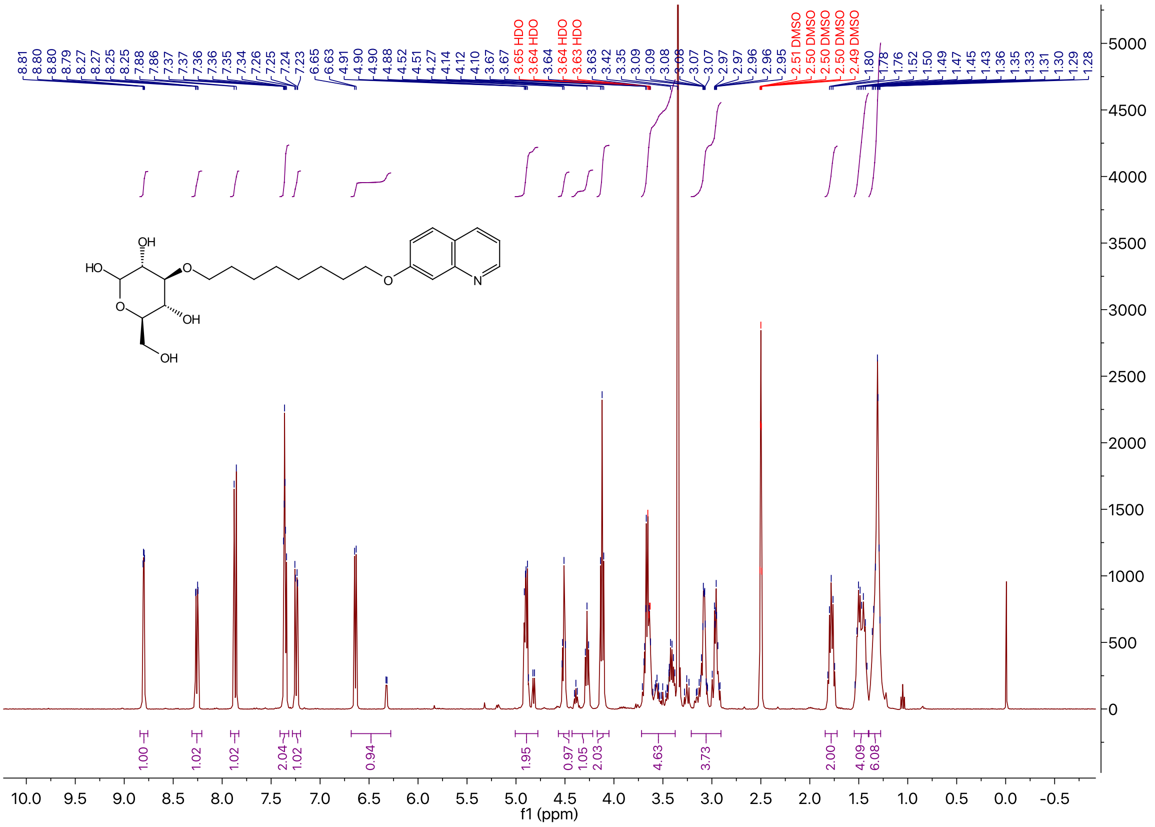

$^{13}C$ NMR of Compound **2j** in DMSO-$d_6$

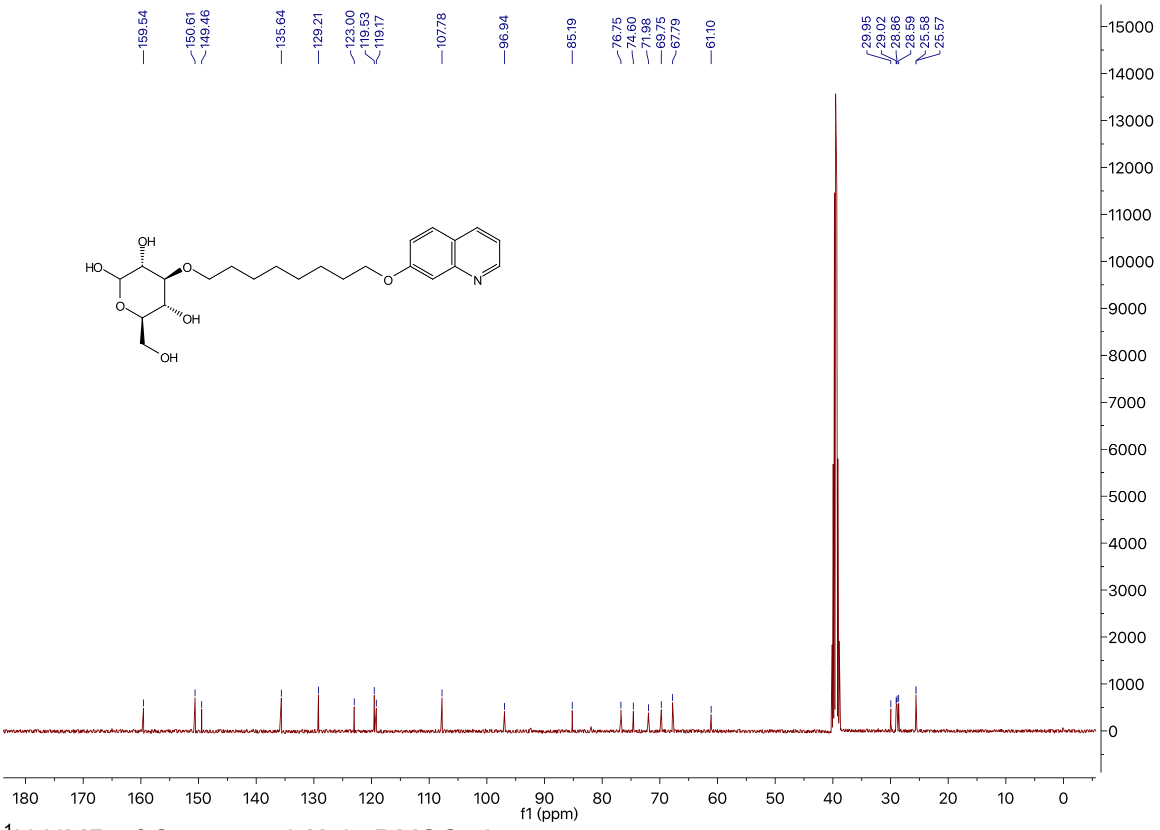

$^{1}H$ NMR of Compound **2k** in DMSO-$d_6$

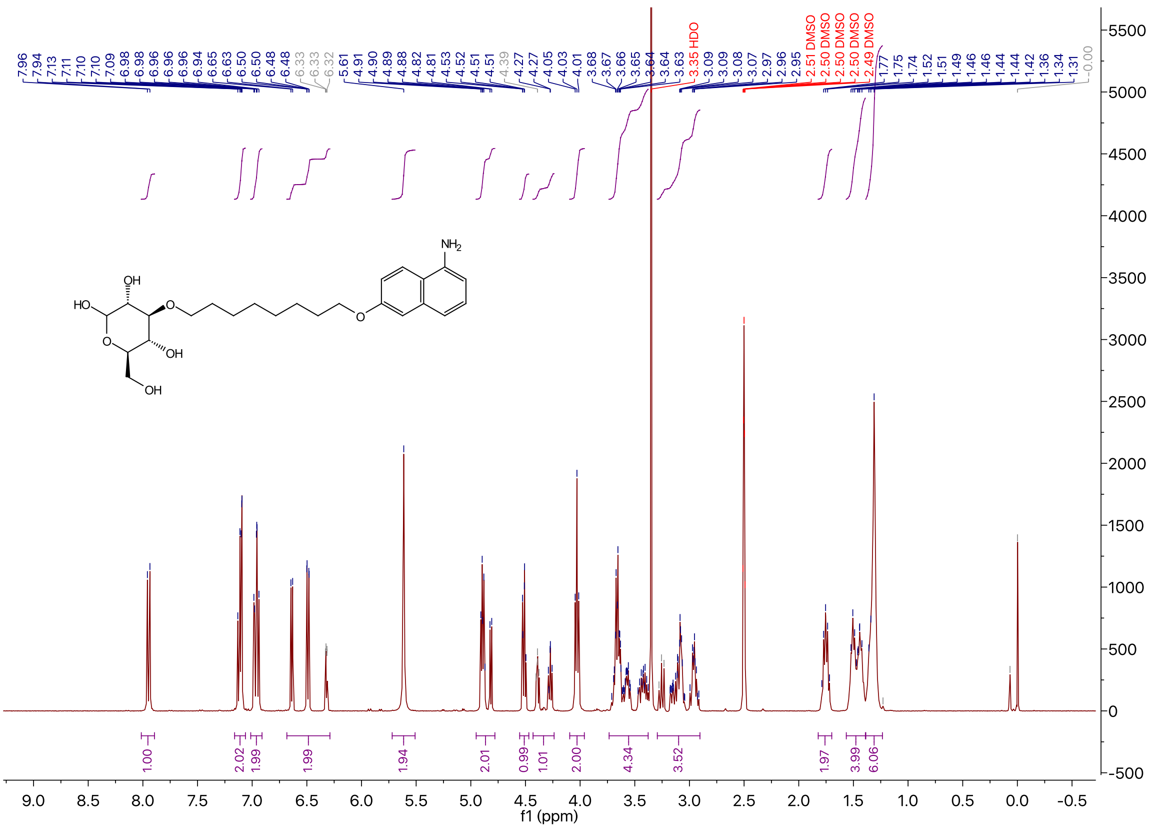

$^{13}C$ NMR of Compound **2k** in DMSO-$d_6$

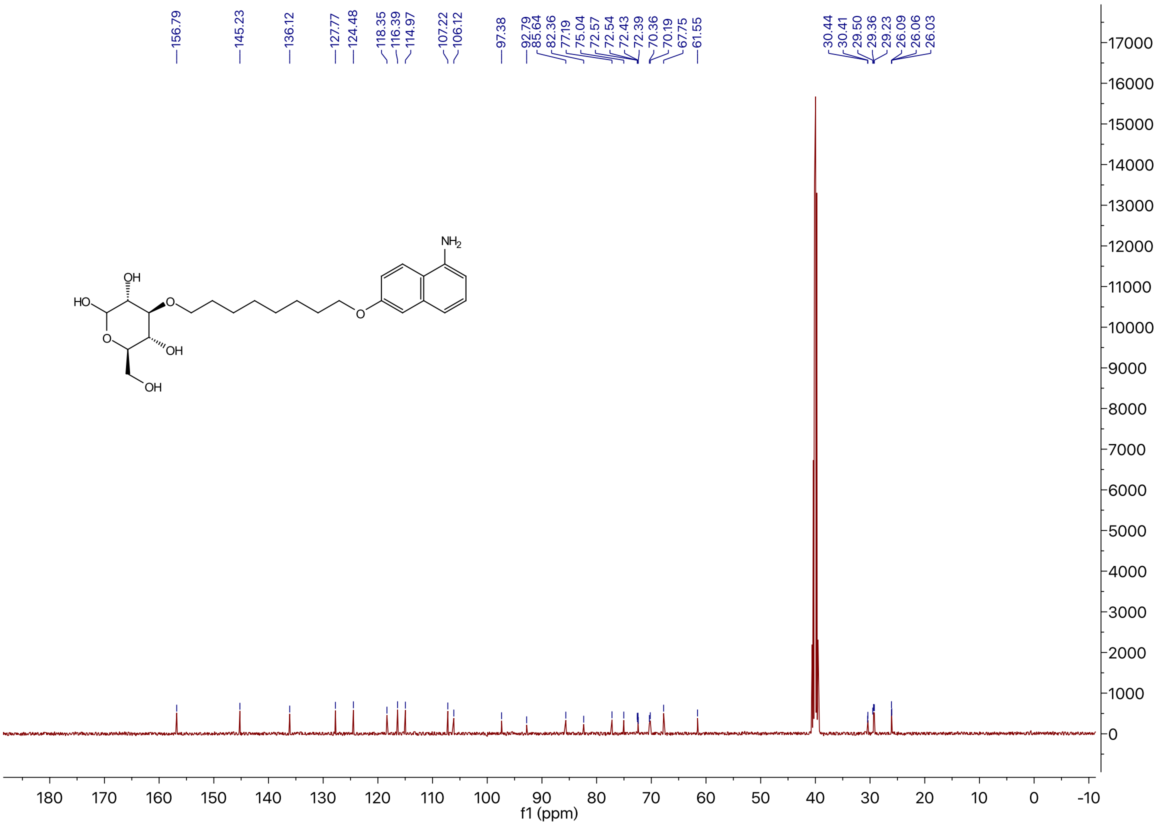

$^{1}H$ NMR of Compound **2l** in DMSO-$d_6$

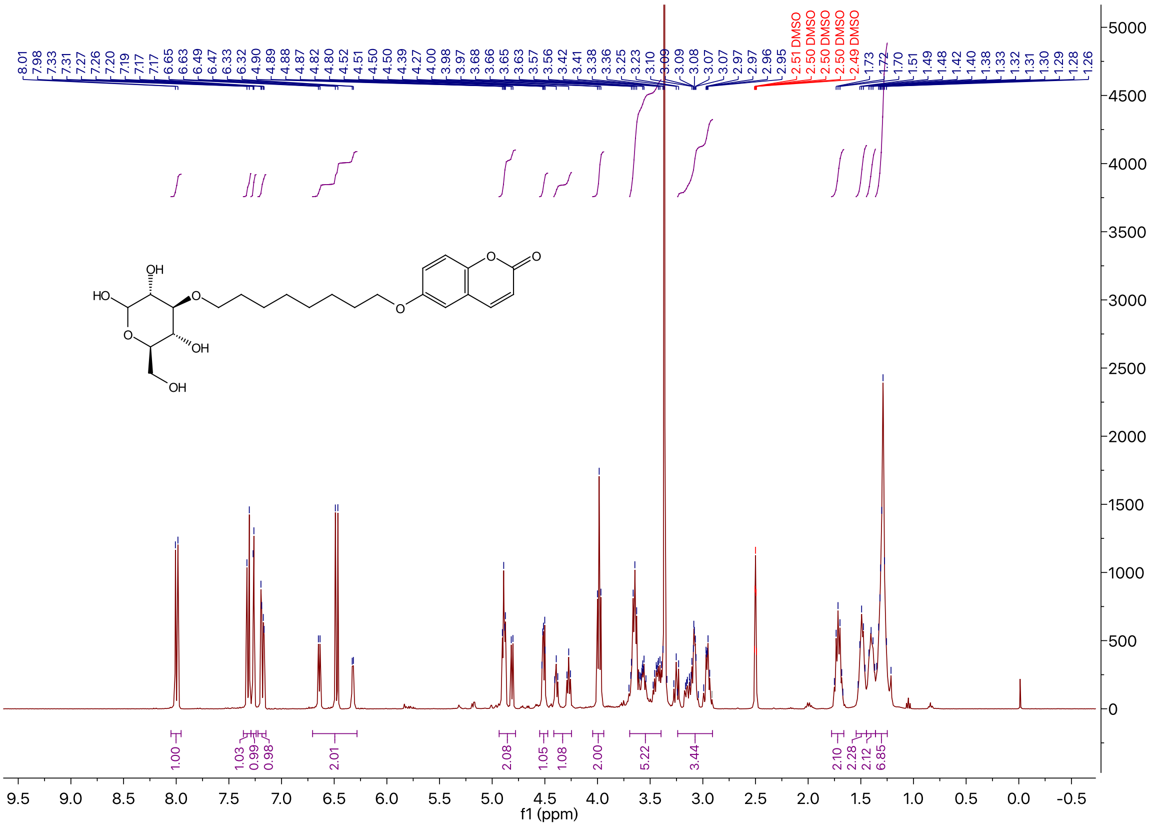

$^{13}C$ NMR of Compound **2l** in DMSO-$d_6$

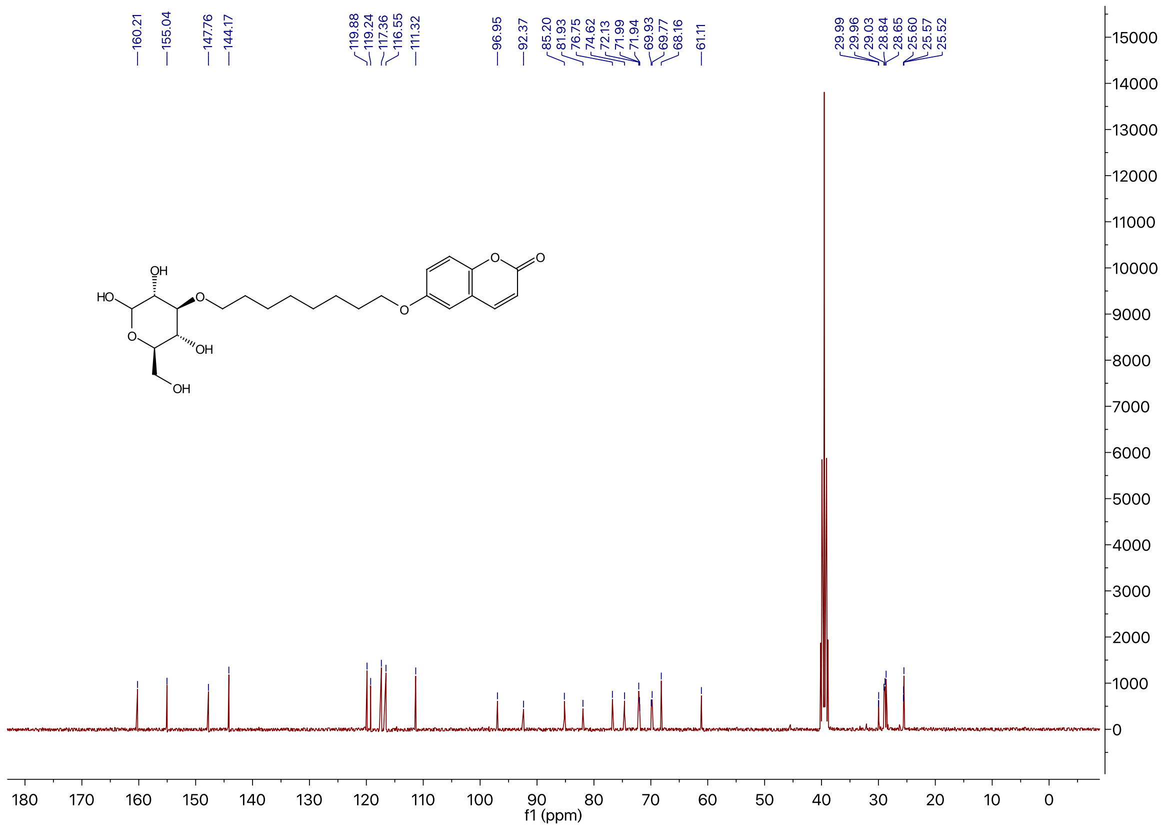

$^{1}H$ NMR of Compound **2m** in DMSO-$d_6$

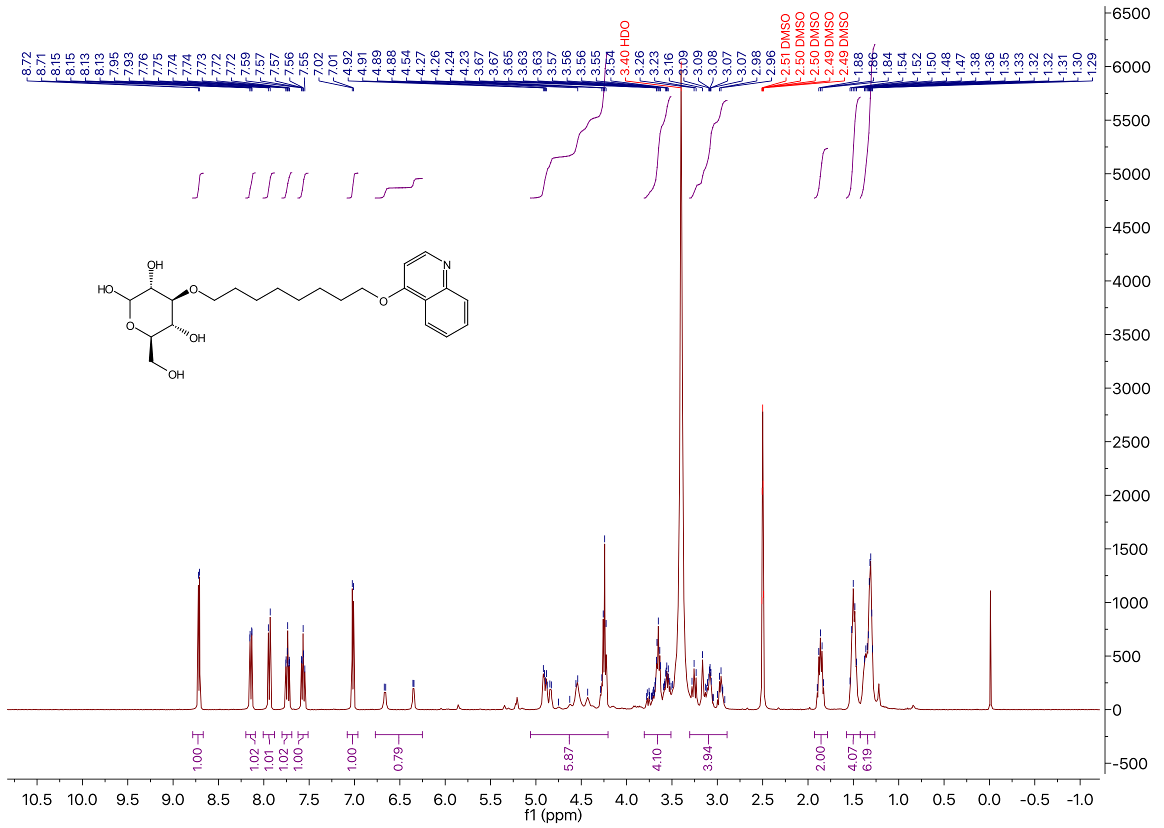

### $^{13}$C NMR of Compound **2m** in DMSO-$d_6$

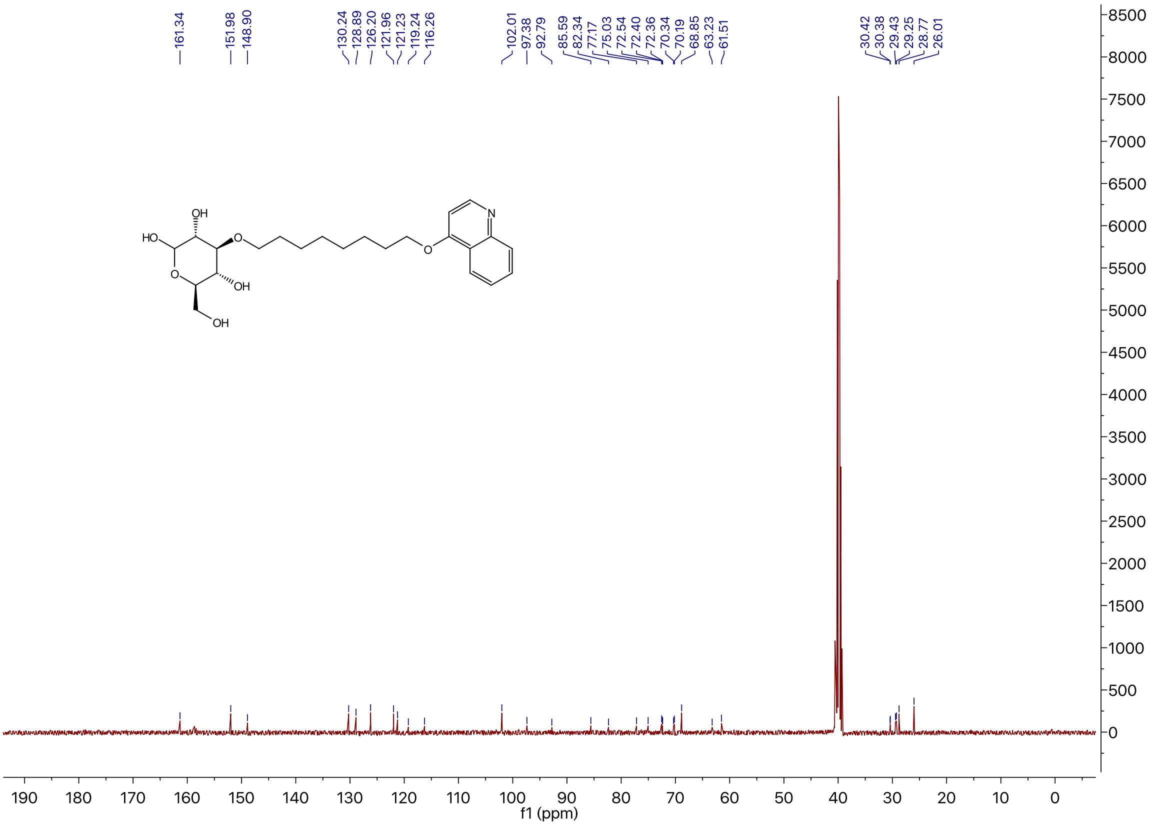

### $^{1}$H NMR of Compound **2n** in DMSO-$d_6$

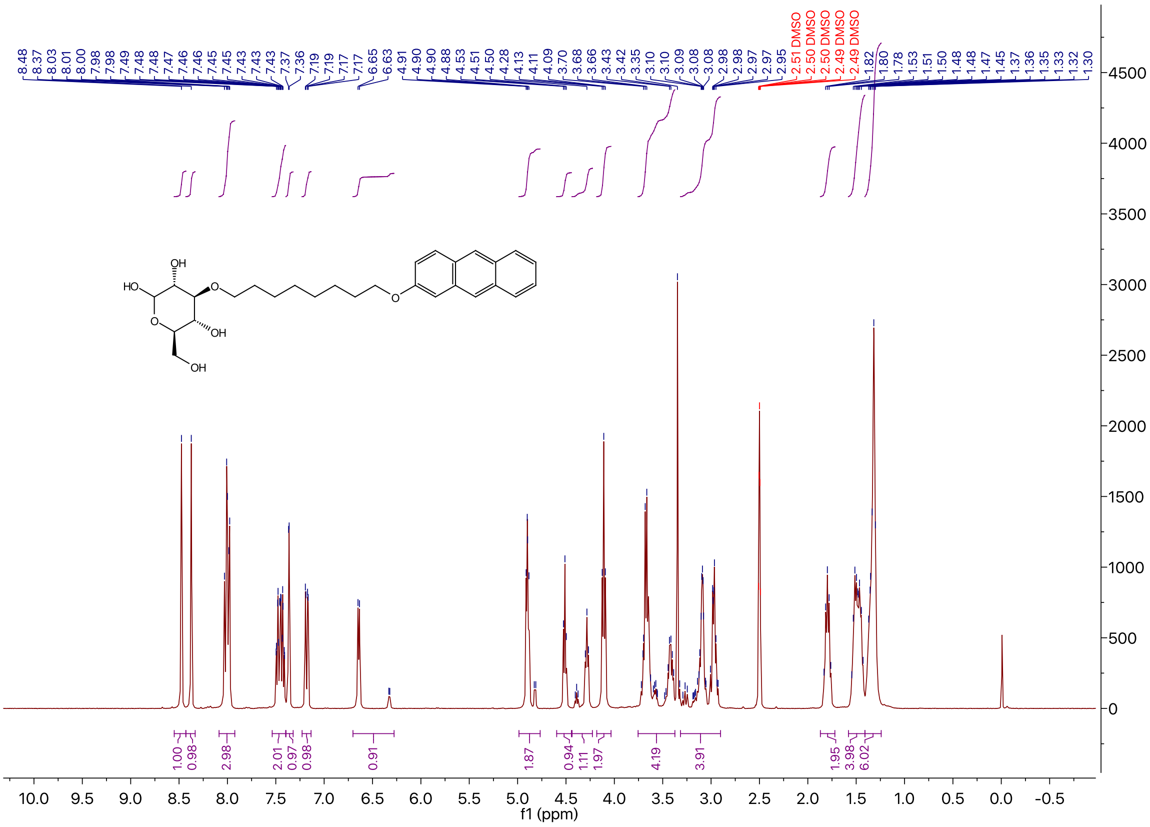

$^{13}$C NMR of Compound **2n** in DMSO-$d_6$

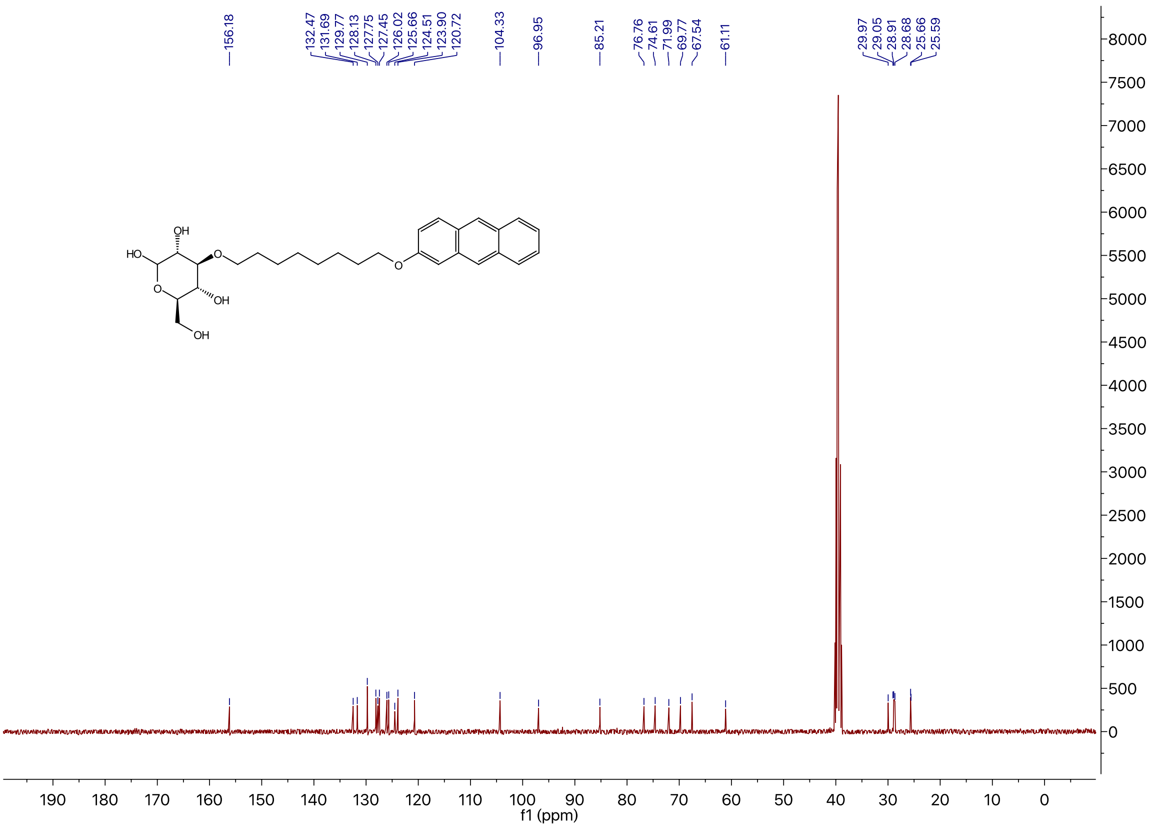

$^{1}$H NMR of Compound **2o** in DMSO-$d_6$

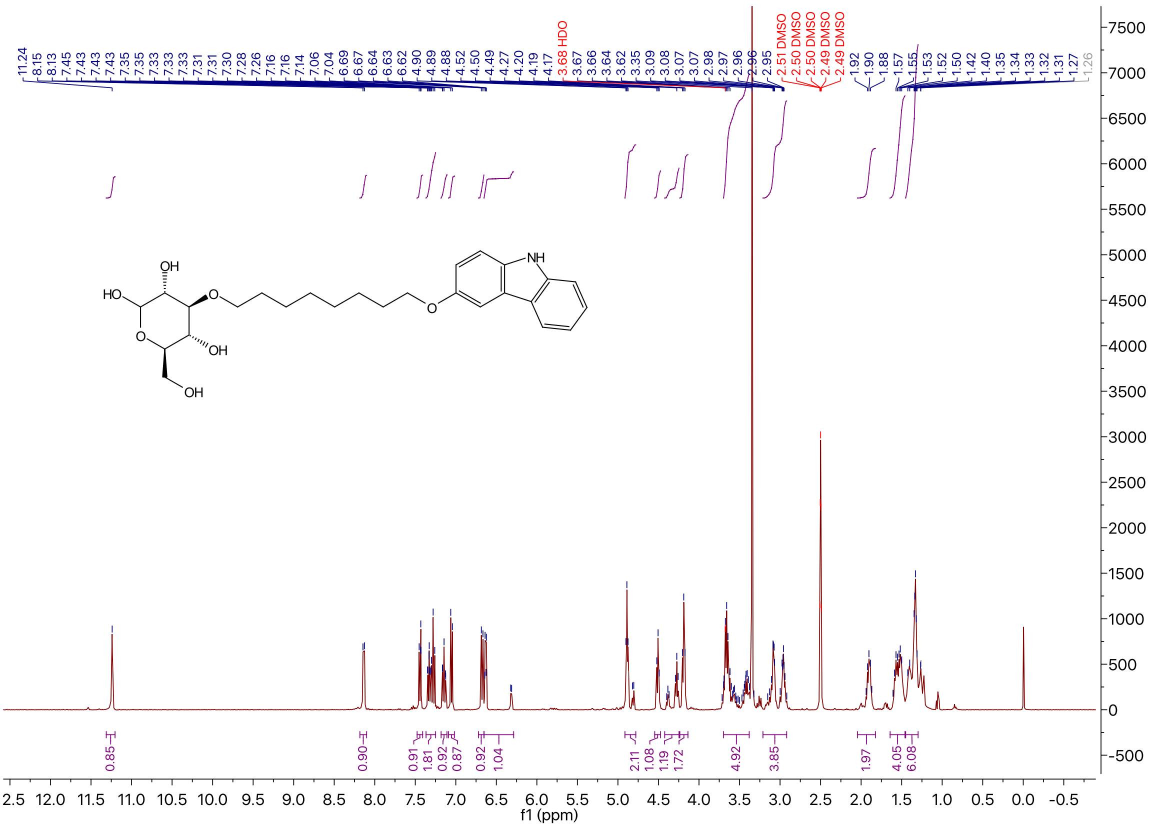

$^{13}$C NMR of Compound **2o** in DMSO-$d_6$

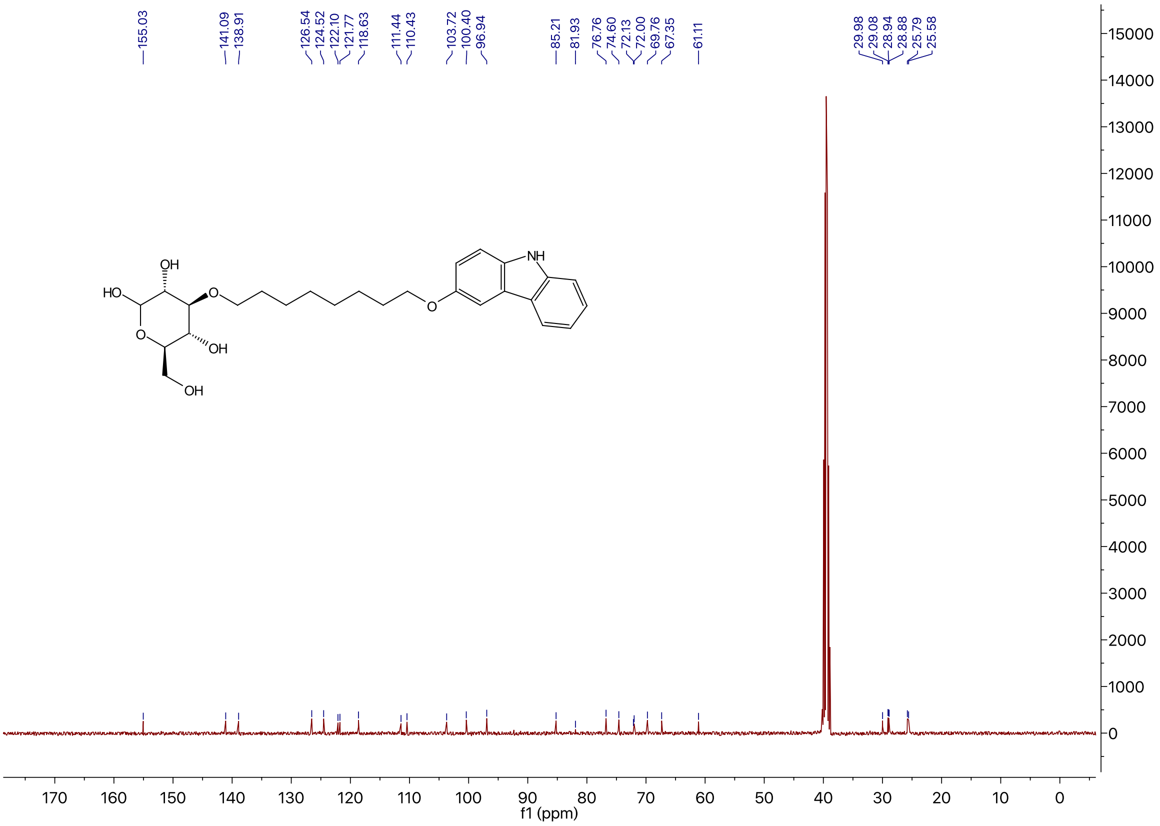

$^{1}$H NMR of Compound **2p** in DMSO-$d_6$

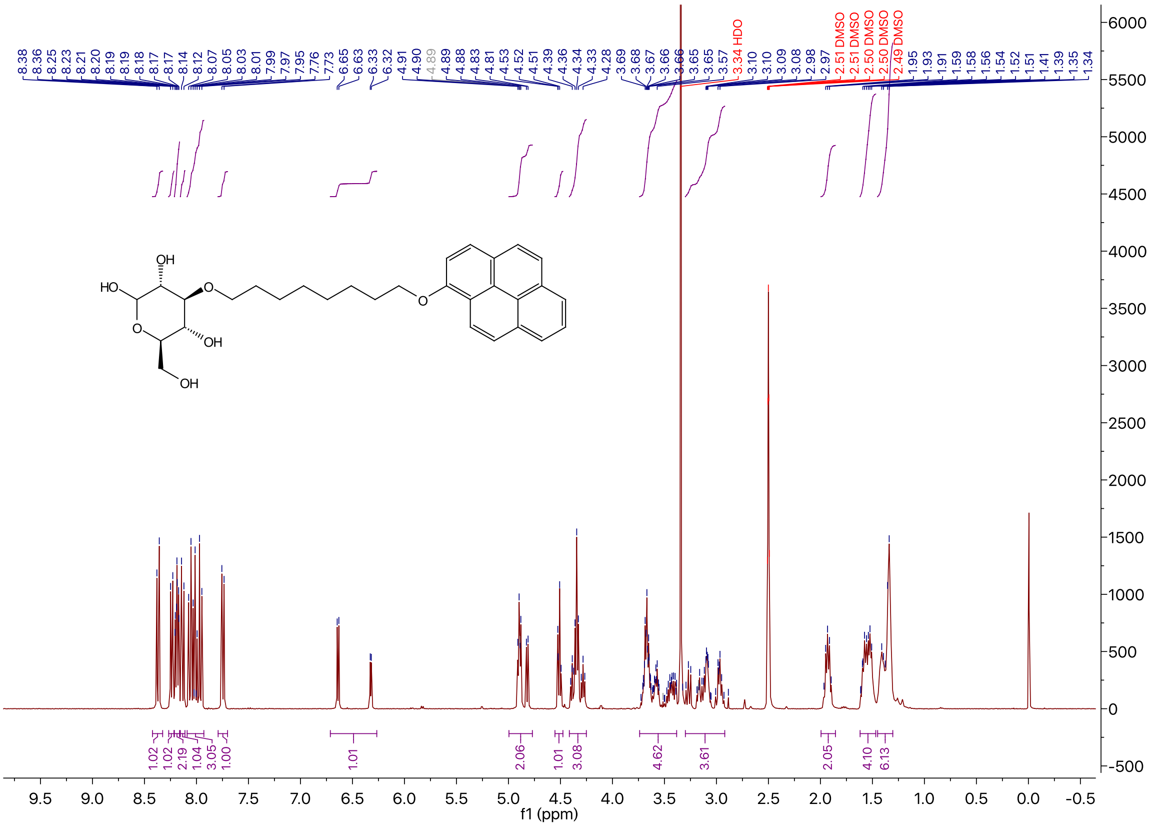

$^{13}$C NMR of Compound **2p** in DMSO-$d_6$

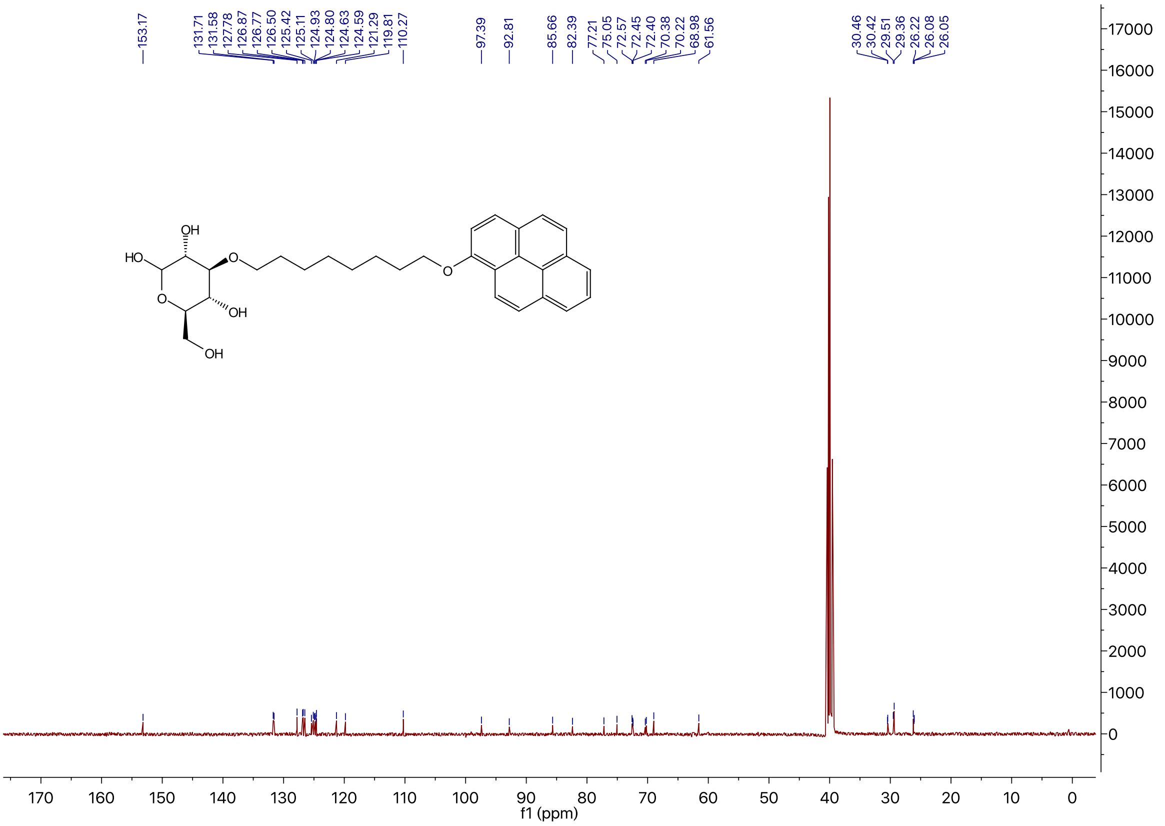

$^{1}$H NMR of Compound **3b** in DMSO-$d_6$

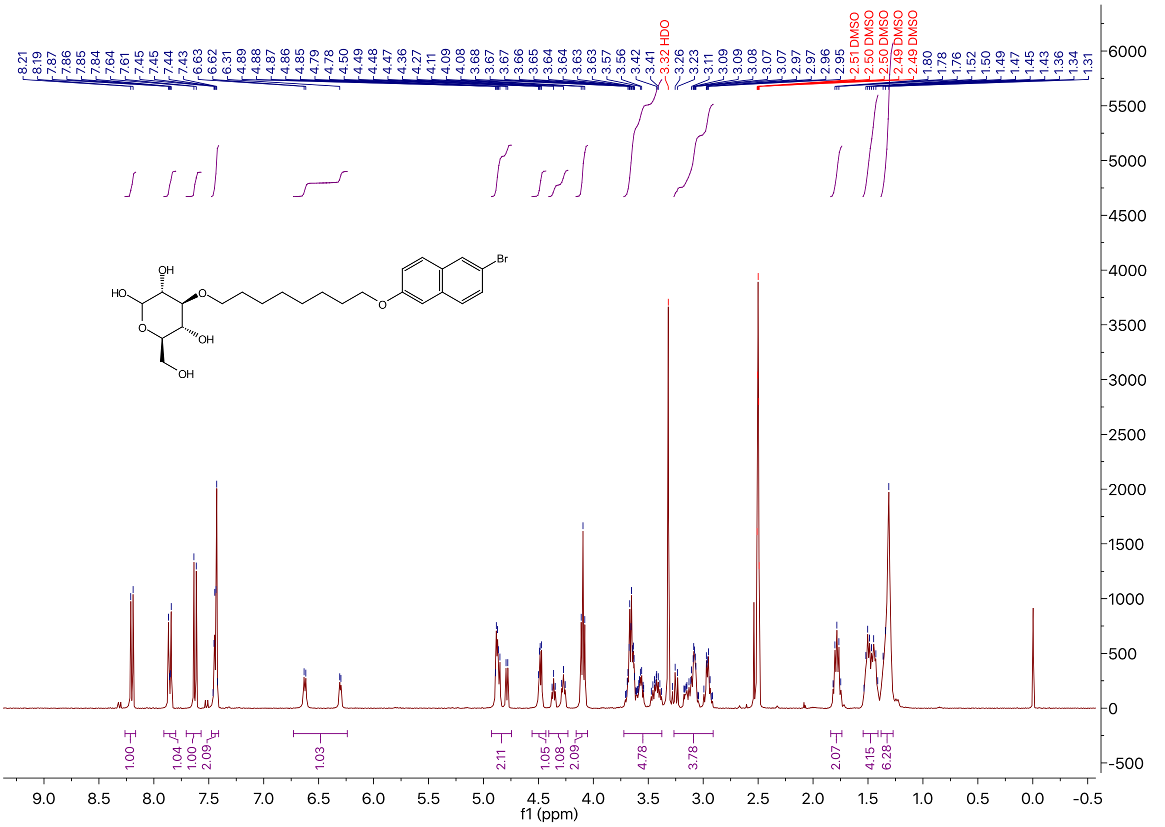

$^{13}C$ NMR of Compound **3b** in DMSO-$d_6$

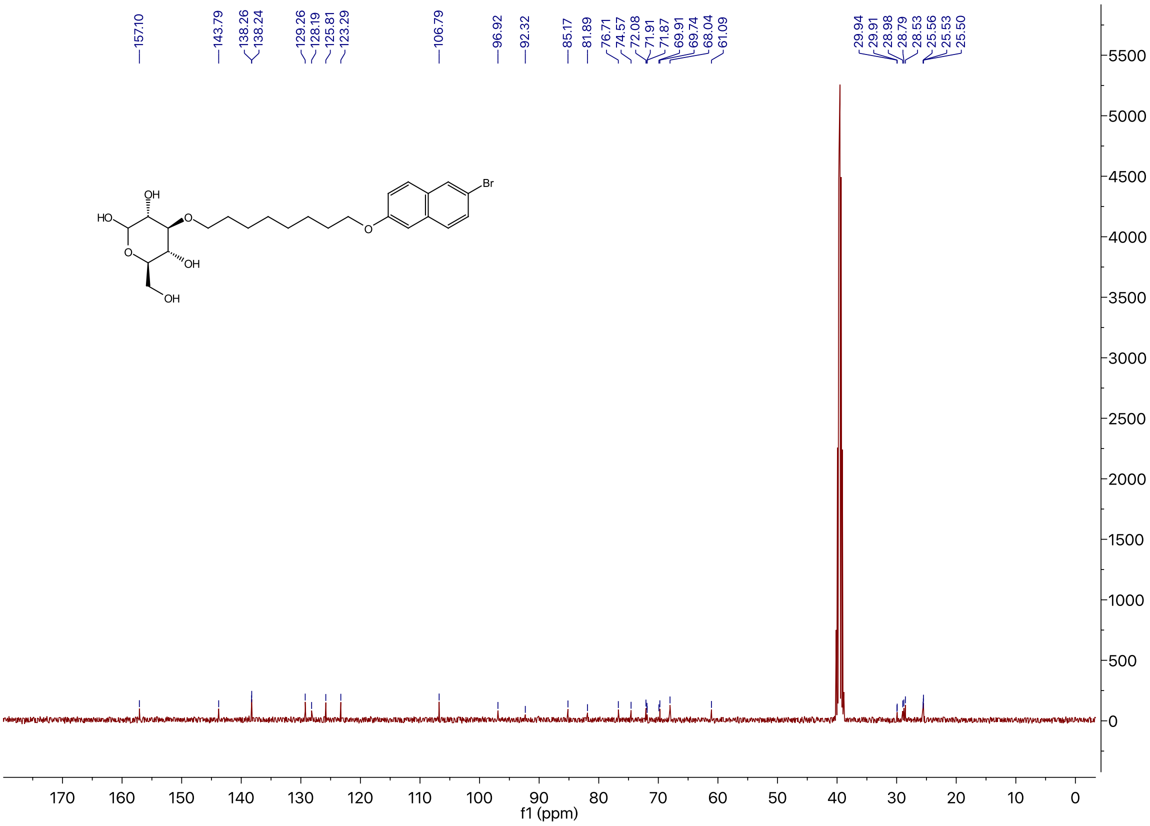

$^{1}H$ NMR of Compound **3c** in DMSO-$d_6$

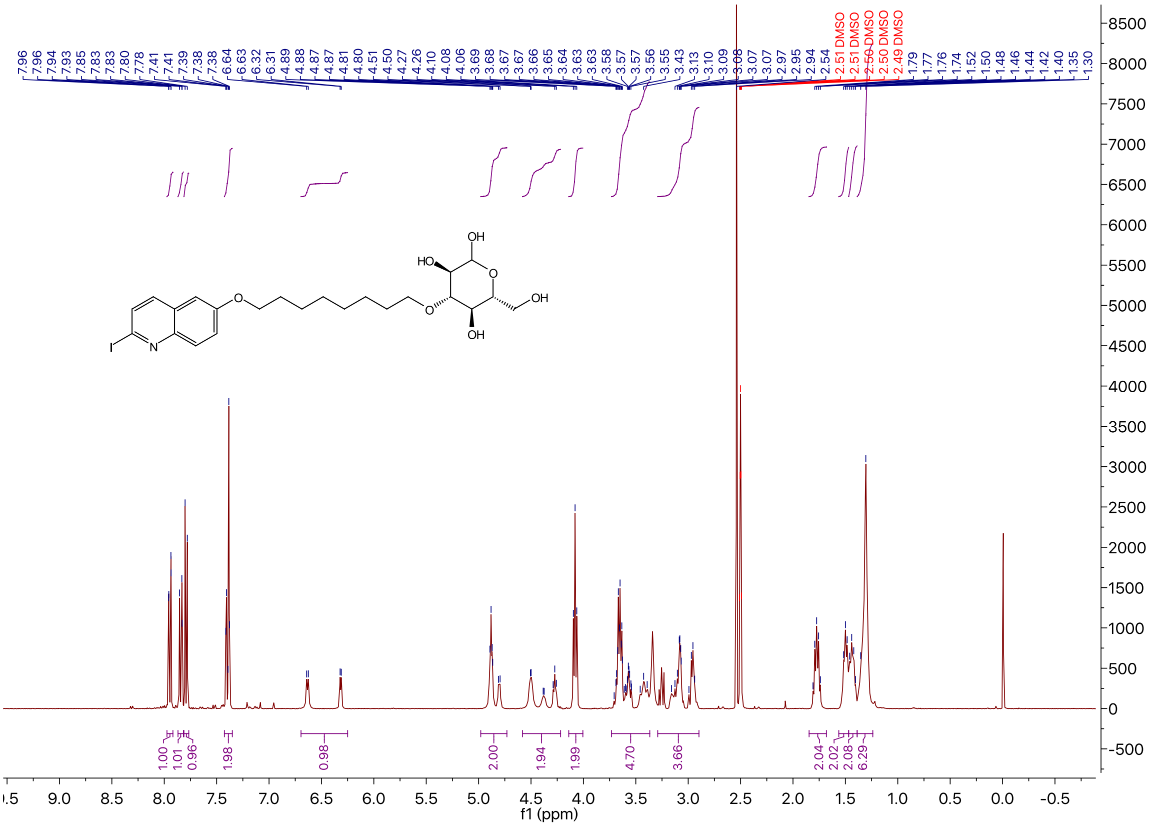

### $^{13}$C NMR of Compound **3c** in DMSO-$d_6$

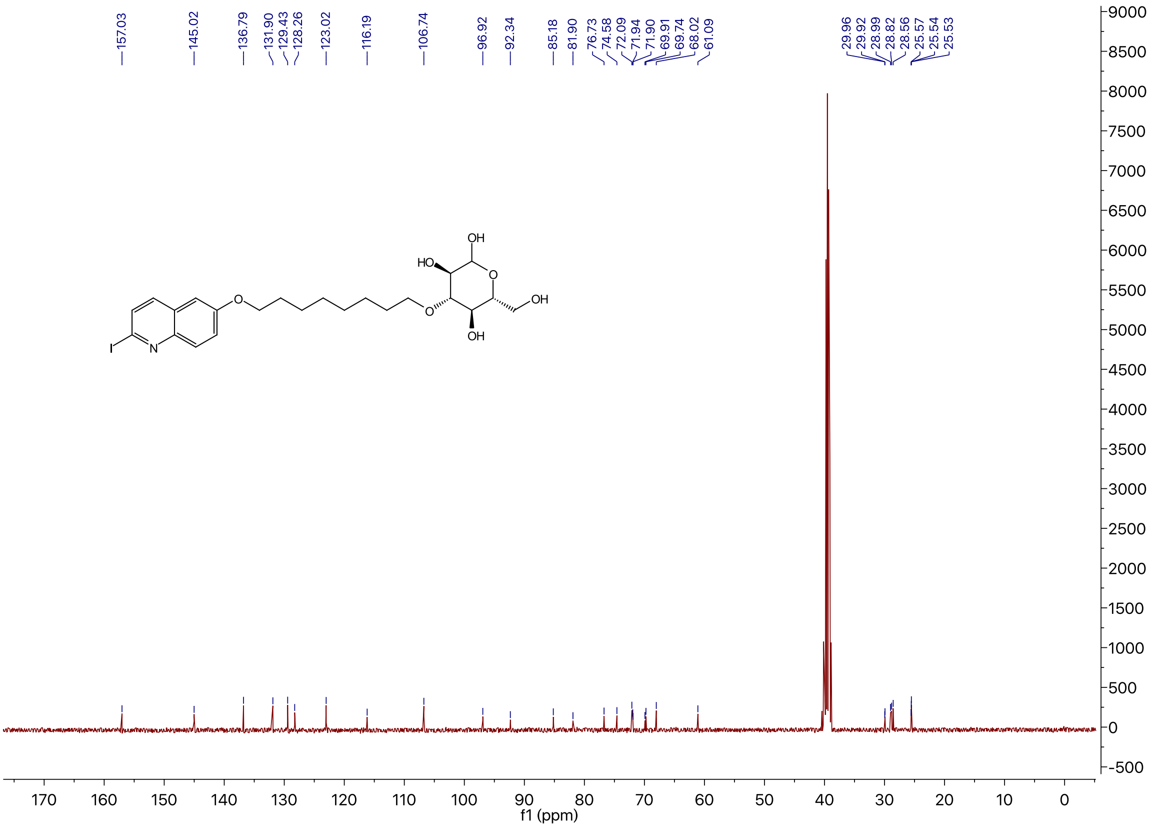

### $^{1}$H NMR of Compound **3d** in DMSO-$d_6$

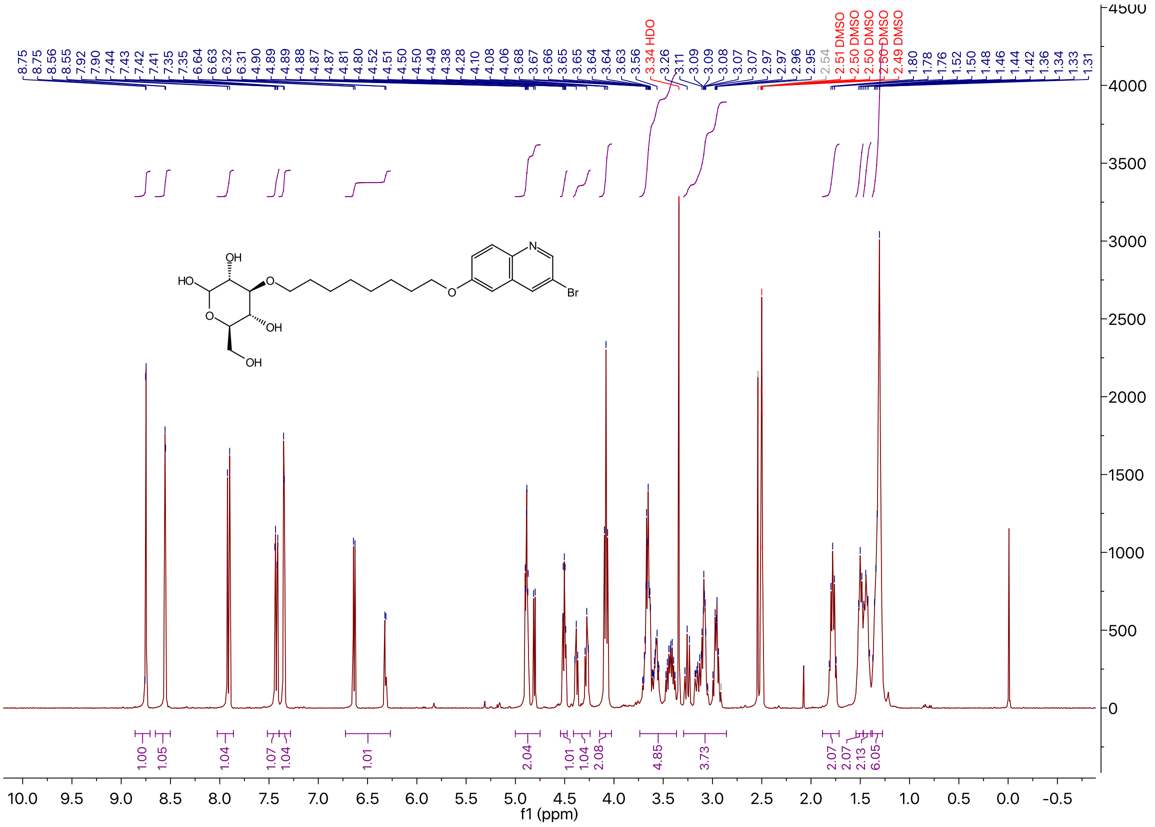

$^{13}C$ NMR of Compound **3d** in DMSO-$d_6$

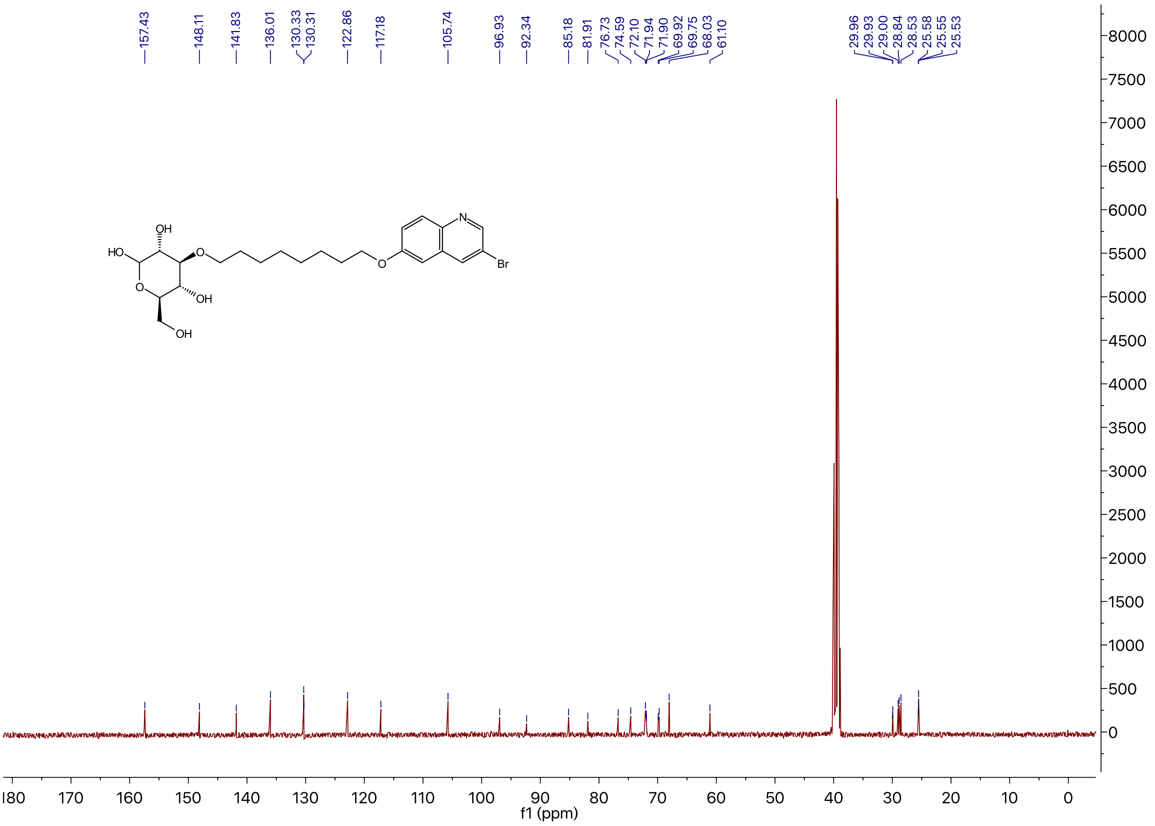

$^{1}H$ NMR of Compound **3e** in DMSO-$d_6$

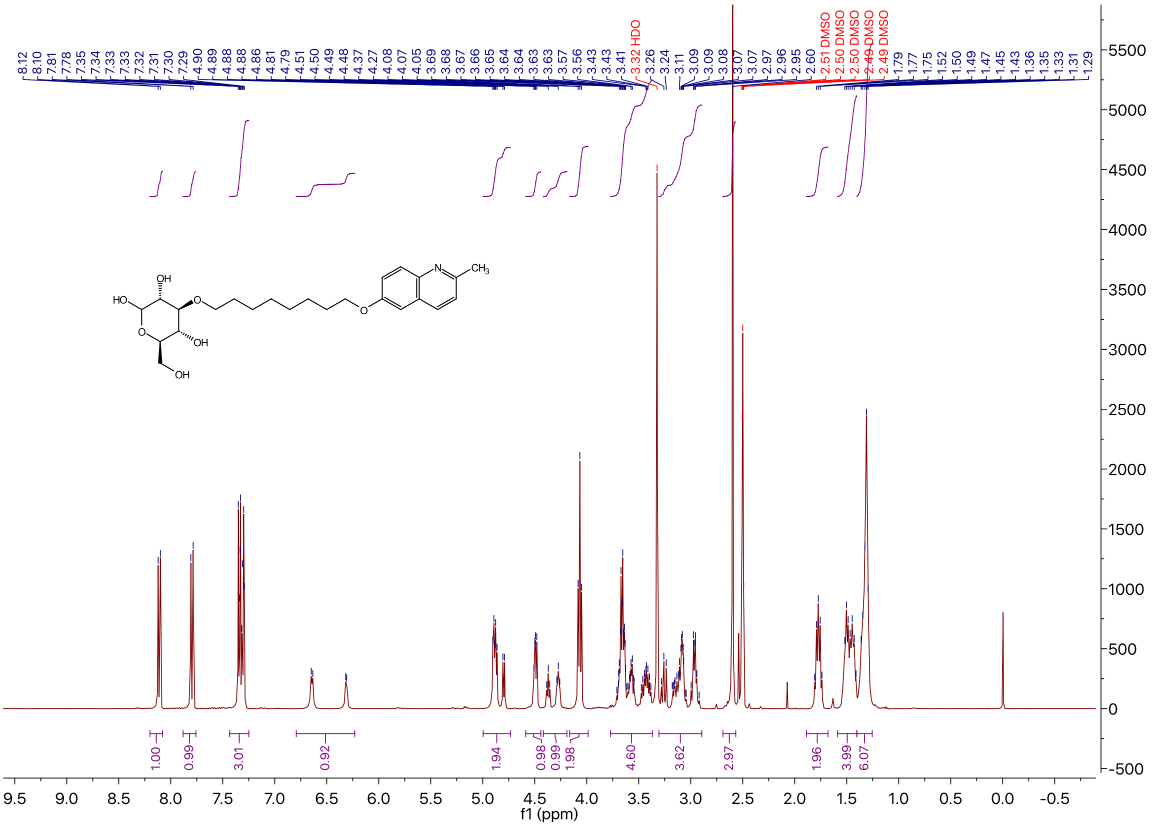

$^{13}$C NMR of Compound **3e** in DMSO-$d_6$

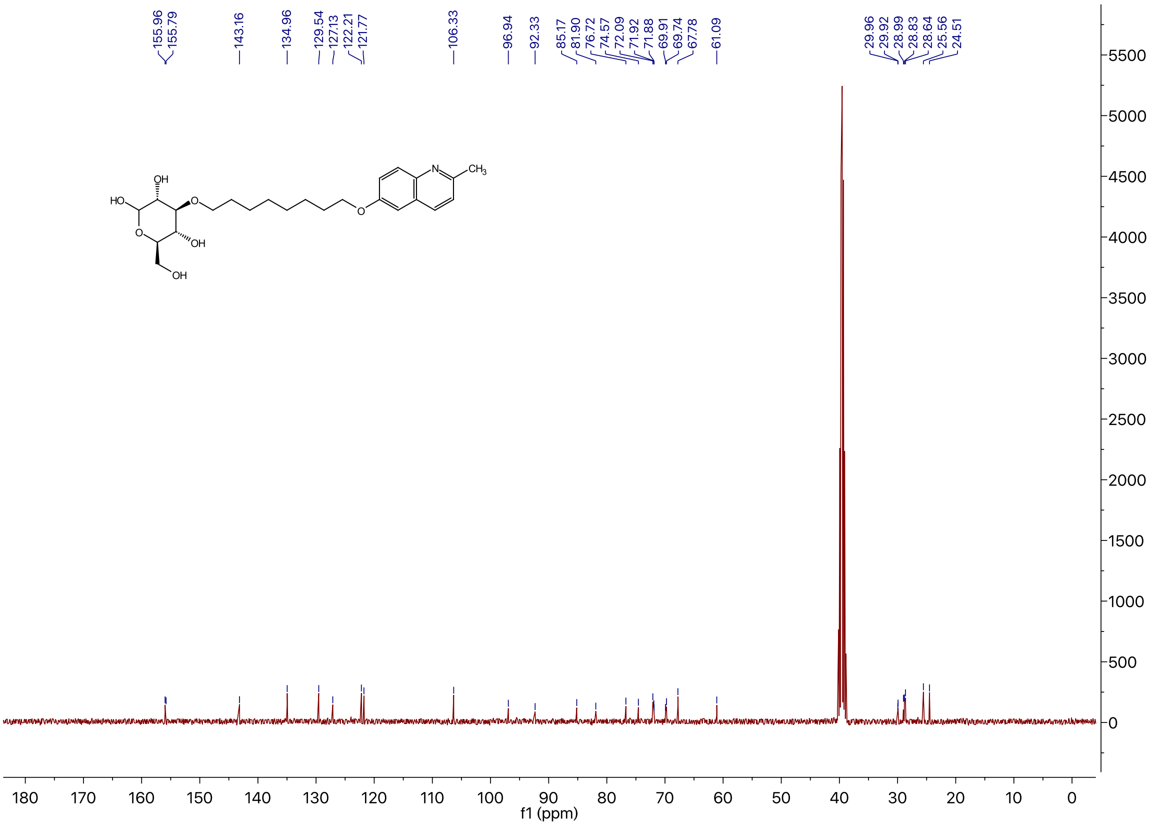

$^{1}$H NMR of Compound **3f** in DMSO-$d_6$

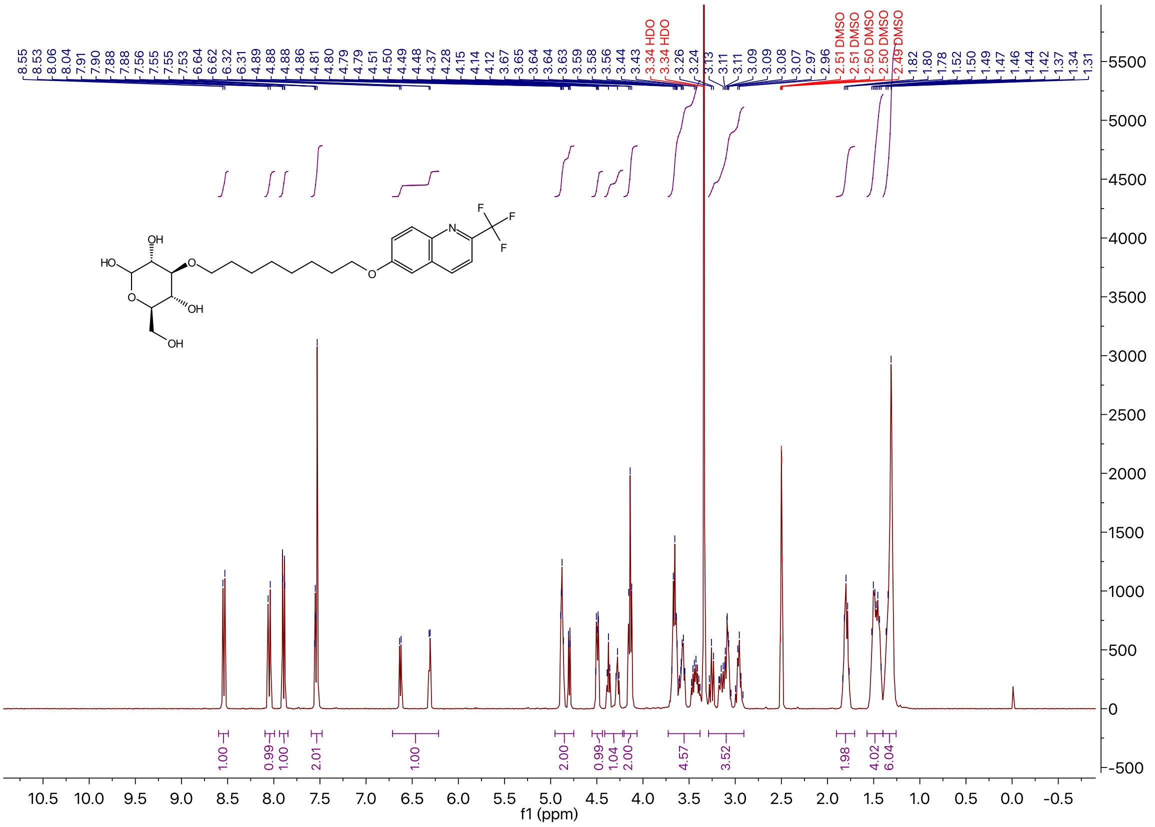

### $^{13}$C NMR of Compound **3f** in DMSO-*d*$_6$

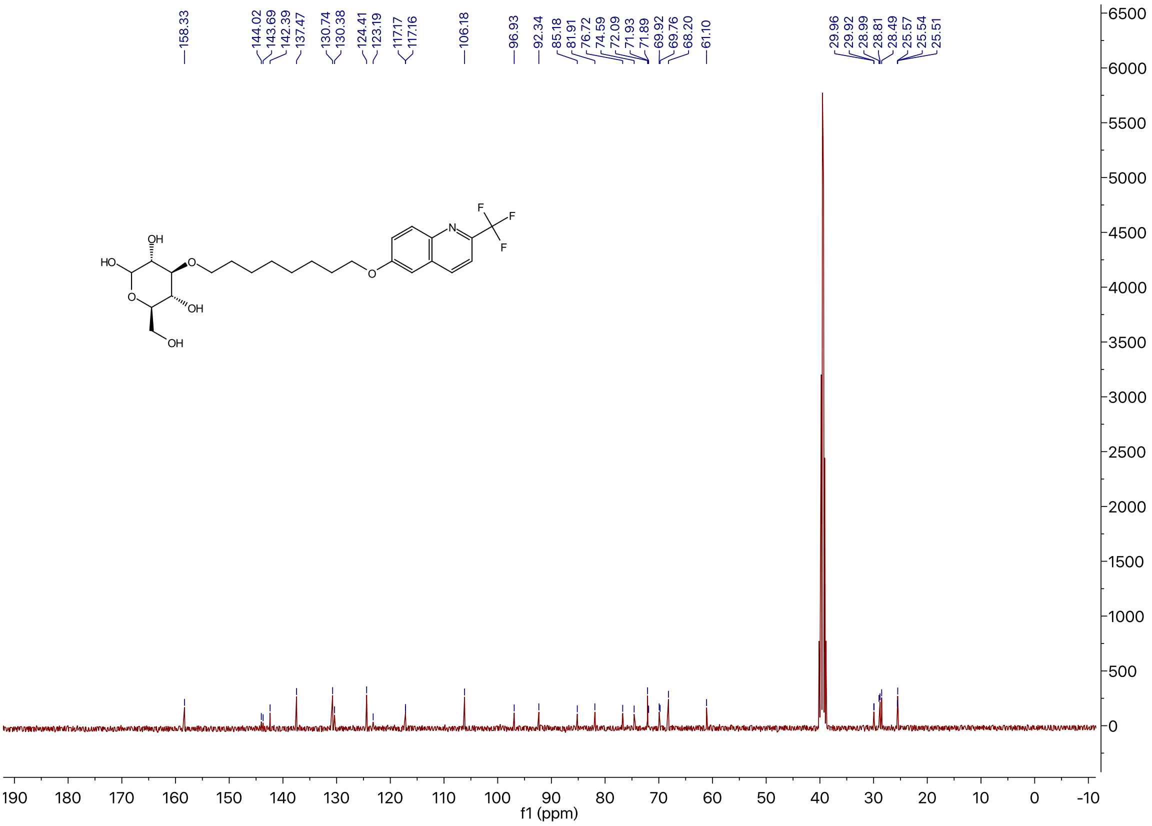

### $^{1}$H NMR of Compound **3g** in DMSO-*d*$_6$

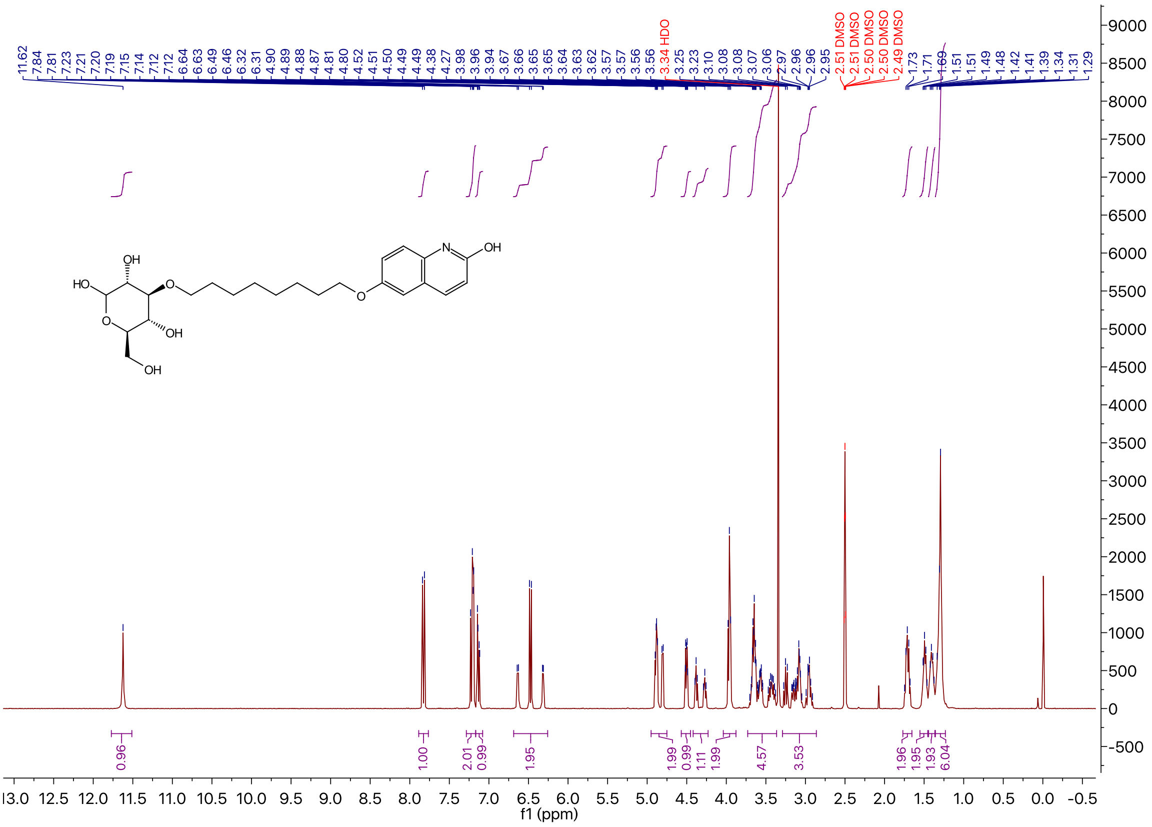

### $^{13}C$ NMR of Compound **3g** in DMSO-$d_6$

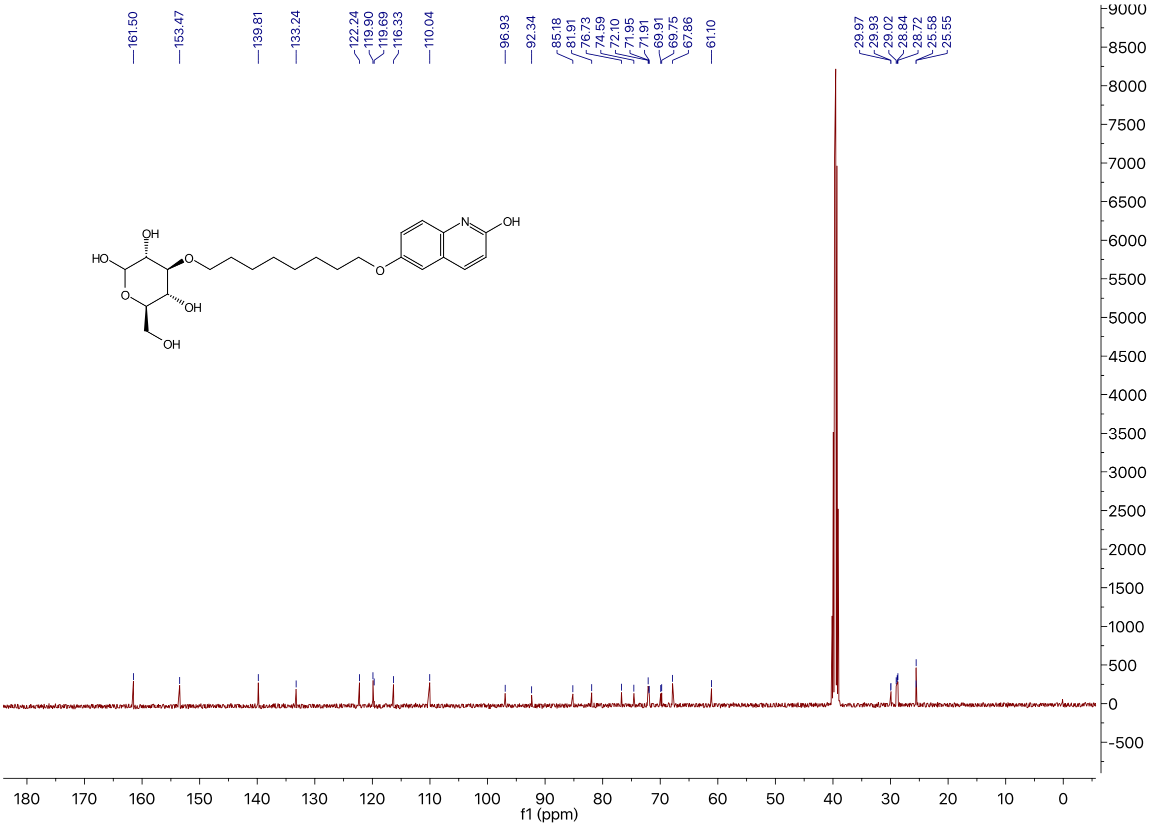

### $^{1}H$ NMR of Compound **3h** in DMSO-$d_6$

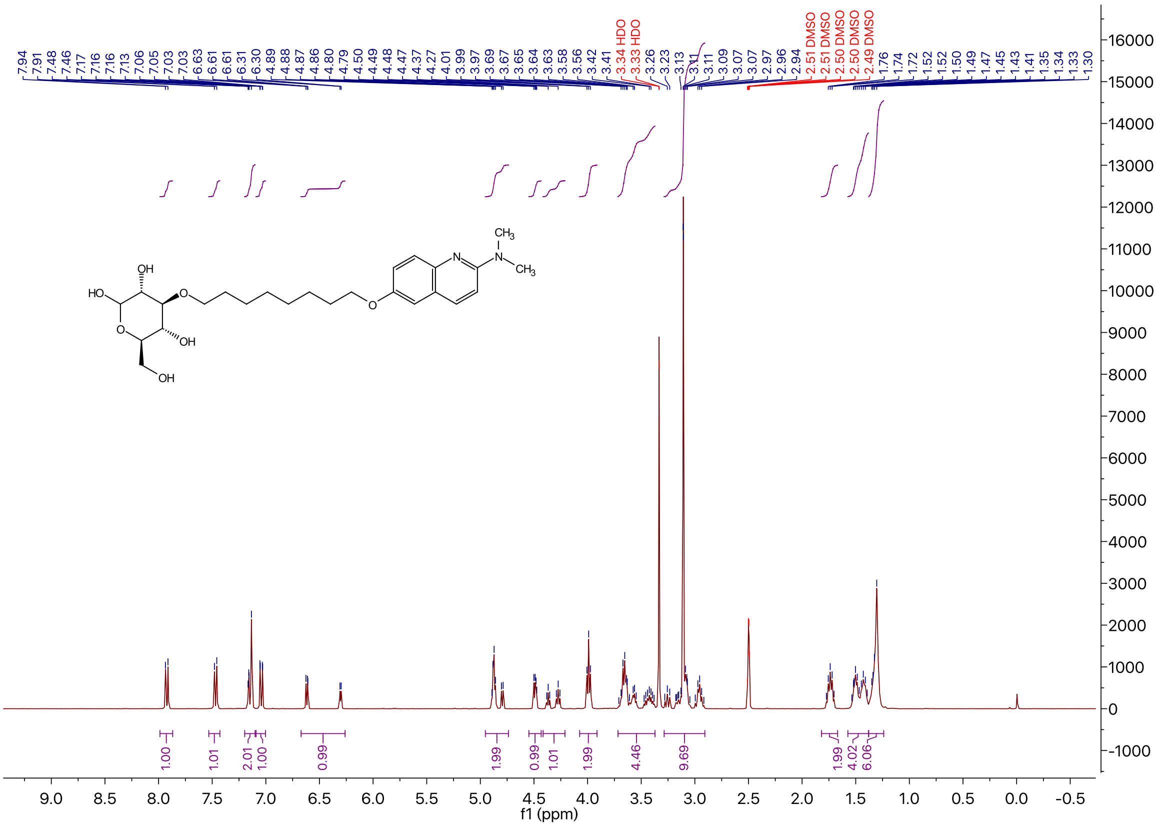

$^{13}$C NMR of Compound **3h** in DMSO-$d_6$

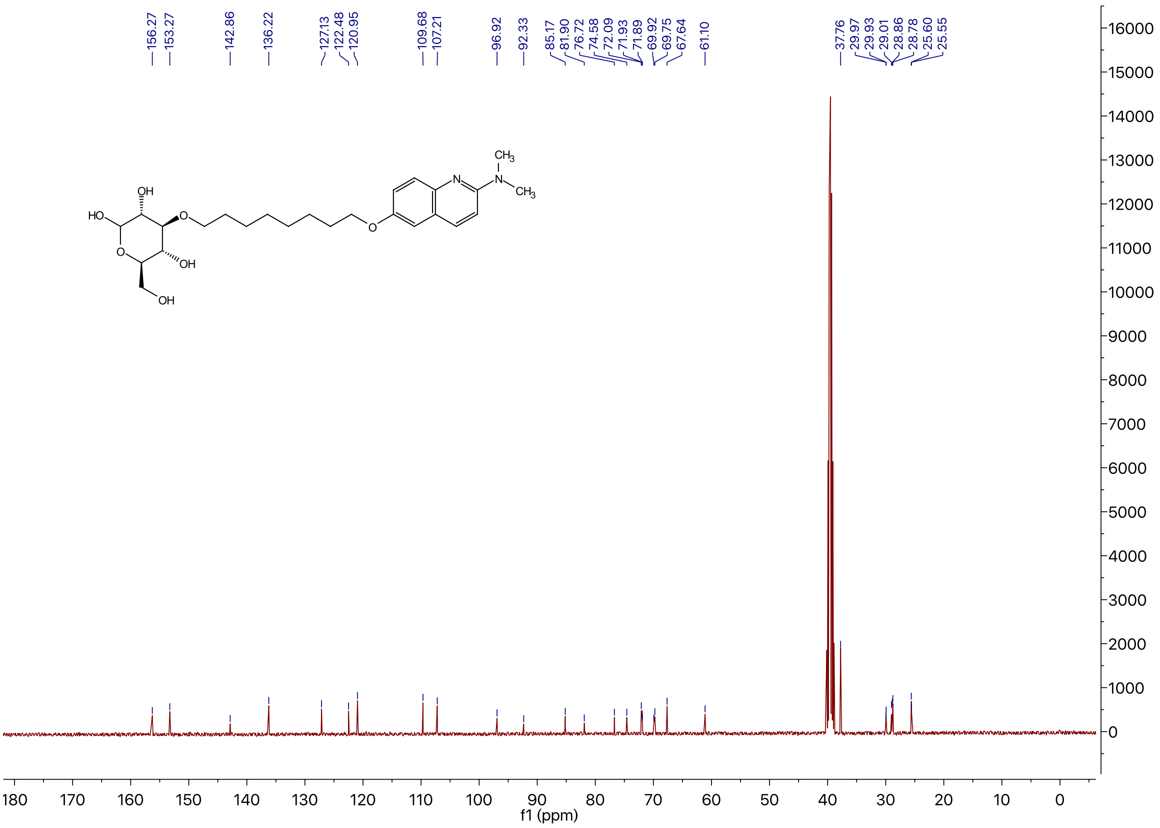

$^{1}$H NMR of Compound **12** in $CDCl_3$

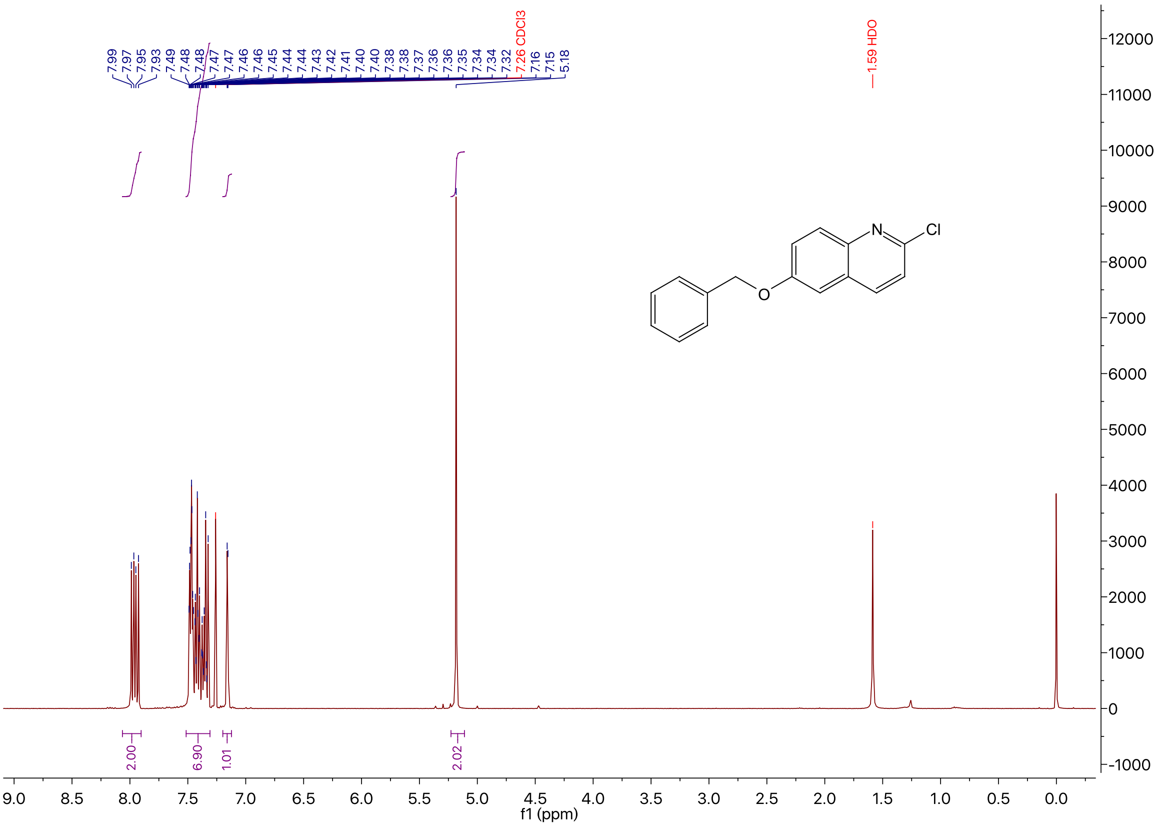

$^{13}C$ NMR of Compound **12** in $CDCl_3$

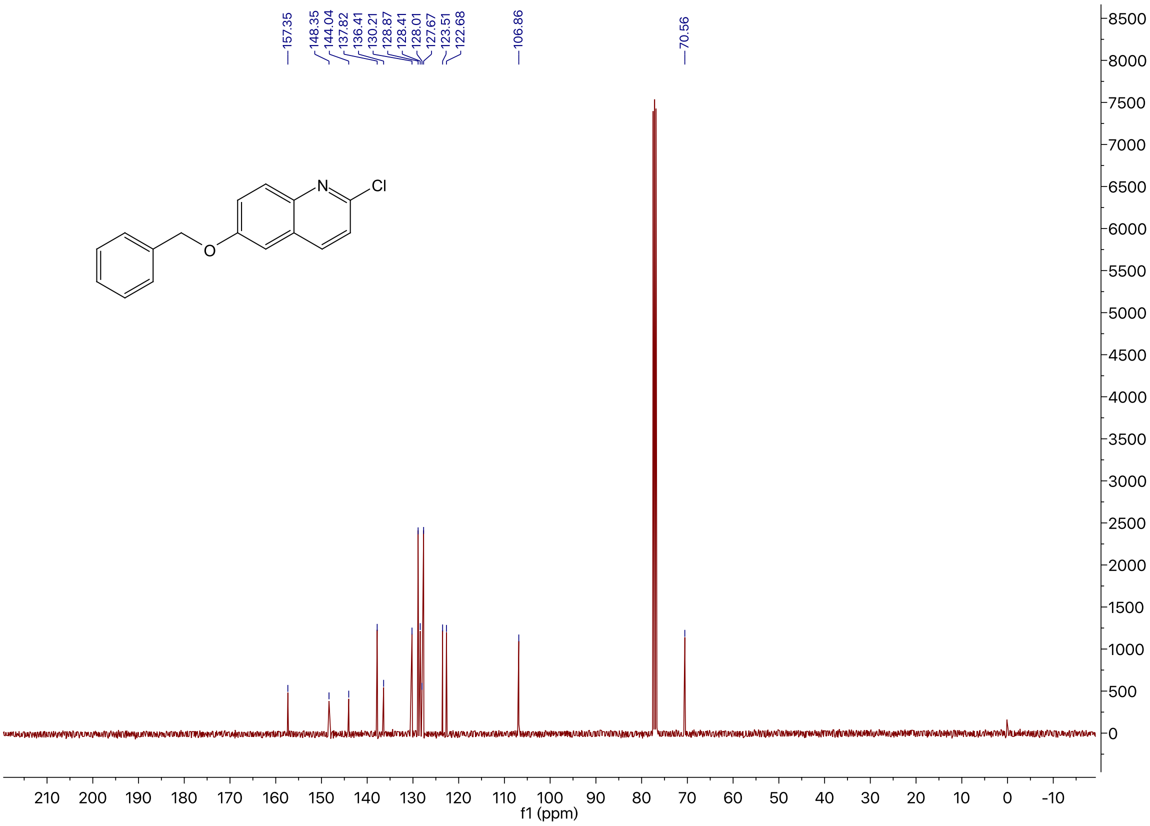

$^{1}H$ NMR of Compound **13** in $CDCl_3$

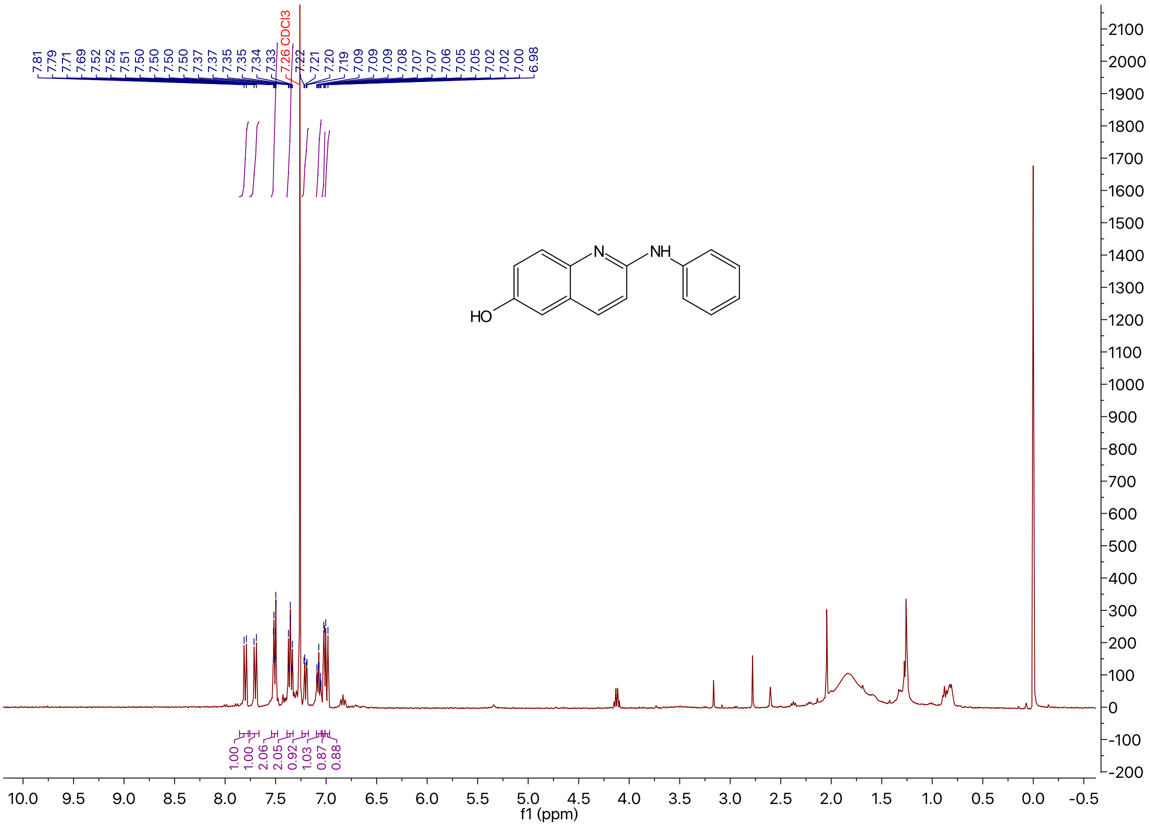

$^{1}H$ NMR of Compound **3m** in DMSO-$d_6$

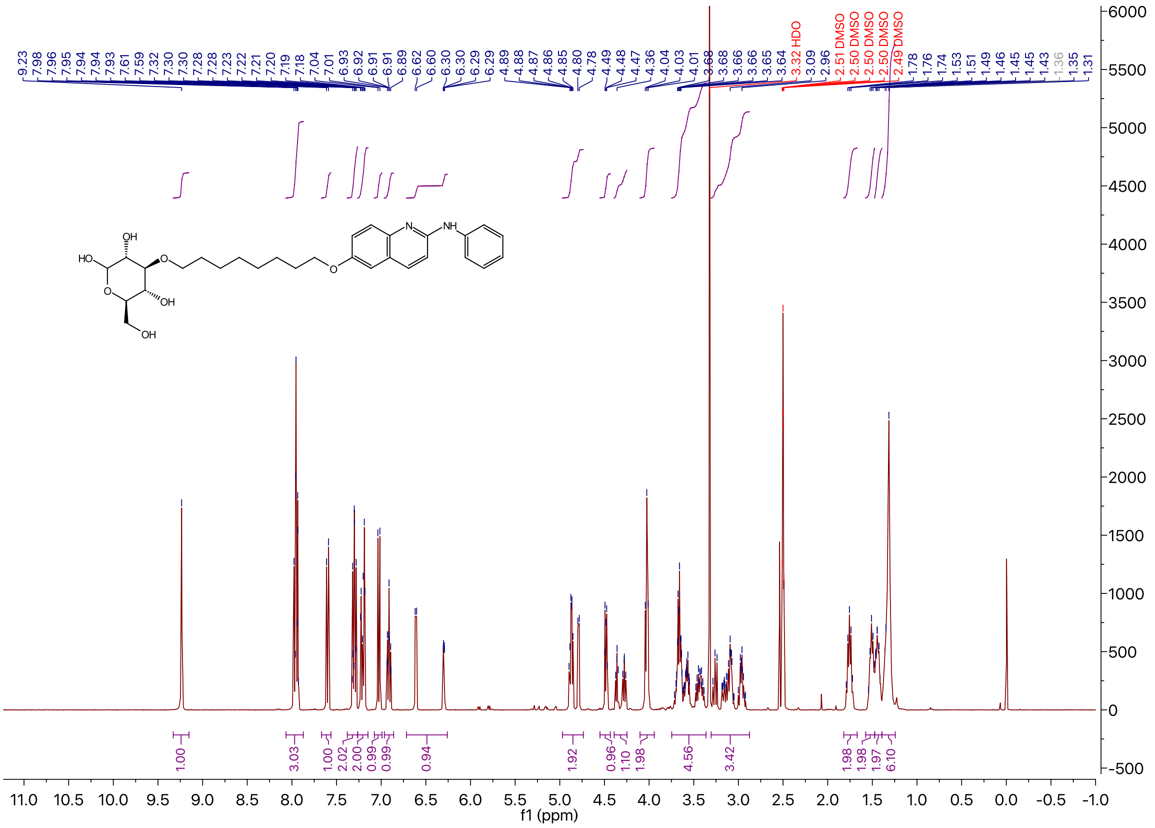

$^{13}C$ NMR of Compound **3m** in DMSO-$d_6$

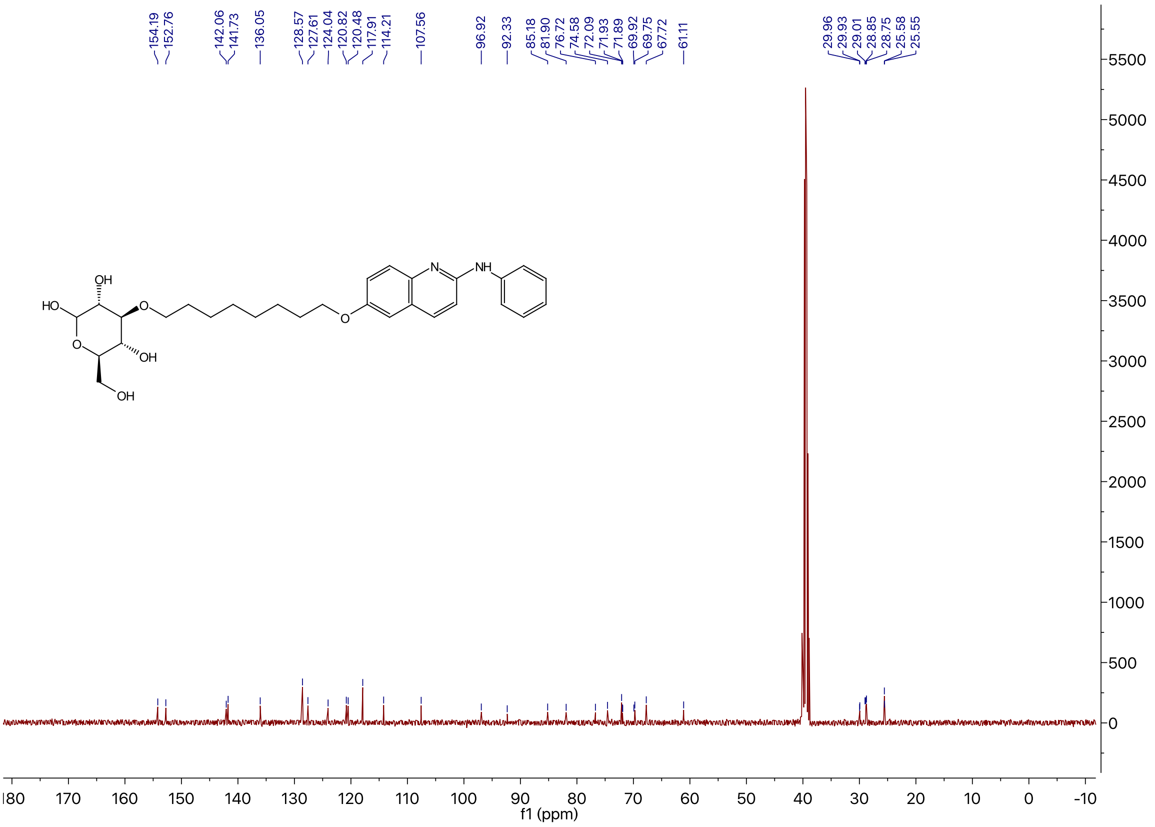

### $^{1}$H NMR of Compound **3l** in DMSO-$d_6$

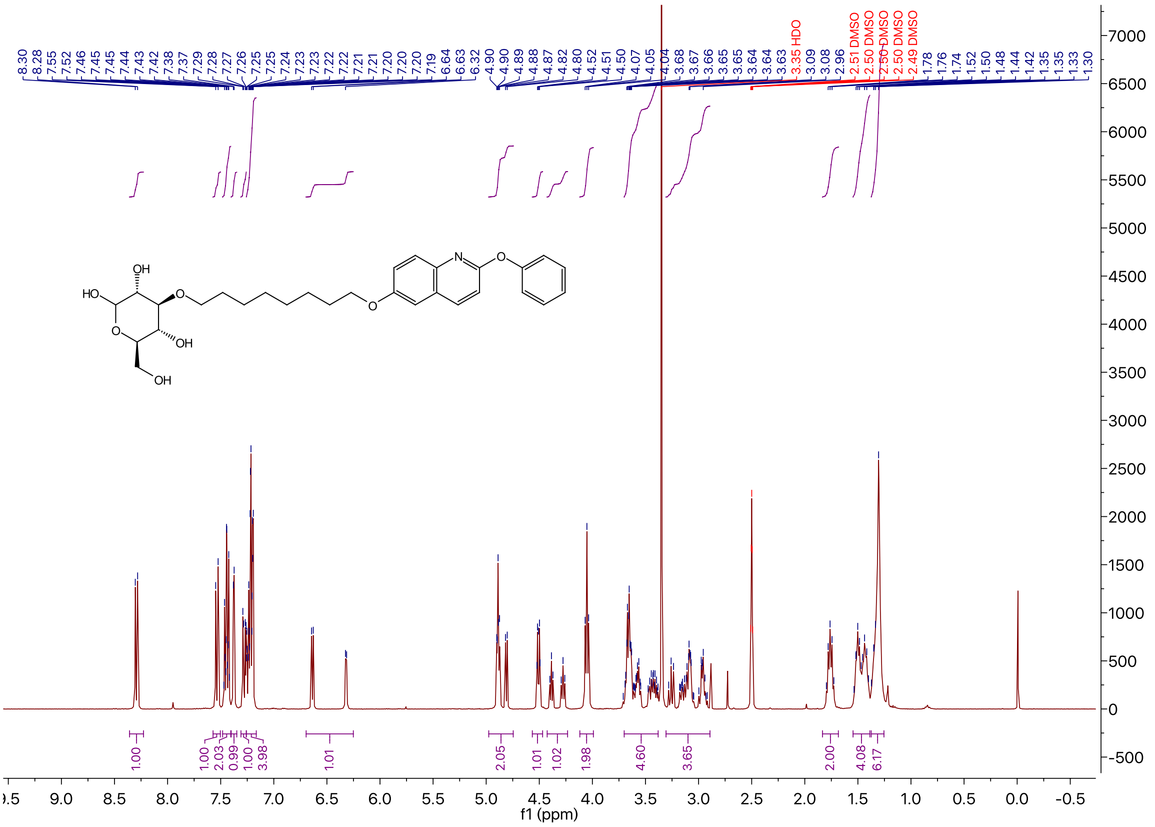

### $^{13}$C NMR of Compound **3l** in DMSO-$d_6$

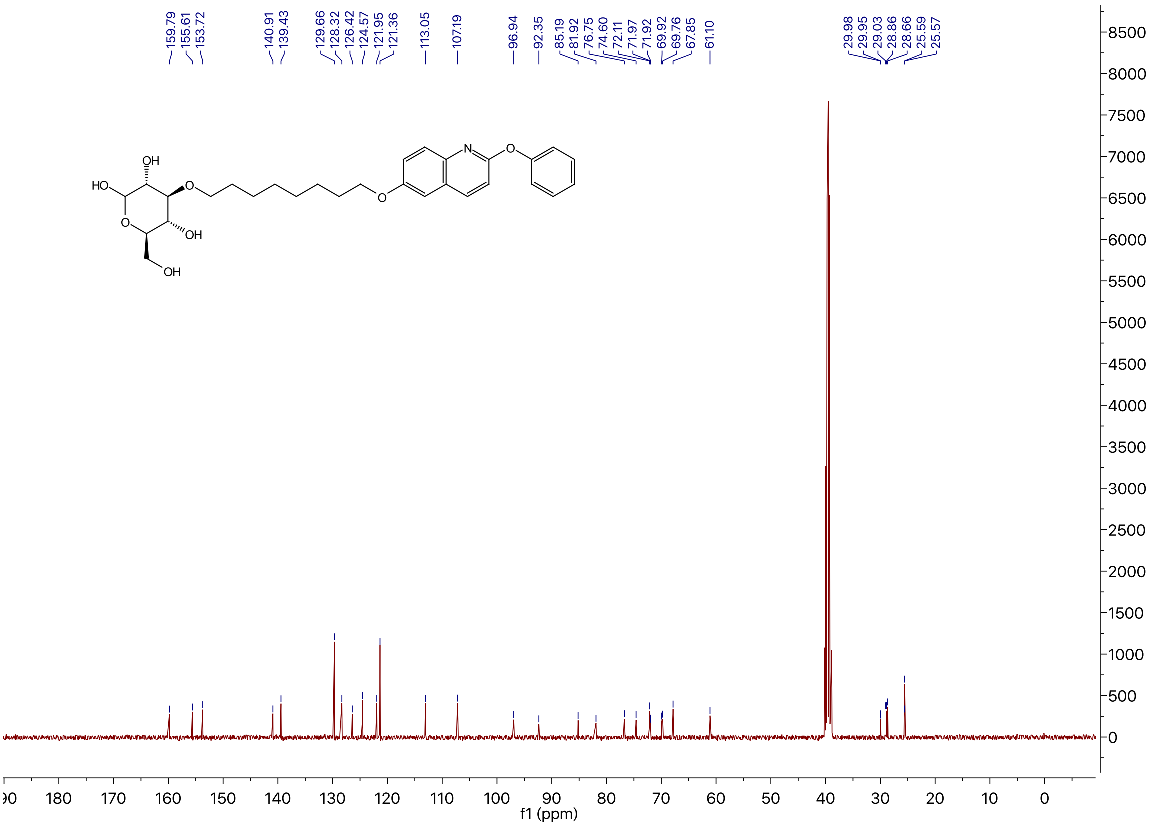

### $^1$H NMR of Compound **3n** in DMSO-$d_6$

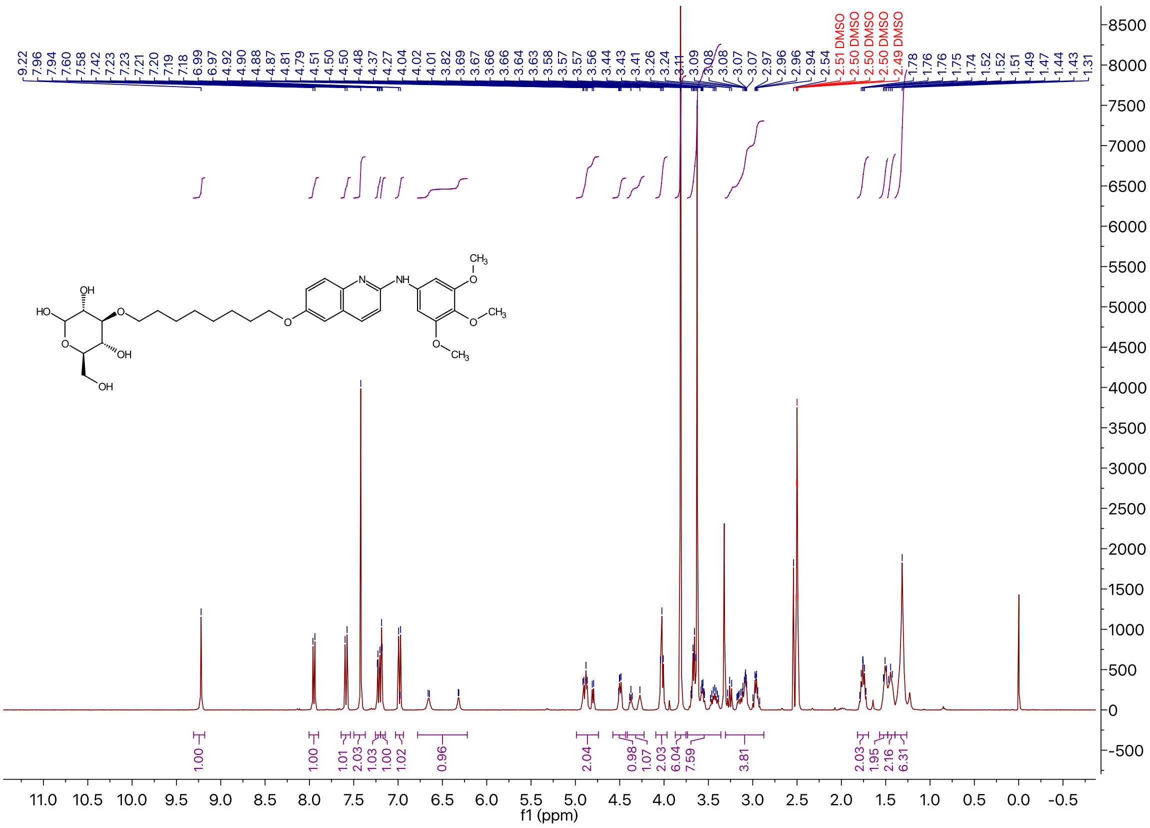

### $^{13}$C NMR of Compound **3n** in DMSO-$d_6$

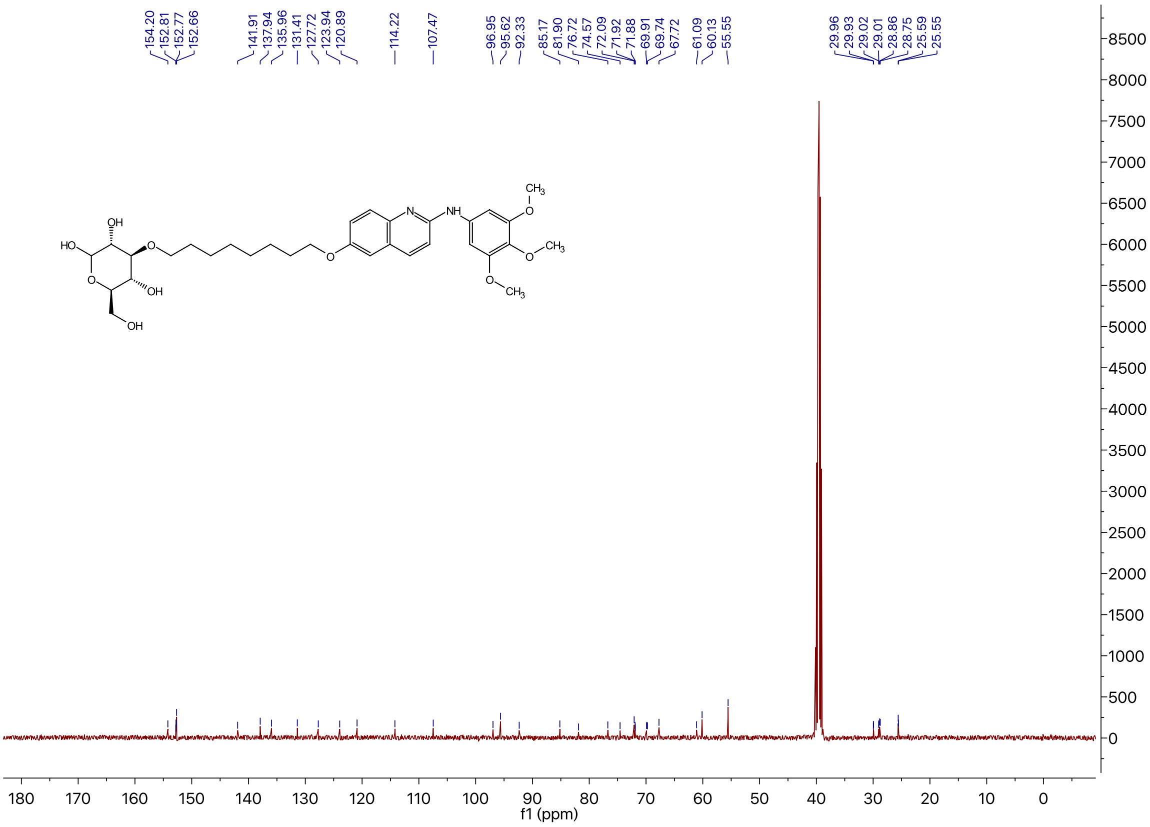

### [1]H NMR of Compound **3i** in DMSO-$d_6$

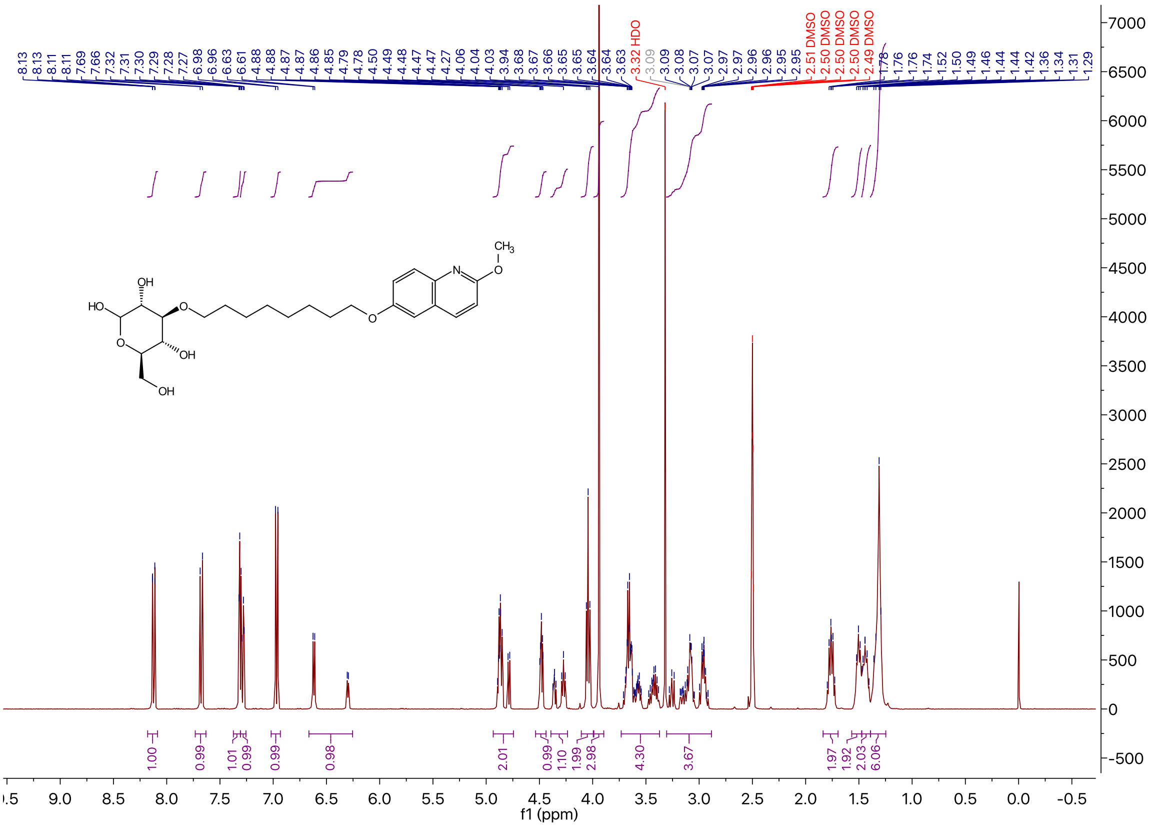

### [13]C NMR of Compound **3i** in DMSO-$d_6$

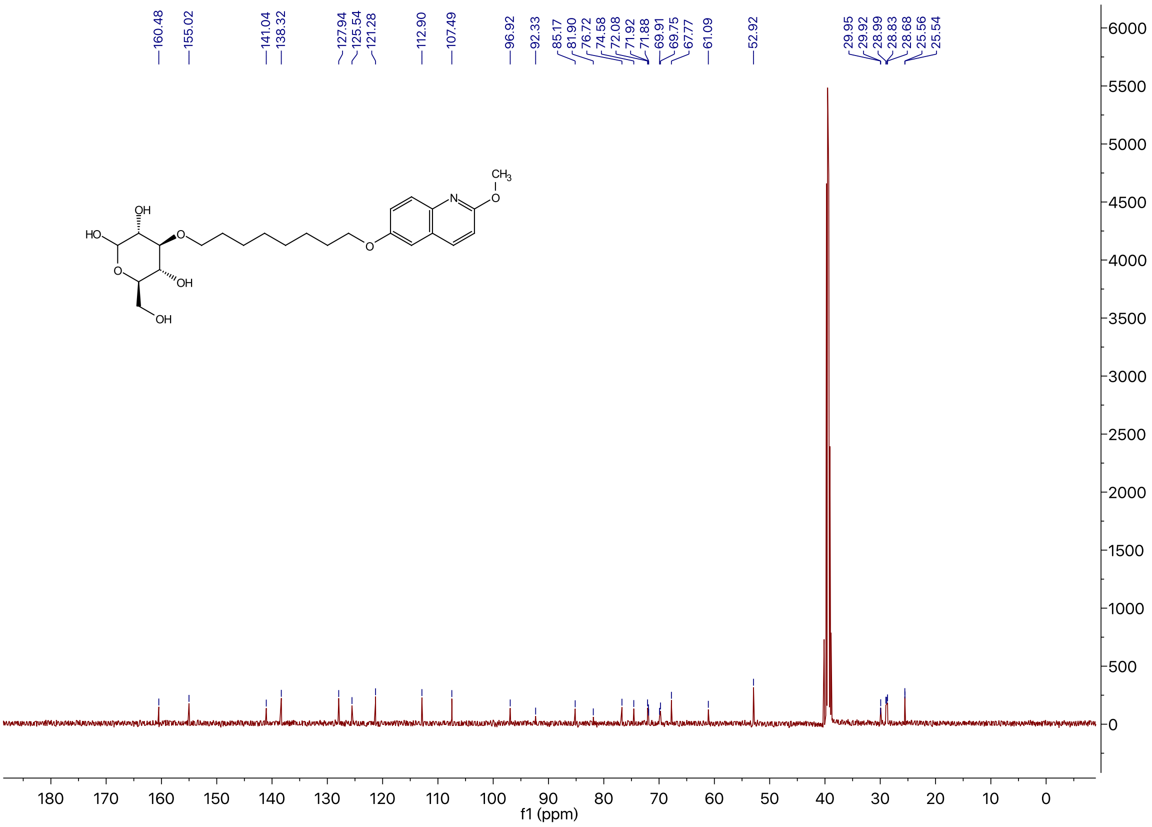

[1]H NMR of Compound **3j** in DMSO-$d_6$

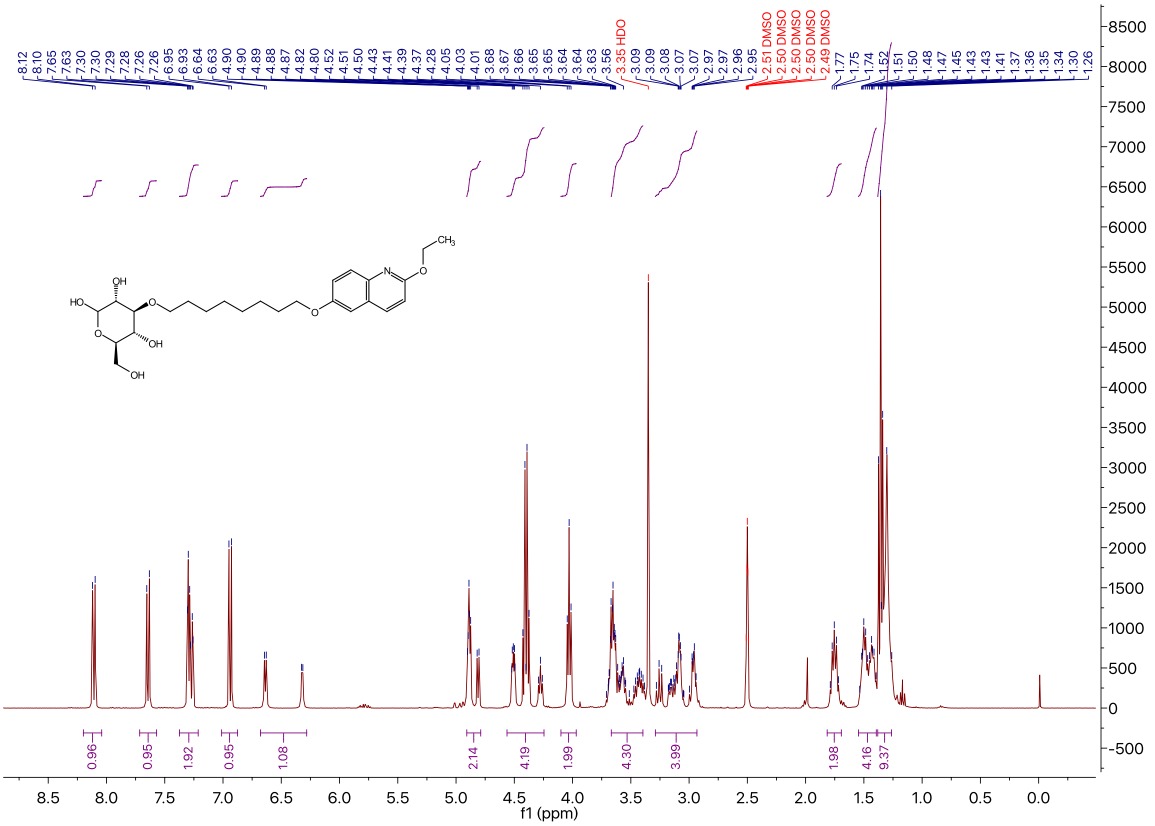

[13]C NMR of Compound **3j** in DMSO-$d_6$

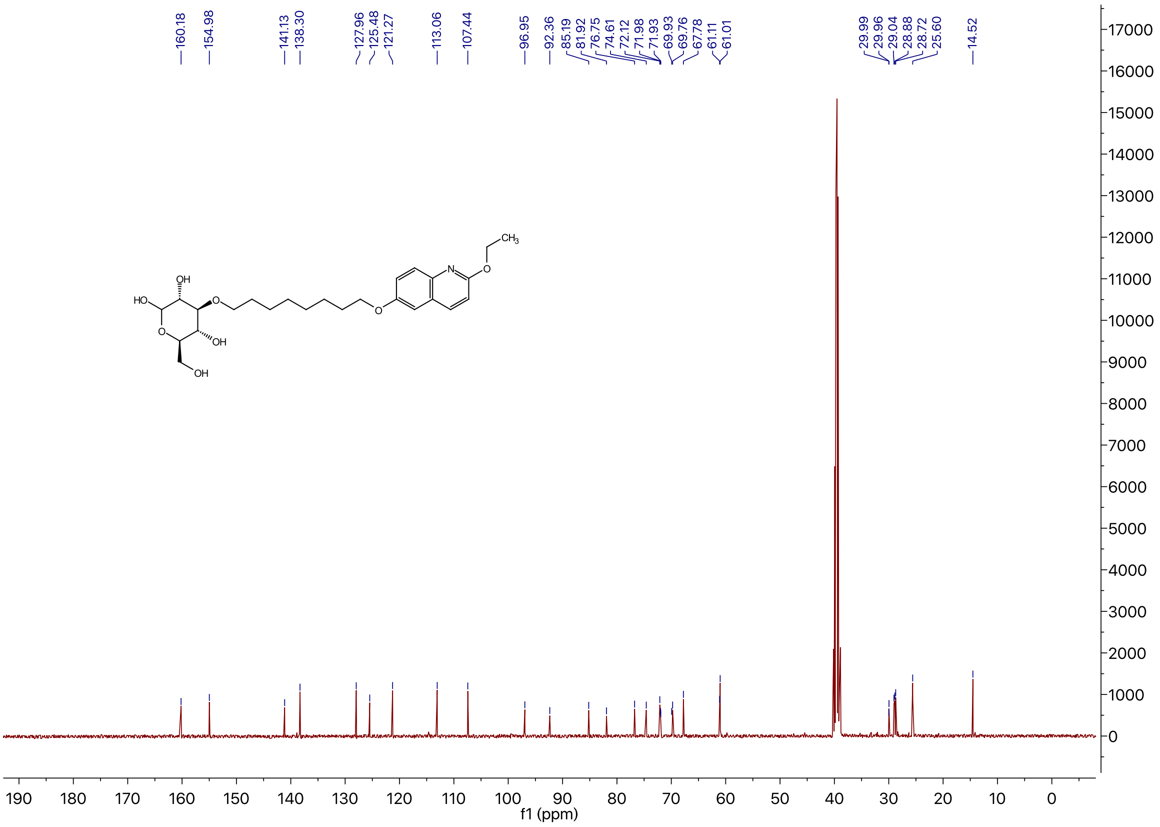

[1]H NMR of Compound **3k** in DMSO-$d_6$

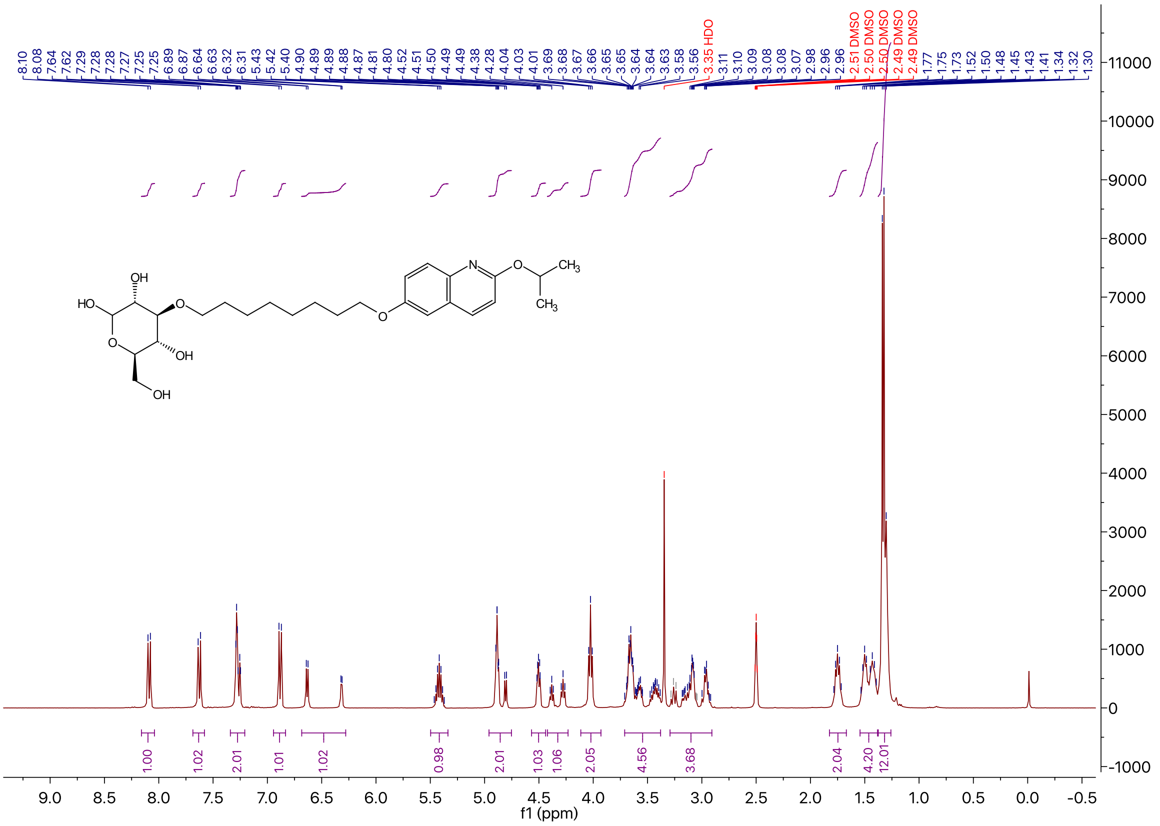

[13]C NMR of Compound **3k** in DMSO-$d_6$

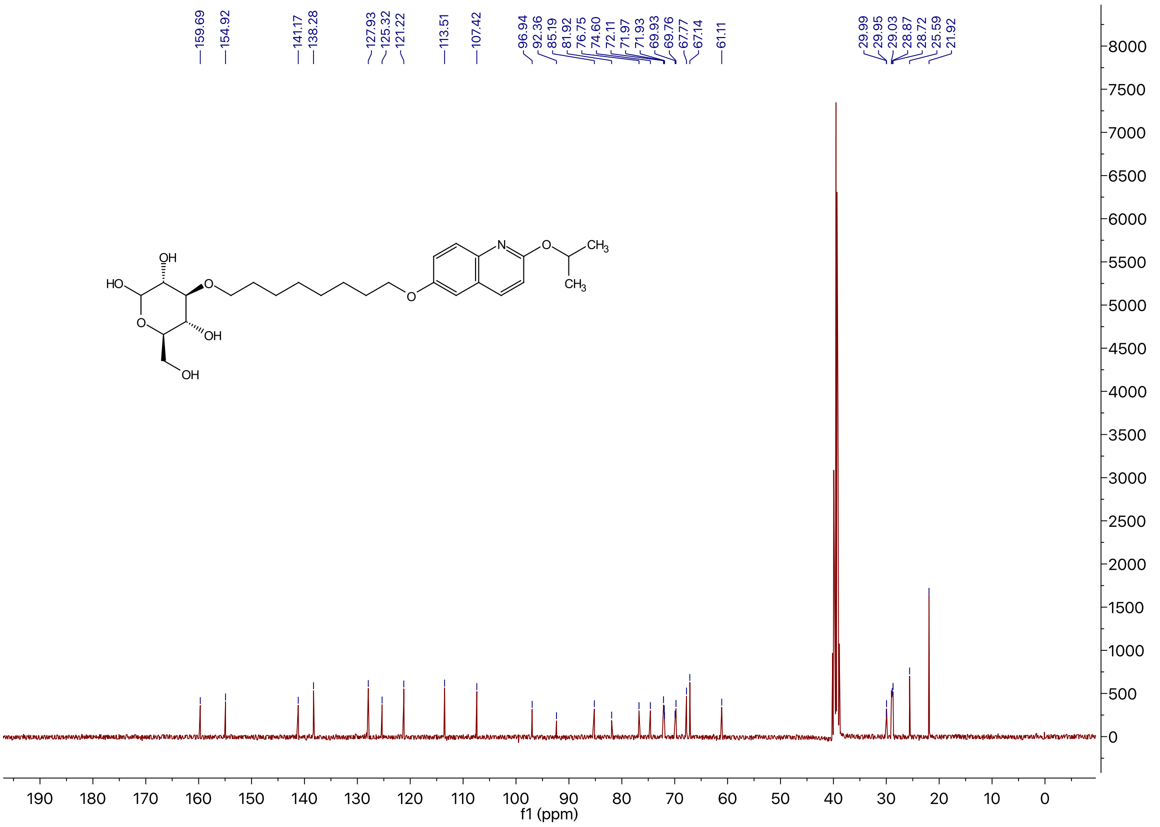

$^{1}H$ NMR of Compound **4a** in DMSO-$d_6$

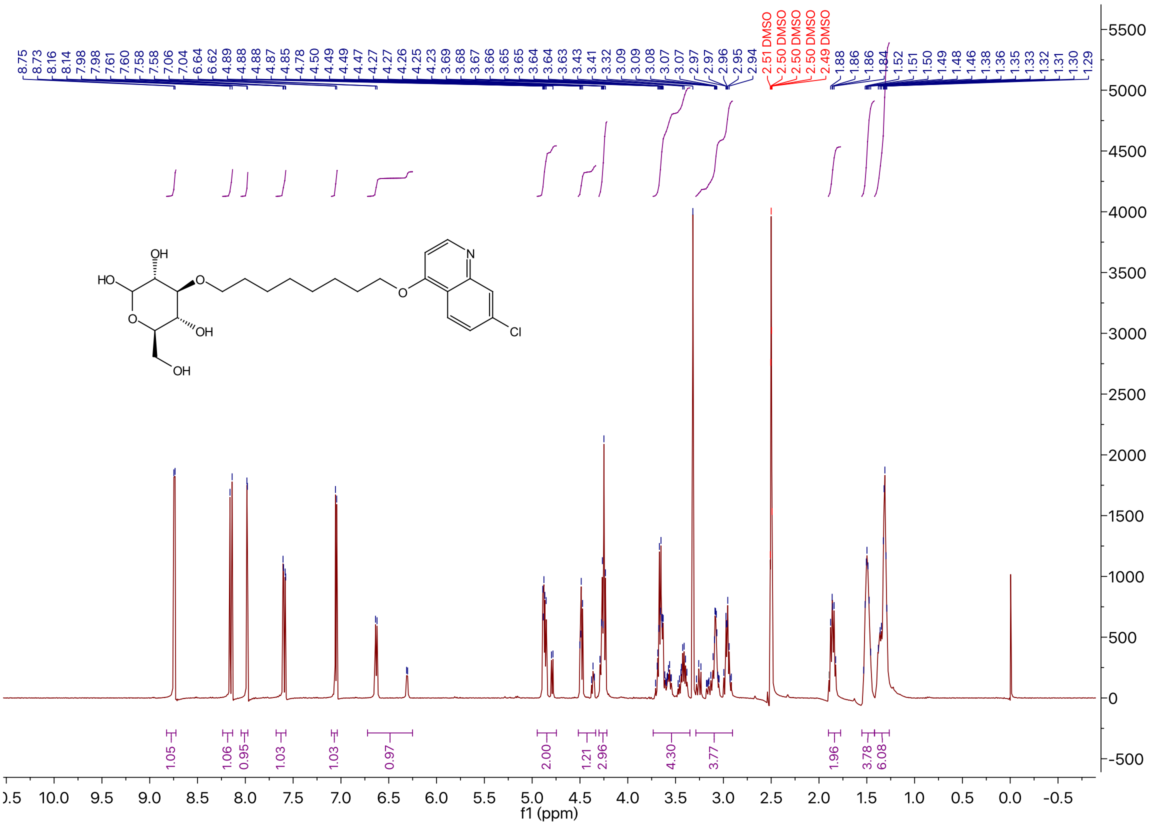

$^{13}C$ NMR of Compound **4a** in DMSO-$d_6$

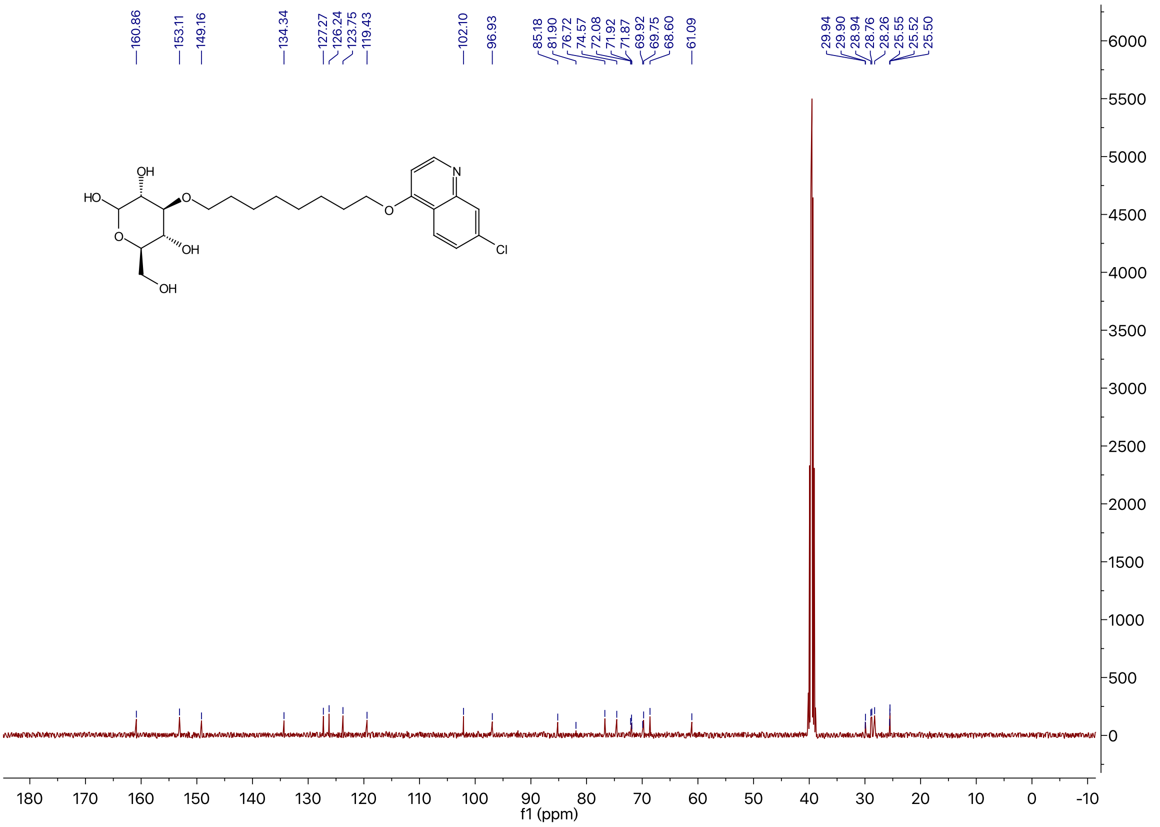

### $^{1}$H NMR of Compound **4b** in DMSO-$d_6$

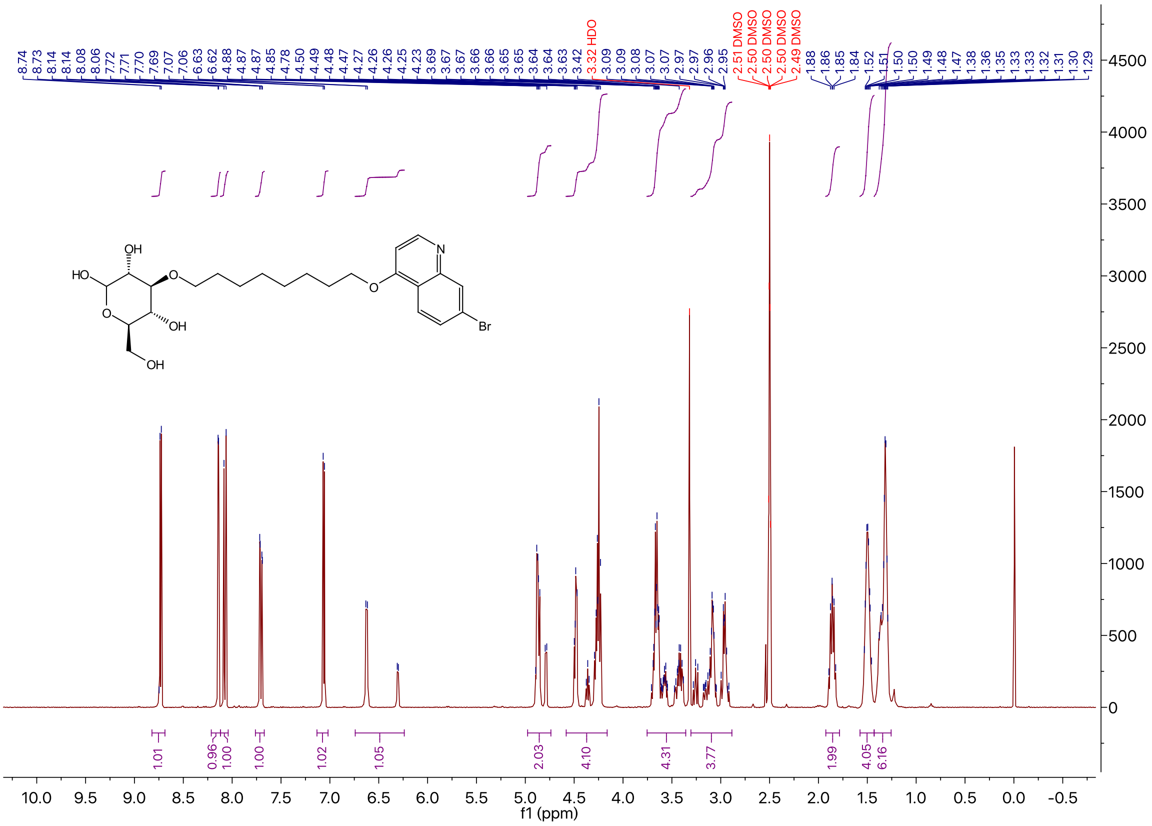

### $^{13}$C NMR of Compound **4b** in DMSO-$d_6$

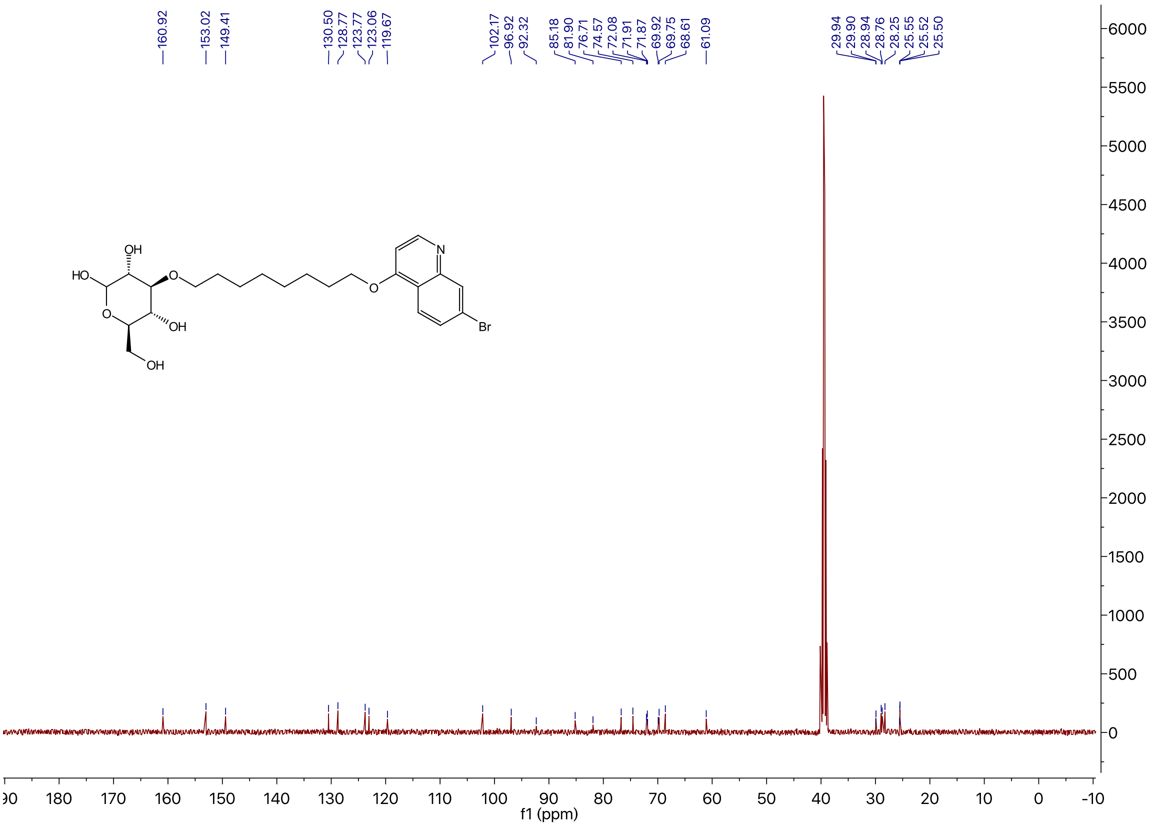

$^{1}H$ NMR of Compound **4c** in DMSO-$d_6$

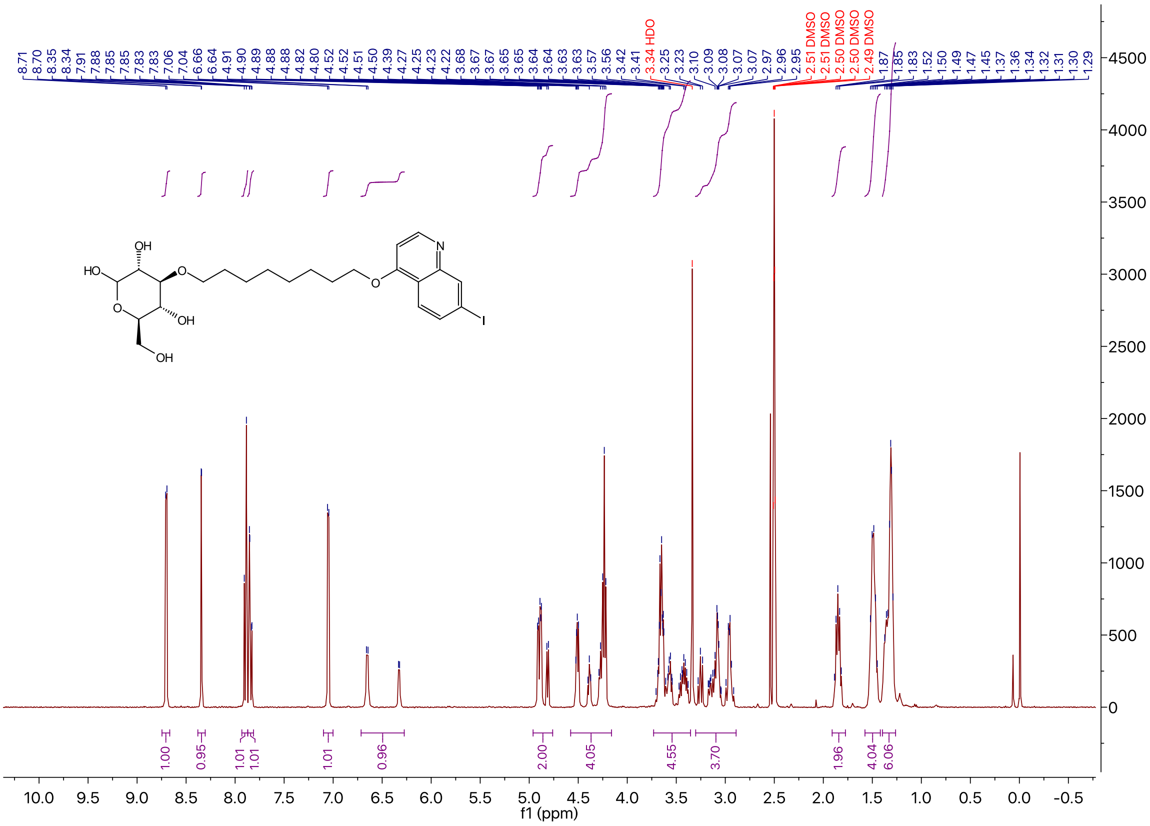

$^{13}C$ NMR of Compound **4c** in DMSO-$d_6$

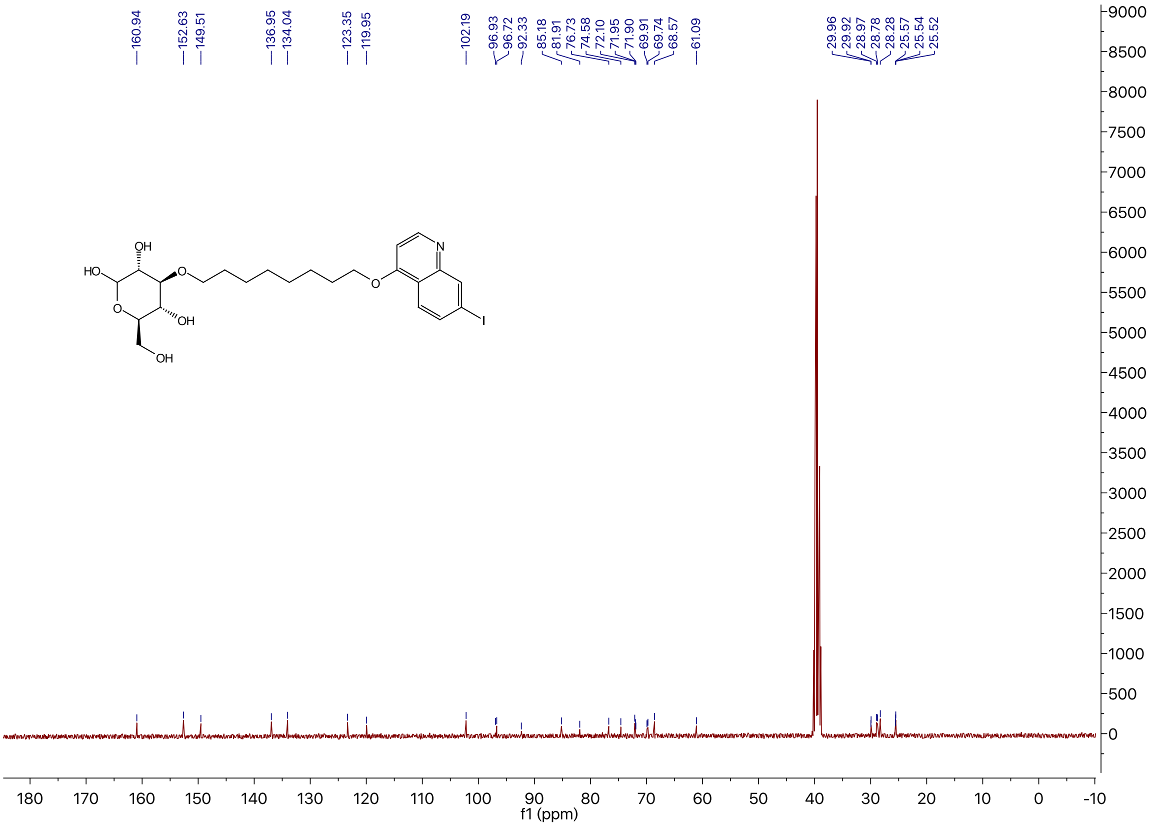

$^1$H NMR of Compound **4d** in DMSO-$d_6$

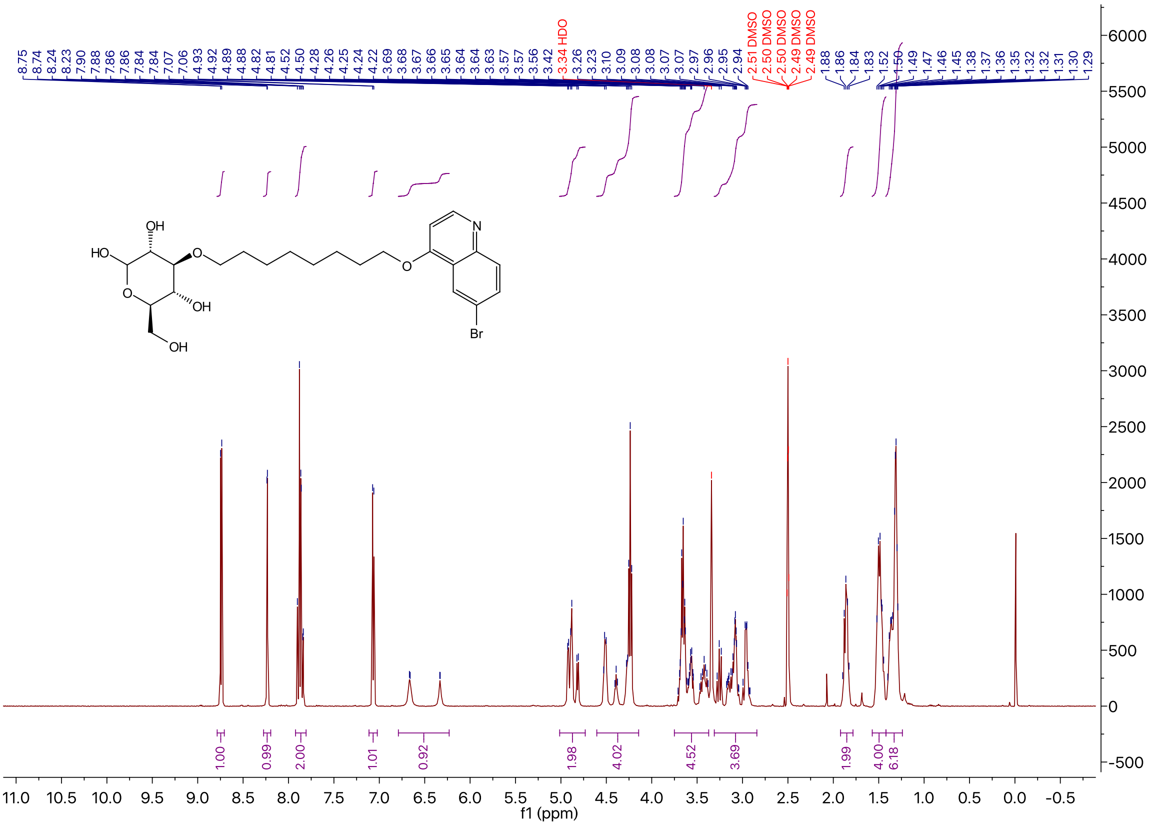

$^{13}$C NMR of Compound **4d** in DMSO-$d_6$

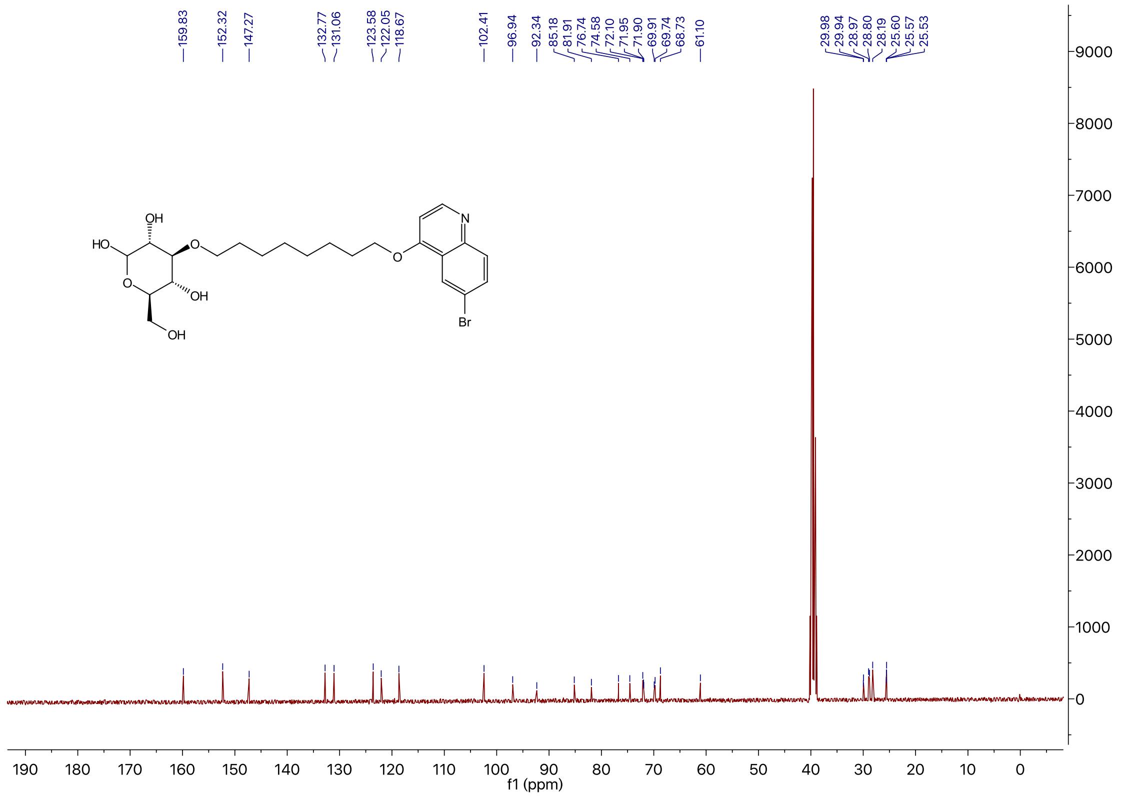

[1]H NMR of Compound **4e** in DMSO-$d_6$

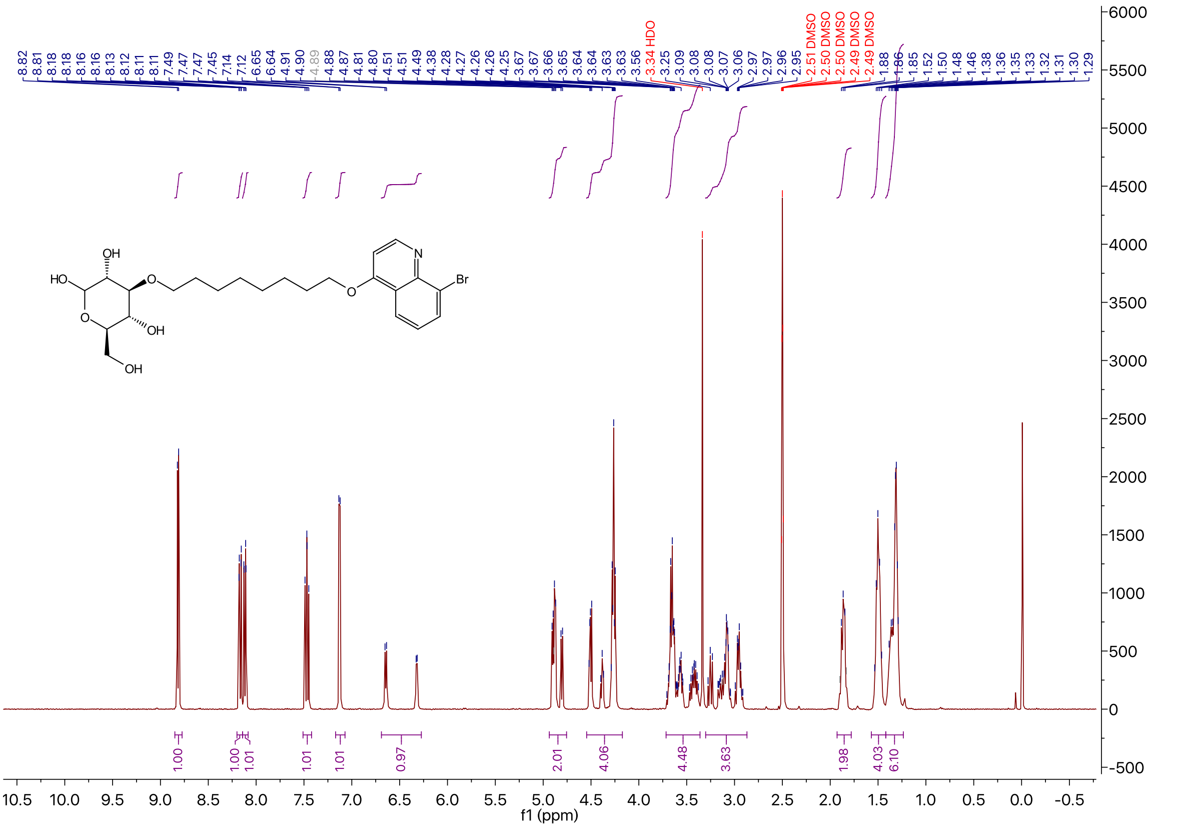

[13]C NMR of Compound **4e** in DMSO-$d_6$

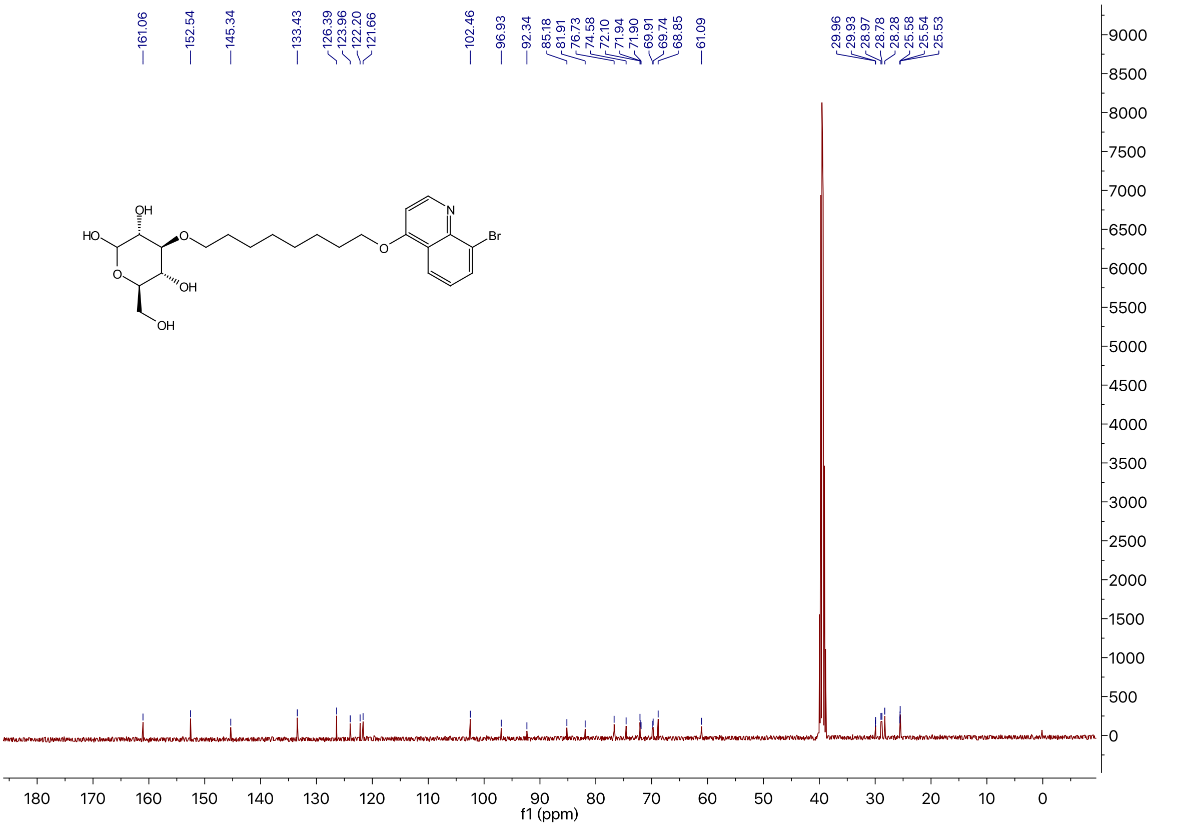

$^1$H NMR of Compound **4f** in DMSO-$d_6$

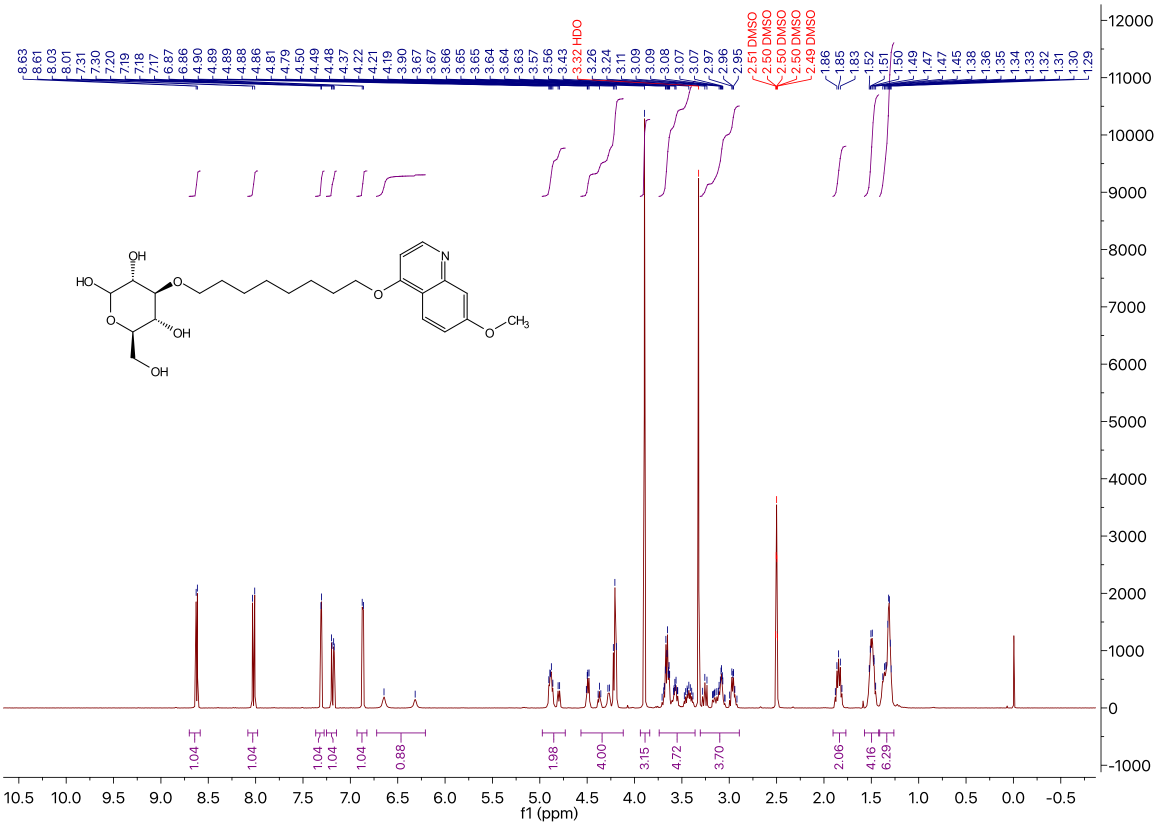

$^{13}$C NMR of Compound **4f** in DMSO-$d_6$

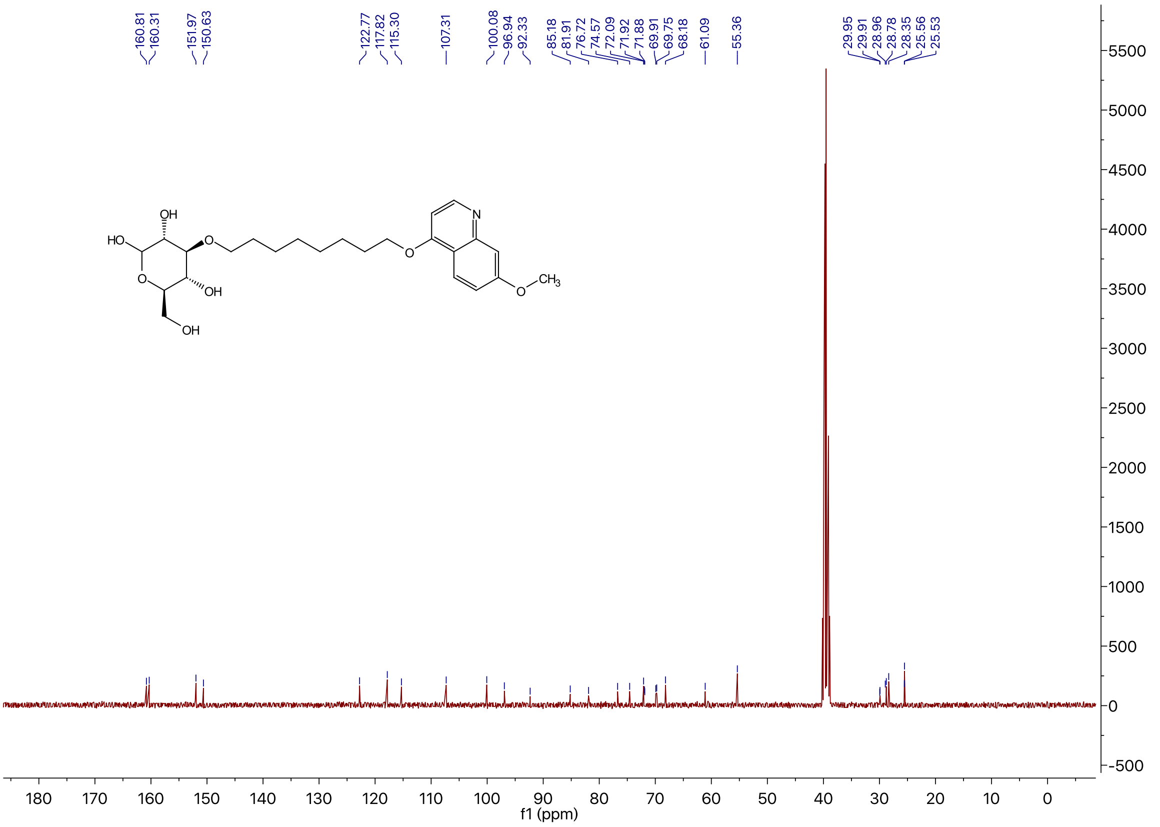

$^{1}$H NMR of Compound **4g** in DMSO-$d_6$

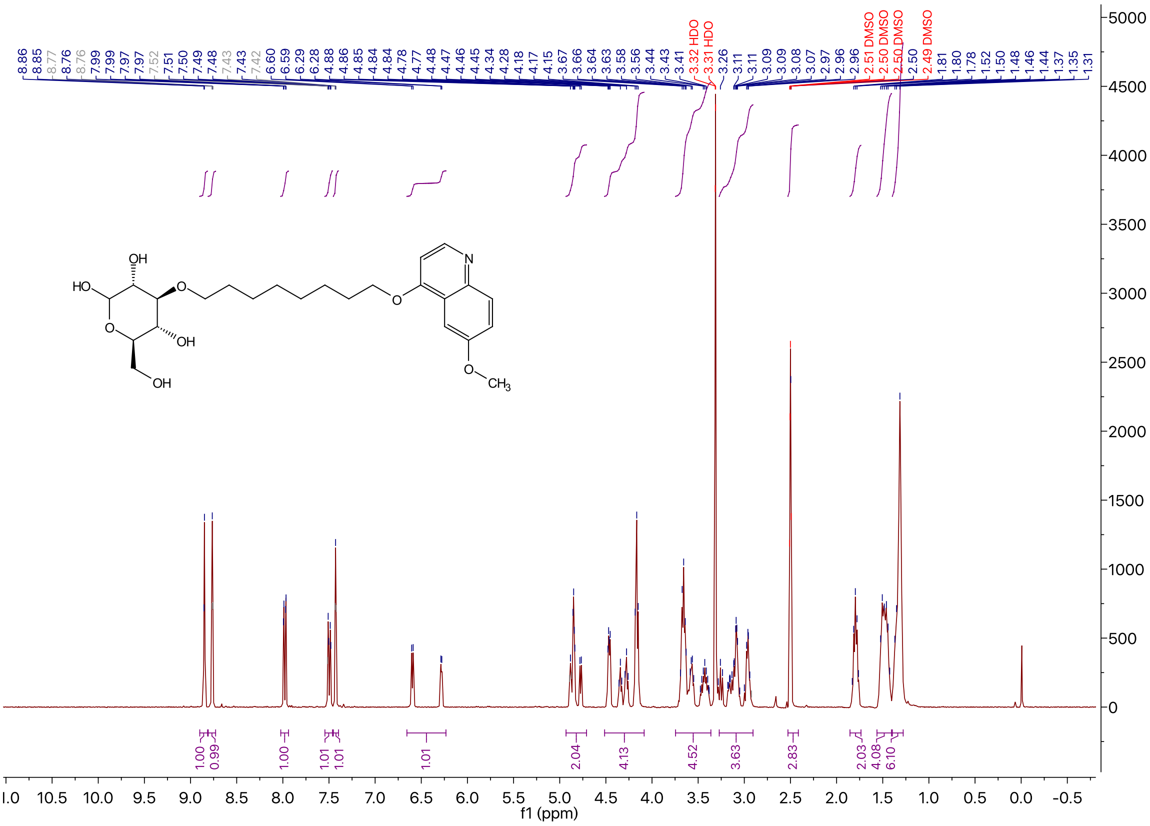

$^{13}$C NMR of Compound **4g** in DMSO-$d_6$

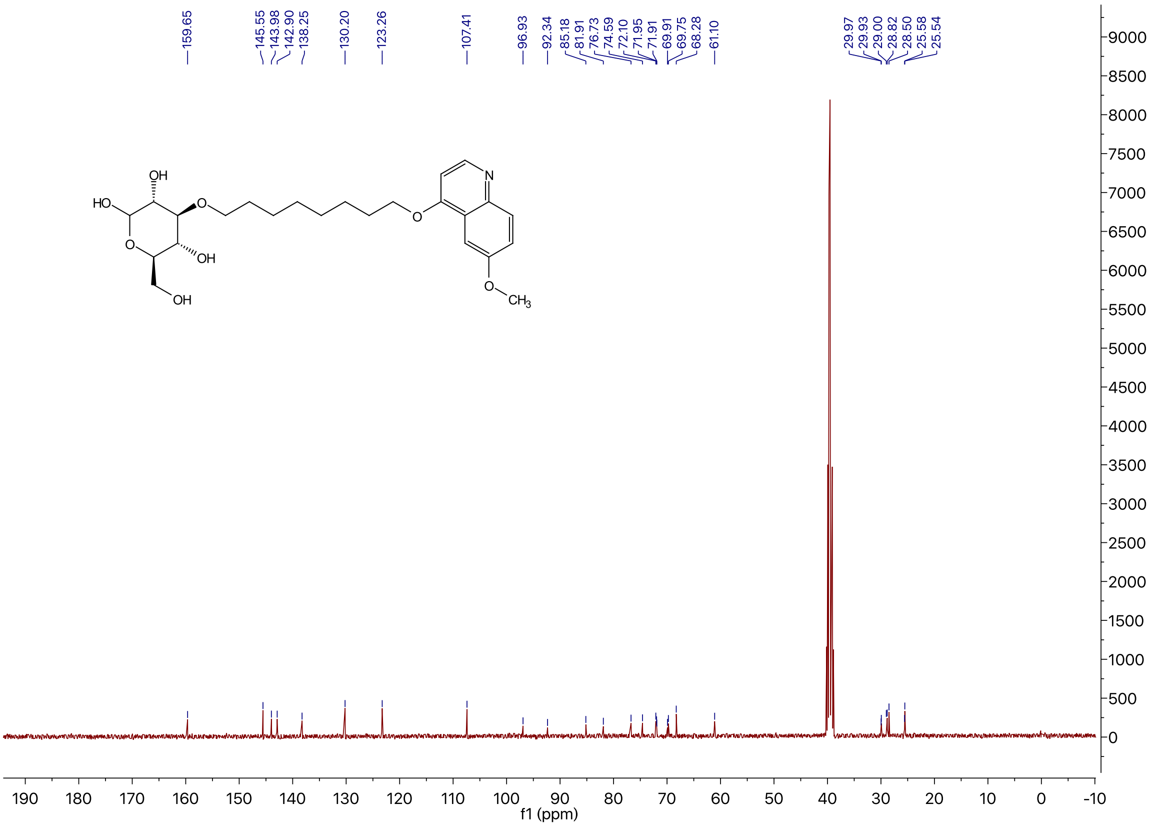

### $^{1}$H NMR of Compound **4h** in DMSO-$d_6$

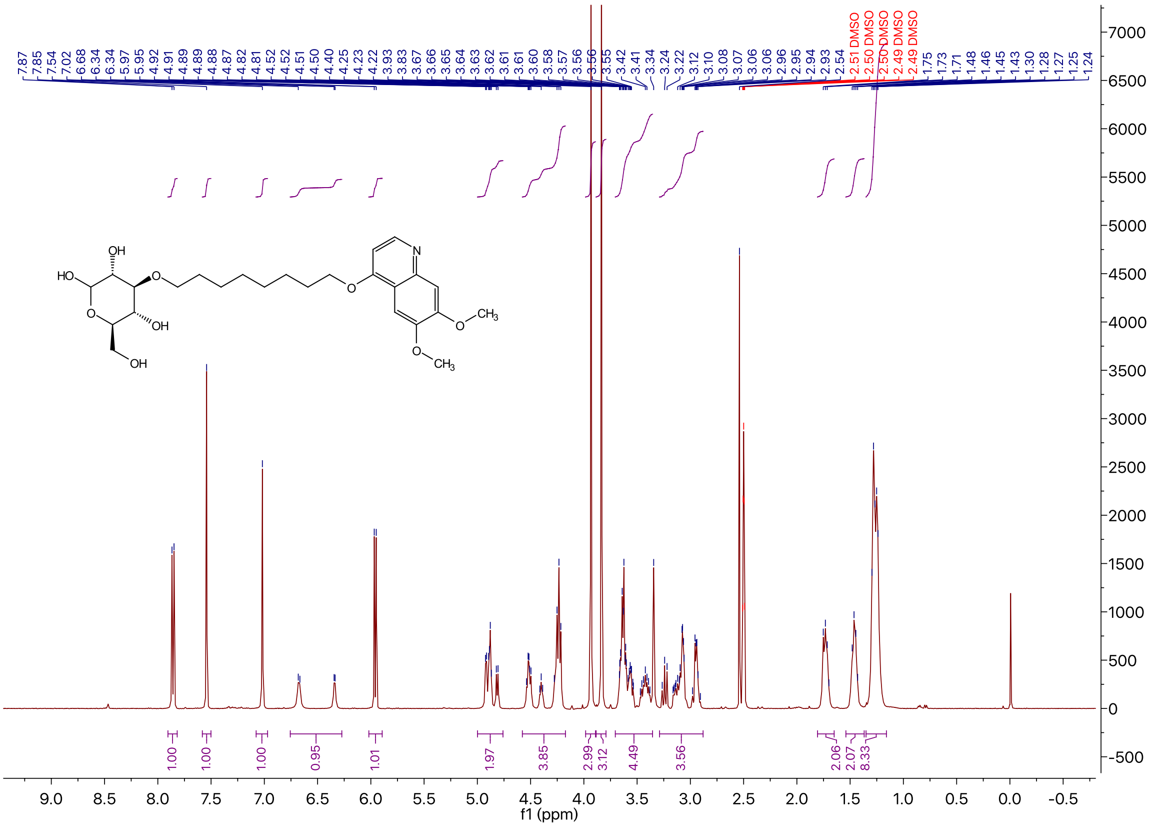

### $^{13}$C NMR of Compound **4h** in DMSO-$d_6$

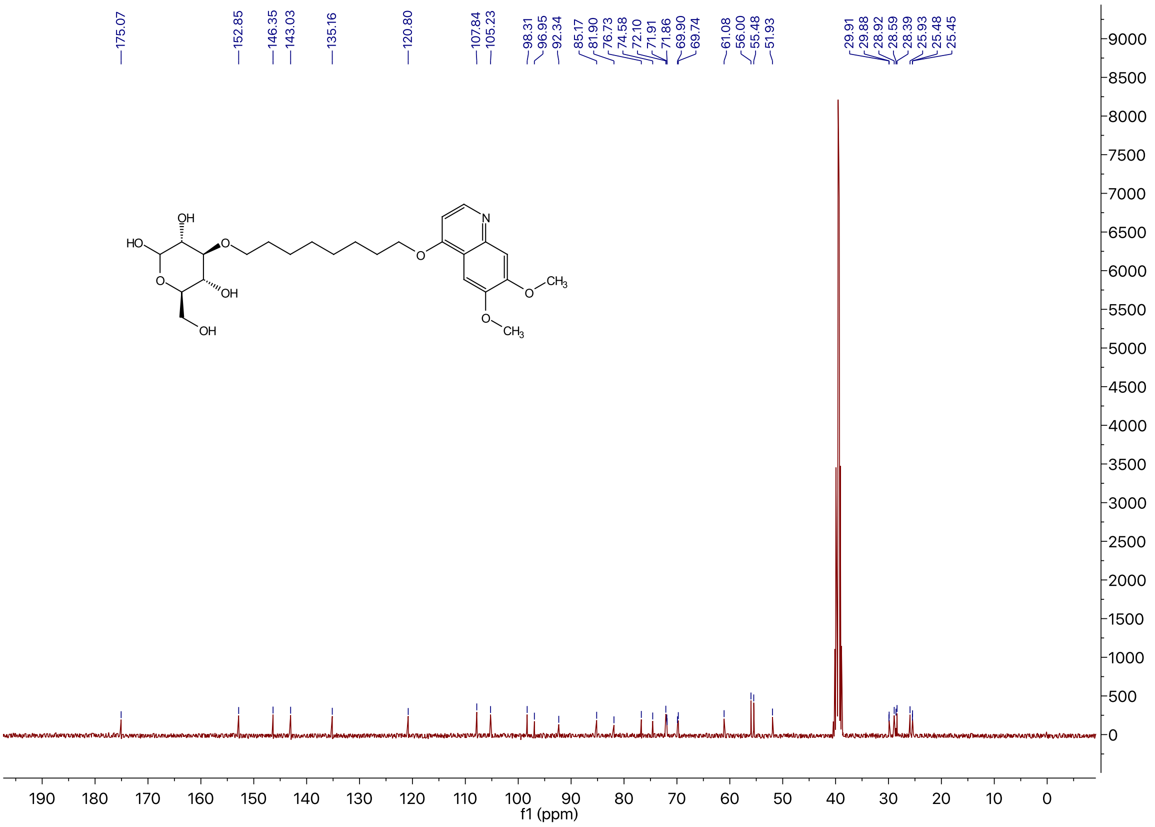

[1]H NMR of Compound **4i** in DMSO-$d_6$

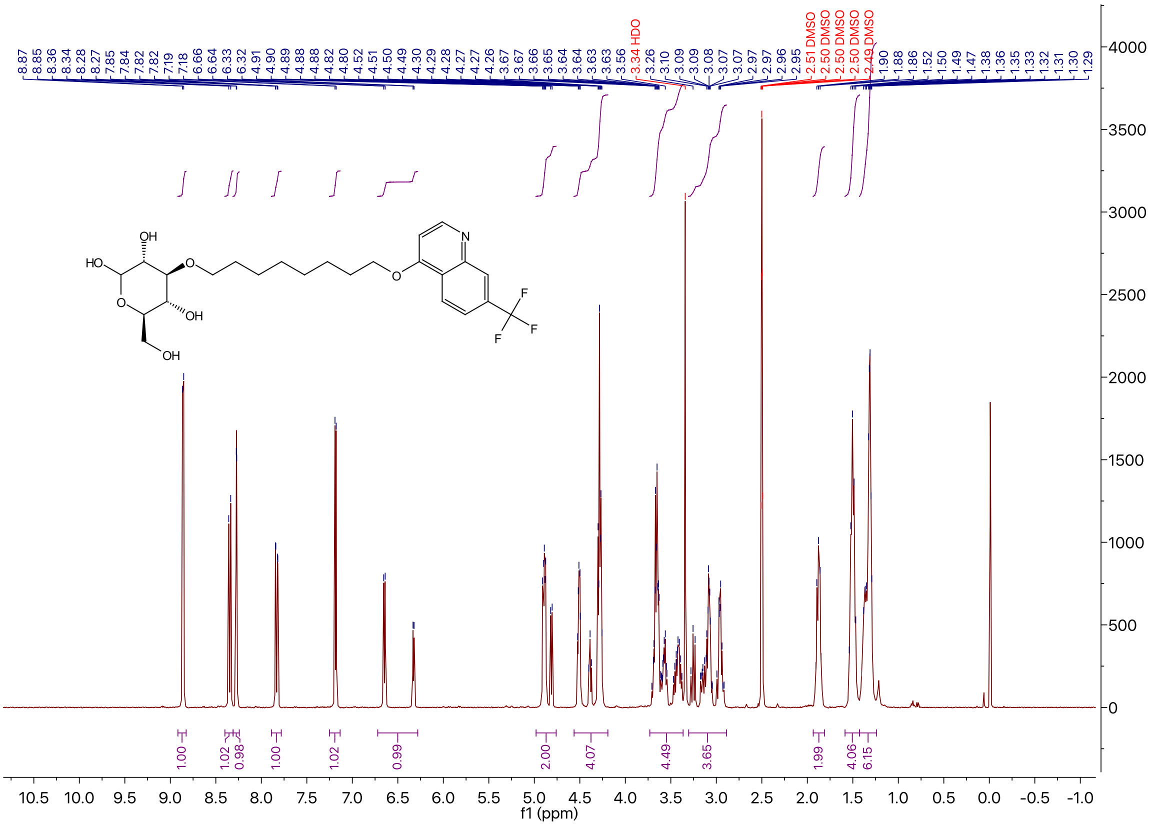

[13]C NMR of Compound **4i** in DMSO-$d_6$

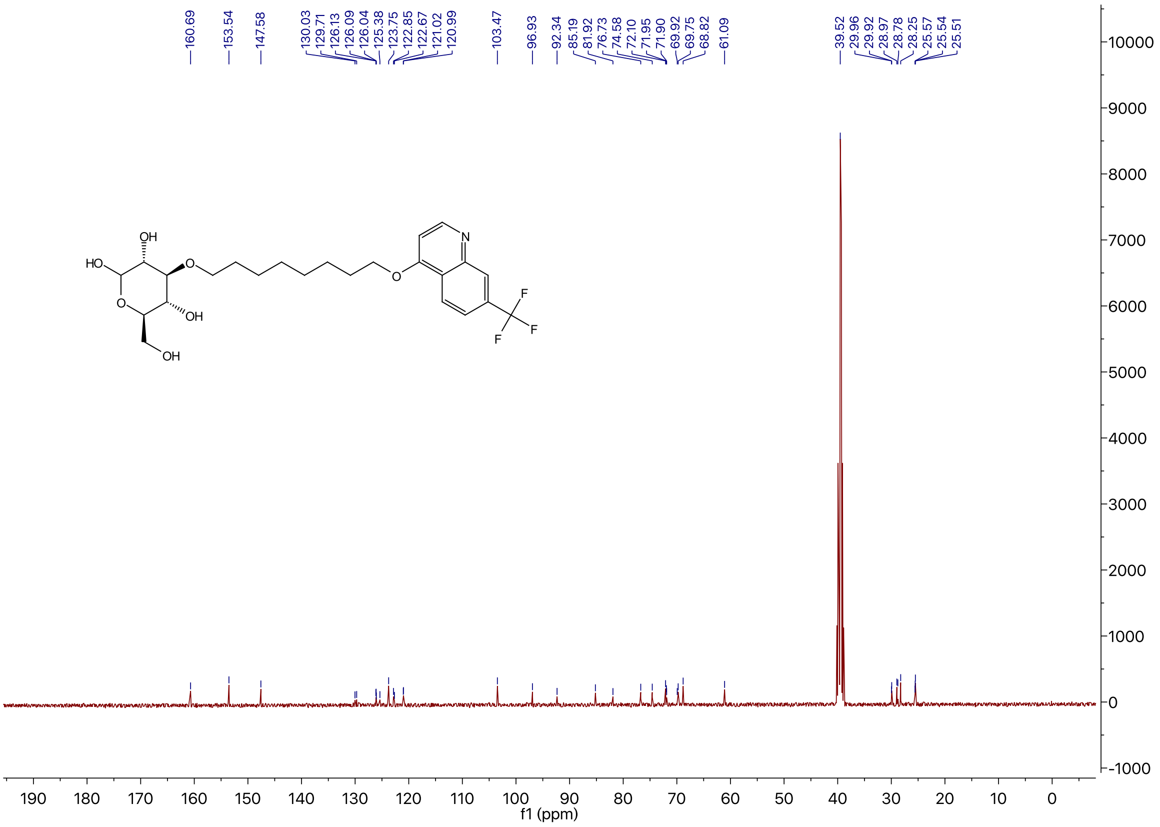

$^{1}H$ NMR of Compound **4j** in DMSO-$d_6$

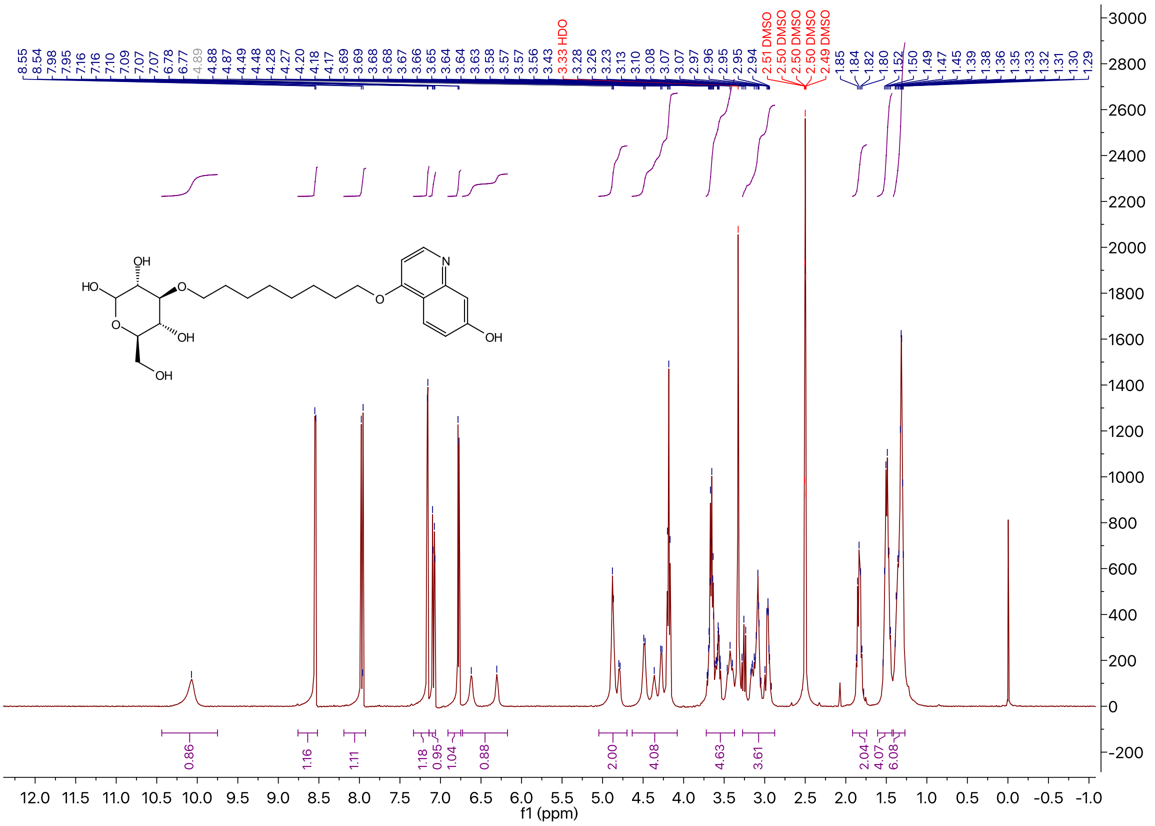

$^{13}C$ NMR of Compound **4j** in DMSO-$d_6$

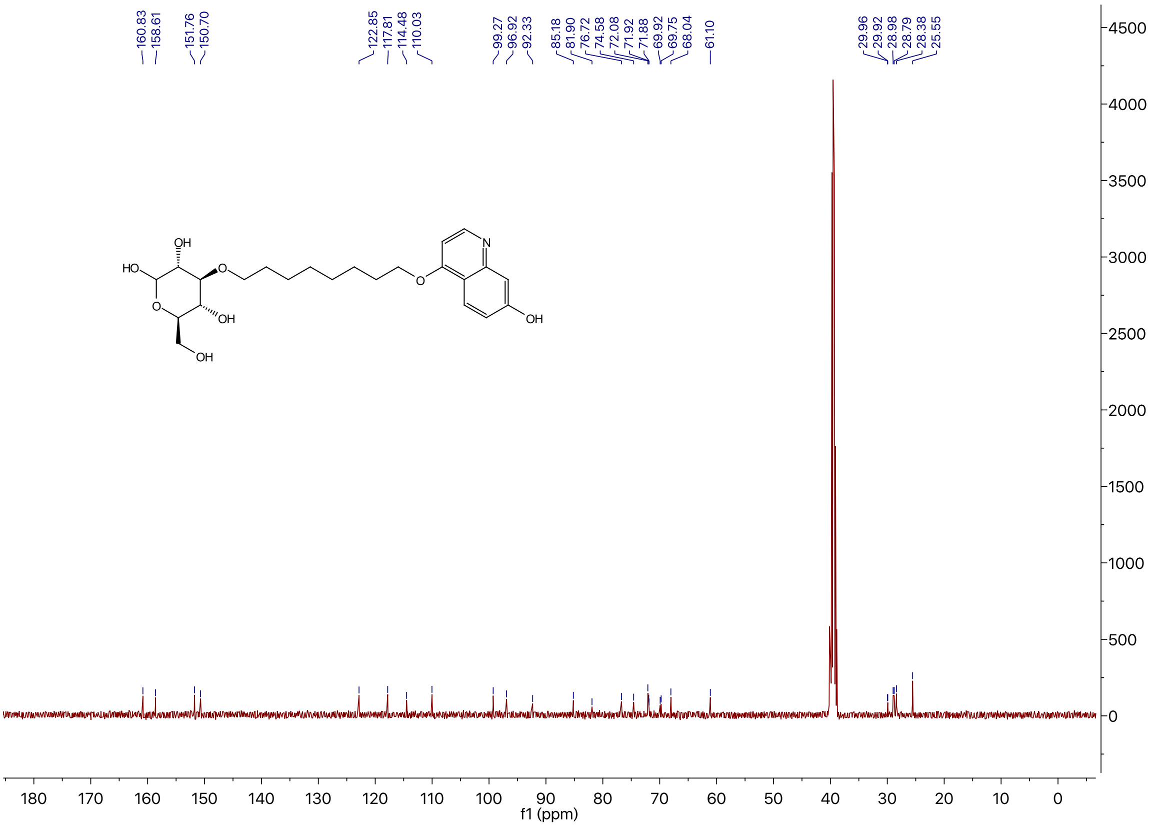

$^{1}H$ NMR of Compound **15** in $CDCl_3$

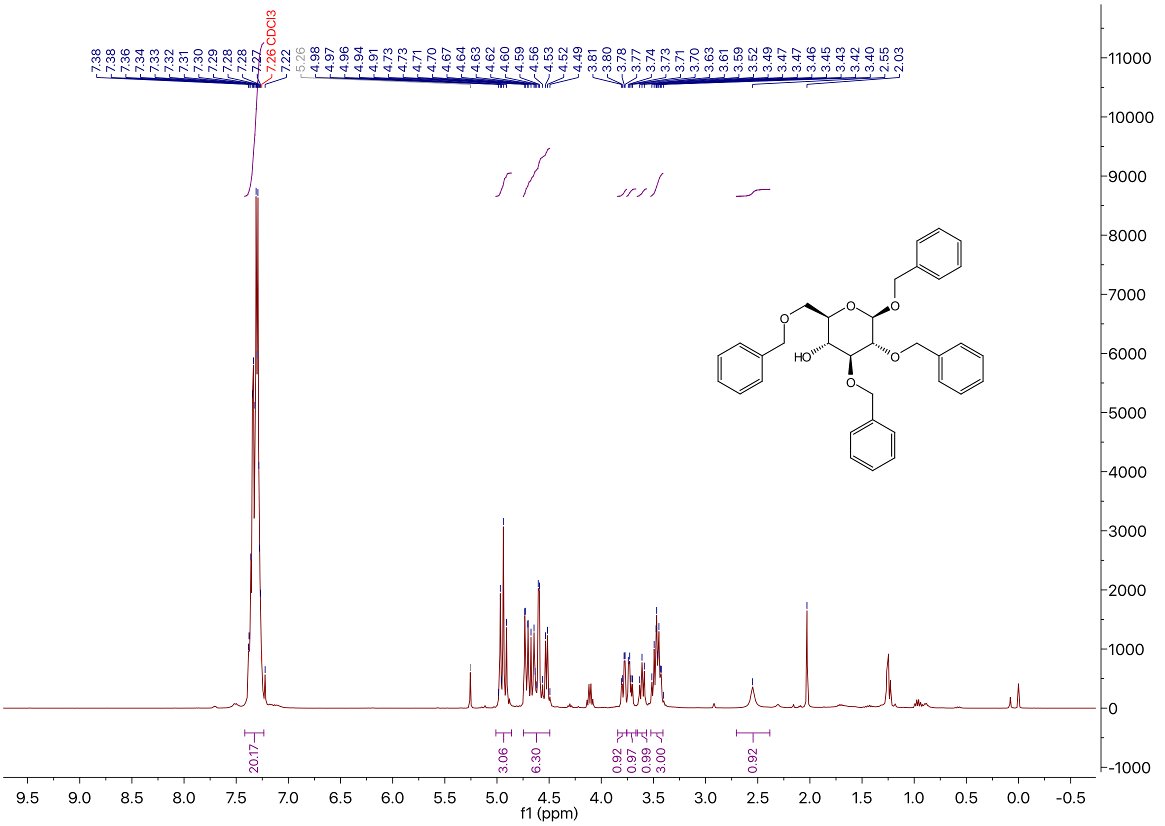

$^{13}C$ NMR of Compound **15** in $CDCl_3$

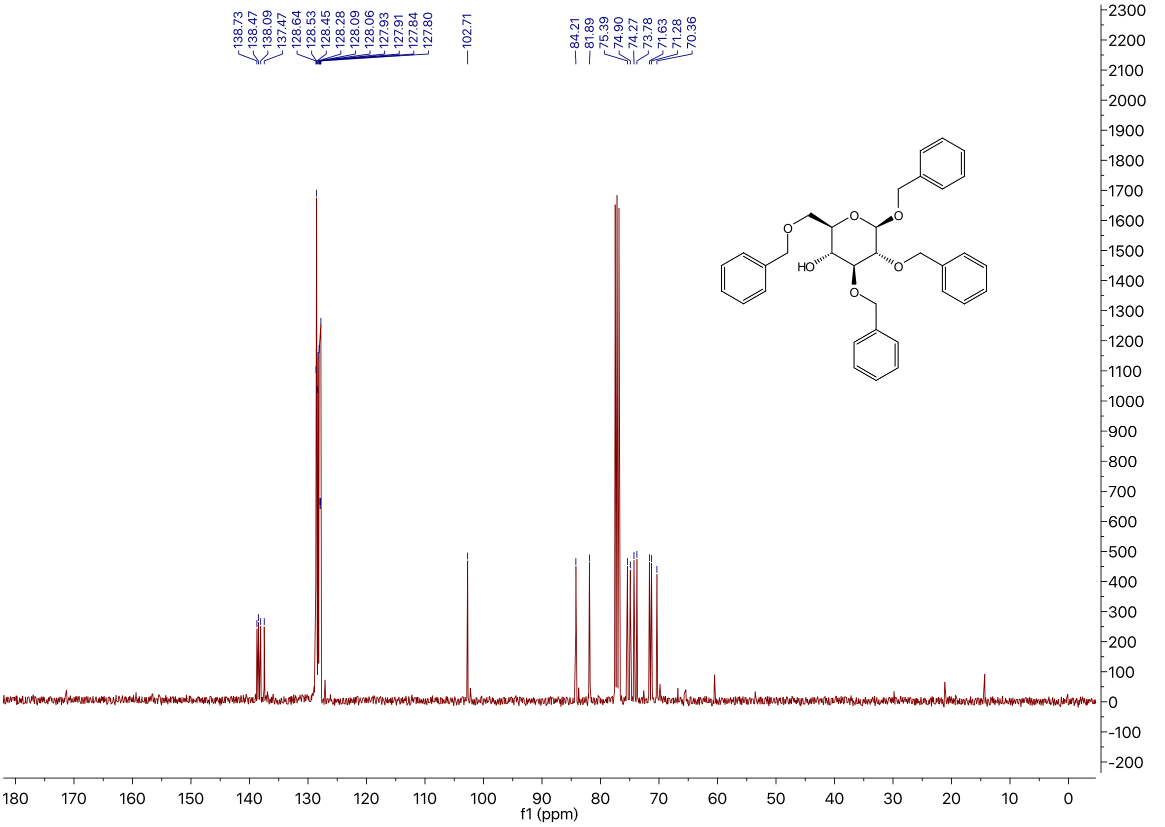

### $^{1}$H NMR of Compound **17** in DMSO-$d_6$

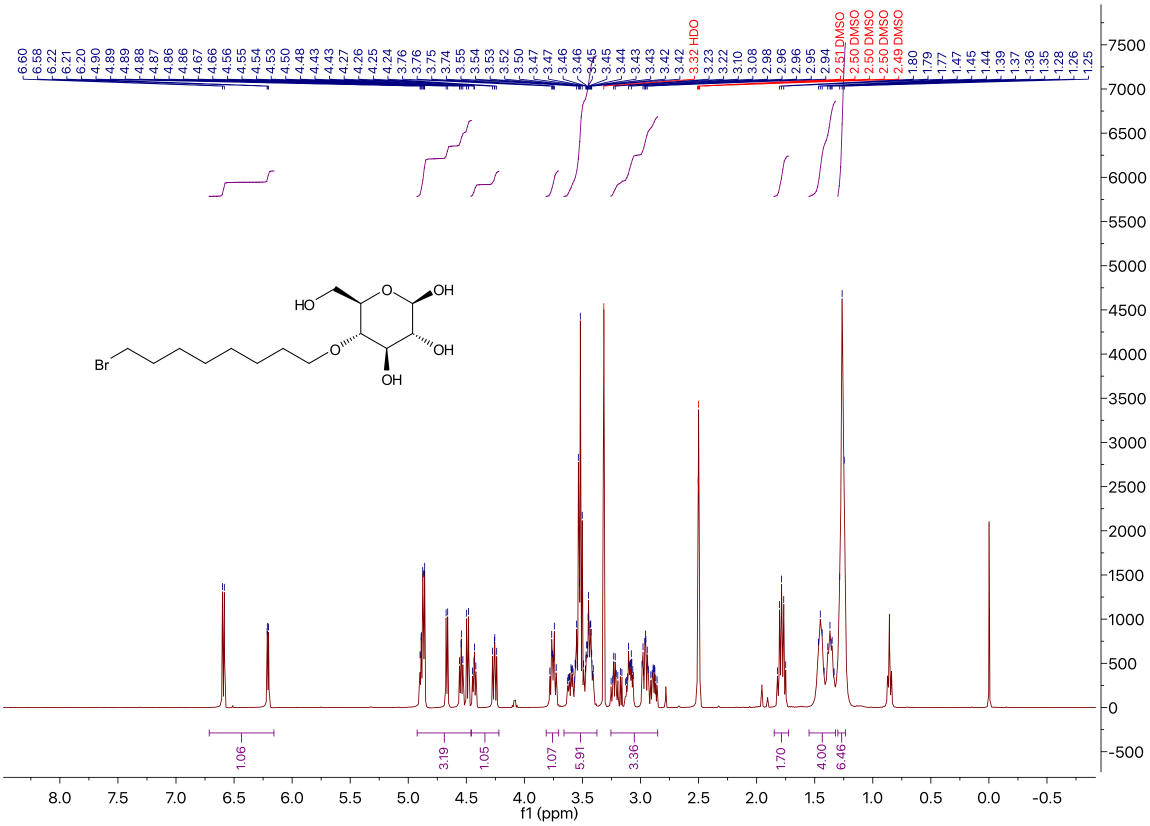

### $^{13}$C NMR of Compound **17** in DMSO-$d_6$

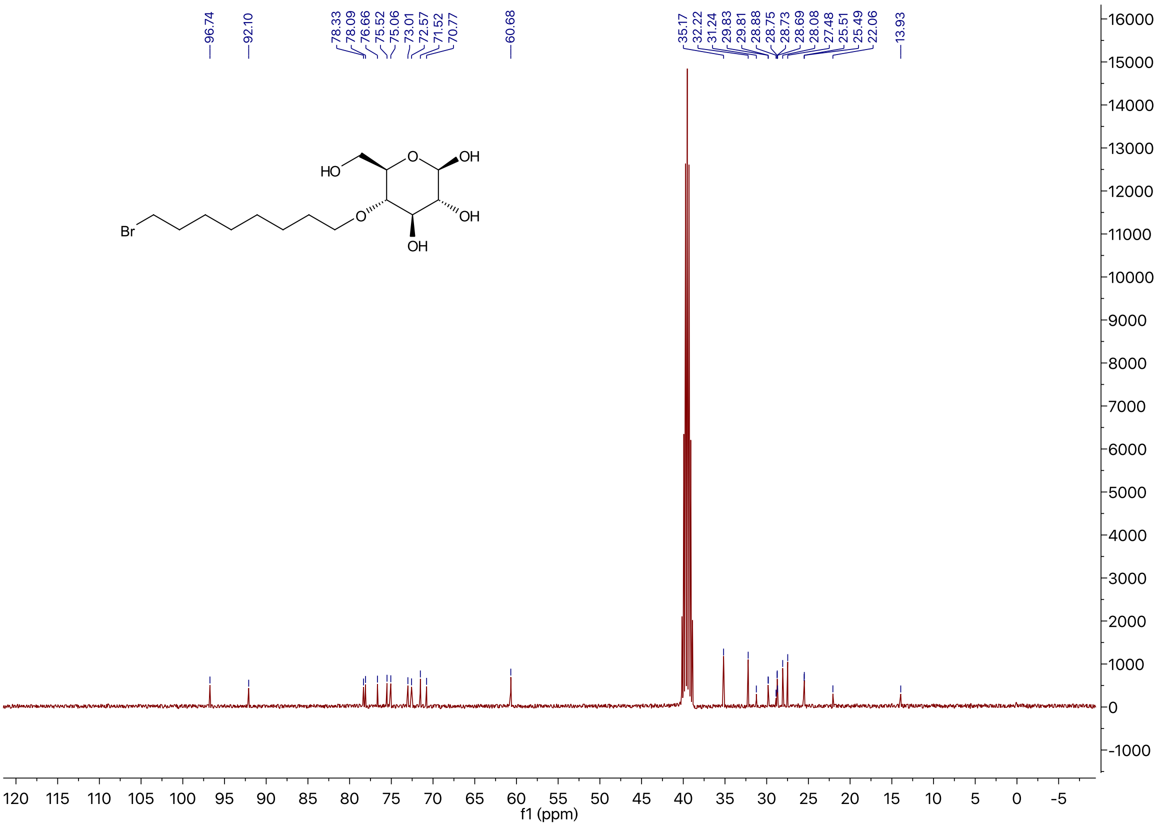

$^1$H NMR of Compound **5c** in DMSO-$d_6$

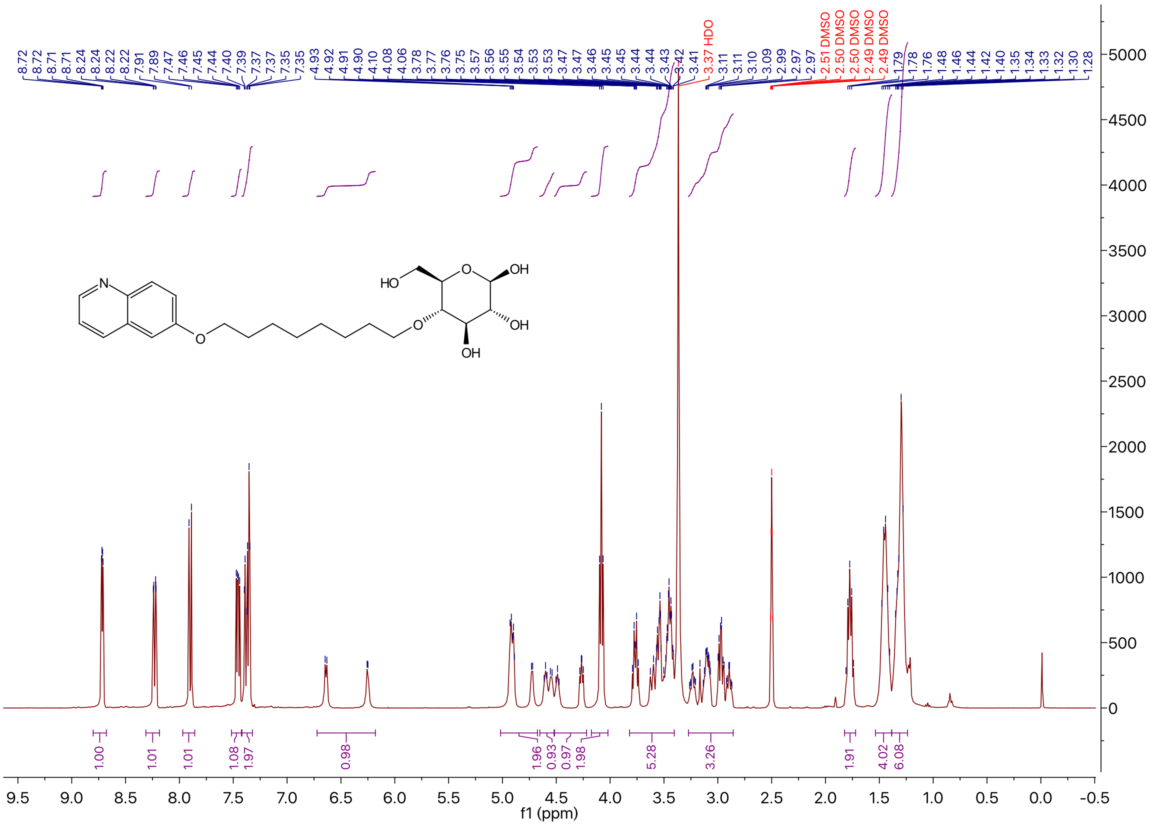

$^{13}$C NMR of Compound **5c** in DMSO-$d_6$

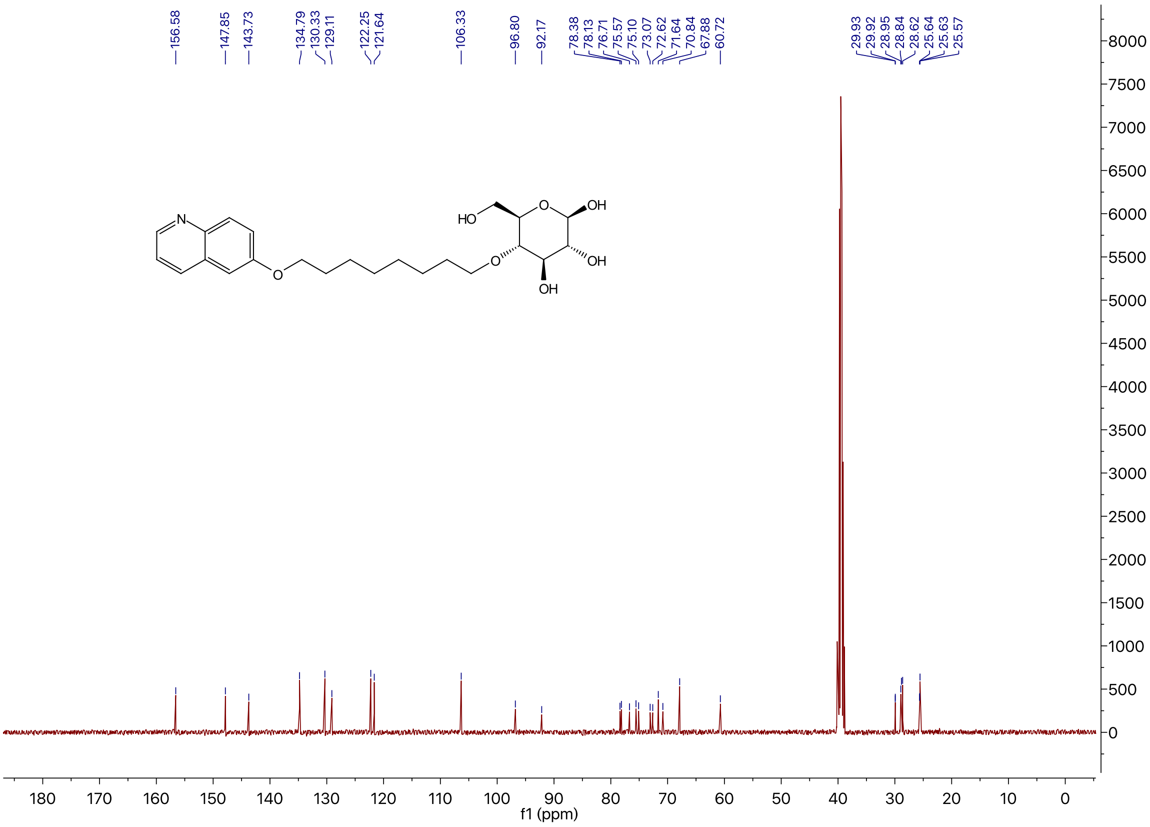

$^1$H NMR of Compound **5a** in Methanol-$d_4$

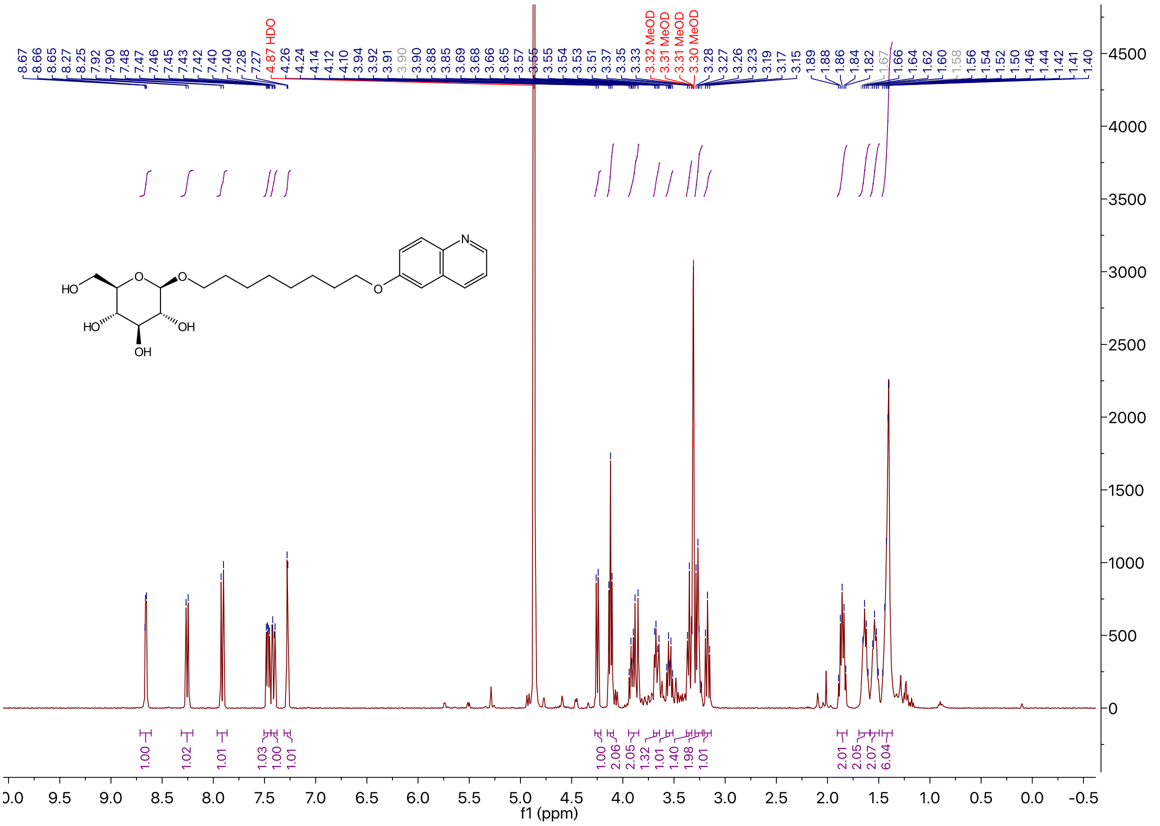

$^{13}$C NMR of Compound **5a** in Methanol-$d_4$

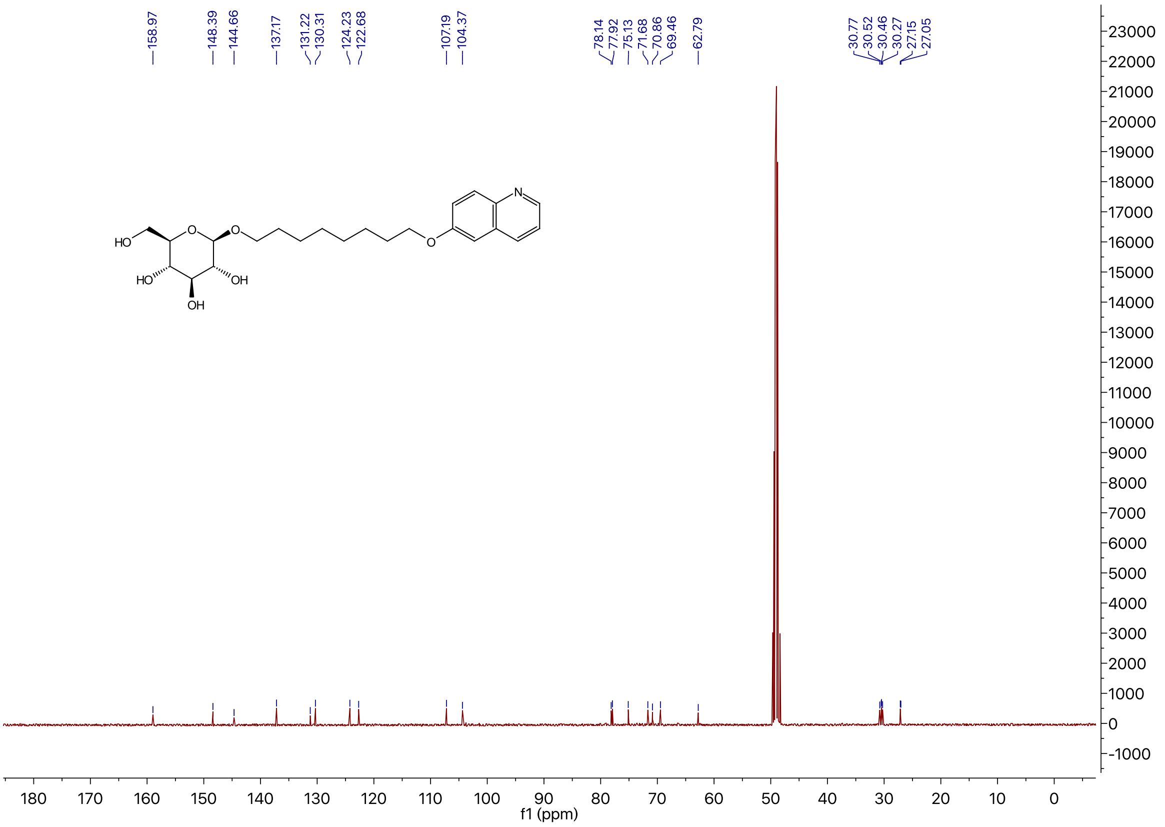

### [1]H NMR of Compound **18** in $CDCl_3$

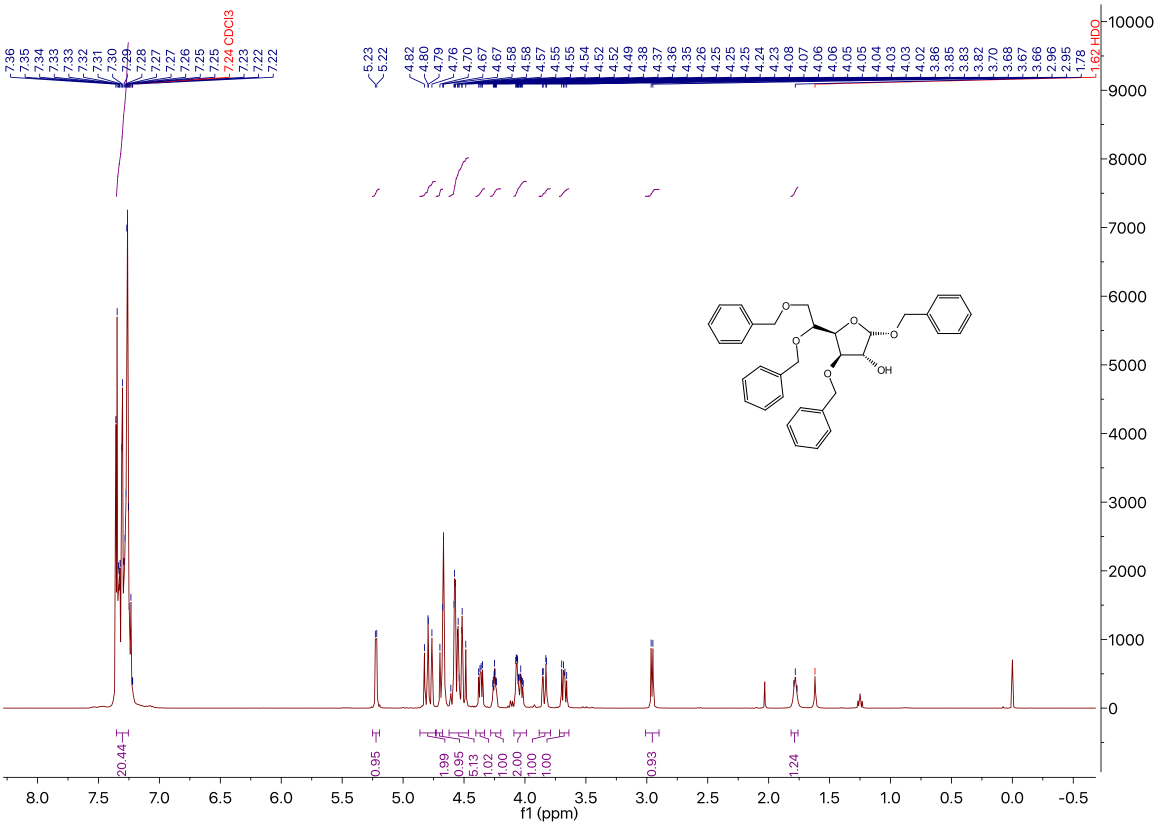

### [13]C NMR of Compound **18** in $CDCl_3$

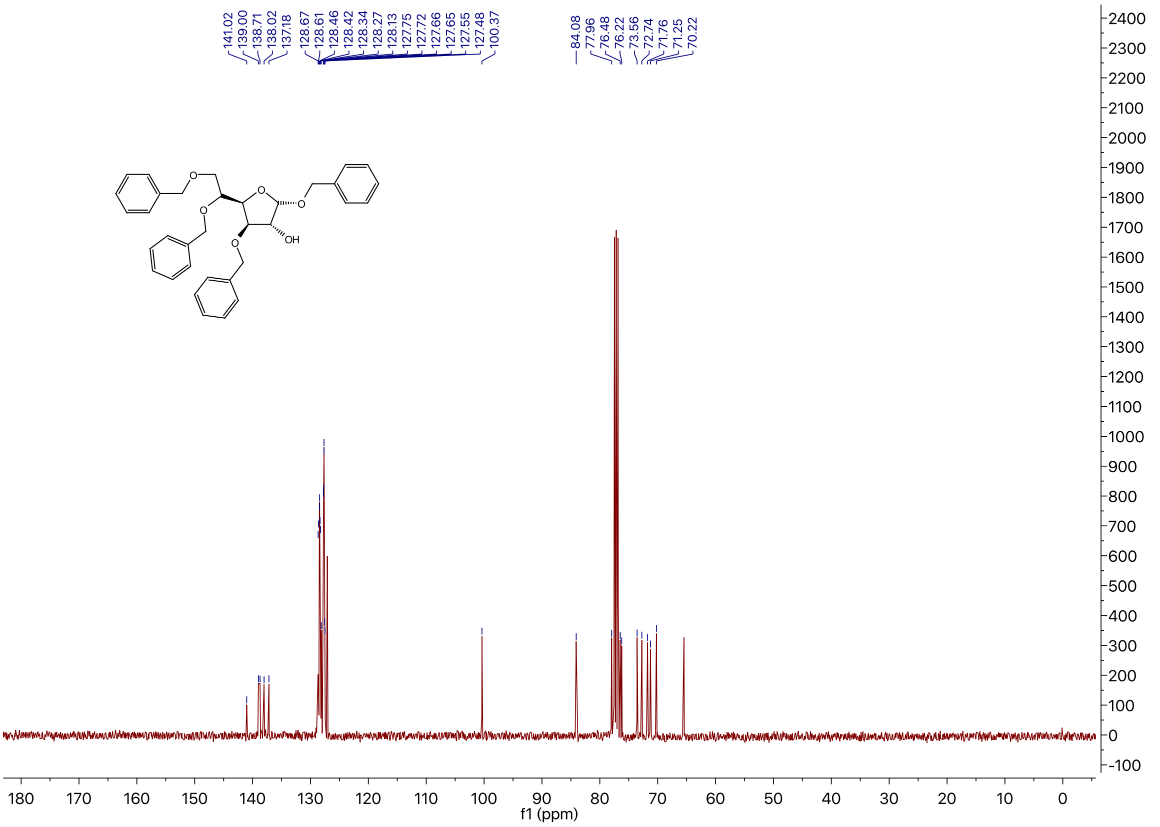

[1]H NMR of Compound **5b** in DMSO-$d_6$

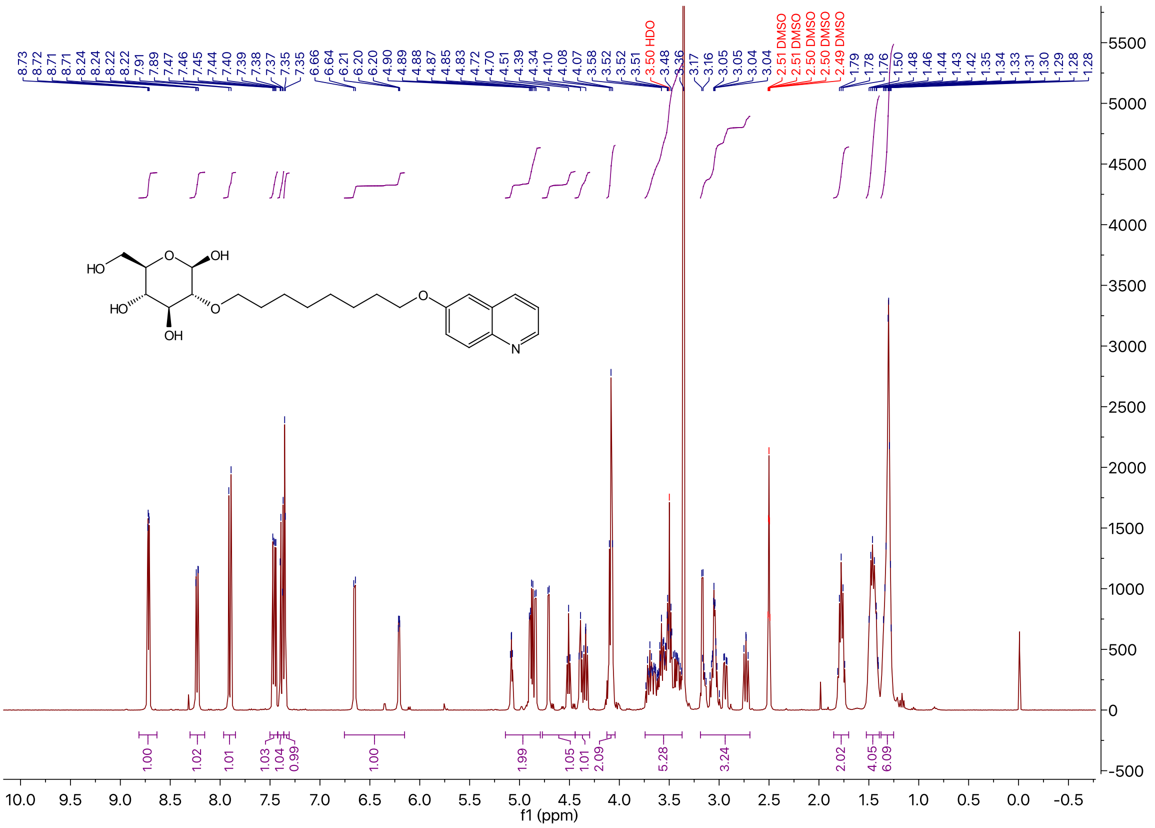

[13]C NMR of Compound **5b** in DMSO-$d_6$

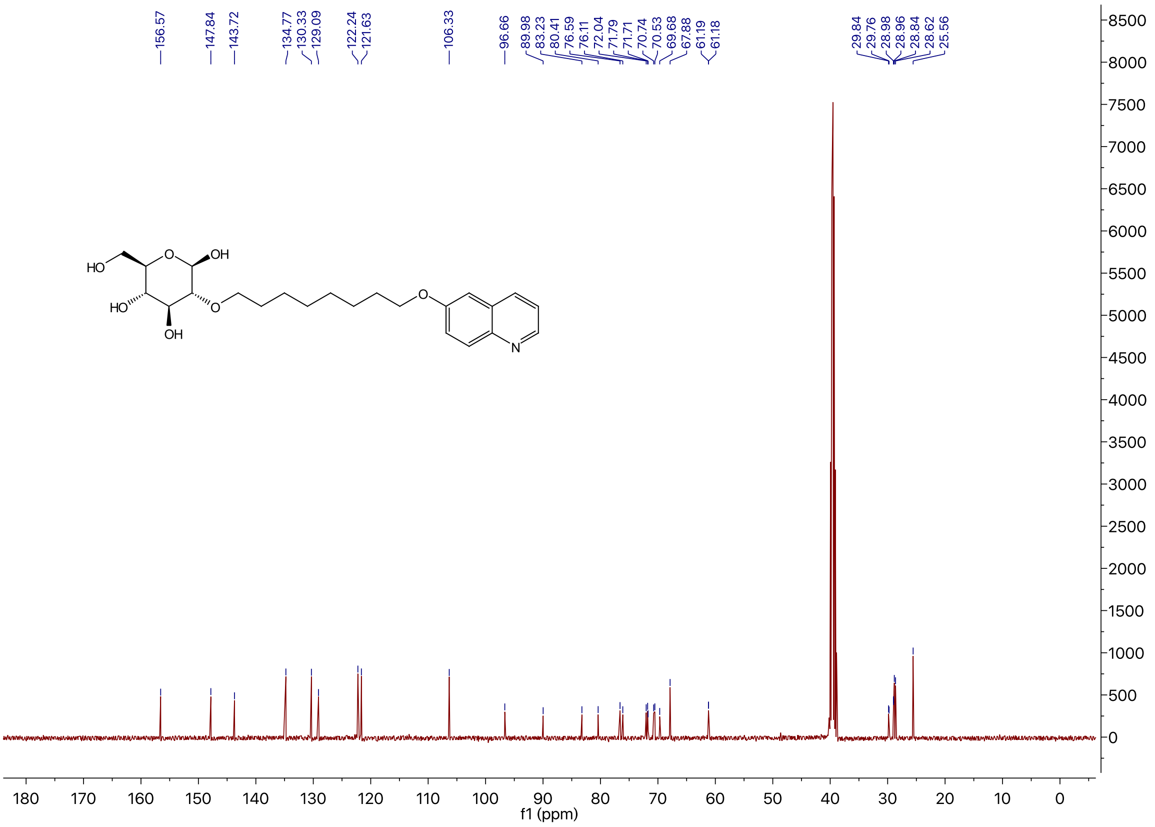

$^1$H NMR of Compound **5d** in DMSO-$d_6$

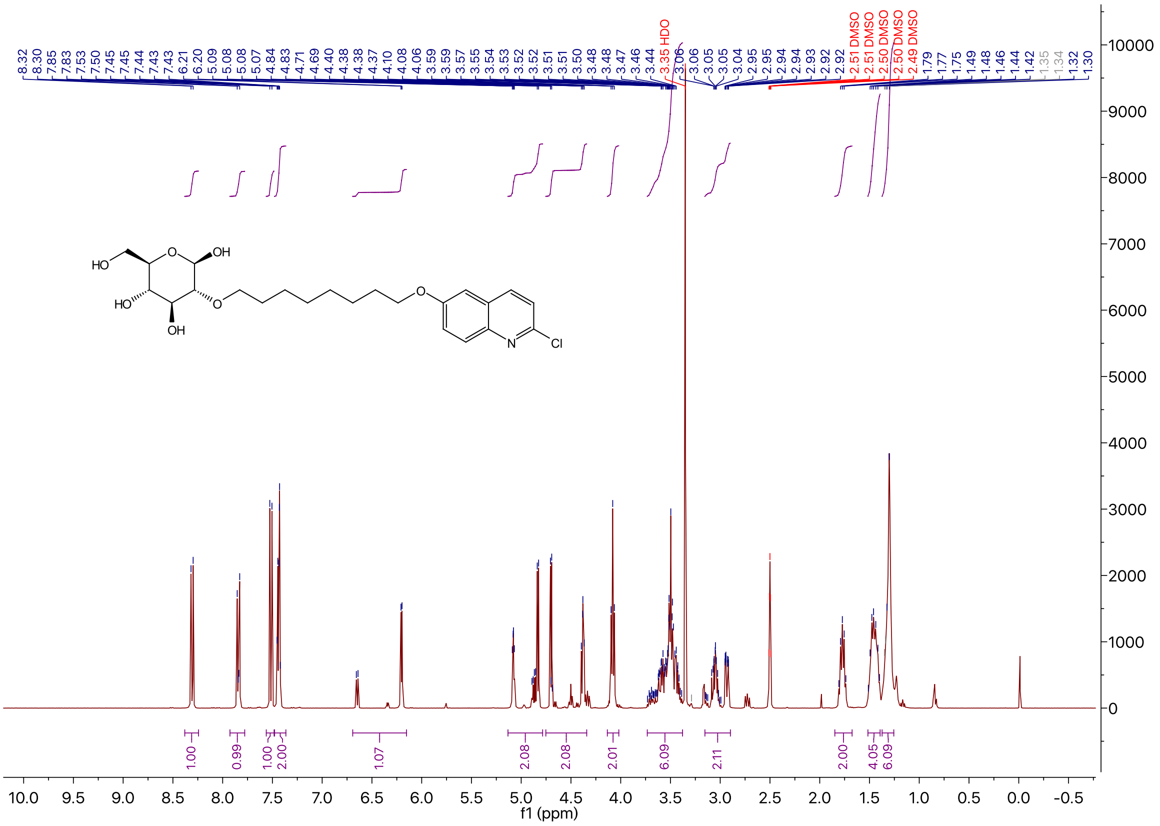

$^{13}$C NMR of Compound **5d** in DMSO-$d_6$

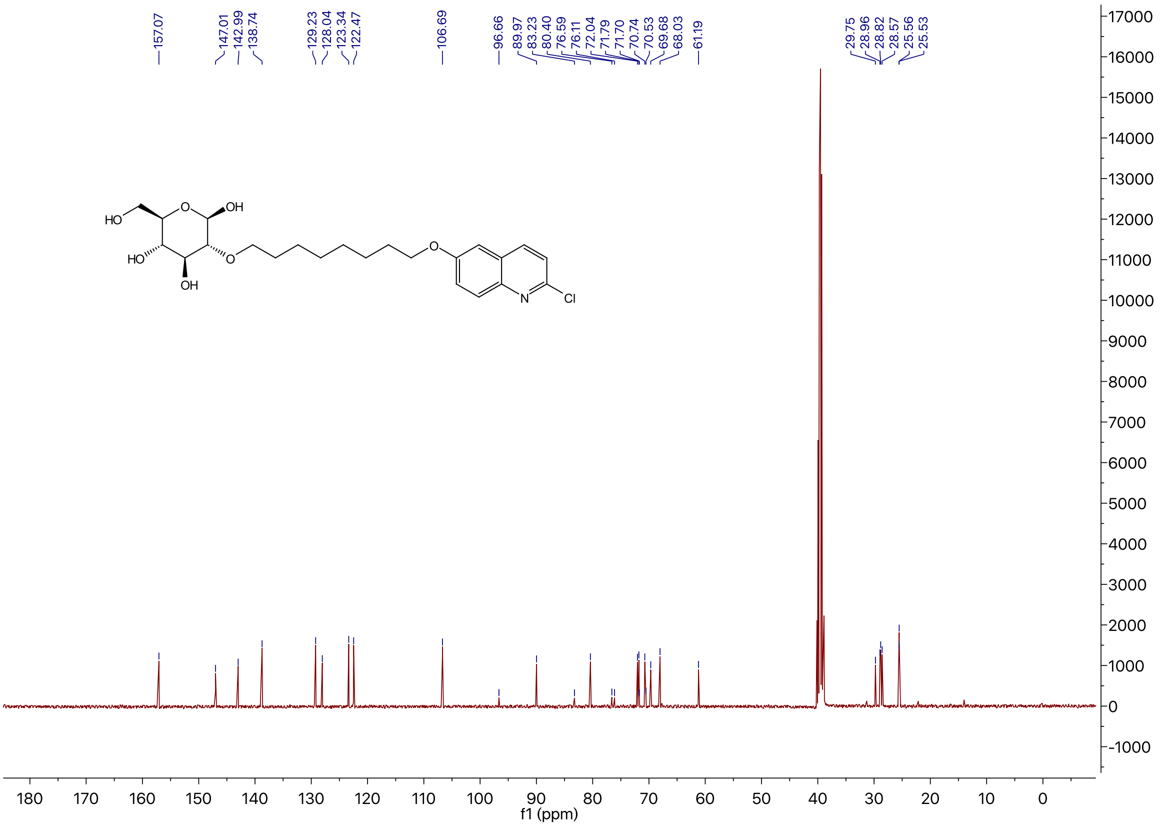

[1]H NMR of Compound **5e** in DMSO-$d_6$

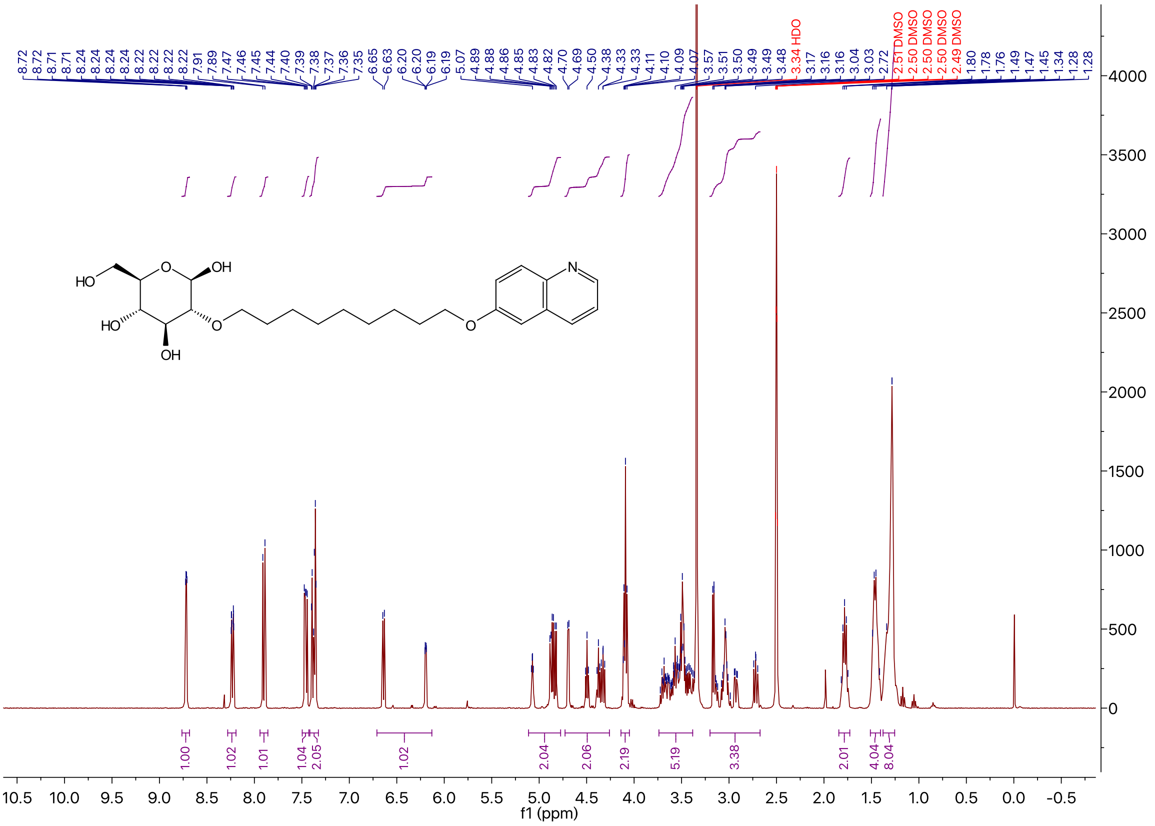

[13]C NMR of Compound **5e** in DMSO-$d_6$

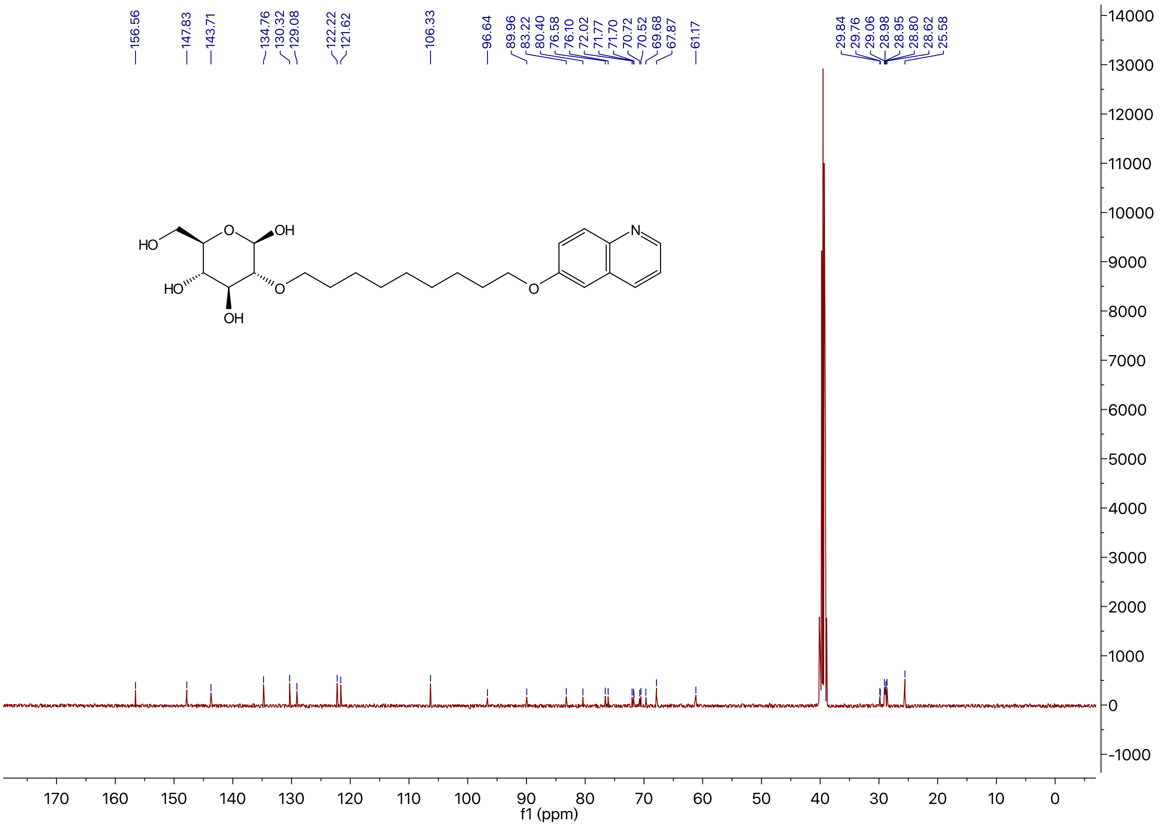
